# Cell surface sculpting using logic-gated protein actuators

**DOI:** 10.1101/2023.12.18.572113

**Authors:** Christian Kofoed, Nicholas E. S. Tay, Xuanjia Ye, Girum Erkalo, Tom W. Muir

## Abstract

Cell differentiation and tissue specialization lead to unique cellular surface landscapes and exacerbated or loss of expression patterns can result in further heterogenicity distinctive of pathological phenotypes^1–3^. Immunotherapies and emerging protein therapeutics seek to exploit such differences by engaging cell populations selectively based on their surface markers. Since a single surface antigen rarely defines a specific cell type^4,5^, the development of programmable molecular systems that integrate multiple cell surface features to convert on-target inputs to user-defined outputs is highly desirable. Here, we describe an autonomous decision-making protein device driven by proximity-gated protein *trans*-splicing that allows local generation of an active protein from two otherwise inactive fragments. We show that this protein actuator platform can perform various Boolean logic operations on cell surfaces, allowing highly selective recruitment of enzymatic and cytotoxic activities to specific cells within mixed populations. Due to its intrinsic modularity and tunability, this technology is expected to be compatible with different types of inputs, targeting modalities and functional outputs, and as such will have broad application in the synthetic biology and biotechnology areas.

## Introduction

Exploiting the heterogeneity of cell surfaces is central to many diagnostic tools and therapeutic strategies. Typically, these approaches employ a targeting vector such as an antibody that engages a cell surface marker to deliver a pharmacological, enzymatic, or cellular activity^6–9^. While this general approach has notable successes, it must contend with the fact that single antigens rarely define cell or tissue types and are often not disease specific. Wrongful commitments of these agents can therefore occur, and in the case of therapeutic modalities may lead to unwanted side-effects^4,5^. One possible solution to this problem is to employ targeting vectors that recognize two or more surface antigens through Boolean logic^10,11^. In line with this, bispecific antibodies partition cell engagement into two concurrent low-affinity binding events^12^; however, achieving complete subpopulation discrimination can be difficult to fine-tune^13^ and the type of logic operations possible with these systems is limited. Recently, *de novo* protein switches and engineered antibody platforms have been reported that couple functional outputs to multiple binding events through Boolean logic gates^14–17^. Though promising, current versions of these tools remain limited in the type of output functions possible. Absolute cell discrimination in terms of binding and activity therefore remains an outstanding challenge with a need for smart protein devices that can perform autonomous decision-making to convert on-target inputs to user-defined outputs in a spatially and temporally controlled manner.

Protein *trans*-splicing (PTS) is intrinsically AND gated: Split intein fragments (IntN and IntC) first undergo complementation followed by an autocatalytic excision-ligation reaction that reconstitutes a protein of interest (POI) in a traceless manner^18^. However, many naturally split inteins conceal their AND gated nature by assembly with pico-to-nanomolar dissociation constants, essentially making them indistinguishable from single chain variants with pseudo first-order reaction kinetics^19^. Recently, we developed caged split inteins where each fragment is fused to truncated versions of the matching partner^20^. This dramatically inhibits fragment complexation of the otherwise ultrafast and ultraefficient naturally split inteins Npu, Gp41-1, GP41-8, and NrdJ-1 and alters their trajectories to follow second-order reaction kinetics. Triggered colocalization can, however, induce spontaneous intermolecular domain swapping, which generates the functional intein fold^21^. Any fragmented POI produced through this scheme is inherently AND gated as it depends on conditional protein splicing (CPS) to recapitulate its function.

Building on advances in CPS, we develop herein the SMART (Splicing-Modulated Actuation upon Recognition of Targets) platform that performs autonomous decision-making based on programmable Boolean logic. Our device relies on two components each with three simple modules: a caged split intein fragment fused to a split-POI and an antigen-binding domain (e.g., nanobodies, DARPin, ligand, etc.). The SMART device passively discriminates among various populations based on their antigen surface presentations and co-engages cells with a correct combinatorial display prompting splicing and POI function (Fig. 1a). Following this outline, we report a generalizable scaffold able to produce a POI based on Boolean AND, OR and NOT logic operations within a heterogeneous cell community at single-cell resolution and without any significant off-target effects.

**Figure 1.**
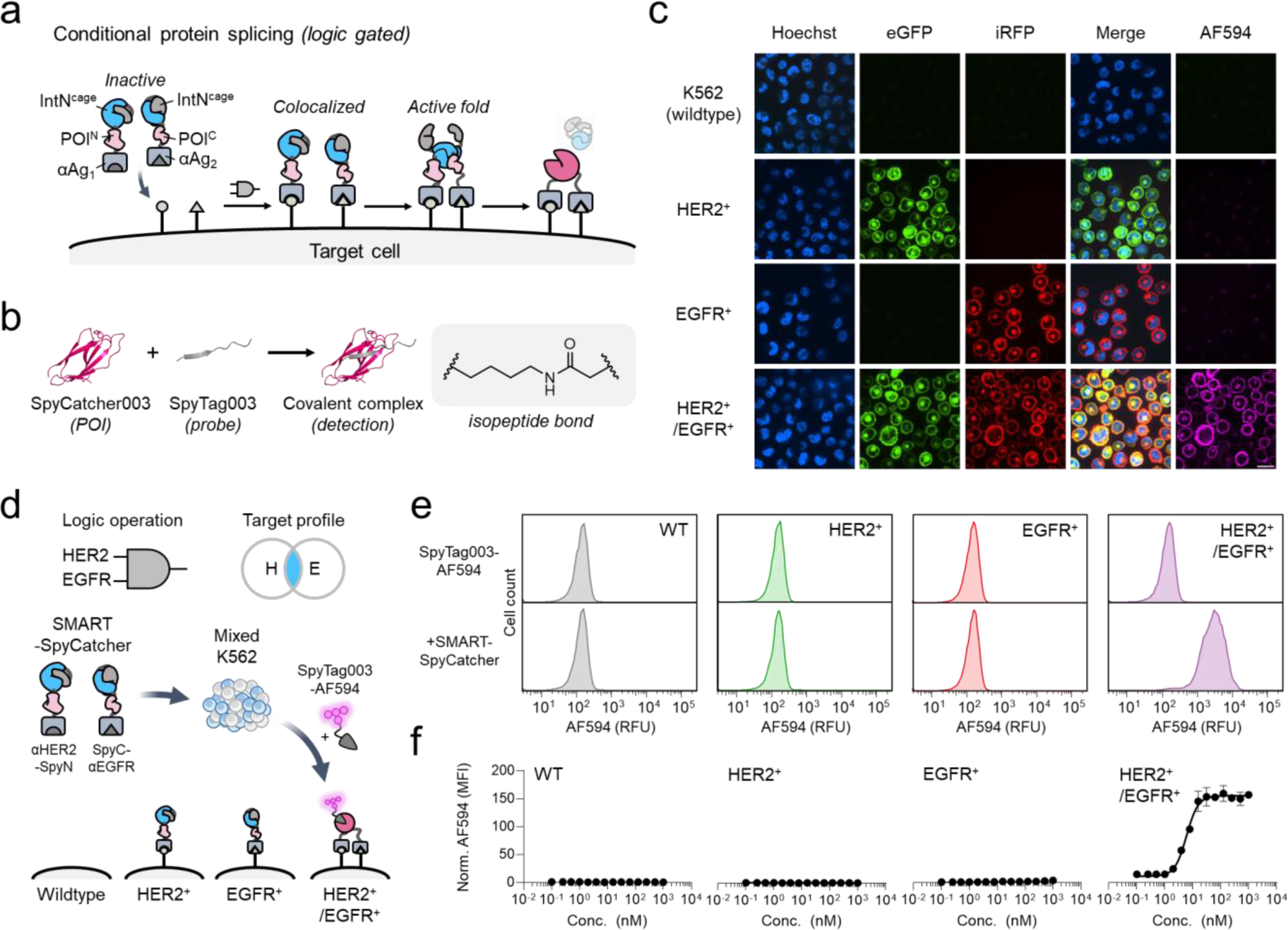
A protein switch that computes Boolean logic on live-cell surfaces using conditional protein splicing. **a**, Schematic of the SMART system. Complementary caged split inteins (IntN^cage^ and IntC^cage^) are fused to targeting vectors (αAg_1_ and αAg_2_) against different cell surface antigens. Colocalization of these proteins on the cell surface leads to protein *trans*-splicing and generation of a functional protein of interest (POI) nested between αAg_1_ and αAg_2_. The system works as an AND-gated input-to-output actuator. **b**, As a functional output SpyCatcher003 was split at position 73-74 and its fragments were fused to IntN^cage^ and IntC^cage^ thereby generating SMART-SpyCatcher. While the split version of SpyCatcher003 is inactive, the reconstituted form can covalently associate with SpyTag003 through the formation of an isopeptide bond. **c**, Individual K562 (wildtype), K562^HER2+^, K562^EGFR+^, and K562^HER2+/EGFR+^ cell lines were treated with αHER2-SpyN (100nM) and SpyC-αEGFR (100 nM) for 2 hr, followed by SpyTag003 labeled with Alexa Fluor 594 (SpyTag003-AF594, 100 nM) for 20 mins. Following washing, the live cells were then analyzed by confocal microscopy. Cell nuclei are stained with Hoechst, while HER2 and EGFR are tagged with eGFP and iRFP, respectively. Scale bar equals 20 μm. **d**, Schematic illustrating use of the SMART-SpyCatcher system operating through [HER2 AND EGFR] logic on a mixed population of the four K562 cell lines combined at equal amounts. **e**, Flow cytometry analysis of the mixed K562 population either treated with SpyTag003-AF594 alone (top) or following treatment with SMART-SpyCatcher [HER2 AND EGFR] system (bottom). The four cell types were gated based on HER2 AND EGFR levels and for each cell type the number of cells versus AF594 signal plotted. RFU, relative fluorescence units. **f**, Dose-response experiment on the mixed K562 population with a dilution series of SMART-SpyCatcher (1 µM – 1 nM) and excess SpyTag003-AF594 (100 nM). The fold change in median fluorescence intensity (MFI) of the AF594 flow cytometry signal at the specified SMART-SpyCatcher concentration compared to the untreated sample is plotted against to the concentration of SMART-SpyCatcher added to the cells. Errors = standard error mean (n = 3 independent biological replicates). The data in panels **c**, **e**, and **f** employed the eNrdJ-1^cage^ pair within SMART-SpyCatcher.

## Results

### Engineering a protein switch

While CPS can in principle generate any desired POI output, we focused our efforts on designing a SMART device that would convert two antigen inputs into a unique protein loading dock when presented on the same cell. This would allow us to both monitor and characterize the device using fluorescence-based methods and later deliver a broader arsenal of functionalities. To achieve this, we concentrated on developing a SMART version of SpyCatcher003^22^, such that the isopeptide bond forming reactivity with the sixteen amino acid peptide SpyTag003 (Fig 1b) would be regulated by AND gated CPS.

We began by systematically screening multiple potential split sites in SpyCatcher003 (designated SpyN and SpyC) for compatibility with CPS (Extended Data Fig. 1a), initially using the caged Npu split intein (NpuN^cage^ and NpuC^cage^) in conjunction with the FKBP/rapamycin/FRB three hybrid system (Supplementary Fig. 1, Supplementary Table 1). Simulating cell-surface colocalization, we induced in-solution dimerization of SpyN-NpuN^cage^-FKBP and FRB-NpuC^cage^-SpyC by the addition of rapamycin (Extended Data Fig. 1b). Our screen revealed several pairs with an ability to fully recapitulate SpyCatcher003-SpyTag003 reactivity upon splicing (Supplementary Fig. 2 and Extended Data Fig. 1c). We elected to move forward with the SpyN^1–73^ and SpyC^74–113^ pair, since this had minimal background reactivity with the peptide probe prior to splicing, i.e., spontaneous complementation was very low (Extended Data Fig. 1c). The Npu split intein contains 4 cysteines in its structure and leaves a cysteine scar in the spliced product. Anticipating that these features could be deleterious to applications on the oxidizing environment of the cell surface, we exchanged Npu^cage^ for a caged version of the NrdJ-1 split intein, which leaves a serine at the splice site. The resulting constructs SpyN-NrdJ-1N^cage^-FKBP and FRB-NrdJ-1C^cage^-SpyC yielded near-complete splicing upon addition of rapamycin (Extended Data Fig. 1d), whereas no splicing nor reactivity with SpyTag003 was observed when the caged split intein was inactivated (Extended Data Fig. 1e). To facilitate further engineering of this system, we generated a fusion of NrdJ-1 and solved its crystal structure at 1.95 Å (Protein Data Bank [PDB] ID 8UBS, Extended Data Fig. 1f, Supplementary Fig. 3 and Supplementary Table 2). Using this structural information, we were able to design a fully active mutant of split NrdJ-1 in which the non-catalytic Cys76 in the IntN fragment is replaced by a valine (Extended Data Fig. 1g-j, Supplementary Fig. 4). We expected that this mutant would be better suited to applications on the cell surface. Hereafter, we refer to the individual components, SpyN-NrdJ-1N(C76V)^cage^ and NrdJ-1C^cage^(C76V)-SpyC as SpyN and SpyC, respectively, and their sum as SMART-SpyCatcher.

### On-cell performance of SMART-SpyCatcher

Next, SMART-SpyCatcher was refitted for on-cell activation by removing FKBP and FRB, while installing DARPins^23–25^ targeting the two model antigens HER2 and EGFR thereby generating αHER2-SpyN and SpyC-αEGFR (Supplementary Fig. 1). The ability of SMART-SpyCatcher to perform [HER2 AND EGFR] logic was assessed on K562 leukemia cells co-expressing HER2-eGFP and EGFR-iRFP^14^. Treatment of these cells with αHER2-SpyN and SpyC-αEGFR followed by SpyTag003 Alexa Fluor 594 conjugate (SpyTag003-AF594) resulted in the expected fluorescence signal, albeit weak, on the cell surface (Extended Data Fig. 2a). With the view to improving the reaction, we tested an enhanced version of NrdJ-1^cage^ (eNrdJ-1^cage^, Supplementary Fig. 4) with an engineered cage to facilitate an increased recruitment of SpyTag003-AF594^26^. This afforded a noticeably brighter AF594 fluorescence on the cell surface (Fig. 1c, Extended Data Fig. 2a). We therefore continued to use this actuator variant for all subsequent experiments unless otherwise stated. To rule out in-solution or at solution-surface interphase activation we also evaluated the activity of SMART-SpyCatcher on K562 cell lines naïve (wildtype), or single positive for either HER2-eGFP or EGFR-iRFP, all of which failed to elicit any response (Fig. 1c, Extended Data Fig. 2b,c). We further tested the on-target/off-target specificity of SMART-SpyCatcher when presented simultaneously with multiple decisions and applied a mixed-population flow cytometry assay for quantification (Fig. 1d, Supplementary Fig. 5,6). When incubated concurrently with equal amounts of the four cell lines in a mixture, SMART-SpyCatcher retained its target specificity (Fig. 1e, Extended Data Fig. 2d, Supplementary Fig. 6) even over a broad concentration range (nanomolar to micromolar), producing a sigmoidal dose-response only for K562^HER2+/EGFR+^ (Fig. 1f). Further experimental evidence supports a mechanism of action, where 1) αHER2-SpyN and SpyC-αEGFR colocalization, 2) splicing of SpyCatcher003, and 3) reactivity with SpyTag003 are all required as blocking any of these steps led to a complete loss of the AF594 signal (Extended Data Fig. 3a–c). Additionally, while SMART-SpyCatcher worked under various conditions (Extended Data Fig. 3d) and was AND gated (Extended Data Fig. 3e), its decision-making ability was lost when the cages were omitted from NrdJ-1, instead leading to massive crosslinking between single- and double-positive K562 cells (Extended Data Fig. 3f).

### Tuning SMART response dynamics

Encouraged by the ability of our protein device to make correct decisions, we created a suite of SMART-SpyCatcher pairs with a dynamic response range. Guided by the crystallographic information on NrdJ-1, we manipulated eNrdJ-1N^cage^ further (Extended Data Fig. 4a-e), while continuing to use eNrdJ-1C^cage^. This afforded multiple pairs with increased or decreased activities (Extended Data Fig. 4f, Supplementary Fig. 7a), without any compromise to target specificity (Extended Data Fig. 4g, Supplementary Table 4). For instance, introducing K114AK116A mutations within the cage sequence increased SpyTag003-AF594 recruitment by an additional 150%, while introducing A119K gave a 25% drop. Alternatively, altering the cage length of eNrdJ-1N^cage^ by either elongating or shortening it led to variants with activities ranging between 13% - 150% that of the standard 35 amino acid toehold while retaining target fidelity (Extended Data Fig. 5, Supplementary Fig. 7b, Supplementary Table 5). The ability to tune the responsiveness of the SMART system through engineering of the caging element expands the types of applications possible using the approach (see below).

### Exploring antigen limitations

HER2 and EGFR are known to engage in homo- and heteromultimeric surface clusters^27–30^ (Fig. 2a). In contrast, Epithelial Cell Adhesion Molecule (EpCAM) has not been reported to preferentially interact with either HER2 or EGFR and actuation would therefore occur from chance encounters. When tested on mixed K562 cell lines expressing either low endogenous levels or induced to overexpress EpCAM in combination with different profiles of HER2 and EGFR (Supplementary Fig. 8,9), we found that SMART-SpyCatcher computed AND functions on cells that fulfilled the assigned logic gate (Fig. 2b, Supplementary Fig. 9). SMART-SpyCatcher can therefore be used in targeting antigens that stochastically encounter each other or casually co-reside. SMART-SpyCatcher had a reduced activation profile on cells with low endogenous levels of EpCAM compared to those stably overexpressing the antigen. For instance, using αHER2-SpyN and SpyC-αEpCAM for [HER2 AND EpCAM] logic resulted in a 15-fold difference in AF594 signal intensity between K562^HER2+^ (which has low EpCAM levels, Supplementary Fig. 8c) and K562^HER2+/EpCAMhi^ (Figure 2b). Yet, the level of actuation on either cell line in response to [HER2 AND EpCAM] logic was adjustable using our tuned versions of eNrdJ-1N^cage^ (Extended Fig. 6a).

**Figure 2.**
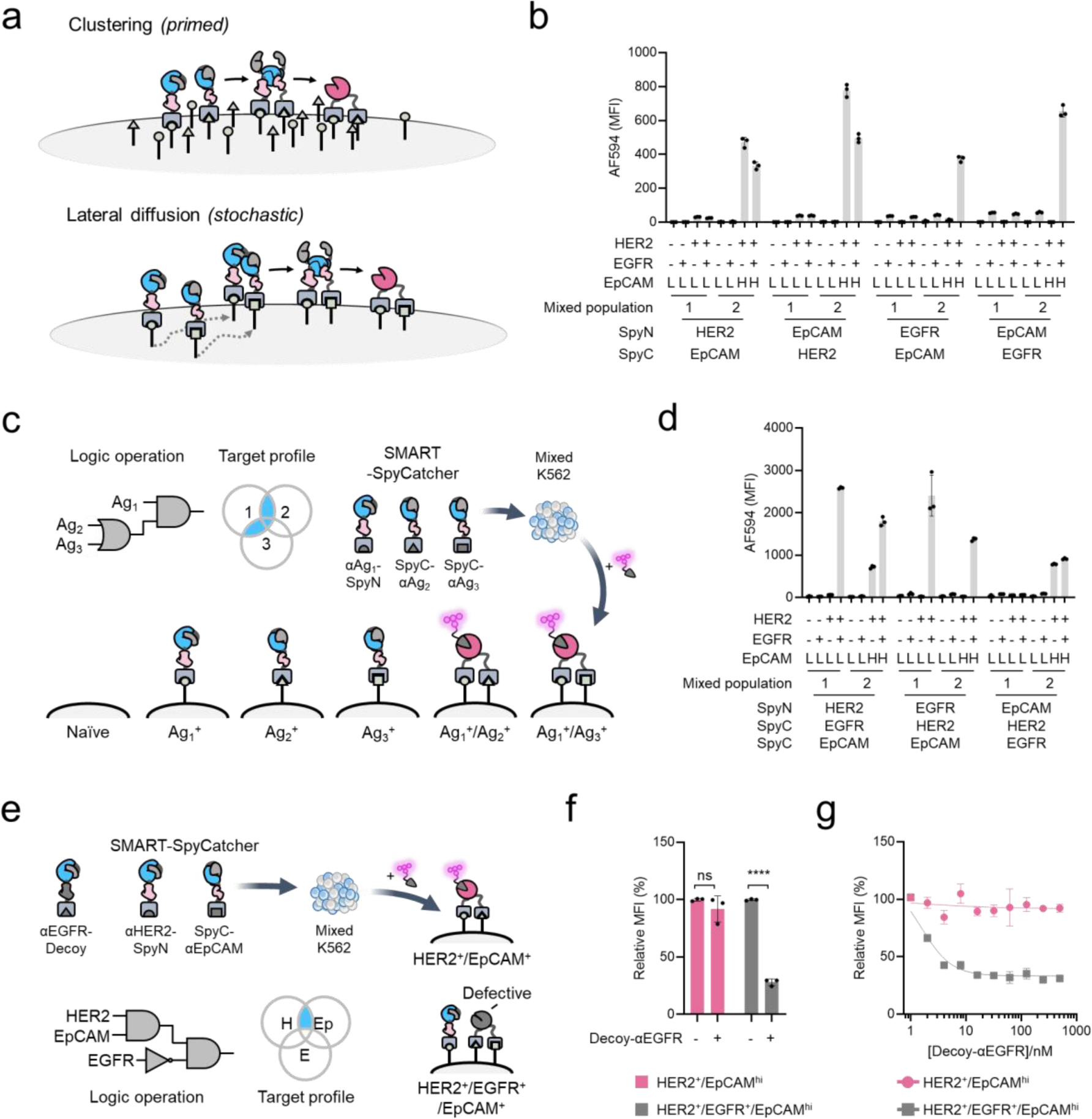
SMART-SpyCatcher computation of 2- and 3-input logic operations on mixed cell populations. **a**, Colocalization and activation of SMART-SpyCatcher can in principle occur from binding to two target antigens that are already closely associated, or which will stochastically encounter each other on the cell membrane. **b**, SMART-SpyCatcher (i.e., SpyN and SpyC) was assigned to operate through AND logic involving combinations of EpCAM with HER2 and EGFR. Two combinations of K562 cell lines were used to test for the actuation of SMART-SpyCatcher and thereby recruitment of SpyTag003-AF594. Mixed-population 1 consisted of equal amounts of K562 (wildtype), K562^EGFR+^, K562^HER2+^, and K562^HER2+/EGFR+^, whereas mixed-population 2 consisted of equal amounts of K562 (wildtype), K562^EGFR+^, K562^HER2+/EpCAMhi^, and K562^HER2+/EGFR+/EpCAMhi^. The antigen profile of each cell line is indicated below each bar plot (L and H designates low endogenous and high ectopic levels respectively for EpCAM), whereas the antigens targeted by the added SpyN and SpyC pairs (i.e. the targeting DARPins employed in the constructs) are indicated at the bottom. Experiments were performed with 100 nM SMART-SpyCatcher (eNrdJ-1^cage^), 100 nM SpyTag003-AF594. Data are presented as the mean of the AF594 median fluorescence intensities (MFI) from flow cytometry analysis with error bars signifying the standard error mean (n = 3 independent biological replicates; see Supplementary Table 3 for statistical one-way ANOVA followed by Dunnett’s test). **c**, Schematic illustrating cell targeting using [Ag1 AND either Ag2 OR Ag3] logic for recruitment using SMART-SpyCatcher. **d**, SMART-SpyCatcher cell targeting employing a 3-input AND/OR function. Experiment was performed as in panel **b** using the two described mixed K562 populations, but employing 3-inputs as indicated. Errors = standard error mean (n = 3 independent biological replicates; see Supplementary Table 7 for statistical one-way ANOVA followed by Dunnett’s test). **e**, The SMART system can encompass NOT operators as illustrated by [HER2 AND EpCAM NOT EGFR]. Here, Decoy-αEGFR obstructs AND logic output on K562^HER2+/EGFR+/EpCAMhi^ since splicing between the Decoy and SpyC generates a defective output unable to recruit SpyTag003-AF594. In contrast K562^HER2+/EpCAMhi^ elicit a normal response as Decoy-αEGFR does not bind to the cell. **f**, SMART-SpyCatcher cell targeting employing a 3-input [HER2 AND EpCAM NOT EGFR] logic and tested on mixed K562 population 2 described above. Decoy-αEGFR (100 nM) was added to the mixed cells followed by αHER2-SpyN (100 mM) and SpyC-αEpCAM (100 nM). Cells were then treated with 100 nM SpyTag003-AF594 and analyzed by flow cytometry. The mean AF594 signal for the indicate cell subpopulation is plotted relative to signal for control cells that had been treated with SpyN-αHER2 and SpyC-αEpCAM (i.e. no decoy). Errors = standard error mean (n = 3 independent biological replicates. Statistical significance was evaluated using a paired t-test (ns denotes not significant; **** denotes P < 0.0001). **g**, A dose-response experiment performed as in **f** with a dilution series of Decoy-αEGFR. Errors = standard error mean (n = 3 independent biological replicates).

### Three-input logic operations

While SMART can perform OR operations (Extended Data Fig. 6b-c), protein switch systems only marginally benefit from this type of logic when used in isolation. However, when combined with AND gates and utilizing three antigen inputs, the OR operator unlocks breadth in an otherwise narrowly defined targeting scheme. For instance, the [Ag_1_ AND either Ag_2_ OR Ag_3_] logic uses one fixed and two variable antigens (Fig. 2c), in which SMART targeting is limited by the fixed antigen. This was illustrated by employing αHER2-SpyN and SpyC-αEGFR/SpyC-αEpCAM, where the SMART-SpyCatcher functions by [HER2 AND either EGFR OR EpCAM] logic. Consistent with the expected AND/OR logic, three subpopulations - K562^HER2+/EGFR+^, K562^HER2+/EpCAMhi^, and K562^HER2+/EGFR+/EpCAMhi^ - responded to these three inputs (Fig. 2d). Extending this, varying the nature of the target input cues led to altered output in a predictable manner; for example, the combination of αEGFR-SpyN, SpyC-αHER2/SpyC-αEpCAM led to activation of only two cell lines, K562^HER2+/EGFR+^ and K562^HER2+/EGFR+/EpCAMhi^ (Fig. 2d). Combining AND/OR logic might therefore be useful in heterogeneous cell communities where a single antigen is shared by target and non-target cell populations while pairs of variable antigens can help define distinct subpopulations.

The lack of a surface marker between closely related cell populations could in principle be used for NOT gating for negative selection upon recognition. We envisioned making a NOT operator, where a decoy would generate a defective POI upon actuation, while outcompeting the reconstitution of the functional SpyCatcher003 output (Fig. 2e). To make such a decoy we used SpyN as a template, and first introduced K31E in its N-terminal SpyCatcher003 fragment, thereby disabling the ability to capture SpyTag003 once the POI was reconstituted (Extended Data Fig. 6d,e). The decoy’s reactivity with SpyC was then enhanced by tapping into the tunability of its split intein actuator component; we manipulated the cage of eNrdJ-1N^cage^ by introducing K114AK116A (Extended Date Fig. 4f) and shortening its C-terminal from 35 to 29 amino acid residues (Extended Data Fig. 5d and 6f). The decoy was then targeted to EGFR. Upon co-incubation of αHER2-SpyN, SpyC-αEpCAM, and αEGFR-Decoy with a mixture of K562 cell lines to assess [HER2 AND EpCAM NOT EGFR] logic, we observed that proper output associated with K562^HER2+/EGFR+/EpCAMhi^ was reduced by 72% whilst K562^HER2+/EpCAMhi^ was unaffected (Fig. 2f). Furthermore, dosing-in Decoy-αEGFR over a broad concentration range only minimally obscured actuation of αHER2-SpyN and SpyC-αEpCAM on K562^HER2/EpCAMhi^ cells, while K562^HER2/EGFR/EpCAMhi^ cells experienced a concentration-dependent decrease of SpyTag003-AF594 association (Fig. 2g).

### AND gated targeting of cancer cell lines

The Boolean reactivity of SMART-SpyCatcher to endogenous surface antigen landscapes was also evaluated on a variety of cancer cell lines using various combinations of SpyN and SpyC. First, we used DARPins labeled with AF594 to estimate the surface levels of HER2, EGFR, and EpCAM for each cell line using flow cytometry analysis (Fig. 3a, Extended Data Fig. 7a, and Supplementary Table 8). Then, the AND gated recruitment of SpyTag003-AF594 to the cells in response to SMART-SpyCatcher equipped with different sets of vectors was recorded (Fig. 3b, Extended Data Fig. 7a, and Supplementary Table 9). Plotting the quantity of the lesser-expressed antigen used in each AND gate against the recruitment of SpyTag003-AF594 achieved by the given logic operation gave a positive linear correlation accounting for 69% of the responsiveness of SMART-SpyCatcher (Fig. 3c). This implies that the level of SMART actuation is dominated by the quantities and possibly densities of the target antigens, where a threshold mechanism acts to protect cells with low or dilute surface levels. We further validated this observation on mixed human mammary cell lines displaying endogenous levels of HER2 and EpCAM (Fig. 3d-e, Extended Data Fig. 7b-d, and Supplementary Fig. 10), thereby mimicking the heterogeneous environment of diseased tissue. Two combinations of cells were tested: Epithelial MCF-10a (HER2^low^, EpCAM^low^) was combined with MCF-7 (HER2^low^, EpCAM^high^) as mixed mammary population 1 or with Sk-br-3 (HER2^high^, EpCAM^high^) as mixed mammary population 2. To enhance the recruitment of SpyTag003-AF594, we used SMART-SpyCatcher employing eNrdJ-1^cage^ with the additional K114AK119A mutations (Extended Data Fig. 4). Neither mixed mammary population effectively recruited SpyTag003-AF594 on their own, however, once αHER2-SpyN and SpyC-αEpCAM were added, recruitment occurred predominately for subpopulation Sk-br-3, with little-to-no probe associated with MCF-10a or MCF-7 (Fig. 3f,g, Supplementary Fig. 11,12). Collectively, these data illustrate how SMART-SpyCatcher can be used to sense and react to endogenous surface antigen landscapes and use this ability to distinguish between normal and cancerous subpopulations (e.g., Sk-br-3 *vs.* MCF-10a) by Boolean logic.

**Figure 3.**
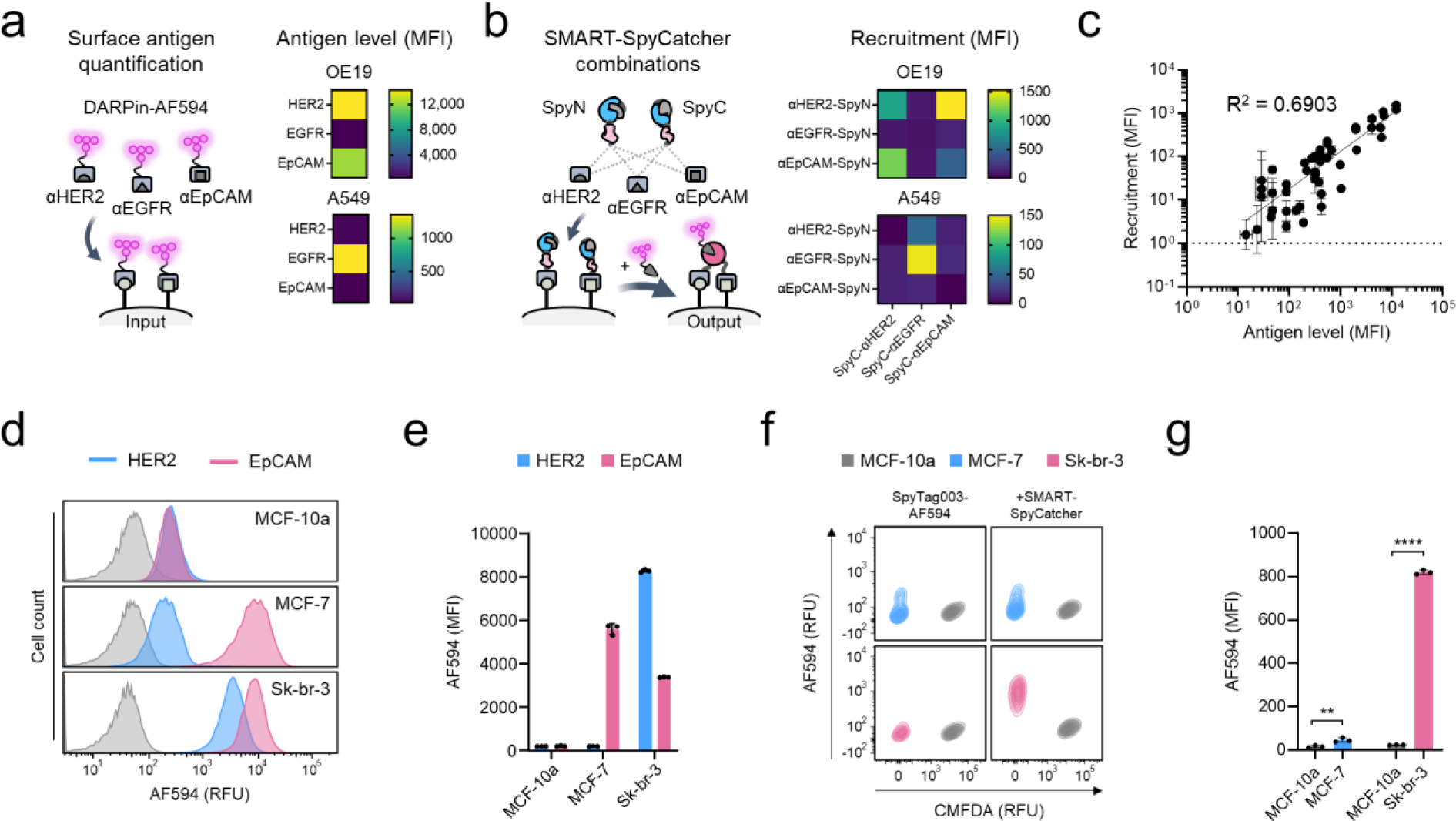
SMART-SpyCatcher performs AND logic on cell lines with endogenous antigen profiles. **a**, Target cells were profiled for their relative surface levels of HER2, EGFR, and EpCAM. Cells were treated with DARPins labeled with AF594 followed by flow cytometry analysis. The results for OE19 and A549 cells are shown on the right in the form of a heatmap. **b**, Combinations of SpyN and SpyC linked to targeting DARPins were used to solve the AND logic matrix for a given cell line. The results for OE19 and A549 cells are shown at right in the form of a heatmap. Cells treated with indicated SpyN and SpyC pairs (100 nM) followed by SpyTag003-AF594 (100 nM) were analyzed by flow cytometry. **c**, Summary of the data across all cancer cell lines tested using the SMART-SpyCatcher system. The amount of the lesser-expressed antigen used in each AND gate is plotted against the observed recruitment of SpyTag003-AF594 (as determine by flow cytometry) by a given SMART-SpyCatcher logic operation. Errors = standard error mean (n = 3 independent biological replicates). **d**-**e**, The endogenous surface levels of HER2 and EpCAM on mammary cell lines MCF-10a, MCF-7, and Sk-br-3 were profiled as in panel **a**. Representative flow cytometry analysis of the three cell lines is in shown in panel **d** (untreated samples are shown in grey) and quantified in panel **e**. The two antigens were detected in all three cell lines and categorized as low (MFI < 1000) or high (MFI ≥ 1000). Errors = standard error mean (n = 3 independent biological replicates). RFU, relative fluorescence units. **f-g**, SMART-SpyCatcher assigned for [HER2 AND EpCAM] logic was tested in flow cytometry experiments using mixed mammary cell population 1 (equal amounts of MCF-10a^HER2lo/EpCAMlo^ and MCF-7^HER2lo/EpCAMhi^) and mixed mammary population 2 (equal amounts of MCF-10a^HER2lo/EpCAMlo^ and Sk-br-3^HER2hi/EpCAMhi^). MCF-10a cells were pre-stained with CMFDA, which labels intracellular proteins. Each population was incubated with 100 nM SpyTag003-AF594 in the absence or presence of 100 nM αHER2-SpyN and SpyC-αEpCAM (employing eNrdJ-1^cage^ with K114AK116A). Shown in panel **f** are representative flow cytometry plots of the recruitment of SpyTag-AF594 by the subpopulations under the different conditions. The data is quantified in panel **g** in which the flow cytometry data are presented as the mean of the individual median fluorescence intensities (MFI) of the AF594 signal with error bars signifying the standard error mean (n = 3 independent biological replicates; paired t-test was performed for statistical analysis, ** denotes P < 0.01, **** denotes P < 0.0001).

### Selective delivery of functionalities

SMART-SpyCatcher generates a unique protein loading dock once activated on the cell surface. In principle, this can be used to recruit a wide array of functionalities to sculpt the cell surface. Of recent interest is the use of bifunctional small molecules to decorate the surface of a selected cell population for the recruitment of antibodies^31,32^. Inspired by this, we asked whether SpyTag003 conjugated to dinitrophenol (SpyTag003-DNP) could recruit anti-DNP antibodies to K562^HER2+/EGFR+^ cells upon SMART-SpyCatcher actuation (Fig. 4a,b). As hoped, incubation of these cells with SMART-SpyCatcher for [HER2 AND EGFR] logic and SpyTag003-DNP led to recruitment of an anti-DNP antibody in a fashion dependent upon the biochemical activity of SMART (Fig. 4c, Extended Fig. 8a). In a mixed K562 population setting, SMART-SpyCatcher successfully led to the recruitment of anti-DNP antibody to the targeted cell lines while avoiding engagement with non-target subpopulations (Fig. 4d and Extended Fig. 8b).

**Figure 4.**
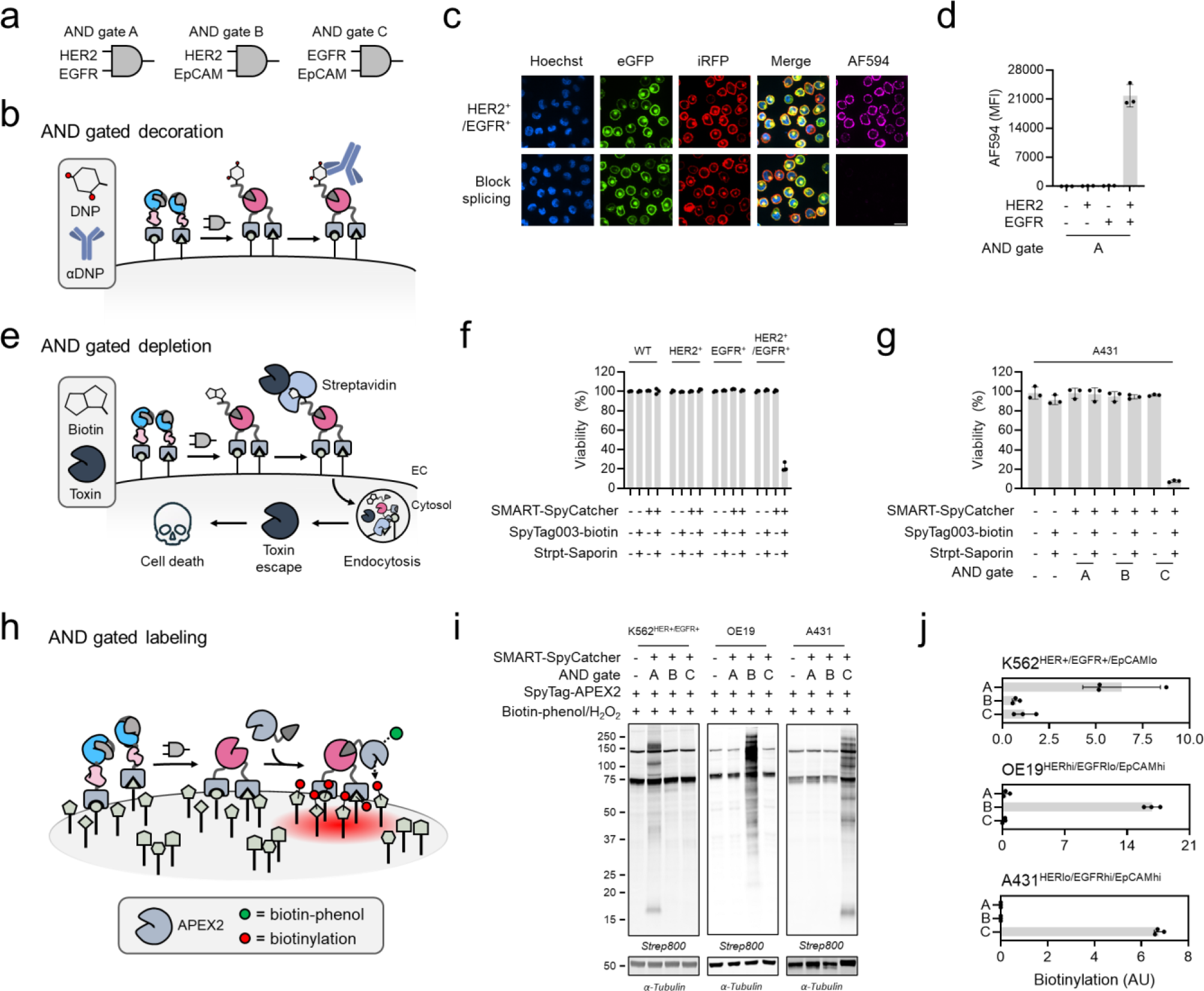
Applications of AND gated cell specific surface sculpting. **a**, Three AND logic gates involving HER2, EGFR, and EpCAM. **b**, Schematic illustrating AND-gated attachment of a small antibody-recruiting molecule such as dinitrophenol (DNP) to a cell surface using the SMART-SpyCatcher system. **c**, K562^HER2+/EGFR+^ cells were treated with 100 nM SMART-SpyCatcher (employing eNrdJ-1^cage^) for [HER2 AND EGFR] logic and SpyTag003-DNP (100 nM). Following washing, the cells were treated with an anti-DNP antibody, followed by an AF594-conjugated secondary antibody and then visualized by confocal microscopy. Cell nuclei are stained with Hoechst, while HER2 and EGFR are tagged with eGFP and iRFP, respectively. The bottom row shows a control experiment in which the protein splicing reaction is blocked by alkylation of a key cysteine residue in the split intein. Scale bar equals 20 μm. **d**, The antibody recruitment workflow was applied to a mixed K562 cell population (equal amounts of K562, K562^HER2+^, K562^EGFR+^, and K562^HER2+/EGFR+^) using AND gate system A (indicated in panel **a**). Following the labeling reaction, cells were analyzed by flow cytometry. Data are presented as the mean of the median AF594 fluorescence intensity (MFI) with error bars signifying the standard error mean (n = 3 independent biological replicates, see Supplementary Table 3 for statistical one-way ANOVA followed by Dunnett’s test). **e**, Schematic illustrating AND gated cell depletion using the SMART-SpyCatcher system. Surface decoration with SpyTag003-biotin is used to recruit a Streptavidin-Saporin disulfide conjugate, which leads to cell death upon internalization and lysosomal escape of the toxin. **f**, The mixed K562 cell population employed in panel **d** was treated with a two-dose regimen of 100 nM SMART-SpyCatcher (employing [HER2 AND EGFR] logic), SpyTag003-biotin (100 nM) and Streptavidin-Saporin (20 nM) employing a 24 hr interval. The cells were then analyzed after an additional 72 hr by flow cytometry and the data presented as % viability relative to untreated wildtype cells (see Methods for details). Errors = standard error mean (n = 3 independent biological replicates). **g**, A431 cells (which have high levels of EGFR and EpCAM, but low HER2) were treated with the indicated SMART-SpyCatcher system (100 nM, AND gates A-C as indicated in panel **a**) followed by SpyTag003-biotin (100 nM) and Streptavidin-Saporin. A two-dose regimen was employed as in panel **f**, and cell viability was determined using an XTT assay, the data plotted relative to the untreated cells. Errors = standard error mean (n = 3 independent biological replicates). **h**, Schematic illustrating the principle of AND gated protein proximity labeling. A SpyTag003-APEX2 fusion protein is delivered to a cell surface modified using the SMART-SpyCatcher system targeting two cell surface antigens. Proteins in the vicinity of these colocalized antigens are then labeled with biotin by a standard APEX2 workflow employing a biotin-phenol probe and activation with H_2_O_2_. **i**, Cell lines were treated with the indicated SMART-SpyCatcher systems (100 nM, AND gates A-C as shown in panel **a**) followed by SpyTag003-APEX2 (300 nM). Cells were then treated with biotin-phenol (250 μM) and H_2_O_2_ (100 mM), before being analyzed by Western blotting. **j**, Quantification of the APEX2 labeling experiment shown in panel **i**. The total biotinylation signal in each lane was quantified by Fiji, normalized to the signal from tubulin, and finally corrected for any endogenously biotinylated proteins and potential background labeling by subtracting the value obtained from the control experiments that had not been treated with SMART-SpyCatcher. Errors = standard error mean (n = 3 independent biological replicates).

As an additional application, we reasoned that stringent delivery and intake of a toxin could allow for the exclusive depletion of a target cell line in a heterogeneous population. While many ribosome-inactivating proteins can facilitate their own cellular intake, the plant toxin Saporin lacks a targeting domain to do so. Whenever internalized, however, Saporin depurinates ribosomes to induce translational arrest and cell death^33^. We used a Streptavidin-Saporin disulfide conjugate to test for cell selective depletion (Fig. 4e). First, however, we validated that AND gated CPS of SpyCatcher003 and delivery of SpyTag003-biotin would recruit a NeutrAvidin Rhodamine Red-X (NA-RRx) conjugate and induce its internalization to endosomal/lysosomal compartments for K562^HER2+/EGFR+^ cells and that the method was selective (Extended Fig. 9a–c). Then we replaced NA-RRx with a Streptavidin-Saporin conjugate in a similar setup. Mixed K562 cells were treated with SMART-SpyCatcher assigned for [HER2 AND EGFR] logic and SpyTag003-biotin and subsequently cultured with Streptavidin-Saporin (Extended Data Fig. 9d,e). After an additional 72 hr the cultures were sampled for variations in cell composition. While a one-dose regimen led to a 53% reduction of K562^HER2+/EGFR+^ cells (Extended Data Fig. 9f and Supplementary Fig. 13), a two-dose regimen led to a 78% depletion of the subpopulation (Fig. 4f and Supplementary Fig. 14). Importantly, neither wildtype, K562^HER2+^ nor K562^EGFR+^ cell lines were depleted to a similar extent in the co-culture following either the one- or two-dose regimens. We also tested depletion of A431 cells, which display low levels of HER2 but high levels of EGFR and EpCAM (Extended Data Fig. 7a). For this, we used SMART-SpyCatcher employing eNrdJ-1^cage^ with the additional K114AK119A mutations to enhance recruitment of SpyTag003-biotin. Compared to an untreated control, A431 cells were depleted by 92% when using a two-dose regimen that employed SpyTag003-biotin, Streptavidin-Saporin and SMART-SpyCatcher operating by [EGFR AND EpCAM] logic (Fig. 4g). As expected, an identical workflow using SMART-SpyCatcher set up for [HER2 AND EGFR] or [HER2 AND EpCAM] logics failed to achieve comparable reductions in these cells, and no reduction was observed when the toxin conjugate was omitted. Hence, SMART-SpyCatcher unlocks logic gated cell depletion strategies^34–36^.

Finally, we established a workflow for performing context-specific protein proximity-labeling (Fig. 4h). For this we generated a fusion between SpyTag003 and APEX2^37^ (Extended Data Fig. 10a). This construct was then used in standard APEX2 labeling studies employing a biotin-phenol probe and activation with H_2_O_2_. Robust cell biotinylation was observed in K562^HER2+EGFR+^ cells that had been treated with SMART-SpyCatcher for [HER2 AND EGFR] logic, whereas no proximity labeling was observed in the various control cell lines (Fig. 4i and Extended Data Fig. 10b). We validated these results further on K562^HER2+/EGFR+^ (HER2^hi^, EGFR^hi^, EpCAM^lo^), OE19 (HER2^hi^, EGFR^lo^, EpCAM^hi^), and A431 (HER2^lo^, EGFR^hi^, EpCAM^hi^) cells lines while performing either [HER2 AND EGFR], [HER2 AND EpCAM], or [EGFR AND EpCAM] logic operations prior to incubation with SpyTag003-APEX2 and biotin-phenol labeling. Only when SMART-SpyCatcher was matched with the correct antigen profile of the individual cell line did we observe APEX2-dependent biotinylation (Fig. 4i,j, Extended Data Fig. 10c). These results indicate that SMART-SpyCatcher can be used to perform AND gated proximity-labeling to study and map the neighborhood for two nearby antigens.

## Discussion

The close similarity in cell surface landscape between various tissue types and their disease states poses a key challenge in securing precision targeting using cell surface antigens. Relatively few surface markers have been recognized as truly disease specific and, thus, reliance on single antigens for therapeutic/diagnostic targeting can be a liability. By contrast, exploiting multiple cell surface features in the targeting step opens the way to Boolean logic operations that should lead to greatly improved specificity^1–3^. For instance, next generation Chimeric Antigen-Receptor T Cells can compute multiple antigen inputs to trigger targeted cell killing with limited adverse effects on non-targets^11,38^. While several protein-based systems have been developed with similar decision-making capabilities^14–17^, the design and optimization of such devices remains a highly complex task. In this study, we have developed a synthetic biology platform, SMART, that at its core, functions as a protein actuator based on conditional protein *trans*-splicing with tunable responsiveness to cellular inputs. This SMART device can sense multiple cell surface features and convert these to a user-defined functional output based on Boolean logic operations. The current version involves restoration of a functional SpyCatcher003 protein dock, which is used to selectively recruit a variety of activities to a target cell population. The existing SMART-SpyCatcher system is highly versatile since it requires fusing a short peptide (the SpyTag003) to whatever cargo is to be delivered to the target cells. As demonstrated, this allows both biological and synthetic activities to be harnessed.

It is important to stress that our SMART system is highly modular. The central CPS-based actuator can, in principle, be wedded to a variety of outputs due to the promiscuity of the NrdJ-1 split intein it employs^26^. Thus, we imagine that split versions of many other proteins could be directly generated on cell surfaces gated by AND, OR or NOT logic operations. The modularity of the system also extends to the sensor components being used; a variety of existing (i.e., ‘off the shelf’) antibodies, nanobodies, lectins, or even peptide ligands should be directly compatible with the platform. Finally, while we currently focus on cell surface applications of SMART-SpyCatcher, we expect that this version of the technology will find application in other contexts, including logic-gated imaging, proximity-proteomics and possibly even genomics within permeabilized cells.

## Data availability

All relevant data are included in the manuscript and the supplemental information and are available upon request. Coordinates and structure files have been deposited to the Protein Data Bank (PDB) under ID 8UBS. Expression plasmids and cell lines are available upon request.

## Acknowledgments

This work was funded by NIH-GMS grant R01 GM086868 and by funds from the Ludwig Institute for Cancer Research. C.K. is supported by EMBO (ALTF 1189-2020). N.E.S.T. is supported by an NIH postdoctoral fellowship (GM149123). X.Y. is supported by a graduate fellowship from the China Scholarship Council (CSC). We thank members of the Muir lab for many helpful discussions during the course of this work. We thank P. Jeffrey for technical assistance with setting up crystal trays, looping, data collection and data processing. We thank C. DeCoste and K. Rittenbach for technical assistance with flow cytometry and cell sorting and instrument use (Princeton University Flow Cytometry Resource Facility, Department of Microbiology; supported, in part, with funding from NCI-CCSG P30CA072720-5921). We thank S. Wang and G. Laevsky for technical assistance with confocal microscopy and instrument use (Princeton University Confocal Imaging Facility, a Nikon Center of Excellence, Department of Molecular Biology). We thank D. Baker (University of Washington) for sharing K562 cell lines, Y. Kang (Princeton University) for sharing MCF-10a, MCF-7, LoVo, and A594 cell lines, and S. Lipkowitz (National Institutes of Health) for sharing the Sk-br-3 cell line.

## Author contributions

C.K. and T.W.M. conceived the work. C.K., N.E.S.T., X.Y., G.E., and T.W.M. designed and executed the experiments. C.K. and T.W.M. prepared this manuscript.

## Materials and Methods

### General materials

Common reagents and chemicals were purchased from MilliporeSigma (Burlington, MA) unless stated otherwise. 2,4-dinitrochlorobenzene (CAS NO 97-00-7), *tert*-butyl bromoacetate (CAS NO. 5292-43-3), *N*-hydroxysuccinimide (CAS NO. 6066-82-6), *N,N’*-Dicyclohexylcarbodiimide (CAS NO. 538-75-0), Celite 545 (CAS NO. 68855-54-9), and 1-(2-aminoethyl)maleimide hydrochloride (CAS NO. 134272-64-3) were purchased from MilliporeSigma (Burlington, MA). 2-[2-(2-aminoethoxy)ethoxy]ethanol (CAS NO 86770-74-3), and Biotin-PEG3-amine (CAS NO 359860-27-8) were purchased from Ambeed (Arlington Heights, IL). Triethylamine (CAS NO 121-44-8) was purchased from Thermo Fischer Scientific (Waltham, MA). Dithiothreitol (DTT, CAS NO 3483-12-3), isopropyl-β-thiogalactopyranoside (IPTG, CAS NO 367-93-1) and bovine serum albumin (BSA) were obtained from Gold Biotechnology (Burlington, MA). Alexa Fluor 594 C5 maleimide was purchased from BroadPharm (San Diego, CA). Biotin maleimide (CAS NO 116919-18-7) was purchased from MilliporeSigma (Burlington, MA).

Oligonucleotide primers were purchased from MilliporeSigma (Burlington, MA). gBlock® Gene Fragments were purchased from Integrated DNA Technologies. PrimeSTAR HS DNA Polymerase was purchased from Takara Bio (Kusatsu, Japan). Gibson Assembly Master Mix was purchased from New England Biolabs (Ipswich, MA). DNA purification kits were purchased from Qiagen (Valencia, CA). PCR Purification & Gel extract columns were purchased from Thomas Scientific (Swedesboro, NJ). All plasmid sequencing was performed by GENEWIZ (South Plainfield, NJ) or alternatively by Plasmidosaurus (Eugene, OR).

Nickel nitrilotriacetic acid (Ni-NTA) resin was obtained from Thermo Fisher Scientific (Waltham, MA). MOPS-SDS running buffer was obtained from Boston Bioproducts (Ashland, MA). Criterion Cassettes, acrylamide, ammonium persulfate, tetramethylethylenediamine, and Econo-Pac® 10DG Columns were obtained from Bio-Rad (Hercules, CA). Nitrocellulose membrane (0.45 μm) for Western blotting was purchased from Thermo Fisher Scientific (Waltham, MA).

NeutrAvidin™ Rhodamine Red™-X conjugate was purchased from Thermo Fischer Scientific (Waltham, MA). Streptavidin-Saporin (Streptavidin-ZAP) was purchased from Advanced Targeting Systems (Carlsbad, CA).

Dulbecco’s Modified Eagle Medium (Gibco), RPMI-1640 Medium (Gibco), McCoy’s 5a Medium Modified (Gibco), F-12K Medium (Gibco), Dulbecco’s phosphate-buffered saline (DPBS, Gibco), Penicillin-Streptomycin (5,000 U/mL), Trypsin-EDTA (0.25%), Trypsin-EDTA (0.05%), and Falcon™ Standard Tissue Culture Dishes were purchased from Thermo Fisher Scientific (Waltham, MA). Mammary Epithelial Cell Growth Medium kit was purchased from MilliporeSigma (Burlington, MA). Fetal Bovine Serum (Heat Inactivated) was purchased from Bio-Techne (Minneapolis, MN). XTT Cell Viability Kit was purchased from Cell Signal Technologies (Danvers, MA). Glass bottom 24 well plates were purchased from Cellvis (Mountain View, CA). Hoechst 33342 solution was obtained from Invitrogen (Carlsbad, CA).

Human cell lines K562^EpCAMlo^ (wildtype, CCL-243), K562^HER2+^, K562^EGFR+^, K562^HER2+/EGFR+^, K562^HER2+/EpCAMhi^, K562^HER2+/EGFR+/EpCAMhi^ were generous gifts from Prof. David Baker (University of Washington, WA). Human cell lines MCF-10a (CRL-10317), MCF-7 (HTB-22), LoVo (CCL-229), A594 (CCL-185) were generous gifts from Prof. Yibin Kang (Princeton University, NJ). Human cell line Sk-br-3 (HTB-30) was a generous gift from Dr. Stan Lipkowitz (National Cancer Institute, Bethesda, MD). Human cell line OE19 (JROECL19) was purchased from MilliporeSigma (Burlington, MA). Human cell lines HCT-116 (CCL-247) and A431 (CRL-1555) were purchased from ATCC (Manassas, VA).

Size exclusion chromatography was performed on an ÄKTA Fast Performance Liquid Chromatography (GE Healthcare) system. Analytical-scale reverse-phase high performance liquid chromatography (RP-HPLC) was conducted on an Agilent 1100 Series or an Agilent 1260 Infinity system equipped with a C18 Vydac column (5 mM, 4.6 x 150 mm) at a flow rate of 1 mL/min. Semi-preparative RP-HPLC was conducted on an Agilent 1260 Infinity system equipped with a Waters XBridge BEH C18 column (5 mM, 10 x 250 mm) at a flow rate of 4 mL/min. Preparative-scale RP-HPLC was conducted on a Waters prep LC system consisting of a Waters 2545 Binary Gradient Module and a Waters 2489 Ultraviolet (UV)-Visible detector equipped with a C18 Vydac column (10 mM, 22 x 250 mm). HPLC solvents were H_2_O with 0.1% TFA (Solvent A) and 90% acetonitrile in water with 0.1% TFA (Solvent B). Proteins and peptides were characterized by electrospray ionization mass spectrometry (ESI-MS) on a Bruker Daltonics MicroTOF-Q II mass spectrometer.

Coomassie-stained SDS-PAGE gels and Western blots were imaged on an Odyssey system (LI-COR). Densitometry measurements were performed using FIJI (National Institutes of Health)^39^. Molecular graphics and analyses were performed with PyMOL v.2.5, developed by Schrödinger, Inc. Graph plots and statistical analysis were made in GraphPad Prism v.9.0.0 (121).

### Chemical Synthesis

Synthetic protocols for DNP-maleimide and biotin phenol are provided in Supplemental Methods.

### DNA cloning

All expression vectors were based on a pET backbone vector with a gene for kanamycin resistance. DNA fragments amplified by Polymerase Chain Reaction (PCR) were inserted into a gene cassette using Gibson Assembly. The gene cassette included an upstream T7 promoter under regulation of a *lac* operator, an ATG start codon in frame with a N-terminal His_6_-SUMO fusion tag when specified and ended with a TAA stop codon and a T7-terminator sequence.

Site-directed mutagenesis (including missense substitutions, insertions, and deletions), in addition to amplifications of gene inserts and plasmid backbones were performed by PCR. To a solution of 10 µL 1 ng/µL plasmid, 1 µM forward primer, 1 µM reverse primer was added 10 µL 2x PrimeSTAR DNA polymerase Master Mix. The mixture was used in a PCR reaction. The completed PCR reaction was restriction digested with 20 units of DpnI for 1 hr at 37 °C before the PCR amplicon was isolated by a general PCR clean up protocol. The purified PCR amplicon was then used in a Gibson assembly reaction (see below). Alternatively, full plasmid amplicons from site directed mutagenesis was used directly in heat-shock transformations of chemically competent *Escherichia coli* DH5α cells, which were plated on LB-agar plates supplemented with 50 µg/mL kanamycin. Single colonies were picked, grown and the plasmid isolated before being verified by Sanger sequencing.

DNA components prepared by PCR using primers designed to have 15-20 base pair overlaps were used in Gibson Assembly reactions. Gibson Assembly reactions were set up by mixing 2 µL DNA fragments (100 ng plasmid backbone amplicon, 3-5 molar excess gene insert amplicon) with 2 µL 2x NEBuilder® HiFi DNA Assembly master mix. The reaction was incubated for 15 min at 50 °C, cooled and thereafter used in a heat-shock transformation of chemically competent *Escherichia coli* DH5α, which were plated on LB-agar plates supplemented with 50 µg/mL kanamycin. Single colonies were picked, grown and the plasmid isolated before being verified by Sanger sequencing.

### Recombinant protein expression and purification

Chemically competent *Escherichia coli* BL21(DE3) cells were heat-shock transformed with a pET vector carrying the gene cassette for the protein of interest. Cells were grown overnight at 37 °C in 8 mL LB medium supplemented with 50 µg/mL kanamycin. This overnight culture was used to inoculate an expression culture of 1 L LB medium supplemented with 50 µg/mL kanamycin. In general, the expression culture was incubated at 37 °C until it reached an OD_600_ of 0.4, whereafter it was cooled for 20 min at 18 °C. Protein expression was then induced by the addition of 0.1 mL 1 M IPTG and the culture left overnight at 18 °C. For expression of full-length NrdJ-1 used in the crystallography study and for any SpyTag003 and stand-alone DARPin constructs, the expression culture was incubated at 37 °C until it reached an OD_600_ of 0.6. The expression was then induced by the addition of 0.1 mL 1 M IPTG and the culture left for 4 hr at 37 °C. In all cases, cells were harvested by centrifugation at 3,500 g, 18 °C for 20 min and suspended in lysis buffer containing 20 mL 50 mM NaH_2_PO_4_ (pH 8.0), 300 mM NaCl, 20 mM imidazole, supplemented with 1 mM dithiothreitol (DTT) and 1 mM phenylmethylsulfonyl fluoride (PMSF). Cell suspensions were then either stored at -20 °C until further use or used directly. Soluble protein was extracted by subjecting the cell suspension to sonication using a duty cycle of 20s on, 30s off, at 30% amplitude while cooled on an ice bath, after which a cleared lysate was produced by centrifugation at 35,000 g, 4 °C for 20 min. The cleared lysate was passed through a preequilibrated Ni^2+^-nitrilotriacetic acid (NTA) column (2 mL resin slurry per liter culture) and the flow-through discarded. The column was then washed with 50 mL lysis buffer, before the protein was eluted using 6 mL lysis buffer supplemented with 250 mM imidazole. His_6_-SUMO tagged proteins were treated overnight with His_6_-Ulp1 protease while being dialyzed against lysis buffer supplemented with 1 mM DTT. The dialyzed sample was then passed through a preequilibrated Ni^2+^-NTA column to remove any cleaved His_6_-SUMO tag and His_6_-Ulp1 protease. The flow-through and an additional 6 mL lysis buffer passed through the column was collected, combined, and concentrated to 0.5 mL. The concentrated sample was filtered through a 0.22 μm spin filter and the flow-through applied to a Superdex 200 10/300 GL (Cytiva Life Sciences) size-exclusion column using 100 mM NaH_2_PO_4_ (pH 7.2), 150 mM NaCl, 1 mM ethylenediaminetetraacetic acid (EDTA), and 1 mM DTT as the eluent at a flowrate of 0.5 mL/min. Pure fractions were identified by SDS-PAGE, validated by analytical RP-HPLC, and the protein mass confirmed by ESI-TOF MS. Pure fractions were supplemented with 10% v/v glycerol, aliquoted and flash-frozen in liquid nitrogen before being stored at -80 °C until further use. Analytical data for all proteins is shown in Supplemental Figure 1 and Supplemental Table 1. The fully annotated amino acid sequences of overexpressed constructs used in this study are given in Supplementary Table 10.

### Western blotting

Samples were run on an SDS-PAGE gel. For Western blotting, the gel was used for a transfer reaction onto a nitrocellulose membrane, which was subsequently blocked with TBS-T (25 mM Tris, 150 mM NaCl, 0.1% v/v Tween-20, pH 7.7) supplemented with 4% w/v skimmed milk powder for 1 hr at RT. The membrane was washed 3 x 5 min with TBS-T and incubated with primary antibodies at specified dilution (Supplementary Table 11) on an orbital shaker for 1 hr at room temperature or alternatively overnight at 4 °C. After washing 3 x 5 min with TBS-T, the IRDye secondary antibody or alternatively IRDye Streptavidin were applied for 30 min to 1 hour at room temperature, before imaging on a Li-Cor Odyssey imager (Li-Cor).

### X-ray crystallography

Crystallography studies employed a fused version of NrdJ-1 containing C1A and N145A inactivating mutations and SGG and SEI as N- and C-terminal extein sequences, respectively. The protein was dialyzed against a buffer containing 25 mM HEPES (pH 7.5), 150 mM NaCl, then concentrated to 40 mg/mL, and finally flash-frozen as aliquots in liquid nitrogen before being stored at -80 °C for further use. Initial crystallization conditions were established using the SaltRx HT screen (Hampton Research) employing a Phoenix crystallization robot (Art Robbins). Crystals were grown at 4 °C by the sitting-drop vapor diffusion method. A focused screen was centered on conditions with increasing concentrations of sodium formate at various pH values. Optimal crystals were obtained after one week using a solution of 4.5 M sodium formate (pH 7.0). Diffraction data was obtained at the National Synchrotron Light Source II (Brookhaven National Laboratory), beamline 17-ID-1. The data was processed using the XDS package^40^. The phase information was determined by molecular replacement using PHASER in the CCP4 suit^41^ and using an *in silico* AlphaFold2^42,43^ model of full-length NrdJ-1 as input. Iterative rounds of model building in Coot^44^ and refinements in PHENIX Refine (version 1.17_3644)^45^ were performed to obtain the final structure. Data collection and refinement statistics are displayed in Supplementary Table 2.

### Protein conjugation reactions

Purified protein containing a C-terminal cysteine (SpyTag003-Cys, SpyTag003^D117A^-Cys, αHER2-Cys DARPin, αEGFR-Cys DARPin, αEpCAM-Cys DARPin) or catalytical cysteine (αHER2-SpyN-eNrdJ-1N^cage^) was reduced with 1 mM DTT for 20 min on ice to prepare it for conjugation chemistry. The sample was washed with buffer (100 mM NaH_2_PO_4_ (pH 7.2), 150 mM NaCl2, 1 mM EDTA) using spin-filtration to remove small molecule thiols. The reduced protein was either flash-frozen in liquid nitrogen and stored at -80 °C for later use or used directly in conjugation reactions as described in the following sections. The reduced protein was mixed with a molar excess of the required alkylating agent (iodoacetamide, Alexa Fluor 594 maleimide, biotin-maleimide, or 2,4-dinitrophenyl-maleimide) and incubated for 20 min at room temperature in the dark. Reactions were monitored by RP-HPLC and ESI-TOF MS and quenched with 1 mM DTT once completed. The reaction mixture was then passed over an Econo-Pac® 10DG Column using 100 mM NaH_2_PO_4_ (pH 7.2), 150 mM NaCl, 1 mM EDTA, 10% v/v glycerol as the eluent. The product was validated by RP-HPLC and ESI-TOF MS, flash-frozen and stored at -80 °C. Analytical data for all protein conjugations is provided in Supplemental Figure 1 and Supplemental Table 1.

### Rapamycin-induced protein *trans*-splicing reactions

Screening for the optimal split site within SpyCatcher003 was performed by combining complementary caged split intein SpyN/SpyC fusion proteins (1 µM of each) in splicing buffer (100 mM NaH_2_PO_4_ (pH 7.2), 150 mM NaCl, 1 mM EDTA, 1 mM DTT, 10% v/v glycerol) in the presence of 2 µM SpyTag003 or conjugates thereof. Protein *trans*-splicing was initiated either by addition of rapamycin (final concentration 10 µM from a 50 µM DMSO stock) or TEV protease (10 units). Reaction mixtures were incubated for 24 hr at 37 °C before being analyzed by SDS-PAGE/Western blotting.

### Mammalian Cell culture

Mammalian cell lines were cultured in media as detailed in the Supplementary Table 12 supplemented with 100 U/mL penicillin/streptomycin in an incubator at 37 ᵒC and 5% CO_2_. All cell lines were regularly tested free of mycoplasma.

### Confocal microscopy

#### Suspension cell lines

The specific cell line or mixture was treated as specified in the relevant section. Post-treatment the cells were suspended in DBPS, 1% w/v BSA, 2 mM CaCl_2_ supplemented with Hoechst 33342 (10 µg/mL) and incubated for 30 min at room temperature before being washed twice in buffer. The cells were then transferred to a glass bottom plate and allowed to settle before being imaged using 40x magnification on a Nikon A1/HD25 microscope (Nikon Instruments, Inc., Melville, NY). Processing of fluorescence microscopy images was performed using FIJI (National Institutes of Health).^1^

#### Adherent cell lines

The specific cell line was plated on a glass bottom plate, cultured, and then treated as specified in the relevant section. Post-treatment DBPS, 1% w/v BSA, 2 mM CaCl_2_ supplemented with Hoechst 33342 (10 µg/mL) was added to the plate and the adherent cells incubated for 30 min at room temperature before being washed twice in buffer. The cells were then transferred to a glass bottom plate and allowed to settle before being imaged as described above.

### Flow cytometry

Cell lines were suspended as 1,000,000 cells/mL in cold in DBPS, 1% w/v BSA, 2 mM CaCl_2_ (supplemented with 50 mM EDTA for adherent cell cultures), filtered through a cell strainer and kept on ice. Flow cytometry data acquisition was obtained on a BD^®^ LSR II Flow Cytometer and results analyzed using FlowJo^TM^ 10.8.1. Single color controls were included for compensation adjustments. Examples for the gating strategy used for flow cytometry analysis of mixed K562 populations are given in Supplementary Fig. 5,6 and Supplementary Fig. 8,9. Examples for the gating strategy used for flow cytometry analysis of mixed mammary populations are given in Supplementary 10-12.

### Cell surface labeling experiments using SMART-SpyCatcher

#### Suspension cell lines

Individually cultured K562 cell lines were spun down and counted to estimate the cell concentration. Then approximately 200,000 cells of an individual K562 cell culture were isolated, washed twice with a buffer containing DPBS, 1% w/v BSA, 2 mM CaCl_2_ and then suspended in 0.4 mL of the same buffer or in media when detailed. To this sample was added the specified SMART-SpyCatcher system at the stated concentration by adding each individual component (SpyN and SpyC) from stock solutions dissolved in 100 mM NaH_2_PO_4_ (pH 7.2), 150 mM NaCl, 1 mM ethylenediaminetetraacetic acid (EDTA), 1 mM DTT. The sample was then incubated for 2 hr at 37 °C, 5% CO_2_ to allow for cell surface receptor binding and protein *trans*-splicing. The specified SpyTag003 conjugate was then added (100 nM final concentration) and the SpyTag003-SpyCatcher003 reaction allowed to proceed in the dark at room temperature for 20 mins. The cells were then washed twice with cold DPBS, 1% w/v BSA, 2 mM CaCl_2_ and incubated on ice until further analysis by confocal microscopy or flow-cytometry. Samples subjected to SDS-PAGE/Western blotting were washed twice with DPBS alone to remove excess BSA. Control experiments examining the mechanism of action in Extended Data Fig. 3a-c included the following steps prior to the addition of SpyTag003-AF594: 1) addition of competing DARPins (500 nM) blocking SMART-SpyCatcher binding, 2) addition of Cys1-alkylated eNrdJ-1N^cage^ component unable to perform splicing, or 3) pre-addition of unlabeled SpyTag003 (500 nM) to block reaction with SpyTag003-AF594. In reactions with SpyTag003^D117A^-AF594, the mutation D117A disables the formation of an isopeptide bond with SpyCatcher003.

#### Adherent cell lines

Individually cultured adherent cell lines were lifted using trypsin, washed with complete medium and counted to estimate the cell concentration. 200,000 cells were then seeded in 24 well plates and allowed to attach and recover for 24 hr in complete media. The assay protocol for SMART-SpyCatcher actuation and SpyTag003 recruitment followed that outlined above for the suspension cell lines. Post-treatment, the adherent cells were washed twice with DPBS, 1% w/v BSA, 2 mM CaCl_2_ and then lifted with a non-enzymatic solution of DPBS, 1% w/v BSA, 50 mM EDTA and incubated on ice before analysis by flow-cytometry. For SDS-PAGE/Western blotting cells were lifted as described and then washed with DPBS alone to remove excess BSA.

### Mixed-population experiments using SMART-SpyCatcher

#### Suspension cell lines

Two distinct mixed populations of K562 cell lines were generated (K562 expresses low endogenous levels of EpCAM):

- Mixed K562 population 1

`o` K562 (wildtype cell line)
`o` _K562EGFR+_
`o` _K562HER2+_
`o` _K562HER2+/EGFR+_
- Mixed K562 population 2

`o` K562 (wildtype cell line)
`o` _K562EGFR+_
`o` _K562HER2+/EpCAMhi_
`o` _K562HER2+/EGFR+/EpCAMhi_

The mixed populations were generated by combining the specified four cell lines (50,000 each) in a reaction tube. The SMART-SpyCatcher system (SpyN and SpyC) was then added at the specified concentration and the sample incubated for 2 hr, 37 °C, 5% CO_2_ to allow for cell surface receptor binding and protein *trans*-splicing. In experiments involving NOT gating, the decoy was added at 100 nM just before the addition of SMART-SpyCatcher. The required SpyTag003 conjugate was then added (100 nM final concentration) and the SpyTag003-SpyCatcher003 reaction was allowed to proceed in the dark at room temperature for 20 mins. The cells were then washed twice with cold DPBS, 1% w/v BSA, 2 mM CaCl_2_ and incubated on ice until further analysis by confocal microscopy or flow-cytometry. When appropriate for data representation, the individual data sets from the two distinct mixed populations were combined into one common bar graph.

#### Adherent cell lines

The three adherent cell lines MCF-10a, MCF7, and Sk-br-3 were cultured individually. Each cell line was lifted with a non-enzymatic solution of DPBS, 1% w/v BSA, 50 mM EDTA and counted to estimate the cell concentrations. MCF-10a was then mixed in equal numbers with either MCF-7 or Sk-br-3, and the two mixed mammary populations washed with either complete DMEM medium (MCF-10a + MCF-7) or complete McCoy’s 5a Medium Modified medium (MCF-10a + Sk-br-3), before being aliquoted at 200,000 cells/well in a 24 well plate. The cells were incubated for 6 hours at 37 °C, 5% CO_2_. The assay protocol for SMART-SpyCatcher actuation and SpyTag003 recruitment follows that outlined above for the suspension cell lines. Post-treatment, the adherent cells were washed twice with DPBS, 1% w/v BSA, 2 mM CaCl_2_ and then lifted with a non-enzymatic solution of DPBS, 1% w/v BSA, 50 mM EDTA and incubated on ice before analysis by flow-cytometry.

### Cell phenotyping

#### Suspension cell lines

Individually cultured K562 cell lines were spun down and counted to estimate the cell concentration. Then approximately 200,000 cells of an individual K562 cell culture were isolated, washed with DBPS, 1% w/v BSA, 2 mM CaCl_2_ and incubated 30 min at room temperature in the same buffer supplemented with 100 nM Alexa Fluor 594 DARPin conjugate targeting the antigen EpCAM. The cells were washed twice with DBPS, 1% w/v BSA, 2 mM CaCl_2_ and incubated on ice until further analysis by flow-cytometry.

#### Adherent cell lines

Plated cells were lifted with trypsin, washed with complete medium and counted to estimate the cell concentration. Then 200,000 cells were seeded in a 24 well plate format and allowed to recover and attach for 24 hr in complete media. The cells were then washed with DBPS, 1% w/v BSA, 2 mM CaCl_2_ and incubated 30 min at room temperature in the same buffer supplemented with 100 nM Alexa Fluor 594 DARPin conjugate targeting either of the three antigens HER2, EGFR, or EpCAM. The cells were washed twice with DBPS, 1% w/v BSA, 2 mM CaCl_2_ before being lifted with a non-enzymatic solution of DPBS, 1% w/v BSA, 50 mM EDTA and incubated on ice until further analysis by flow-cytometry.

### Anti-DNP monoclonal IgG recruitment assay

The cell sample was prepared following the assay protocol detailed for experiments involving a single population or mixed populations of suspension cells as described above. Following incubation with SMART-SpyCatcher (SpyN and SpyC each at 100 nM final concentration) for 2 hr, SpyTag003 labeled with 2,4-dinitrophenol (SpyTag003-DNP, 100 nM) was added to the cells and the SpyTag003-SpyCatcher003 reaction allowed to proceed for 20 minutes at room temperature in the dark. The cells were then washed twice with DPBS, 1% w/v BSA, 2 mM CaCl_2_ and then incubated with primary anti-DNP IgG antibody (1:1000) for 30 min at room temperature. Excess antibody was washed away with DBPS, 1% w/v BSA, 2 mM CaCl_2_ and the cells then incubated with excess secondary Goat anti-rabbit IgG antibody Alexa Fluor 594 conjugate (1:4000) for 30 minutes. The cells were then washed with DBPS, 1% w/v BSA, 2 mM CaCl_2_ before being imaged by confocal microscopy or analyzed by flow cytometry. Control experiments validating that anti-DNP IgG engagement was caused by SpyTag003-DNP conjugation to the cell surface included the following steps prior to addition of SpyTag003-DNP: by 1) addition of competing DARPins (500 nM) blocking SMART-SpyCatcher binding, 2) addition of Cys1-alkylated eNrdJ-1N^cage^ component unable to perform splicing, or 3) addition of unlabeled SpyTag003 (500 nM) to block reaction with the SpyTag003-DNP variant.

### NeutrAvidin Rhodamine Red-X recruitment assay

The cell sample was prepared following the assay protocol detailed for single population or mixed populations of suspension cells described above. Following incubation with SMART-SpyCatcher (SpyN and SpyC each at 100 nM final concentration) for 2 hr, SpyTag003 labeled with biotin (SpyTag003-Biotin, 100 nM) was added to the cells and the SpyTag003-SpyCatcher003 reaction allowed to proceed for 20 minutes at room temperature in the dark. The cells were then washed twice with cold DPBS, 1% w/v BSA, 2 mM CaCl_2_ and then incubated with NeutrAvidin Rhodamine Red-X (1:500) for 30 min. The cells were then washed with DBPS, 1% w/v BSA, 2 mM CaCl_2_ before being imaged by confocal microscopy or analyzed by flow cytometry. For the experiment studying the intake of NeutrAvidin Rhodamine Red-X the sample was imaged immediately and after an additional incubation period of 4 hr at the specified temperature.

### Cell depletion assay

#### Mixed K562 population depletion

The sample was prepared following the assay protocol detailed for mixed populations of suspension cells as described above. Following incubation with SMART-SpyCatcher (SpyN and SpyC each at 100 nM final concentration) for 2 hr, SpyTag003 labeled with biotin (SpyTag003-Biotin, 100 nM) was added to the cells and the SpyTag003-SpyCatcher003 reaction allowed to proceed for 30 minutes at room temperature. The cells were washed with complete RPMI-1640 medium. Cells were then aliquoted into a 96 well plate wells (12,500 cells/well) and an additional 100 μL complete RPMI-1640 media containing Streptavidin-Saporin (20 nM final concentration) was added. The cells were then cultured for 72 hr for the single dose regimen. For the two-dose regiment the cells were cultured for 24 hr before being subjected to the treatment described above a second time, and then cultured for an additional 72 hr. Samples were then analyzed by flow cytometry.

Subpopulation percentages obtained from the flow cytometry analysis were normalized using the following formula:

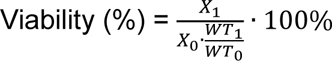

where *X_0_* is the percentage of the specific subpopulation in the negative untreated sample, *X_1_* is the percentage of the specific subpopulation the relevant sample, *WT_0_* is the percentage of the wildtype subpopulation in the negative untreated sample, and *WT_1_* is the percentage of the wildtype subpopulation in the relevant sample. The derived percentage is taken as the viability of the specified subpopulation.

#### Single A431 population depletion

For the A431 cell depletion assay, the cultured cells were lifted with trypsin, counted, and then seeded at 5,000 cells per well in a 96 well plate before further culturing for 24 hr at 37 °C, 5% CO_2_. The cells were then washed with DPBS, 1% w/v BSA, 2 mM CaCl_2_ and incubated with the indicated SMART-SpyCatcher (SpyN and SpyC each at 100 nM final concentration) for 2 hr at 37 °C, 5% CO_2_. SpyTag003-Biotin (100 nM) was added, and the sample incubated for 30 min at room temperature. The cells were gently washed with complete DMEM medium before being suspended in 200 μL DMEM medium containing 20 nM Streptavidin-Saporin conjugate as indicated. The cells were cultured for 24 hr at 37 °C, 5% CO_2_, before the complete treatment described above was repeated to give a two-dose regimen. Finally, the cells were cultured for an additional 72 hr. Each well was then washed twice with complete DMEM medium and 200 μL complete DMEM medium added. An XTT cell viability assay was used to quantify the metabolic activity of each sample according to the protocol prescribed by the manufacturer. The conversion of XTT to formazan was measured on a SpectraMax® iD5 Multi-Mode Microplate Reader (Molecular Devices) using well-scan mode at 465 nm at room temperature. The obtained absorbance values were normalized to the negative untreated control.

### APEX2 proximity labeling

The cell sample, prepared following the assay protocol detailed for single populations, was treated with the SMART-SpyCatcher reactants (SpyN and SpyC each at 100 nM final concentration) for 2 hr. The cells were then incubated with SpyTag003-APEX2 (300 nM) for 30 minutes at room temperature before being washed twice with DPBS. To initiate APEX proximity labeling, the cells were treated with 1 mL DPBS containing biotin-phenol (250 µM final concentration) followed by the addition of 10 µL freshly prepared DPBS containing 100 mM H_2_O_2_, and the reaction allowed to proceed at room temperature under gentle swirling for the indicated time. The reaction was quenched by the addition of 200 µL quencher buffer (DPBS, 10 mM sodium ascorbate, 5 mM Trolox) and the cells further washed twice with 1 mL of the quencher buffer. The cells were lifted (when necessary) by the addition of 1 mL DPBS, 50 mM EDTA, pelleted and suspended in 200 µL–500 µL lysis buffer (50 mM Tris-HCl (pH 8.0), 150 mM NaCl, 1% v/v NP-40, 0.5% v/v sodium deoxycholate, 0.1% w/v sodium dodecyl sulfate supplemented with 1x Halt™ Protease Inhibitor Cocktail and 1 mM PMSF), then sonicated briefly and centrifuged for 20 min at 15,000 g, 4 °C. The supernatant was isolated and analyzed by SDS-PAGE/Western blotting.

### Statistics and Reproducibility

All statistical analyses were conducted in GraphPad Prism v.9.2.0. P values were determined by either paired t-test or one-way ANOVA followed by Dunnett’s test as listed in the figure legends. The statistical significances of differences (ns denotes not significant; * denotes P < 0.05, ** denotes P < 0.01, *** denotes P < 0.001, **** denotes P < 0.0001) are specified throughout the figures and legends. Heatmaps were generated using GraphPad Prism v.9.2.0. All experiments analyzed by SDS-PAGE/Western blotting and/or flow cytometry were repeated at least three times (independent biological replicates).

---

**Extended Data Figure 1.**
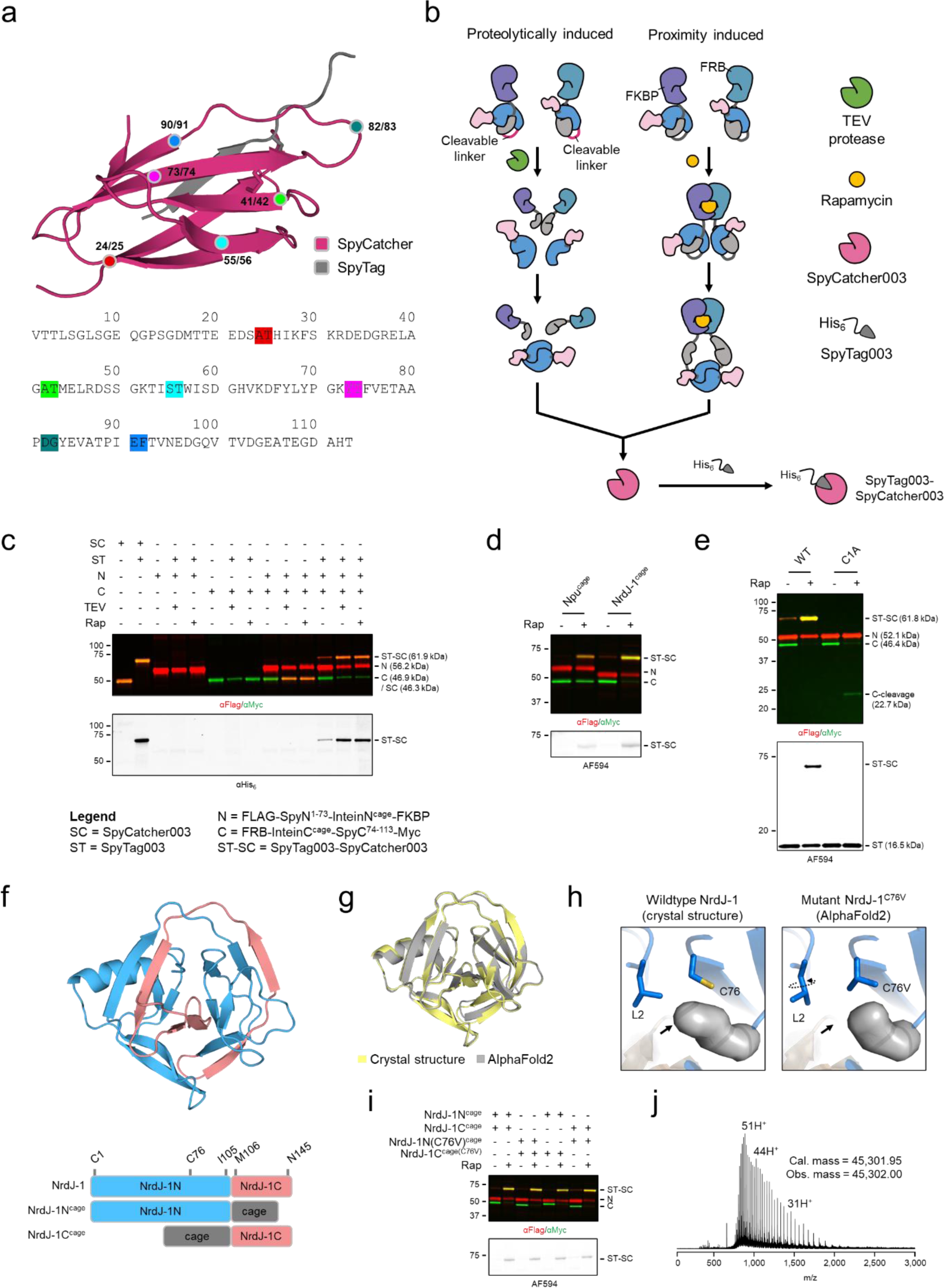
Systematic screen to identify SpyCatcher003 split site and optimization of SMART-SpyCatcher. **a**, SpyCatcher003 was split at various sites and the cognate pairs used to generate FLAG-SpyN^1-x^-NpuN^cage^-FKBP and FRB-NpuC^cage^-SpyC^y-113^–Myc, with x and y denoting the last and first residue of the two fragments (i.e., the split site). The isopeptide bond (formed between D117 of SpyTag003 and K31 of SpyCatcher003) is shown in sticks in the structure of SpyTag-SpyCatcher (PDB ID: 4MLI). The primary structure of SpyCatcher003 is shown with split sites indicated. **b**, Schematic of the cell-free *in vitro* screen used to identify the optimal split site. Conditional protein splicing was induced either by proteolytic decaging using TEV protease or through chemically induced FKBP-rapamycin-FRB heterodimerization, which simulates ideal colocalization on a target cell surface. **c**, The result of screening FLAG-SpyN^1–74^-NpuN^cage^-FKBP and FRB-NpuC^cage^-SpyC^74–113^–Myc generated from splitting SpyCatcher003 at position 73-74. Reactions were performed with FLAG-SpyN^1–74^-NpuN^cage^-FKBP (1 μM), FRB-NpuC^cage^-SpyC^74–113^–Myc, (1 μM) and His_6_-SpyTag003 (2 μM). CPS was induced by the addition of either TEV protease (10 units) or rapamycin (10 μM). The reactions were analyzed by Western blot after 24 hr incubation at 37 °C. FLAG-SpyCatcher003-Myc was used as a size standard for the spliced product; FLAG-SpyCatcher003-Myc (1 μM) reacted with His_6_-SpyTag003 (2 μM) was used as a size standard for the covalent complex between the two. The legend and experimental conditions apply for subsequent panels with alterations when noted. **d**, Npu^cage^ was swapped with NrdJ-1^cage^ in SMART-SpyCatcher thereby giving FLAG-SpyN^1–73^-NrdJ-1N^cage^-FKBP and FRB-NrdJ-1C^cage^-SpyC^74–113^–Myc. A SpyTag003 Alexa Fluor 594 conjugate (SpyTag003-AF594) was used as the activity probe. The reactions were analyzed by Western blot. **e**, SpyN^1–73^-NrdJ-1N^cage^-FKBP with the WT or a catalytically dead C1A variant of NrdJ-1N^cage^ was mixed with FRB-NrdJ-1C^cage^-SpyC^74–113^–Myc and SpyTag003-AF594. The reactions were analyzed by Western blot. **f**, The crystal structure of fusion NrdJ-1 (PDB ID: 8UBS), with sequences corresponding to NrdJ-1N and NrdJ-1C colored in light blue and light red respectively. A schematic representation of the fusion protein and the caged split intein is shown as well to indicate relevant domains in addition to their N- and C-termini and the position of the non-catalytic Cys76. **g-h**, A structural analysis was performed to predict the impact of introducing a C76V mutation in fusion NrdJ-1. **g**, The backbone of wildtype NrdJ-1 (crystal structure) and that of the NrdJ-1^C76V^ mutant (*in silico* AlphaFold2 model) are superimposable with a mean square deviation of 0.56 Å. **h**, Residue sidechains of Cys76 and Leu2 are shown as sticks and cavities in solid grey. The experimentally determined local environment of Cys76 is shown on the left. The *in silico* predicted local environment of C76V is shown on the right. Changes are indicated with arrows. **i**, Combinations of FLAG-SpyN^1–73^-NrdJ-1N^cage^-FKBP or FLAG-SpyN^1–74^-NrdJ-1NC76V^cage^-FKBP with FRB-NrdJ-1C^cage^-SpyC^74–113^–Myc or FRB-NrdJ-1C^cage^(C76V)-SpyC^74–113^–Myc were tested as indicated using the reaction conditions described above. The reactions were analyzed by Western blot. **j**, ESI-TOF MS characterization of the SpyCatcher003 spliced product produced by rapamycin triggered CPS between FLAG-SpyN^1–74^-NrdJ-1NC76V^cage^-FKBP and FRB-NrdJ-1C^cage^(C76V)-SpyC^74–113^–Myc. Reaction conditions followed those described above.

**Extended Data Figure 2.**
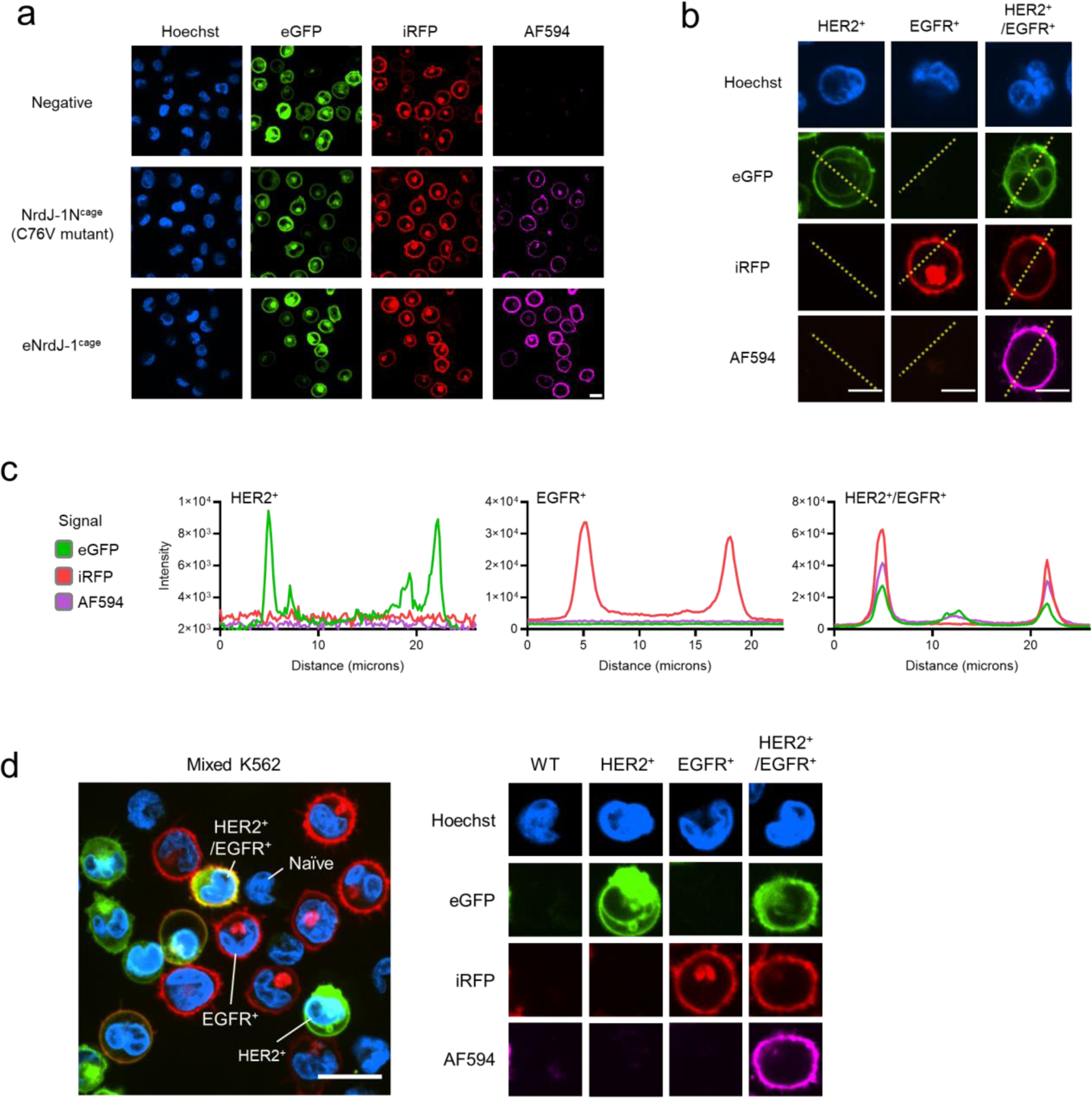
Confocal microscopy of K562 cell lines treated with SMART-SpyCatcher for [HER2 AND EGFR] logic. **a**, K562^HER2+/EGFR+^ cells were treated with αHER2-SpyN (100nM) and SpyC-αEGFR (100 nM) for 2 hr, followed by SpyTag003 labeled with Alexa Fluor 594 (SpyTag003-AF594, 100 nM) for 20 mins. SMART-SpyCatcher employed either NrdJ-1N(C76V)^cage^/NrdJ-1C^cage^(C76V) or NrdJ-1N(C76V)^cage^(K104EK119A)/NrdJ-1C^cage^(C76VD66K) (referred to as eNrdJ-1^cage^). Following washing, the live cells were then analyzed by confocal microscopy. Cell nuclei were stained with Hoechst, while HER2 and EGFR were tagged with eGFP and iRFP, respectively. Scale bar equals 20 μm. All subsequent panels apply similar reaction and analysis conditions as in panel **a** using the SMART-SpyCatcher (eNrdJ-1^cage^) [HER2 AND EGFR] system. **b**, K562^HER2+^, K562^EGFR+^, and K562^HER2+/EGFR+^ cells were treated and analyzed individually. Scale bars equal 10 µm. **c**, To determine the colocalization of HER2, EGFR and SpyTag003-AF594 the signal intensities of eGFP, iRFP, and AF594 derived from the cells in panel **b** were plotted (yellow dotted lines). **d**, A mixed population consisting of equal amounts of K562 (wildtype), K562^HER2+^, K562^EGFR+^, and K562^HER2+/EGFR+^ cells were treated and imaged. Representative single cells of each cell line from the treated mixture and their associated fluorescence signals are shown on the right. Scale bar equals 20 μm.

**Extended Data Figure 3.**
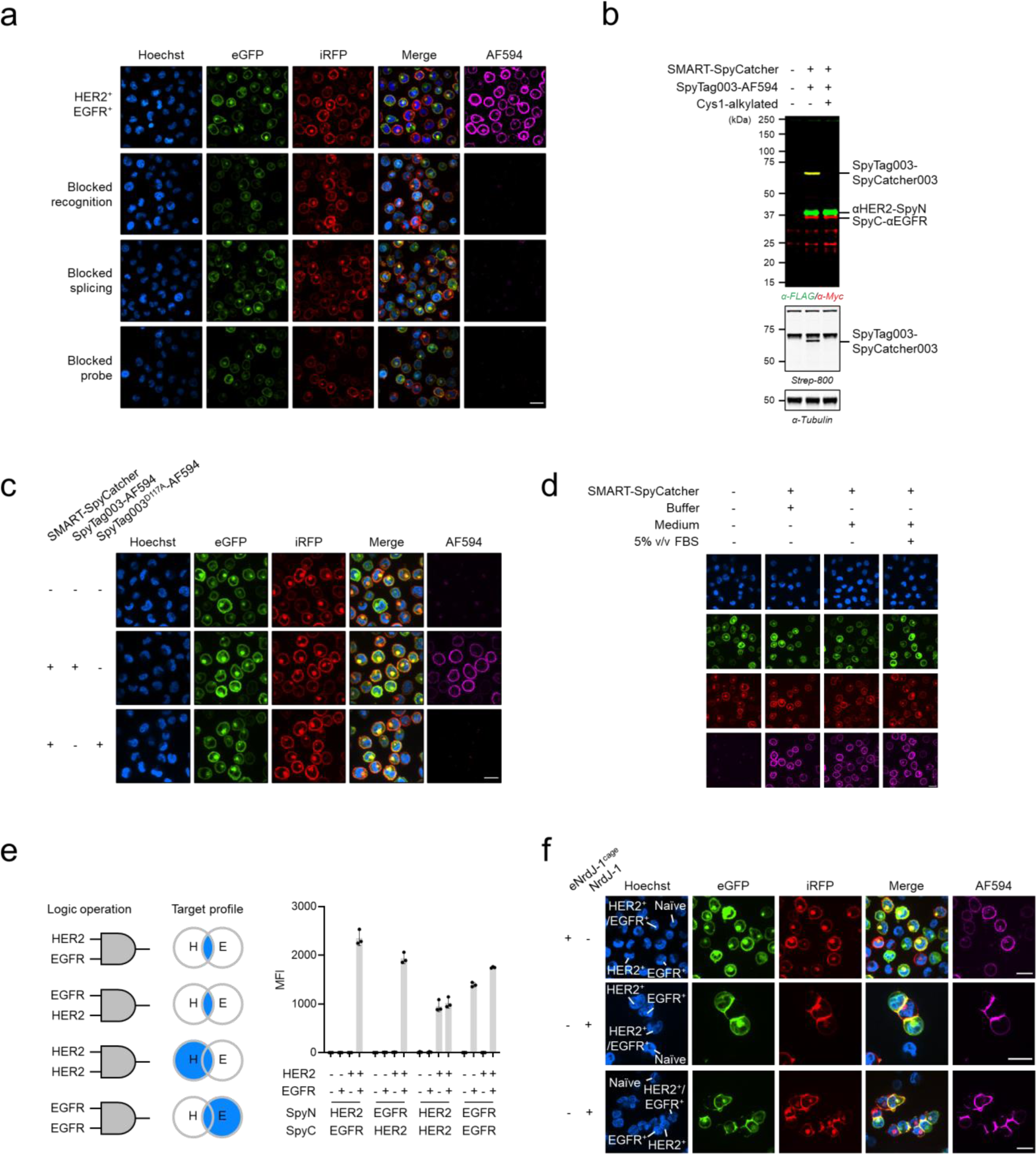
The mechanism of action of SMART-SpyCatcher. **a**, K562^HER2+/EGFR+^ cells were treated with αHER2-SpyN (100nM) and SpyC-αEGFR (100 nM) for 2 hr, followed by SpyTag003 labeled with Alexa Fluor 594 (SpyTag003-AF594, 100 nM) for 20 mins. SMART-SpyCatcher employed eNrdJ-1^cage^. Following washing, the live cells were then analyzed by confocal microscopy. Cell nuclei were stained with Hoechst, while HER2 and EGFR were tagged with eGFP and iRFP, respectively. The cells in row 1 was treated as described above, whereas reaction conditions in the subsequent experiments were supplemented with DARPins targeting HER2 and EGFR (500 nM each, row 2), used an inactivated version of NrdJ-1N^cage^ (Cys1-alkylated, row 3), or was supplemented with unlabeled SpyTag003 (500 nM, row 4). Scale bar equals 20 μm. All subsequent panels apply similar reaction and analysis conditions as in panel **a** (row 1) using the SMART-SpyCatcher (eNrdJ-1^cage^) system with any alterations as noted. **b**, Western-blot analysis of K562^HER2+/EGFR+^ cells treated with αHER2-SpyN and SpyC-αEGFR (identified by FLAG and Myc respectively), and SpyTag003 labeled with biotin (SpyTag003-biotin; identified by Strep-800). **c**, Experiments were performed on K562^HER2+/EGFR+^ cells with the SMART-SpyCatcher (eNrdJ-1^cage^) [HER2 AND EGFR] system, and either SpyTag003-AF594 or inactive SpyTag003^D117A^ labeled with Alexa Fluor 594 (SpyTag003^D117A^-AF594). Scale bar equals 20 μm. **d**, K562^HER2+/EGFR+^ cells were treated with 100 nM SMART-SpyCatcher (eNrdJ-1^cage^) in buffer (DPBS, 1% w/v BSA, 2 mM CaCl_2_; medium), RPMI 1640 medium (cystine-free), or in medium supplemented with 5% v/v fetal bovine serum (5% v/v FBS) for 2 hr, followed by SpyTag003-AF594 for 20 min. Further image analysis was performed as described above. Scale bar equals 20 μm. **e**, Flow cytometry analysis of a mixed population consisting of equal amounts of K562 (wildtype), K562^HER2+^, K562^EGFR+^, and K562^HER2+/EGFR+^ cells following treatment with SMART-SpyCatcher (eNrdJ-1^cage^) operating through the AND logic indicated. The added SpyN and SpyC pairs (i.e., the targeting DARPins employed in the constructs) are indicated at the bottom. Data are presented as the mean of the AF594 median fluorescence intensities (MFI) from flow cytometry analysis with error bars signifying the standard error mean (n = 3 independent biological replicates; see Supplementary Table 3 for statistical one-way ANOVA followed by Dunnett’s test). **f**, The mixed K562 population described above were treated with αHER2-SpyN (100nM) and SpyC-αEGFR (100 nM) employing eNrdJ-1^cage^ or the uncaged split intein NrdJ-1, for 2 hr, followed by SpyTag003-AF594 (100 nM) for 20 min. Further image analysis was performed as described above. Scale bars equal 20 μm.

**Extended Data Figure 4.**
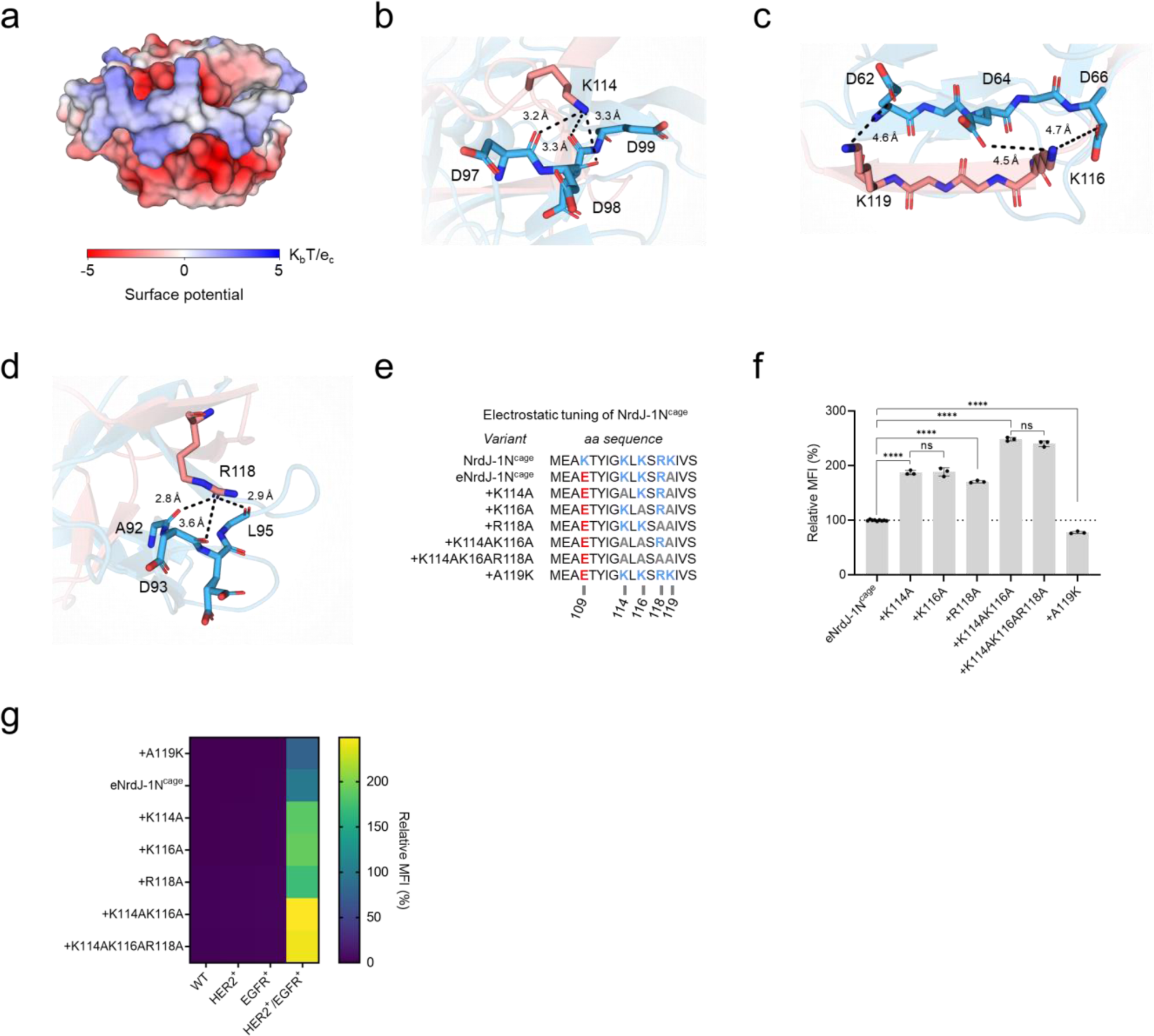
Structural analysis of NrdJ-1 and electrostatic tuning of eNrdJ-1N^cage^. **a**-**e**, Summary of the electrostatic interactions involved in the fragment-fragment association between the split intein fragments NrdJ-1N and NrdJ-1C. **a**, Surface potential map calculated using the Adaptive Poisson-Boltzmann Solver (PyMOL) using the crystal structure of fused NrdJ-1 as the template. **b**-**d**, The residues involved in potential electrostatic and H-bonding interactions are represented (light blue for NrdJ-1N and light red for NrdJ-1C). The distances between indicated heavy atoms are shown. **e**, The N-terminal amino acid sequence of the various cages employed with NrdJ-1N. **f**-**g**, Flow cytometry analysis of a mixed population consisting of equal amounts of K562 (wildtype), K562^HER2+^, K562^EGFR+^, and K562^HER2+/EGFR+^ cells following treatment with αHER2-SpyN (using the indicated eNrdJ-1N^cage^ variants at 100 nM), SpyC-αEGFR (using the standard eNrdJ-1C^cage^ variant at 100 nM), and SpyTag003-AF594 (100 nM). Data are presented as the mean of the AF594 median fluorescence intensities (MFI) normalized to that measured for the K562^HER2+/EGFR+^ cells treated with SMART-SpyCatcher employing the standard version of eNrdJ-1 with error bars signifying the standard error mean (n = 3 independent biological replicates). **f**, Statistical significance was evaluated using a paired t-test (ns denotes not significant; * denotes P < 0.05; ** denotes P < 0.01; *** denotes P < 0.001; **** denotes P < 0.0001). **g**, The normalized AF594 intensities associated with the individual four cell lines across the different experiments is summarized in the form of a heatmap (see Supplementary Table 4 for individual values and Supplementary Table 3 for statistical one-way ANOVA followed by Dunnett’s test).

**Extended Data Figure 5.**
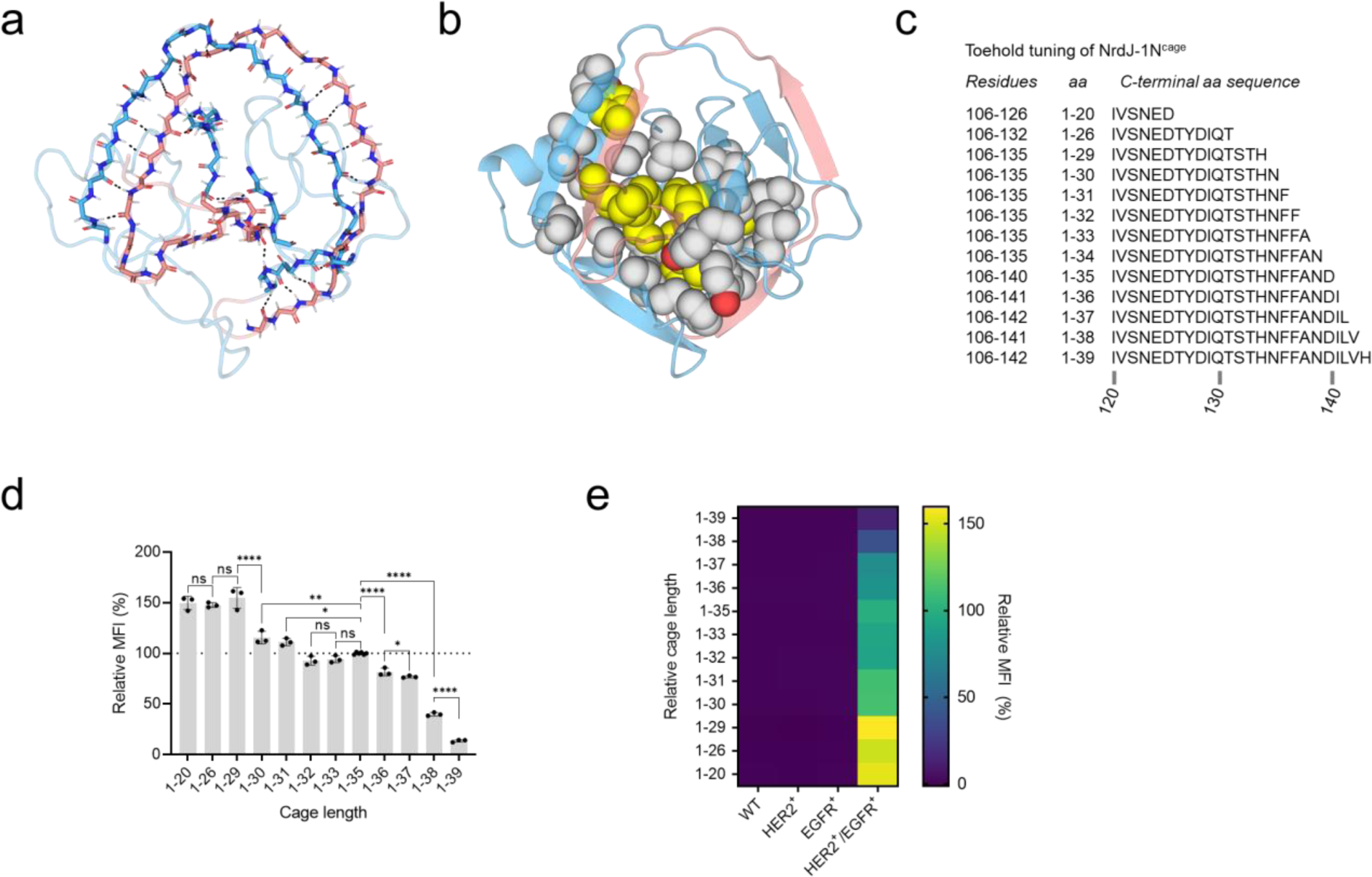
Structural analysis of NrdJ-1 and cage toehold tuning of eNrdJ-1N^cage^. **a**-**b**, The backbone of fused NrdJ-1 intertwine to form a horseshoe-like fold stabilized by **a**, β-strand hydrogen bonds and **b**, hydrophobic packing (grey and yellow space fill indicate residues of NrdJ-1N and NrdJ-1C respectively). Domains corresponding to NrdJ-1N and NrdJ-1C in light blue and light red respectively. **c**, The C-terminal amino acid sequences of the various truncated cages employed with NrdJ-1N. For reference, eNrdJ-1N^cage^ is denoted as 1-35. **d**-**e**, Flow cytometry analysis of a mixed population consisting of equal amounts of K562 (wildtype), K562^HER2+^, K562^EGFR+^, and K562^HER2+/EGFR+^ cells) following treatment with αHER2-SpyN (using the indicated eNrdJ-1N^cage^ variants at 100 nM), SpyC-αEGFR (using the standard eNrdJ-1C^cage^ variant at 100 nM), and SpyTag003-AF594 (100 nM). Data are presented as the mean of the AF594 median fluorescence intensities (MFI) normalized to that measured for the K562^HER2+/EGFR+^ cells treated with SMART-SpyCatcher employing the standard version of eNrdJ-1 with error bars signifying the standard error mean (n = 3 independent biological replicates). Statistical significance was evaluated using a paired t-test (ns denotes not significant; * denotes P < 0.05; ** denotes P < 0.01; *** denotes P < 0.001; **** denotes P < 0.0001). **e**, The normalized AF594 intensities associated with the individual four cell lines across the different experiments is summarized in the form of a heatmap (see Supplementary Table 5 for individual values and Supplementary Table 3 for statistical one-way ANOVA followed by Dunnett’s test).

**Extended Data Figure 6.**
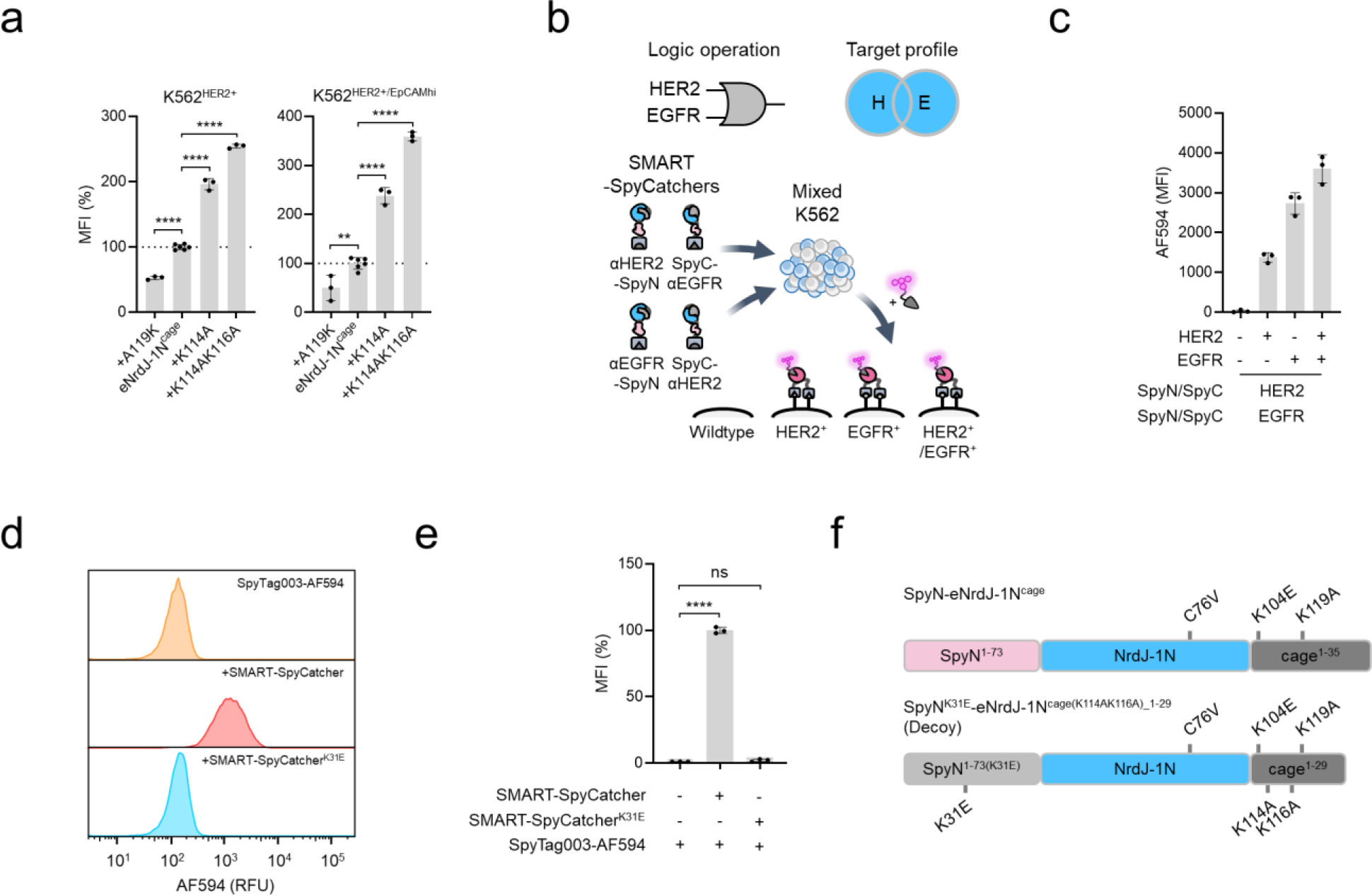
Developing three-input logics for SMART-SpyCatcher. **a**, Flow cytometry analysis of a mixed population consisting of equal amounts of K562 (wildtype), K562^HER2+^, K562^EGFR+^, and K562^HER2+/EGFR+/EpCAMhi^ cells following treatment with αHER2-SpyN (using the indicated tuned eNrdJ-1N^cage^ variant at 100 nM), SpyC-αEGFR (100 nM), and SpyTag003-AF594 (100 nM). Data are presented as the mean of the AF594 median fluorescence intensities (MFI) normalized to that measured for the K562^HER2+/EGFR+^ cells treated with SMART-SpyCatcher employing the standard version of eNrdJ-1. Error bars signify the standard error mean (n = 3 independent biological replicates). Statistical significance was evaluated using a paired t-test (ns denotes not significant; * denotes P < 0.05; ** denotes P < 0.01; *** denotes P < 0.001; **** denotes P < 0.0001). **b**, Schematic illustrating use of the SMART-SpyCatcher system operating through [HER2 OR EGFR] logic on a mixed K562 population. **c**, Flow cytometry analysis for a mixed population consisting of equal amounst of K562 (wildtype), K562^HER2+^, K562^EGFR+^, and K562^HER2+/EGFR+^ cells following treatment with αHER2-SpyN/αEGFR-SpyN/SpyC-αHER2/SpyC-αEGFR (each at 100 nM, eNrdJ-1^cage^), and SpyTag003-AF594 (100 nM). Errors = standard error mean (n = 3 independent biological replicates; see Supplementary Table 6 for statistical one-way ANOVA followed by Dunnett’s test). **d**-**e**, Flow cytometry data of K562^HER2+/EGFR+^ treated as in panel **a**, with SMART-SpyCatcher (eNrdJ-1^cage^) [HER2 AND EGFR] system using αHER2-SpyN with K31 or K31E in the N-terminal fragment of SpyCatcher003. Panel **d** gives an example of the measured AF594 signals, with the normalized MFI quantified in panel **e**. Errors = standard error mean (n = 3 independent biological replicates) with statistical significance evaluated using a paired t-test (**** denotes P < 0.0001). RFU, relative fluorescence units. **f**, Schematic illustrating the changes made to SpyN to generate the Decoy construct.

**Extended Data Fig. 7.**
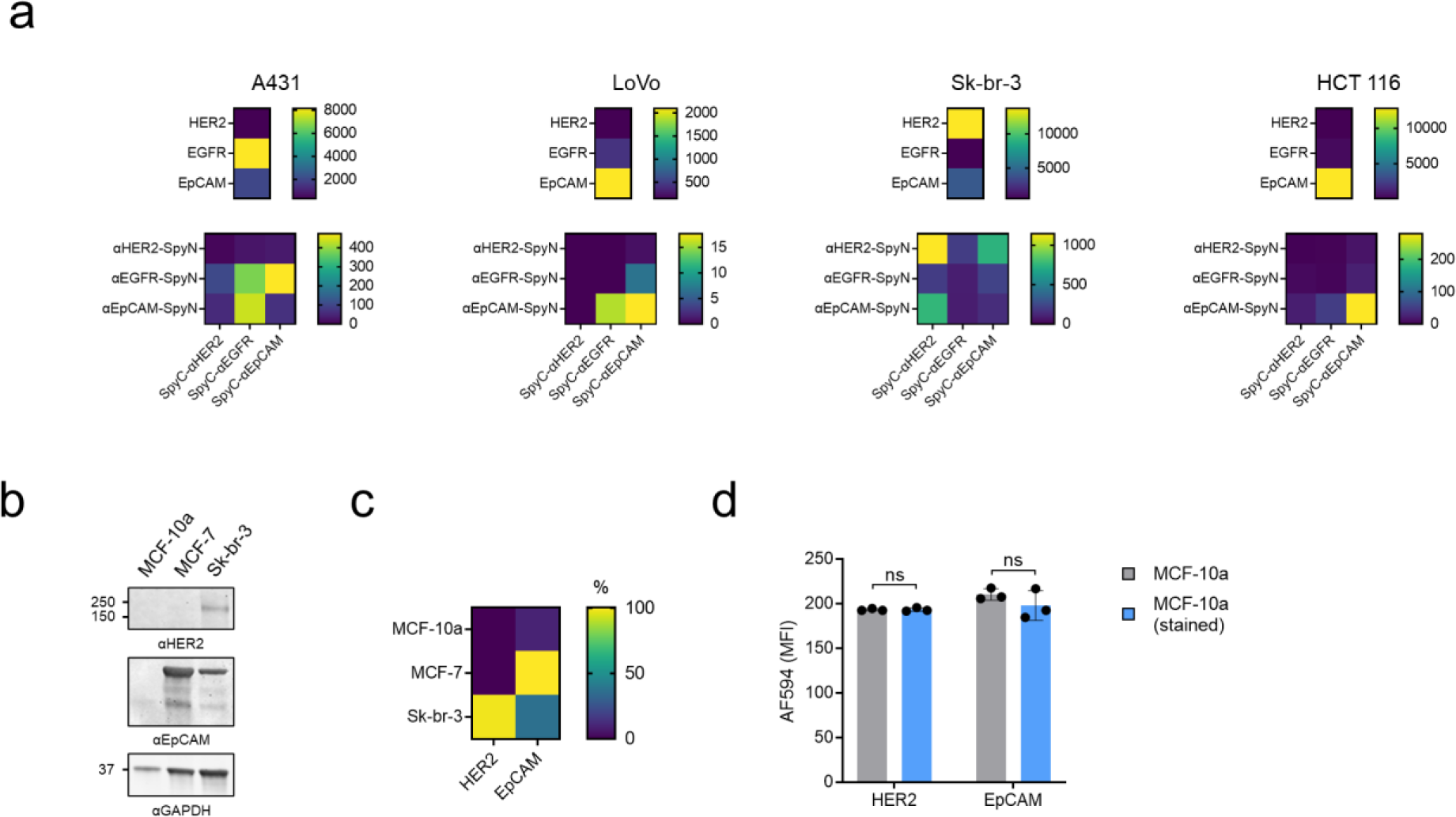
SMART-SpyCatcher uses endogenous surface antigen levels for target discrimination. **a**, Top: indicated cells were profiled for their relative relative surface levels of HER2, EGFR, and EpCAM. Cells were treated with DARPins labeled with AF594 followed by flow cytometry analysis. The results are shown in the form of heatmaps (see Supplementary Table 8 for individual values). Bottom: Cells treated with indicated SpyN and SpyC pairs (100 nM) followed by SpyTag003-AF594 (100 nM) were analyzed by flow cytometry. The results are shown in the form of heatmaps (see Supplementary Table 9 for individual values). **b**-**c**, The total cellular levels of HER2 and EpCAM for MCF-10a, MCF-7, and Sk-br-3 cell lines were detected by Western blotting. Panel **b** gives an example of the Western blot results, while panel **c** gives the quantification of the values corrected against the GAPDH signal for each cell line and further normalized to the overall highest level (Sk-br-3 for HER2, MCF-7 for EpCAM). **d**, MCF-10a cells were phenotyped as described in **a** to determine any differences in the levels of detectable surface HER2 and EpCAM as a consequence of CMFDA staining. Errors = standard error mean (n = 3 independent biological replicates). Statistical significance was evaluated using a paired t-test (ns denotes not significant).

**Extended Data Figure 8.**
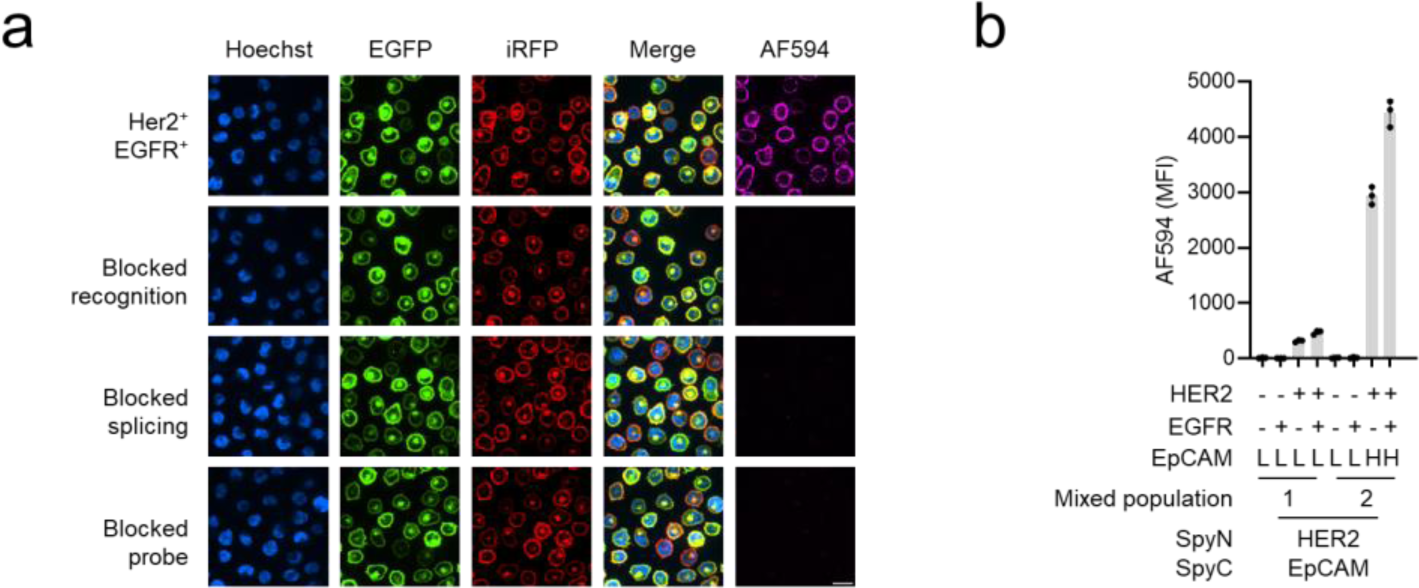
Highly selective hapten delivery and recruitment of anti-hapten antibodies. **a**, K562^HER2+/EGFR+^ cells were treated with 100 nM SMART-SpyCatcher (employing eNrdJ-1^cage^) for [HER2 AND EGFR] logic and SpyTag003 labeled with 2,4-dinitrophenol (SpyTag003-DNP, 100 nM). Following washing, the cells were treated with an anti-DNP antibody, followed by an AF594-conjugated secondary antibody, and then visualized by confocal microscopy. Cell nuclei were stained with Hoechst, while HER2 and EGFR were tagged with eGFP and iRFP, respectively. The cells in row 1 were treated as described above, whereas reaction conditions in the subsequent experiments were supplemented with DARPins targeting HER2 and EGFR (500 nM each, row 2), used an inactivated version of NrdJ-1N^cage^ (Cys1-alkylated, row 3), or were supplemented with unlabeled SpyTag003 (500 nM, row 4). Scale bar equals 20 μm. **b**, Two combinations of K562 cell lines were used to test for the actuation of SMART-SpyCatcher for [HER2 AND EpCAM] logic and recruitment of SpyTag003-DNP in a complex cell setting. Mixed-population 1 consisted of equal amounts of K562 (wildtype), K562^EGFR+^, K562^HER2+^, and K562^HER2+/EGFR+^, whereas mixed-population 2 consisted of equal amounts of K562 (wildtype), K562^EGFR+^, K562^HER2+/EpCAMhi^, and K562^HER2+/EGFR+/EpCAMhi^. The antigen profile of each cell line is indicated below each bar plot (L and H designates low endogenous and high ectopic levels respectively for EpCAM). Experiments were performed with 100 nM SMART-SpyCatcher (eNrdJ-1^cage^), 100 nM SpyTag003-AF594. Data are presented as the mean of the AF594 median fluorescence intensities (MFI) from flow cytometry analysis with error bars signifying the standard error mean (n = 3 independent biological replicates; see Supplementary Table 3 for statistical one-way ANOVA followed by Dunnett’s test).

**Extended Data Figure 9.**
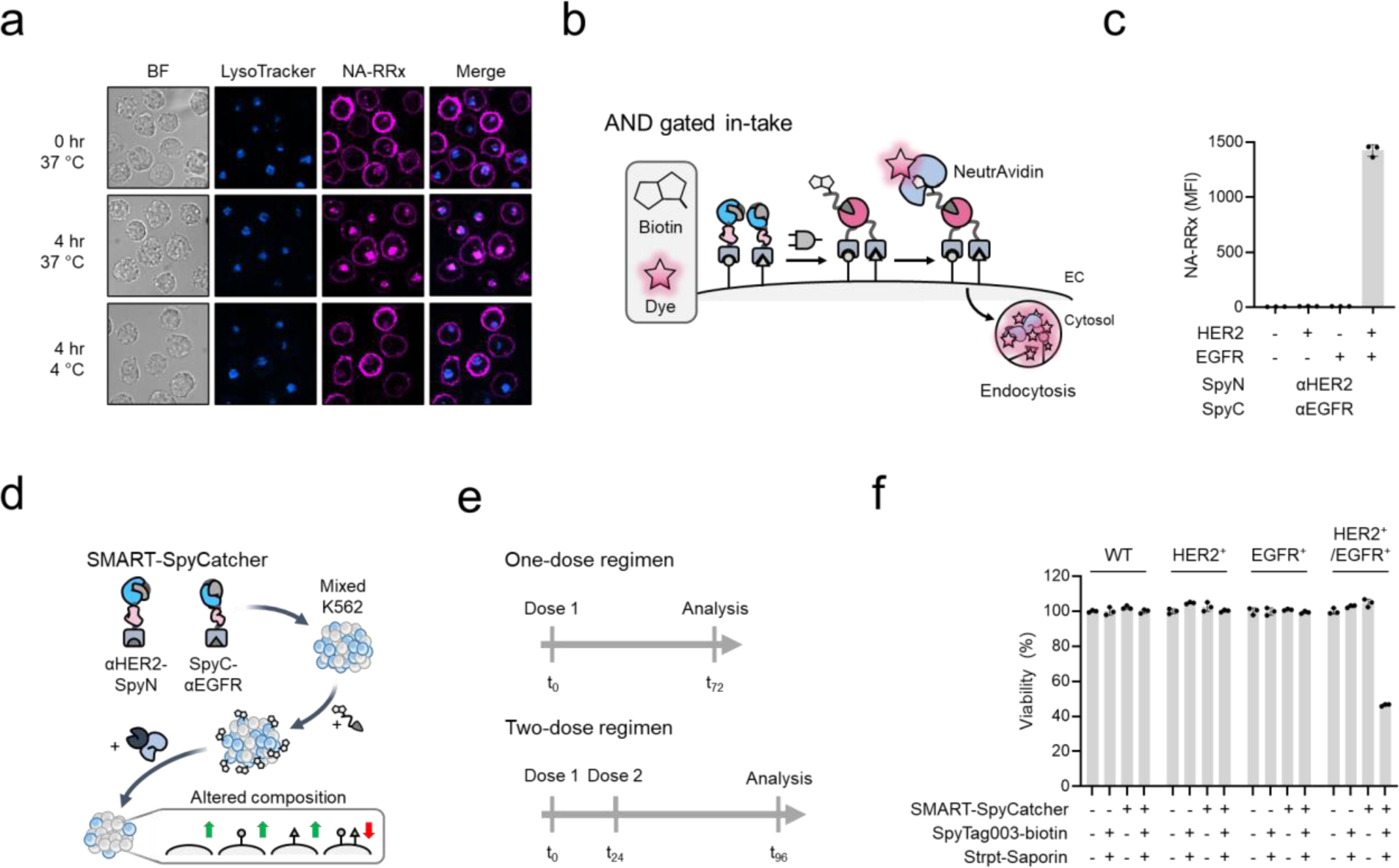
Targeted cell depletion using Boolean logic. **a**, K562^HER2+/EGFR+^ cells were treated with 100 nM SMART-SpyCatcher (employing eNrdJ-1^cage^) for [HER2 AND EGFR] logic and SpyTag003 labeled with biotin (SpyTag003-biotin, 100 nM). Following washing, the cells were treated with a NeutrAvidin Rhodamine Red-X conjugate (NA-RRx, magenta) and then either visualized by confocal microscopy immediately or after further incubation for 4 hr at 37 °C or 4 °C. Cell nuclei were stained with Hoechst, while lysosomal compartments were stained with LysoTracker. **b**, Schematic illustrating the proposed recruitment and internalization of NA-RRx enabled by the SMART-SpyCatcher system. **c**, A mixed population consisting of equal amounts of K562 (wildtype), K562^EGFR+^, K562^HER2+^, and K562^HER2+/EGFR^ cells was used to test for the selective recruitment of SpyTag003-biotin and subsequent recruitment of NA-RRx. The cell mixture was treated as in panel **a**. The NA-RRx signal associated with the individual subpopulations was quantified by flow cytometry with the data presented as the mean of the NA-RRx median fluorescence intensities (MFI) with error bars signifying the standard error mean (n = 3 independent biological replicates; see Supplementary Table 3 for statistical one-way ANOVA followed by Dunnett’s test). **d**, Schematic illustrating the selective cell depletion of a double positive cell line in a complex cell mixture using SMART-SpyCatcher, SpyTag003-biotin and a Streptavidin-Saporin disulfide conjugate. **e**, Summary of how the one-dose and two-dose regimens were performed. **f**, The mixed K562 cell population employed in panel **c** was treated with 100 nM SMART-SpyCatcher (employing eNrdJ-1^cage^ and [HER2 AND EGFR] logic), SpyTag003-biotin (100 nM), and Streptavidin-Saporin (20 nM). Cells were then analyzed by flow cytometry and the data presented as % viability relative to untreated wildtype cells (see Methods for details). Errors = standard error mean (n = 3 independent biological replicates).

**Extended Data Figure 10.**
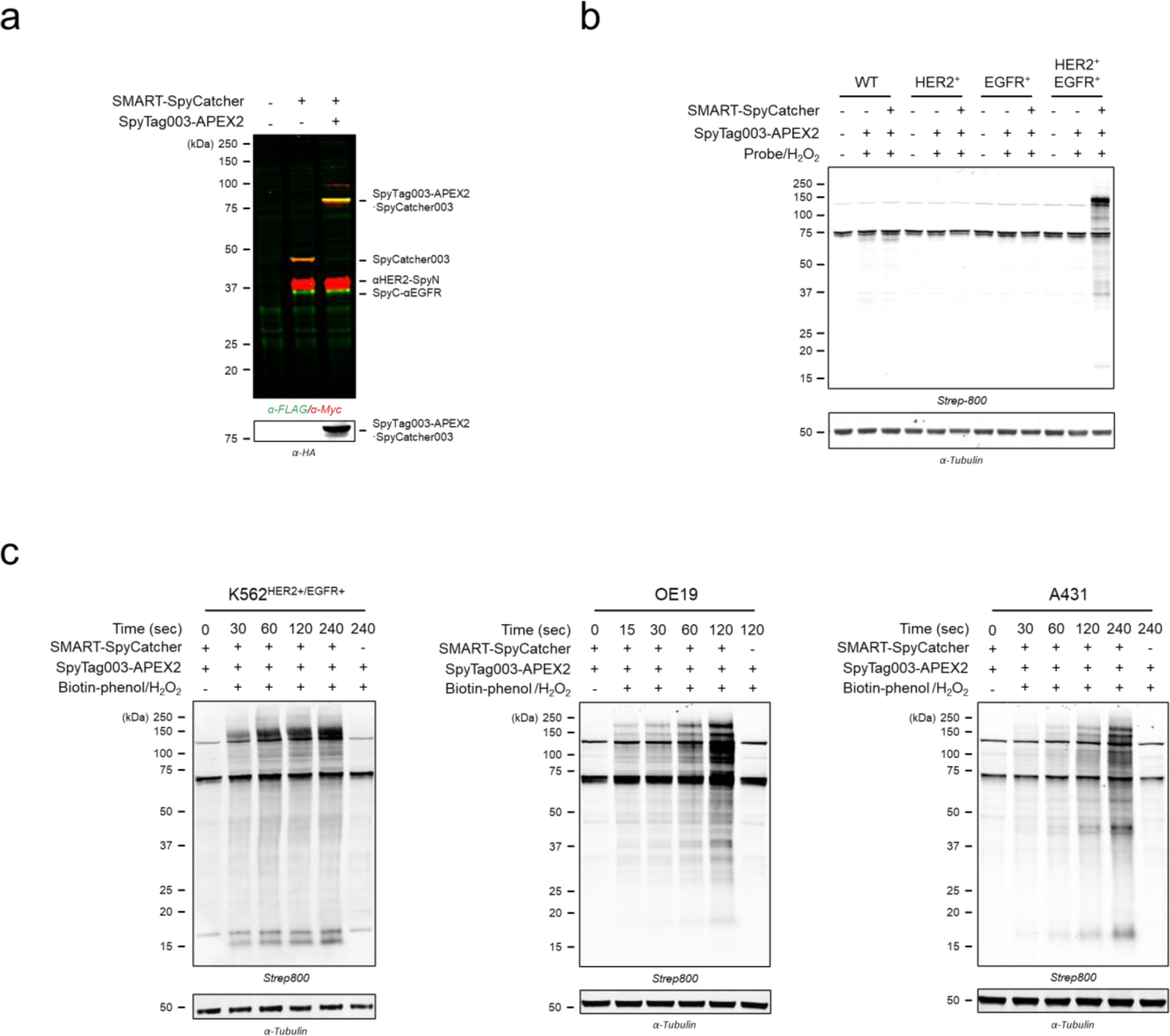
APEX2 delivery and protein proximity labeling. **a**, Western blot analysis of K562^HER2+/EGFR+^ treated with the SMART-SpyCatcher (eNrdJ-1) [HER2 AND EGFR] system for 2 hr, followed by HA-tagged SpyTag003-APEX2 for 20 min. **b**, K562 (wildtype), K562^HER2+^, K562^EGFR+^, and K562^HER2+/EGFR+^ cells were treated individually as described in panel **a**. Following washing, APEX2 proximity labeling was induced by addition of biotin-phenol and H_2_O_2_. The reactions were quenched with 10 mM sodium ascorbate and 5 mM Trolox at room temperature after two minutes. Samples were then analyzed by Western blot using Strep-800 to detect biotinylation. **c**, Time-course study of APEX2 labeling. The cells were treated as in panel **b** with labeling quenched at the indicated time points. SMART-SpyCatcher assigned for [HER2 AND EGFR] logic was used with the K562^HER2+/EGFR+^ cells; SMART-SpyCatcher assigned for [HER2 AND EpCAM] logic was used with the OE19 cells; SMART-SpyCatcher assigned for [EGFR AND EpCAM] logic was used with the A431 cells. In all cases, SMART-SpyCatcher employed eNrdJ-1.

