## Supplementary information for "Cell surface sculpting using logic-gated protein actuators"

### Table of contents

### Supplementary Figures and Tables

Supplementary Figure 1: Protein purification and characterization

Supplementary Figure 2: Conditional protein splicing of SpyN-NpuN$^{cage}$-FKBP and FRB-NpuC$^{cage}$-SpyC

Supplementary Figure 3: Preparation and characterization of the NrdJ-1 fusion

Supplementary Figure 4: Development of an optimal NrdJ-1 actuator

Supplementary Figure 5: Flow cytometry of individual K562 cell lines

Supplementary Figure 6: Flow cytometry gating strategy for mixed K562 population 1 flow cytometry experiments

Supplementary Figure 7: The elution profiles of αHER2-SpyN employing tuned eNrdJ-1Ncage variants

Supplementary Figure 8: Phenotyping K562 cell lines with high ectopic levels of EpCAM

Supplementary Figure 9: Flow cytometry gating strategy for mixed K562 population 2 flow cytometry experiments

Supplementary Figure 10: Flow cytometry of individual mammary cell lines

Supplementary Figure 11: Flow cytometry gating strategy for mixed mammary population 1 flow cytometry experiments

Supplementary Figure 12: Flow cytometry gating strategy for mixed mammary population 2 flow cytometry experiments

Supplementary Figure 13: K562 depletion experiments with a one-dose regimen

Supplementary Figure 14: K562 depletion experiments with a two-dose regimen

Supplementary Table 1: Proteins used in this study

Supplementary Table 2: Crystallization data collection and refinement statistics

Supplementary Table 3: Statistical analysis of flow cytometry data for AND gate experiments

Supplementary Table 4: Flow cytometry data for electrostatically tuned eNrdJ-1cage

Supplementary Table 5: Flow cytometry data for toehold tuned eNrdJ-1cage

Supplementary Table 6: Statistical analysis of flow cytometry data for OR gate experiments

Supplementary Table 7: Statistical analysis of flow cytometry data for AND/OR gate experiments

Supplementary Table 8: Flow cytometry data for antigen phenotyping

Supplementary Table 9: Flow cytometry data for SpyTag003-AF594 recruitment

Supplementary Table 10: Primary structures of expressed proteins

Supplementary Table 11: Antibodies used for in this study

Supplementary Table 12: Mammalian cell culture media conditions

### Supplementary Synthetic Methods

Synthesis of DNP-maleimide

Synthesis of biotin-phenol

### References

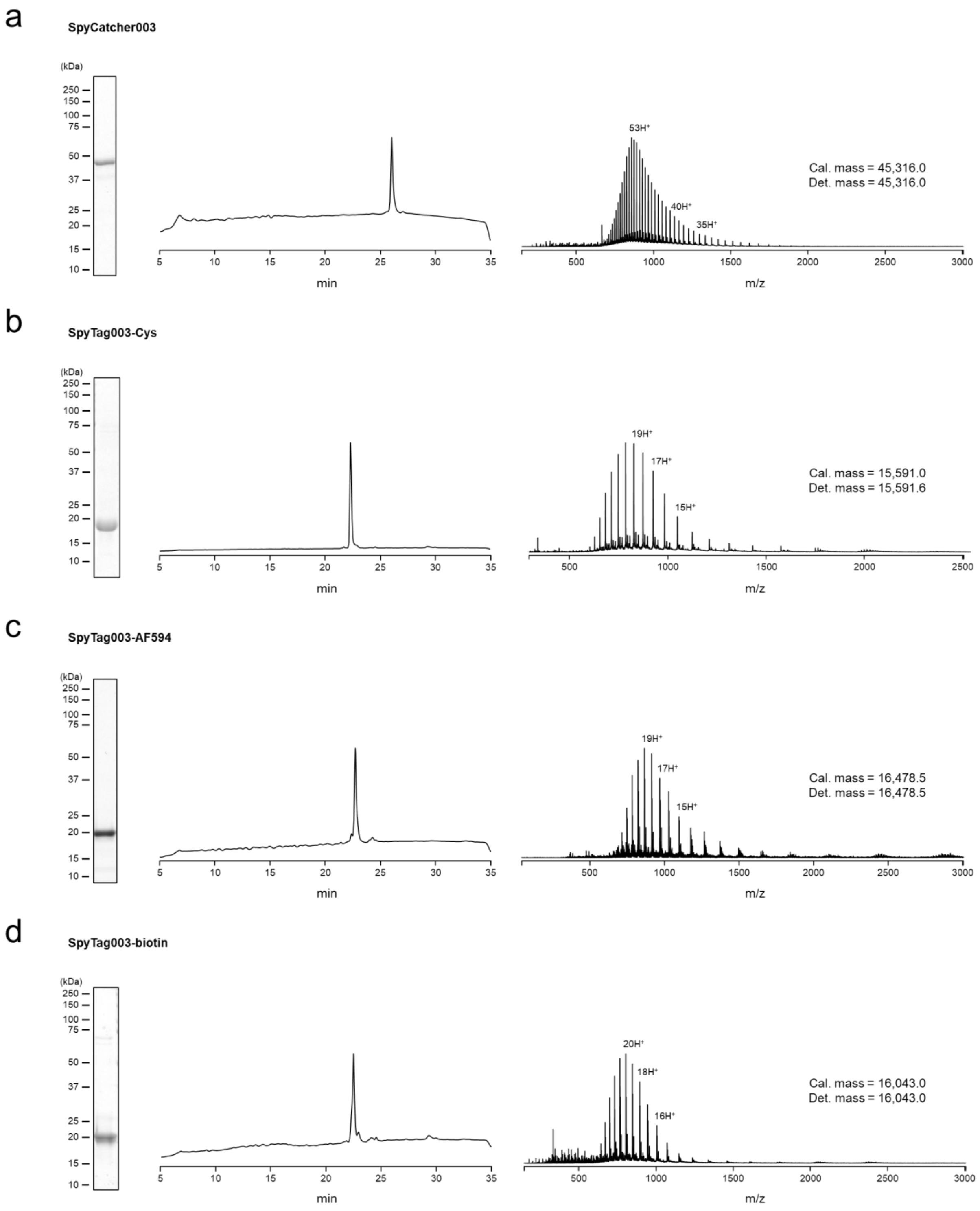
a
SpyCatcher003
(kDa)
250
150
100
75
50
37
25
20
15
10
min
m/z
53H+
40H+
35H+
Cal. mass = 45,316.0
Det. mass = 45,316.0
b
SpyTag003-Cys
19H+
17H+
15H+
Cal. mass = 15,591.0
Det. mass = 15,591.6
c
SpyTag003-AF594
19H+
17H+
15H+
Cal. mass = 16,478.5
Det. mass = 16,478.5
d
SpyTag003-biotin
20H+
18H+
16H+
Cal. mass = 16,043.0
Det. mass = 16,043.0

e

**SpyTag003-DNP**

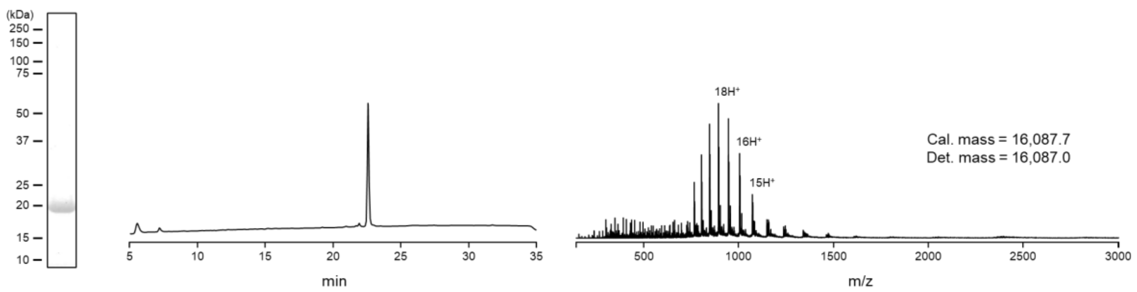


f

**SpyTag003$^{D117A}$-Cys**

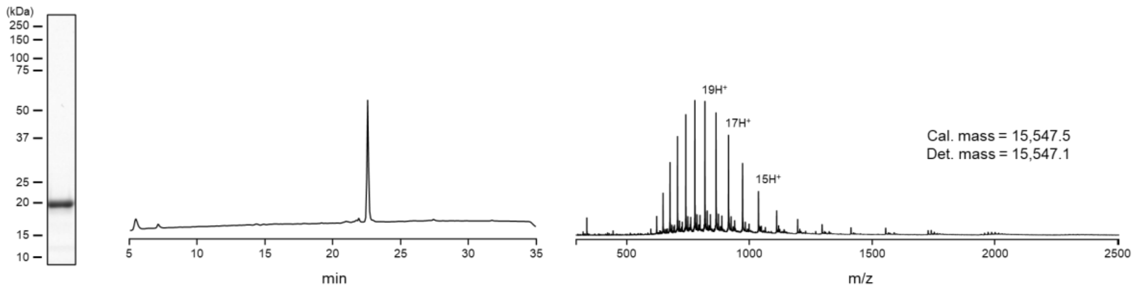


g

**SpyTag003$^{D117A}$-AF594**

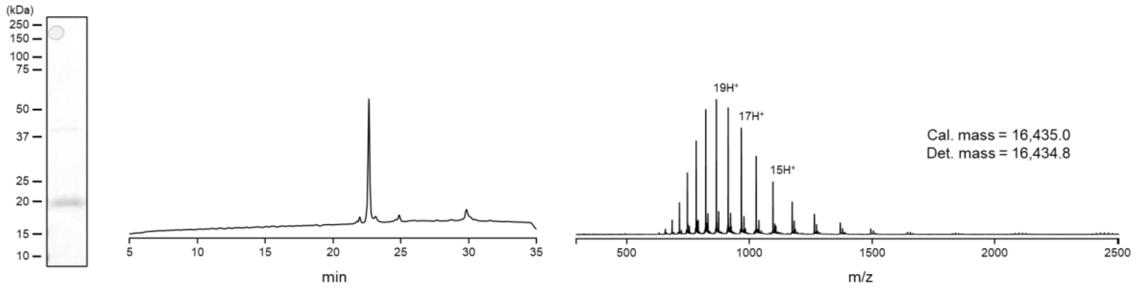


h

**SpyTag003-APEX2**

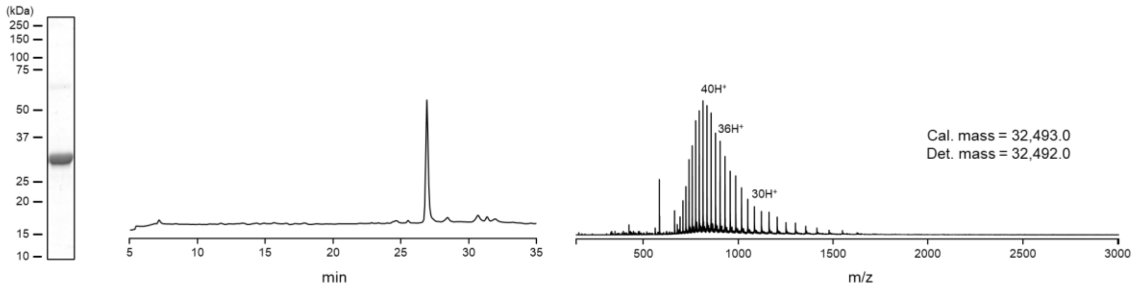

i

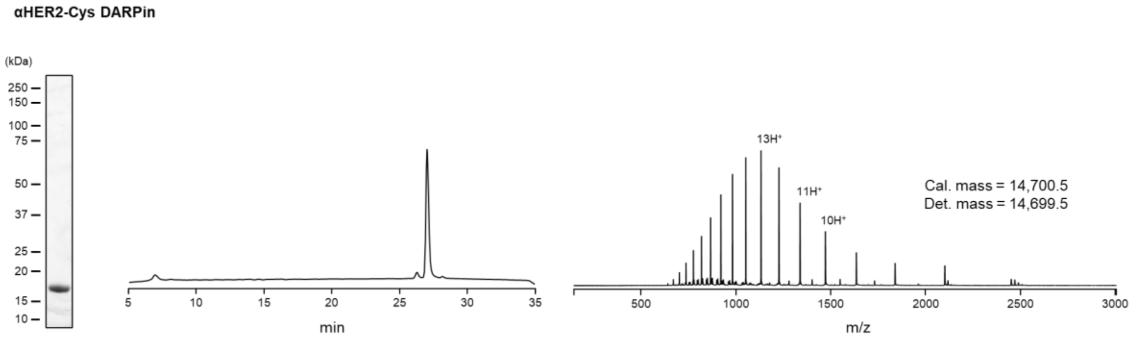
αHER2-Cys DARPin
(kDa)
250
150
100
75
50
37
25
20
15
10
min
13H⁺
11H⁺
10H⁺
Cal. mass = 14,700.5
Det. mass = 14,699.5
m/z

j

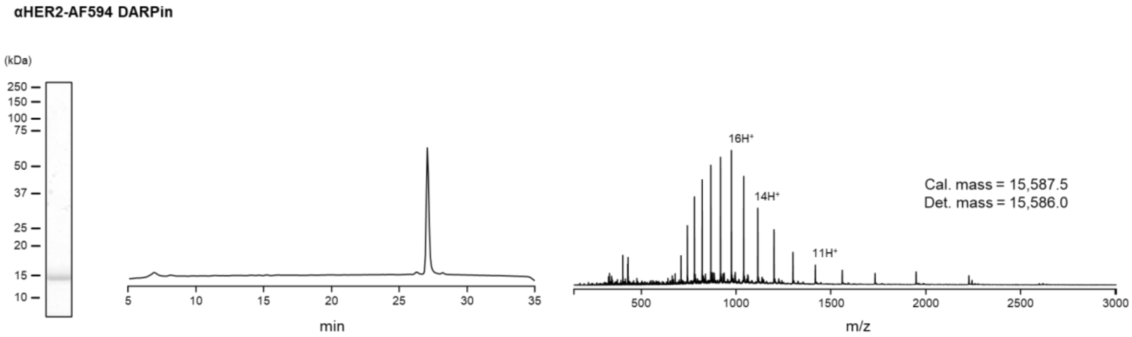
αHER2-AF594 DARPin
(kDa)
250
150
100
75
50
37
25
20
15
10
min
16H⁺
14H⁺
11H⁺
Cal. mass = 15,587.5
Det. mass = 15,586.0
m/z

k

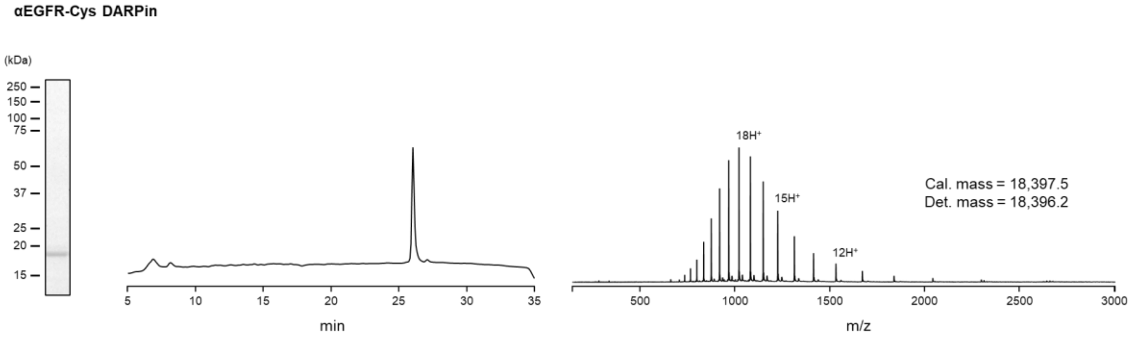
αEGFR-Cys DARPin
(kDa)
250
150
100
75
50
37
25
20
15
min
18H⁺
15H⁺
12H⁺
Cal. mass = 18,397.5
Det. mass = 18,396.2
m/z

l

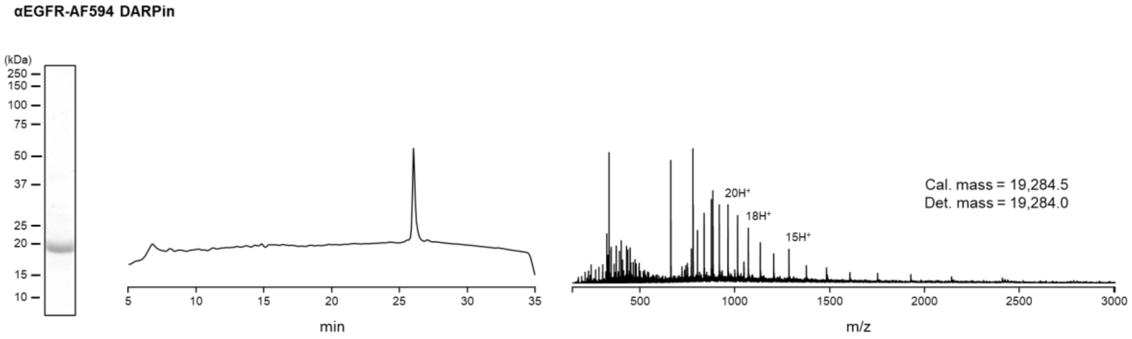
αEGFR-AF594 DARPin
(kDa)
250
150
100
75
50
37
25
20
15
10
min
20H⁺
18H⁺
15H⁺
Cal. mass = 19,284.5
Det. mass = 19,284.0
m/z

m

**αEpCAM-Cys DARPin**

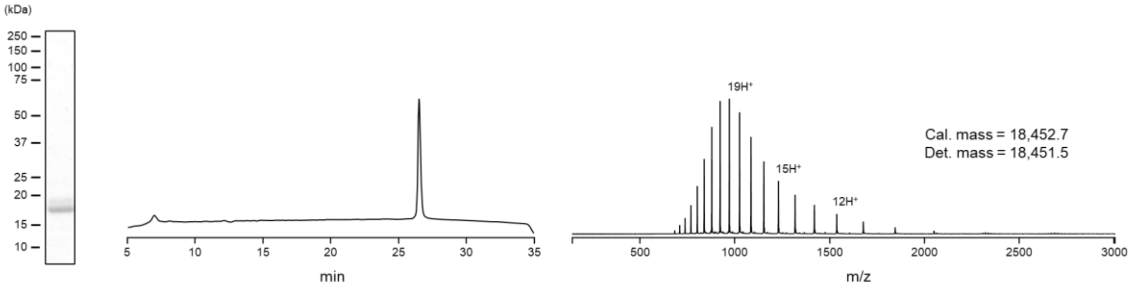


n

**αEpCAM-AF594 DARPin**

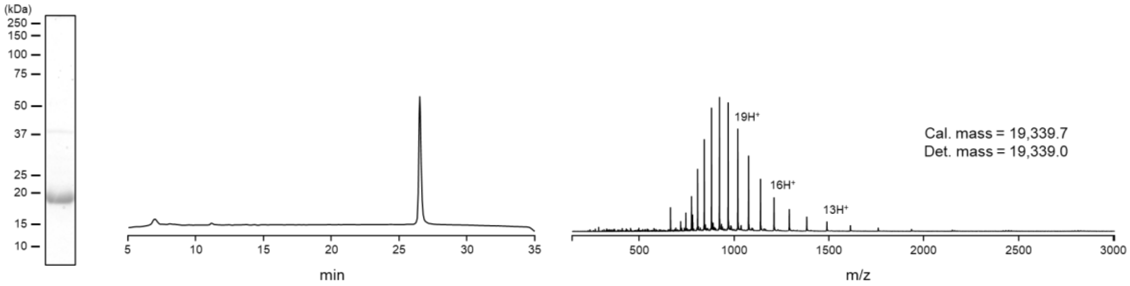


o

**FLAG-SpyN$^{1-24}$-NpuNcage-FKBP**

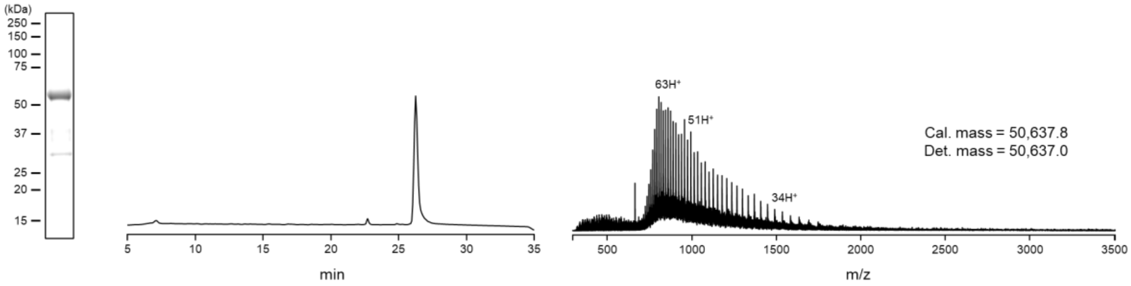


p

**FLAG-SpyN$^{1-42}$-NpuNcage-FKBP**

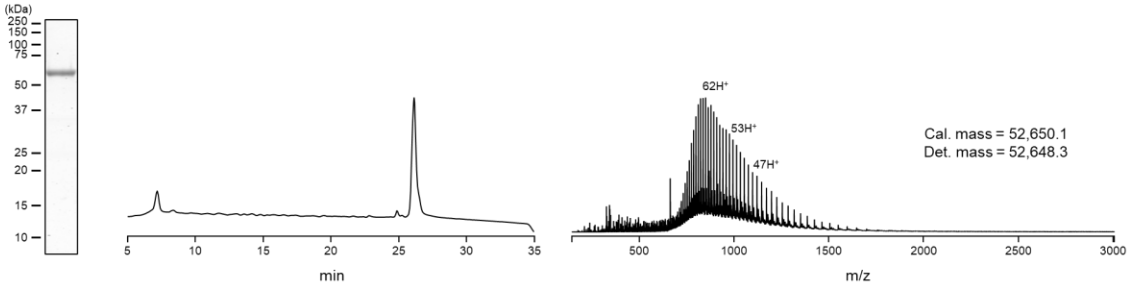

q

**FLAG-SpyN$^{1-55}$-NpuNcage-FKBP**

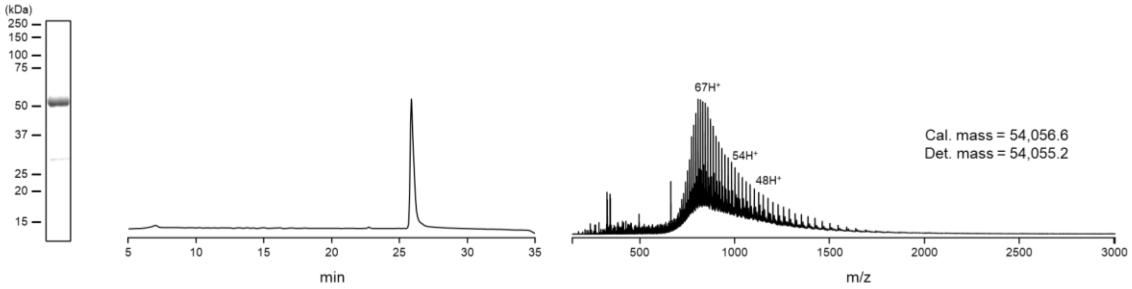


r

**FLAG-SpyN$^{1-73}$-NpuNcage-FKBP**

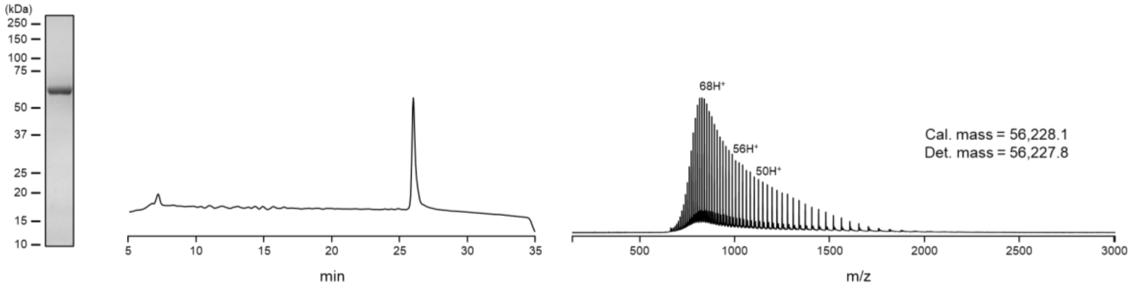


s

**FLAG-SpyN$^{1-82}$-NpuNcage-FKBP**

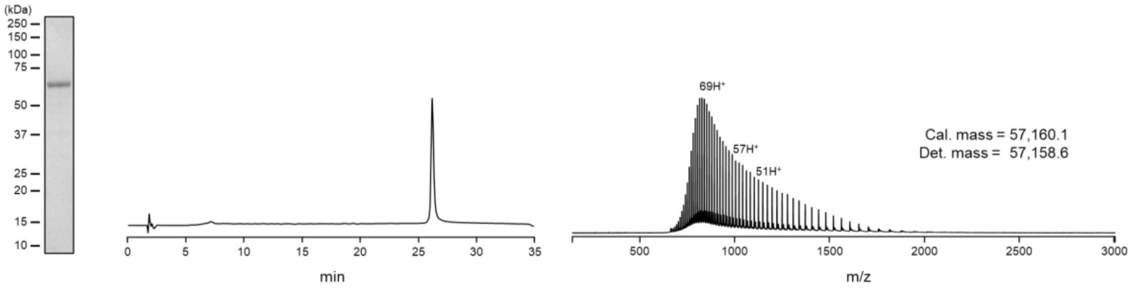


t

**FLAG-SpyN$^{1-90}$-NpuNcage-FKBP**

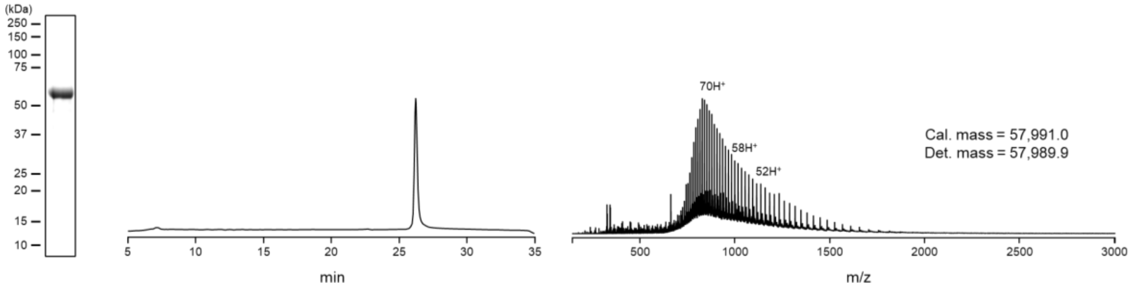

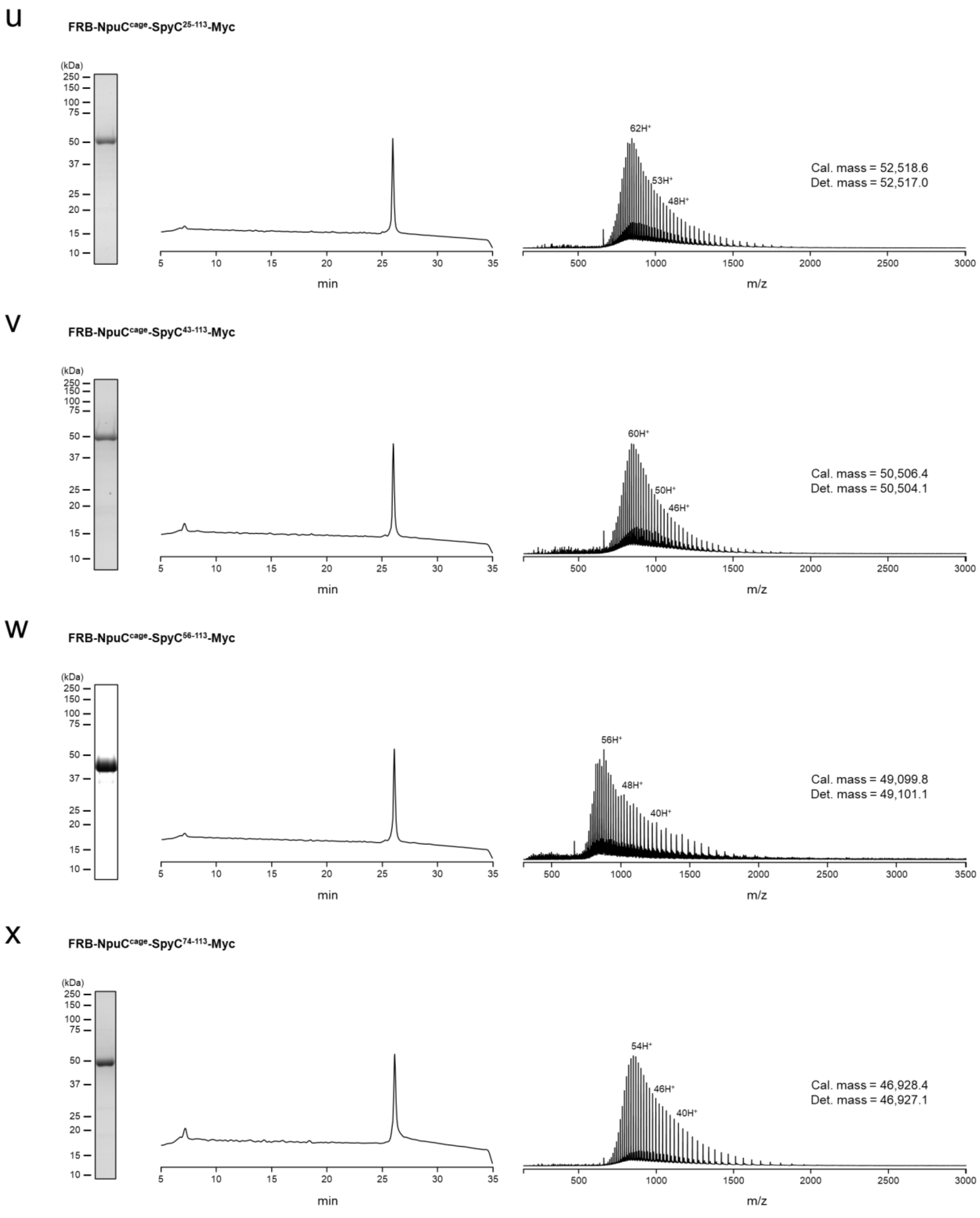
u
FRB-NpuC$^{cage}$-SpyC$^{25-113}$-Myc
Cal. mass = 52,518.6
Det. mass = 52,517.0
62H$^+$
53H$^+$
48H$^+$
v
FRB-NpuC$^{cage}$-SpyC$^{43-113}$-Myc
Cal. mass = 50,506.4
Det. mass = 50,504.1
60H$^+$
50H$^+$
46H$^+$
w
FRB-NpuC$^{cage}$-SpyC$^{56-113}$-Myc
Cal. mass = 49,099.8
Det. mass = 49,101.1
56H$^+$
48H$^+$
40H$^+$
x
FRB-NpuC$^{cage}$-SpyC$^{74-113}$-Myc
Cal. mass = 46,928.4
Det. mass = 46,927.1
54H$^+$
46H$^+$
40H$^+$
(kDa)
min
m/z

**y** FRB-NpuC$^{cage}$-SpyC$^{83\text{-}113}$-Myc

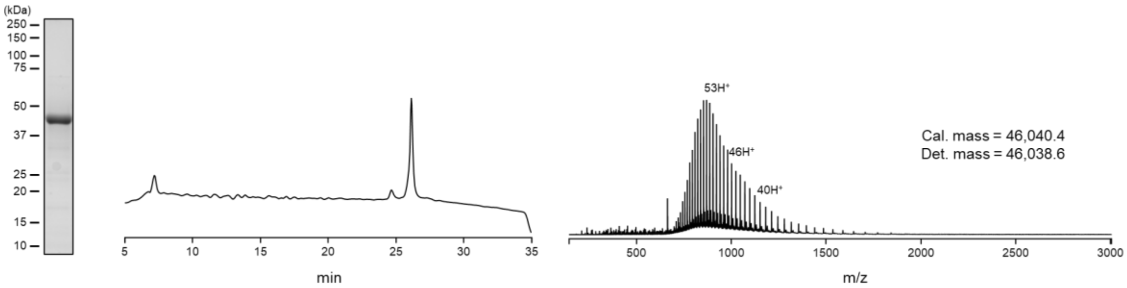


**z** FRB-NpuC$^{cage}$-SpyC$^{91\text{-}113}$-Myc

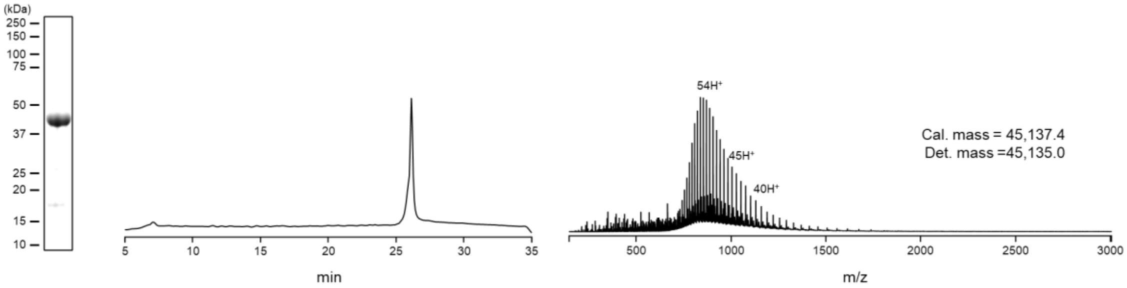


**aa** FLAG-SpyN$^{1\text{-}73}$-NrdJ-1N$^{cage}$-FKBP

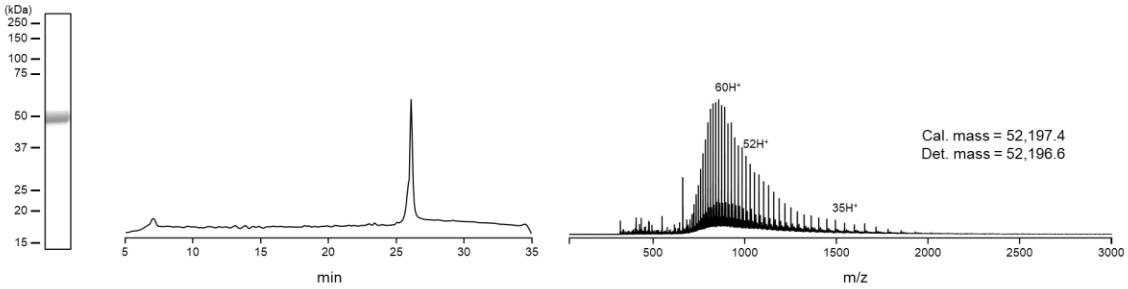


**ab** FLAG-SpyN$^{1\text{-}73}$-NrdJ-1N(C1A)$^{cage}$-FKBP

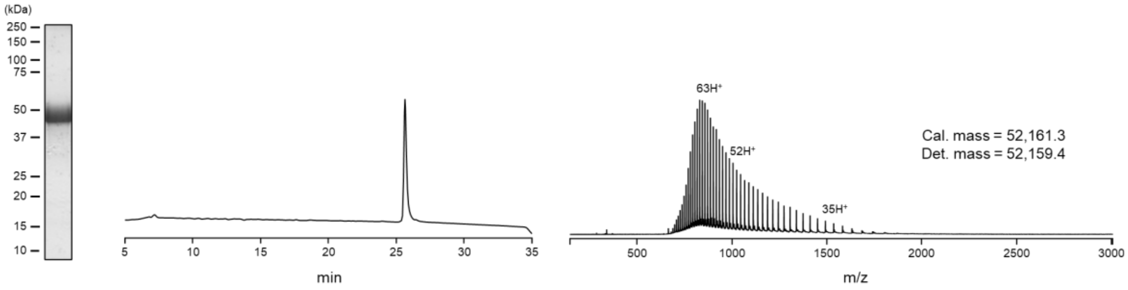

ac

**FLAG-SpyN$^{1-73}$-NrdJ-1N(C76V)$^{cage}$-FKBP**

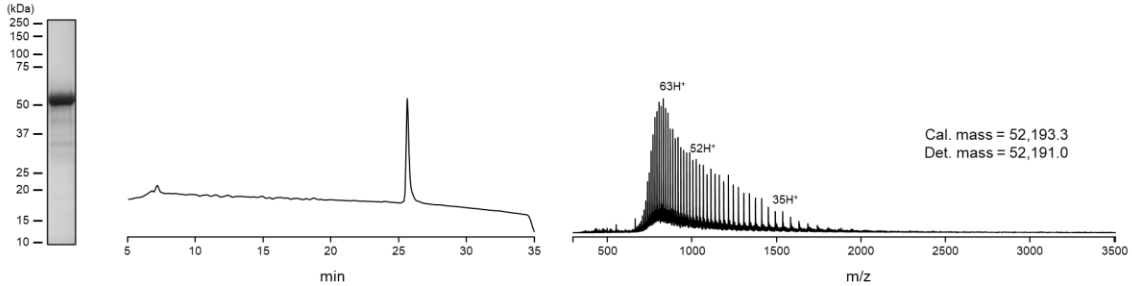


ad

**FRB-NrdJ-1C$^{cage}$-SpyC$^{74-113}$-Myc**

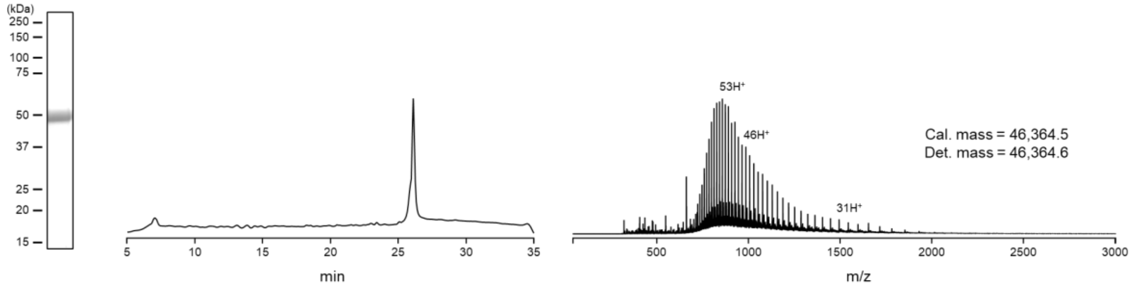


ae

**FRB-NrdJ-1C$^{cage(C76V)}$-SpyC$^{74-113}$**

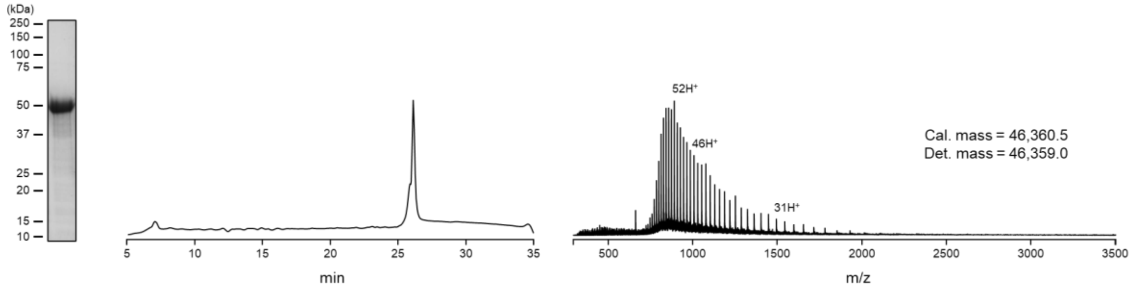


af

**αHER2-SpyN-NrdJ-1N$^{cage(C76V)}$**

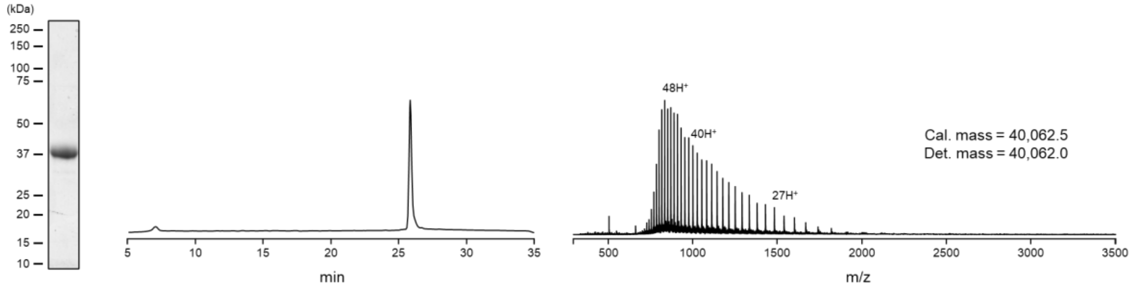

**ag** **αHER2-SpyN-eNrdJ-1$N^{cage}$**

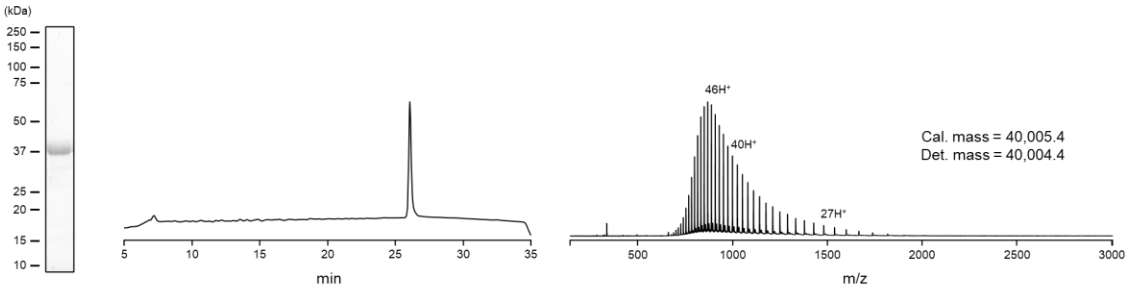


**ah** **αHER2-SpyN-eNrdJ-1$N^{cage}$ C1-alkylated**

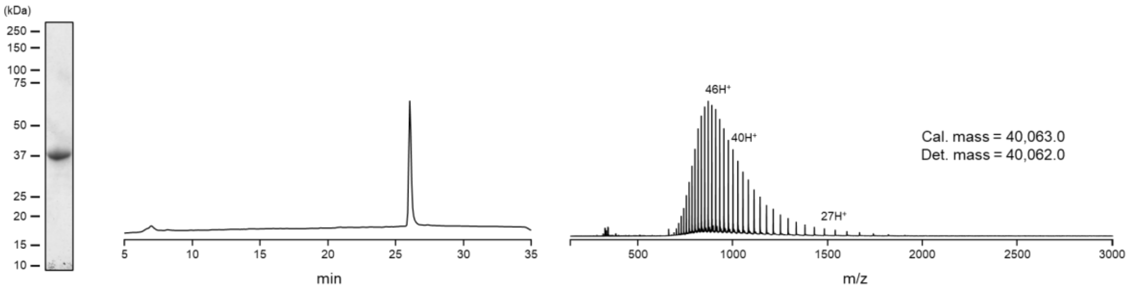


**ai** **αEGFR-SpyN-eNrdJ-1$N^{cage}$**

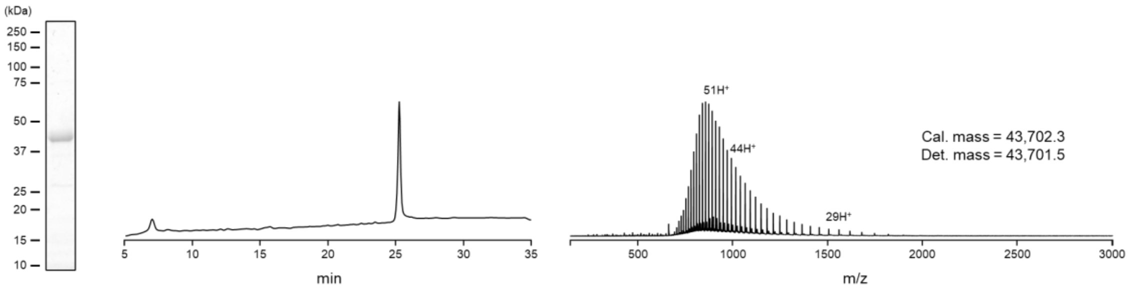


**aj** **αEpCAM-SpyN-eNrdJ-1$N^{cage}$**

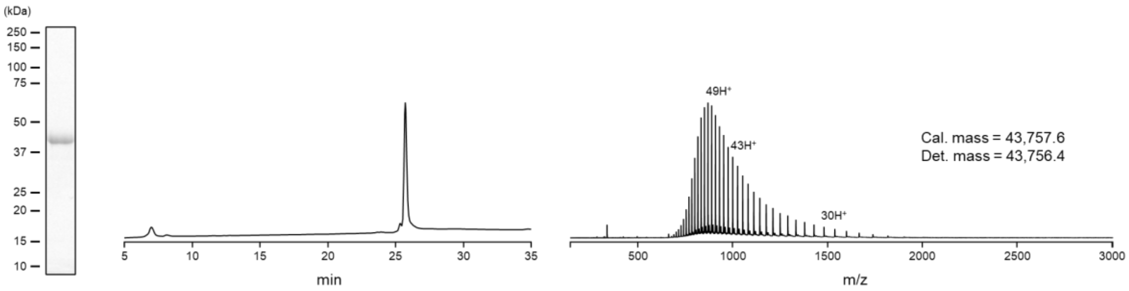

ak

**NrdJ-1C$^{cage(C76V)}$-SpyC-αEGFR**

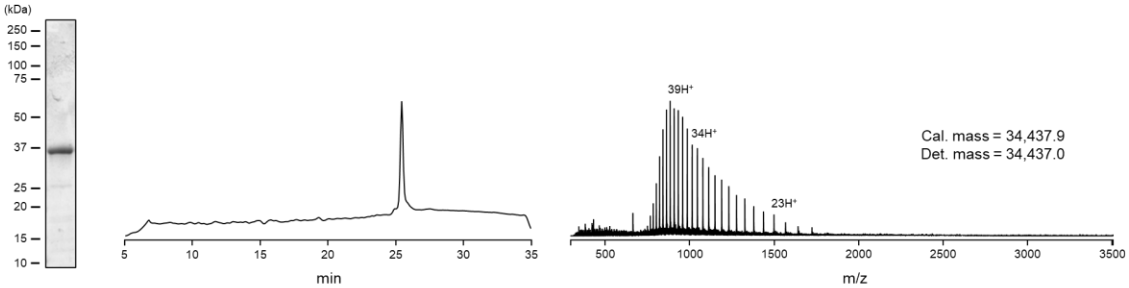


al

**eNrdJ-1C$^{cage}$-SpyC-αHER2**

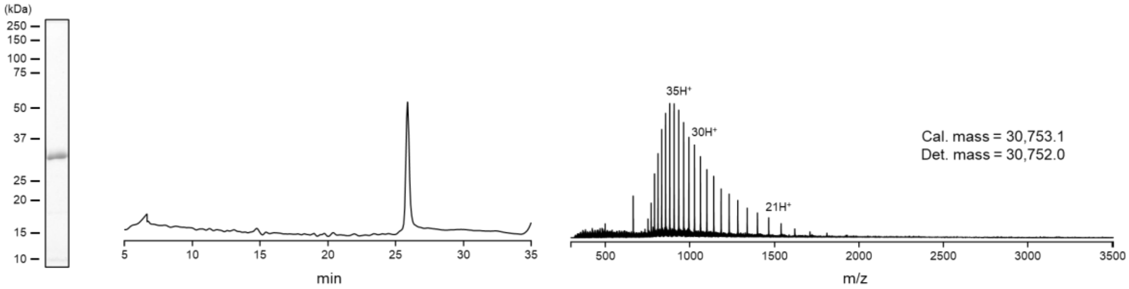


am

**eNrdJ-1C$^{cage}$-SpyC-αEGFR**

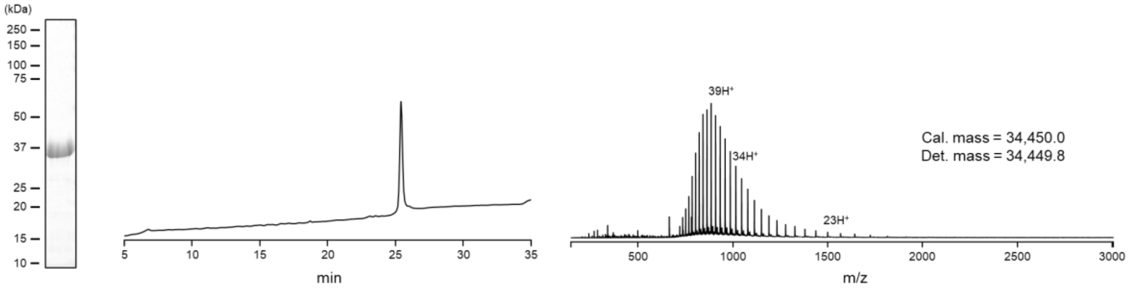


an

**eNrdJ-1C$^{cage}$-SpyC-αEpCAM**

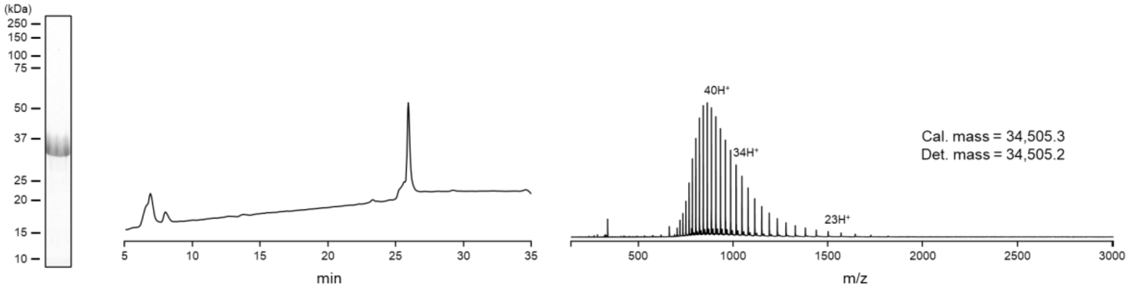

ao

**αHER2-SpyN-eNrdJ-1N$^{cage(K114A)}$**

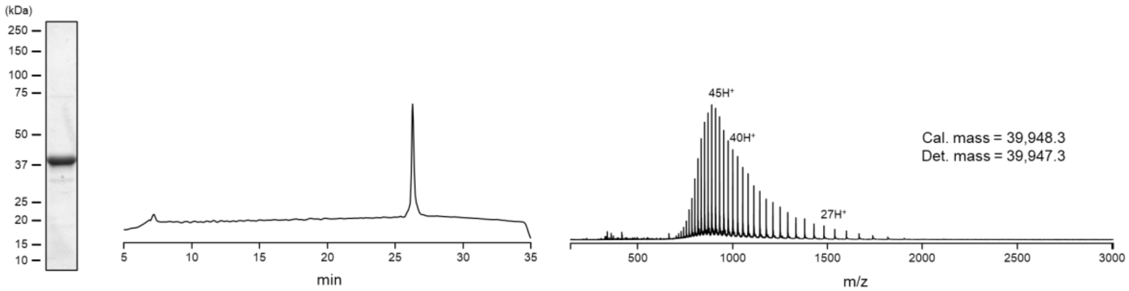


ap

**αHER2-SpyN-eNrdJ-1N$^{cage(K116A)}$**

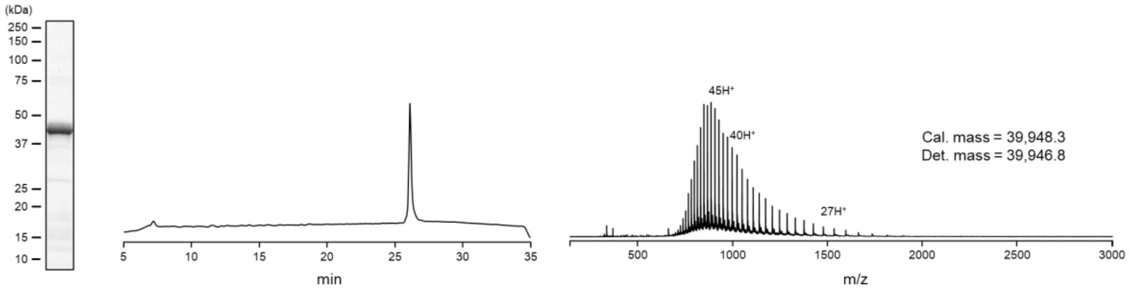


aq

**αHER2-SpyN-eNrdJ-1N$^{cage(K118A)}$**

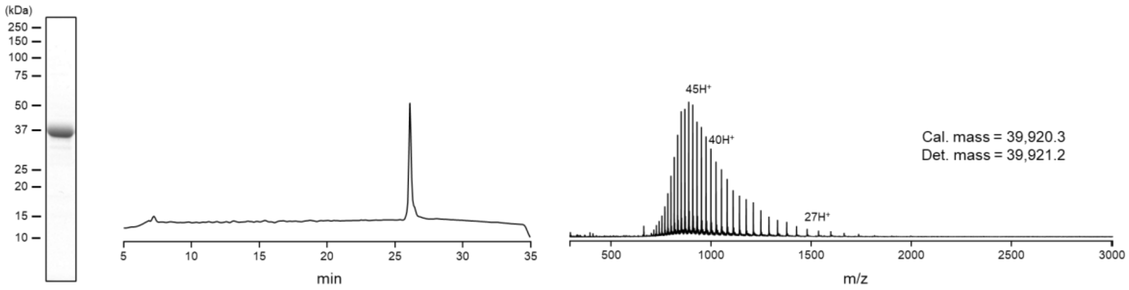


ar

**αHER2-SpyN-eNrdJ-1N$^{cage(K114AK116A)}$**

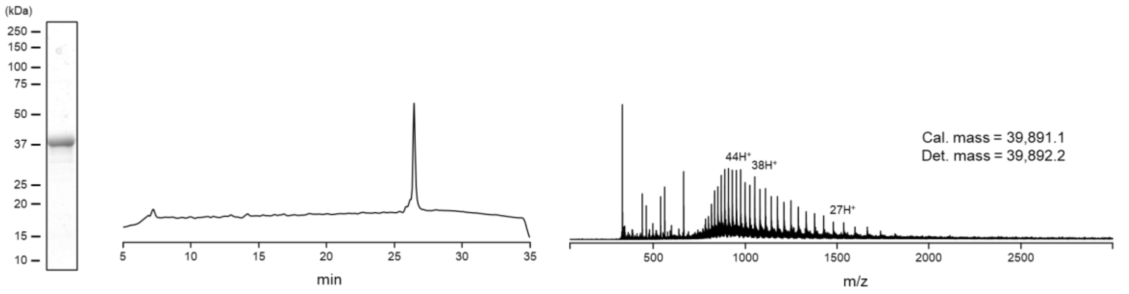

as

**αHER2-SpyN-eNrdJ-1N$^{cage(K114AK116AK118A)}$**

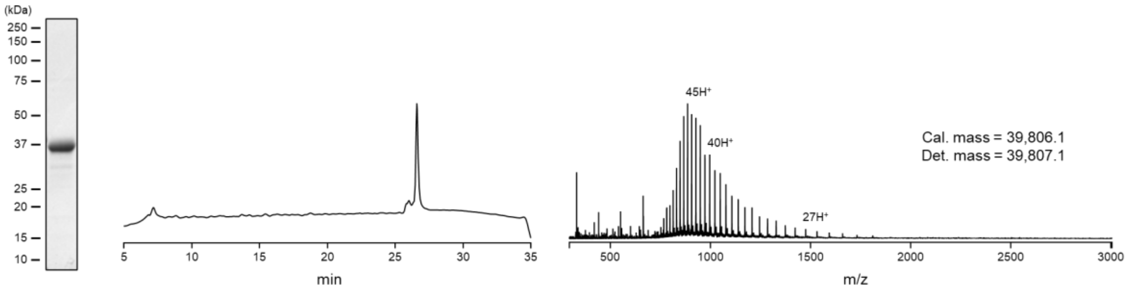


at

**αHER2-SpyN-eNrdJ-1N$^{cage(A119K)}$**

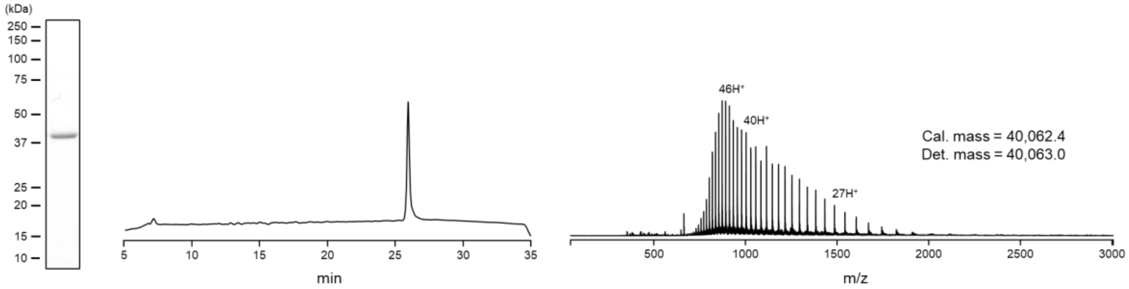


au

**αEGFR-SpyN-eNrdJ-1N$^{cage(K114AK116A)}$**

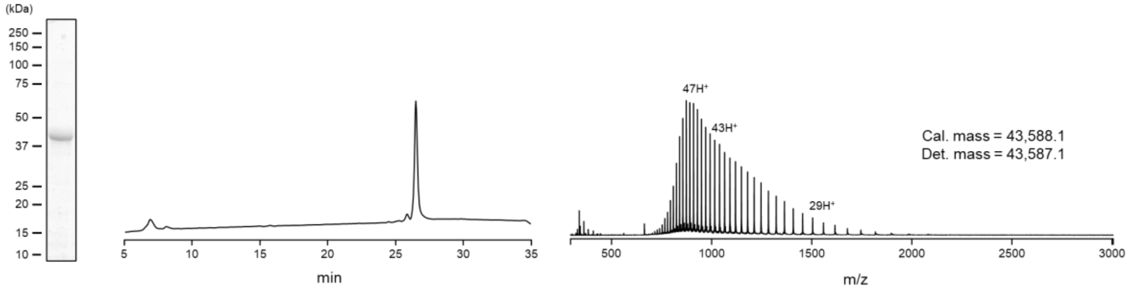


av

**αEpCAM-SpyN-eNrdJ-1N$^{cage(K114AK116A)}$**

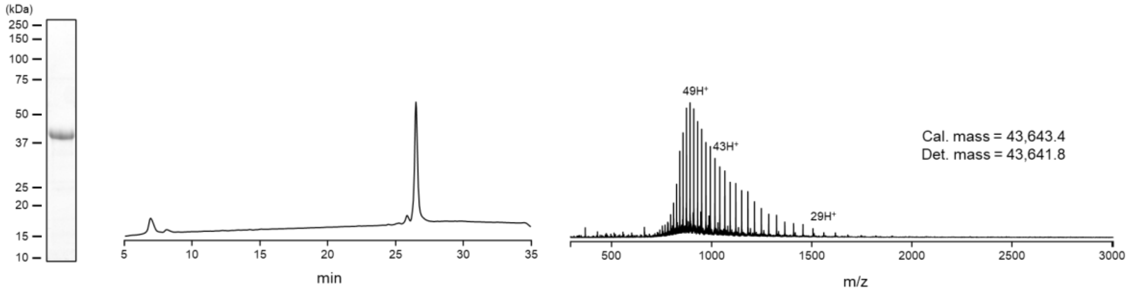

aw

αHER2-SpyN-eNrdJ-1Ncage$^{(1-39)}$

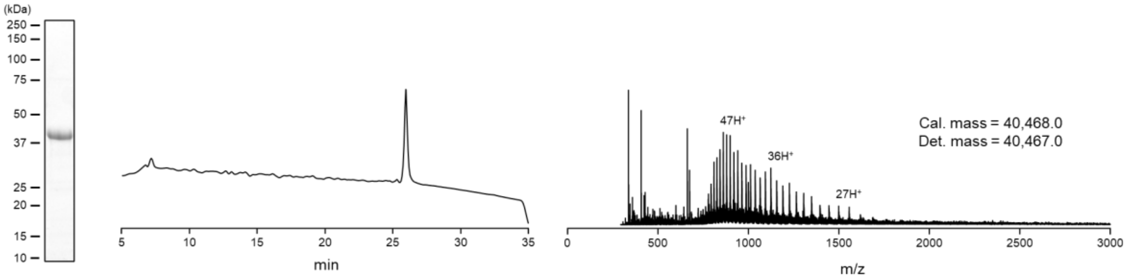


ax

αHER2-SpyN-eNrdJ-1Ncage$^{(1-38)}$

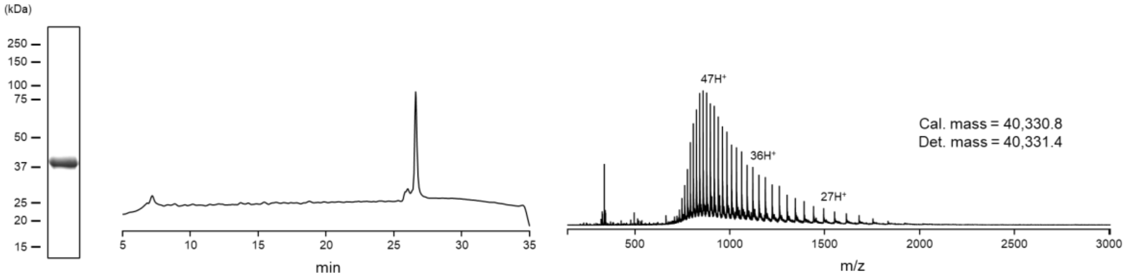


ay

αHER2-SpyN-eNrdJ-1Ncage$^{(1-37)}$

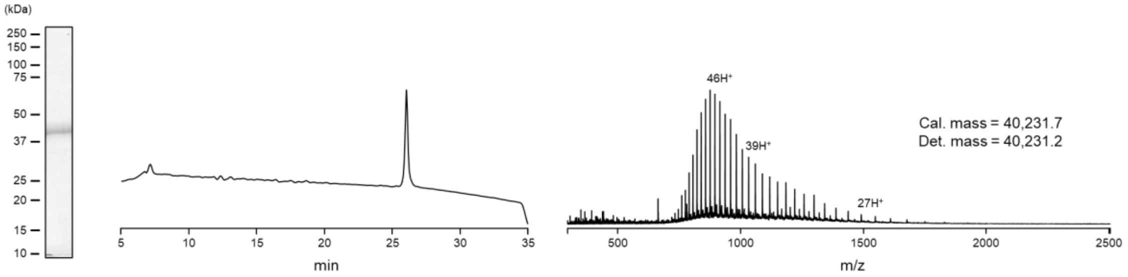


az

αHER2-SpyN-eNrdJ-1N$^{cage(1-36)}$

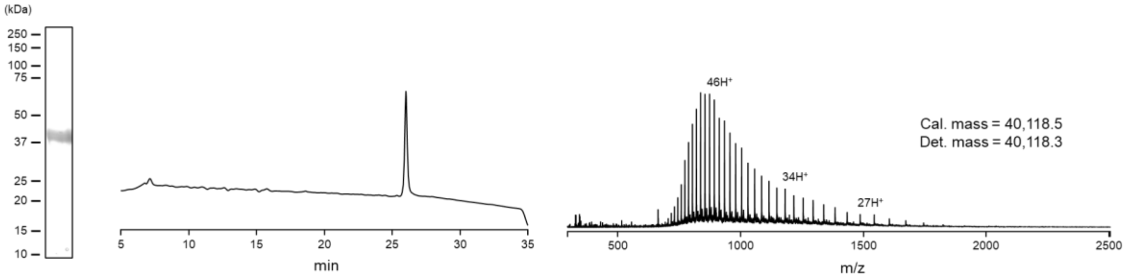

ba αHER2-SpyN-eNrdJ-1N$^{cage(1-33)}$

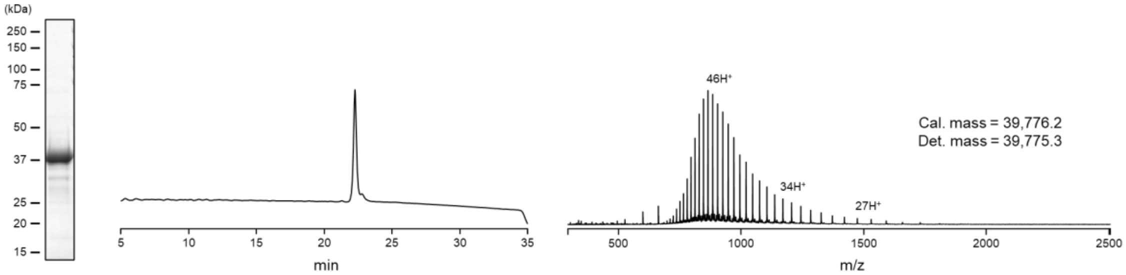


bb αHER2-SpyN-eNrdJ-1N$^{cage(1-32)}$

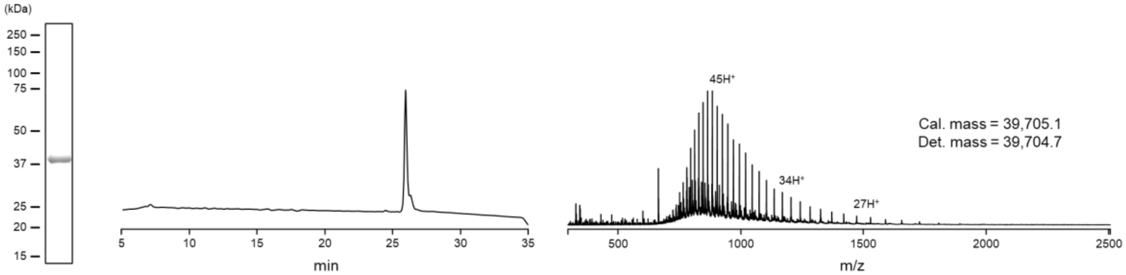


bc αHER2-SpyN-eNrdJ-1N$^{cage(1-31)}$

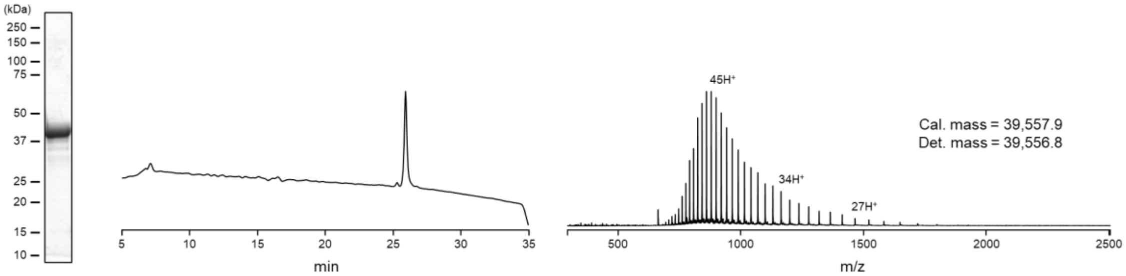


bd αHER2-SpyN-eNrdJ-1N$^{cage(1-30)}$

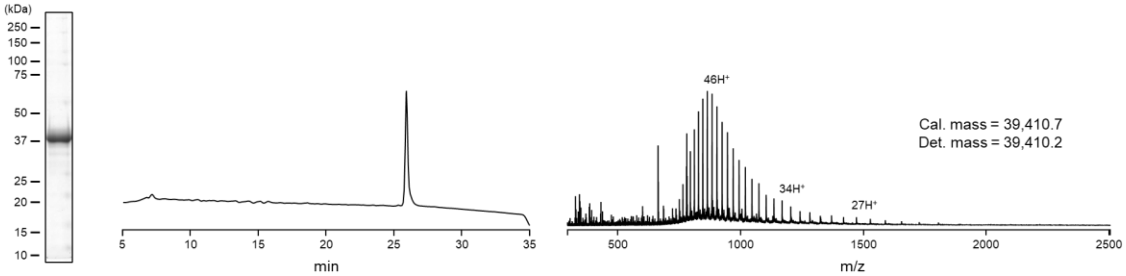

be

**αHER2-SpyN-eNrdJ-1N$^{cage(1-29)}$**

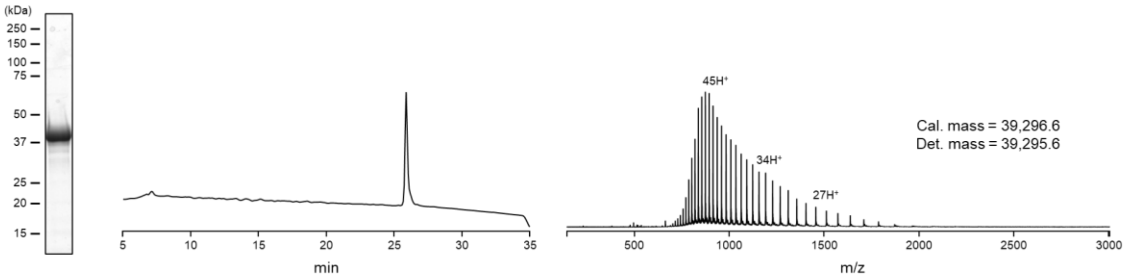


bf

**αHER2-SpyN-eNrdJ-1N$^{cage(1-26)}$**

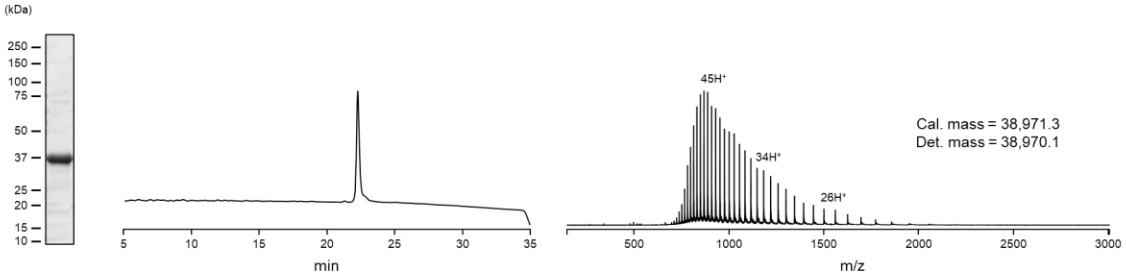


bg

**αHER2-SpyN-eNrdJ-1N$^{cage(1-20)}$**

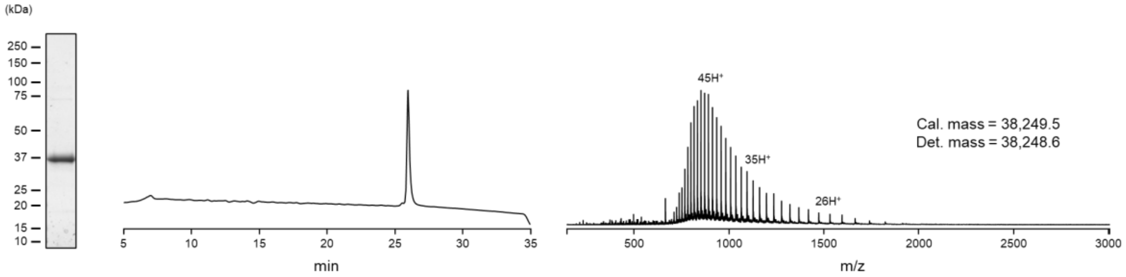


bh

**αHER2-SpyN-NrdJ-1N**

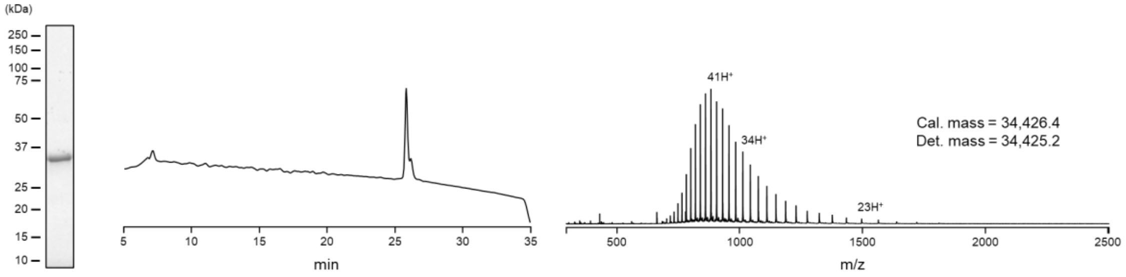

bi

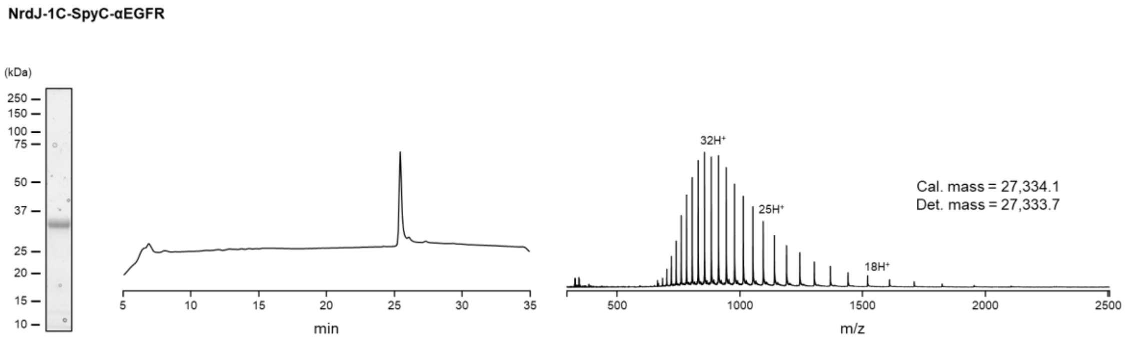


bj

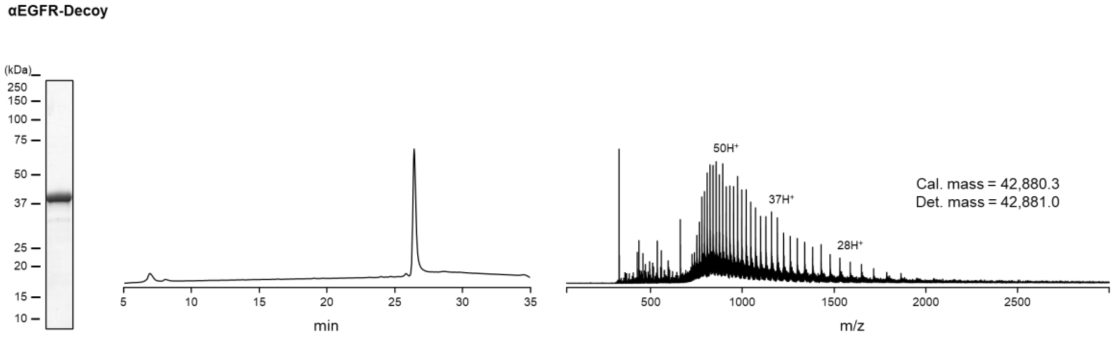


**Supplementary Figure 1. Protein purification and characterization.** Protein preparations were analyzed by SDS-PAGE (left), RP-HPLC (0-70% B gradient over 30 min; middle), and ESI-TOF MS (right): **a**, SpyCatcher003, **b**, SpyTag003-Cys, **c**, SpyTag003-AF594, **d**, SpyTag003-biotin, **e**, SpyTag003-DNP, **f**, SpyTag003$^{D117A}$-Cys, **g**, SpyTag003$^{D117A}$-AF594, **h**,SpyTag003-APEX2, **i**, αHER2-Cys DARPin, **j**, αHER2-AF594 DARPin, **k**, αEGFR-Cys DARPin, **l**, αEGFR-AF594 DARPin, **m**, αEpCAM-Cys DARPin, **n**, αEpCAM-AF594 DARPin, **o**, FLAG-SpyN$^{1-24}$-NpuN$^{cage}$-FKBP, **p**, FLAG-SpyN$^{1-42}$-NpuN$^{cage}$-FKBP, **q**, FLAG-SpyN$^{1-55}$-NpuN$^{cage}$-FKBP, **r**, FLAG-SpyN$^{1-73}$-NpuN$^{cage}$-FKBP, **s**, FLAG-SpyN$^{1-82}$-NpuN$^{cage}$-FKBP, **t**, FLAG-SpyN$^{1-90}$-NpuN$^{cage}$-FKBP, **u**, FRB-NpuC$^{cage}$-SpyC$^{25-113}$-Myc, **v**, FRB-NpuC$^{cage}$-SpyC$^{43-113}$-Myc, **w**, FRB-NpuC$^{cage}$-SpyC$^{56-113}$-Myc, **x**, FRB-NpuC$^{cage}$-SpyC$^{74-113}$-Myc, **y**, FRB-NpuC$^{cage}$-SpyC$^{83-113}$-Myc, **z**, FRB-NpuC$^{cage}$-SpyC$^{91-113}$-Myc, **aa**, FLAG-SpyN$^{1-73}$-NrdJ-1N$^{cage}$-FKBP, **ab**, FLAG-SpyN$^{1-73}$-NrdJ-1N(C1A)$^{cage}$-FKBP, **ac**, FLAG-SpyN$^{1-73}$-NrdJ-1N(C76V)$^{cage}$-FKBP, **ad**, FRB-NrdJ-1C$^{cage}$-SpyC$^{74-113}$-Myc, **ae**, FRB-NrdJ-1C$^{cage(C76V)}$-SpyC$^{74-113}$-Myc, **af**, αHER2-SpyN-NrdJ-1N(C76V)$^{cage}$, **ag**, αHER2-SpyN-eNrdJ-1N$^{cage}$, **ah**, αHER2-SpyN-eNrdJ-1N$^{cage}$ Cys1-alkylated, **ai**, αEGFR-SpyN-eNrdJ-1N$^{cage}$, **aj**, αEpCAM-SpyN-eNrdJ-1N$^{cage}$, **ak**, NrdJ-1C$^{cage(C76V)}$-SpyC-αEGFR, **al**, eNrdJ-1C$^{cage}$-SpyC-αHER2, **am**, eNrdJ-1C$^{cage}$-SpyC-αEGFR, **an**, eNrdJ-1C$^{cage}$-SpyC-αEpCAM, **ao**, αHER2-SpyN-eNrdJ-1N$^{cage(K114A)}$, **ap**, αHER2-SpyN-eNrdJ-1N$^{cage(K116A)}$, **aq**, αHER2-SpyN-eNrdJ-1Ncage$^{(R118A)}$, **ar**, αHER2-SpyN-eNrdJ-1Ncage$^{(K114AK116A)}$, **as**, αHER2-SpyN-eNrdJ-1Ncage$^{(K114AK116AR118A)}$, **at**, αHER2-

SpyN-eNrdJ-1Ncage$^{(A119K)}$, **au**, αEGFR-SpyN-eNrdJ-1Ncage$^{(K114AK116A)}$, **av**, αEpCAM-SpyN-eNrdJ-1Ncage$^{(K114AK116A)}$, **aw**, αHER2-SpyN-eNrdJ-1Ncage$^{(1-39)}$, **ax**, αHER2-SpyN-eNrdJ-1Ncage$^{(1-38)}$, **ay**, αHER2-SpyN-eNrdJ-1Ncage$^{(1-37)}$, **az**, αHER2-SpyN-eNrdJ-1Ncage$^{(1-36)}$, **ba**, αHER2-SpyN-eNrdJ-1Ncage$^{(1-33)}$, **bb**, αHER2-SpyN-eNrdJ-1Ncage$^{(1-32)}$, **bc**, αHER2-SpyN-eNrdJ-1Ncage$^{(1-31)}$, **bd**, αHER2-SpyN-eNrdJ-1Ncage$^{(1-30)}$, **be**, αHER2-SpyN-eNrdJ-1Ncage$^{(1-29)}$, **bf**, αHER2-SpyN-eNrdJ-1Ncage$^{(1-26)}$, **bg**, αHER2-SpyN-eNrdJ-1Ncage$^{(1-20)}$, **bh**, αHER2-SpyN-NrdJ-1N, **bi**, NrdJ-1C-SpyC-αEGFR, **bj**, αEGFR-Decoy.

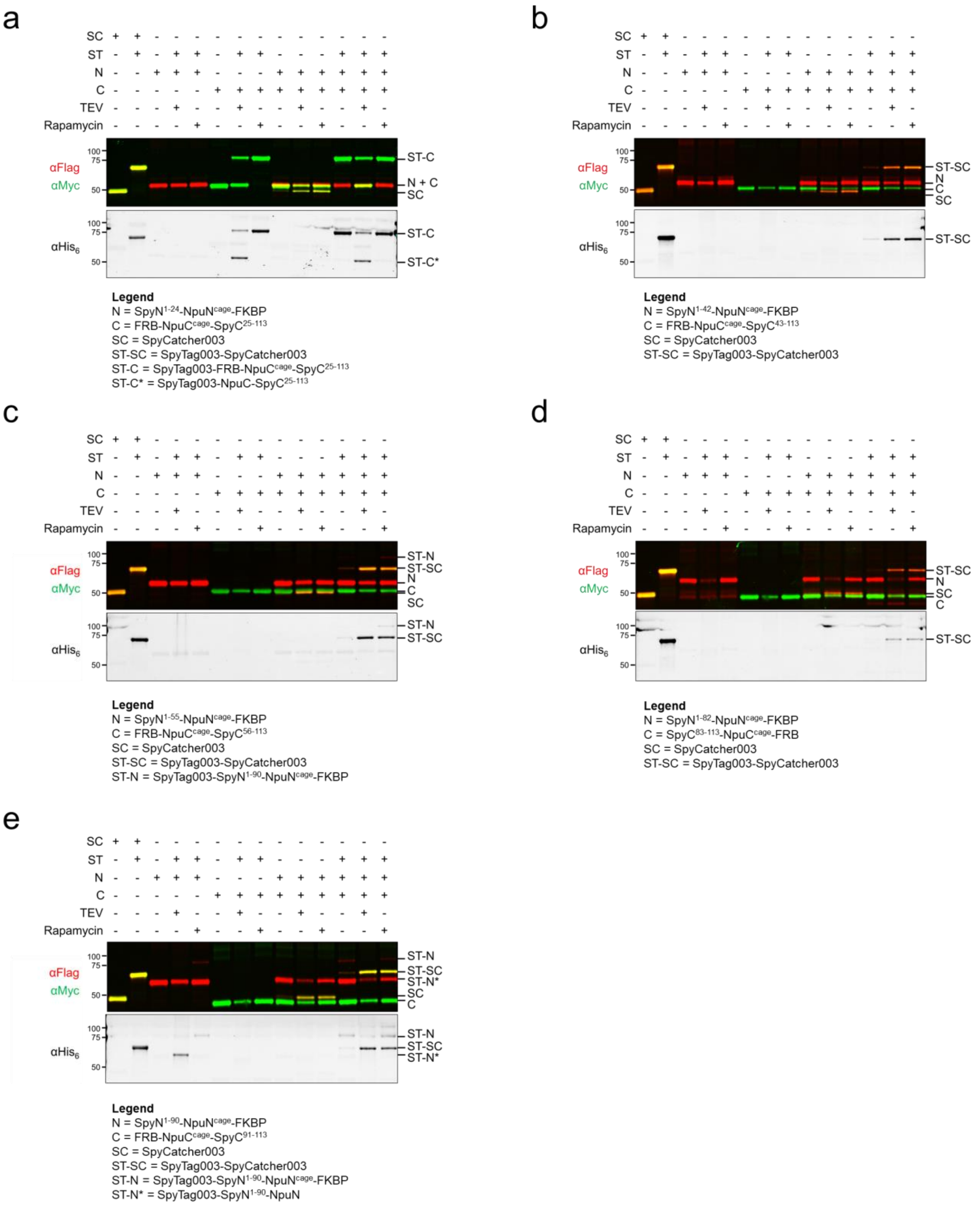


**Supplementary Figure 2. Conditional protein splicing of SpyN-NpuN$^{cage}$-FKBP and FRB-NpuC$^{cage}$-SpyC. a-e**, The results of screening FLAG-SpyN$^{1-x}$-NpuN$^{cage}$-FKBP and FRB-NpuC$^{cage}$-SpyC$^{y-113}$-Myc with x and y designating last and first residue of the N- and C-terminal fragments generated by splitting SpyCatcher003 at site x/y. Reactions were performed with FLAG-SpyN$^{1\text{-}x}$-NpuN$^{cage}$-FKBP (1 μM), FRB-NpuC$^{cage}$-SpyC$^{y\text{-}113}$-Myc, (1 μM) and His$_6$-SpyTag003 (2 μM). CPS was induced by the addition of either TEV protease (10 units) or rapamycin (10 μM). The reactions were analyzed by Western blot after 24 hr incubation at 37 °C. FLAG-SpyCatcher003-Myc (1 μM) was used as a size standard for the spliced product; FLAG-SpyCatcher003-Myc (1 μM) reacted with His$_6$-SpyTag003 (2 μM) was used as a size standard for the covalent complex between the two.

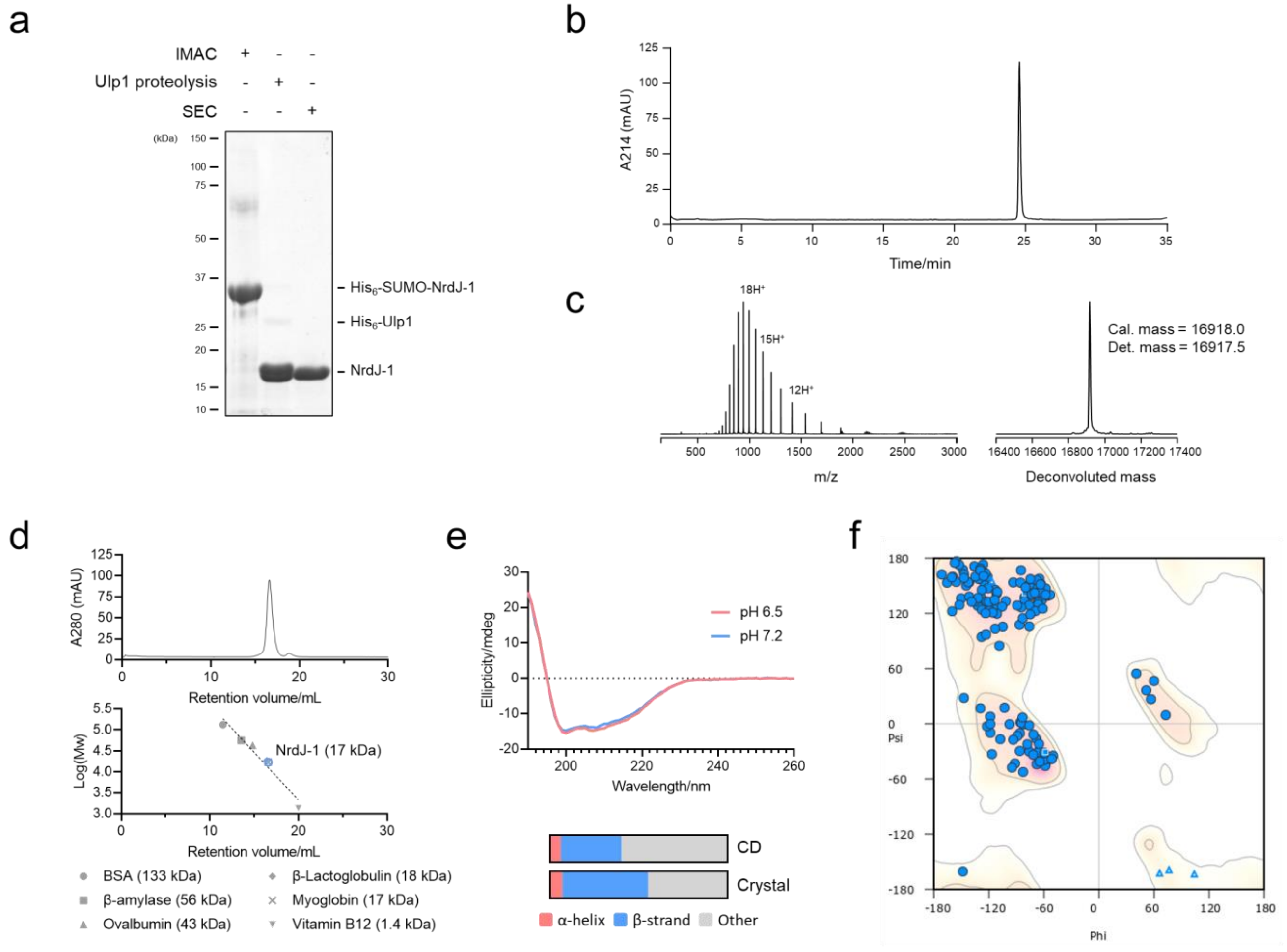


**Supplementary Figure 3. Preparation and characterization of the NrdJ-1 fusion. a**, SDS-PAGE, and **b**, RP-HPLC analysis (0-70% B gradient over 30 min) of NrdJ-1 characterizing the purity of the preparation. The protein was further characterized by **c**, ESI-TOF MS for mass confirmation and **d**, analytical gel-filtration chromatography validating its monomeric status. **e**, Circular dichroism spectra displaying the secondary structure of NrdJ-1 sampled at pH 6.5 and 7.2, which are largely superimposable (top). The spectrum at pH 7.2 was used to quantify the ratios of secondary structure elements (assessed by DichroWeb[1]) and compared to that estimated from the crystal structure (bottom). **f**, Ramachandran plot generated in Coot of the crystal structure of NrdJ-1 (A chain) showing no outliers.

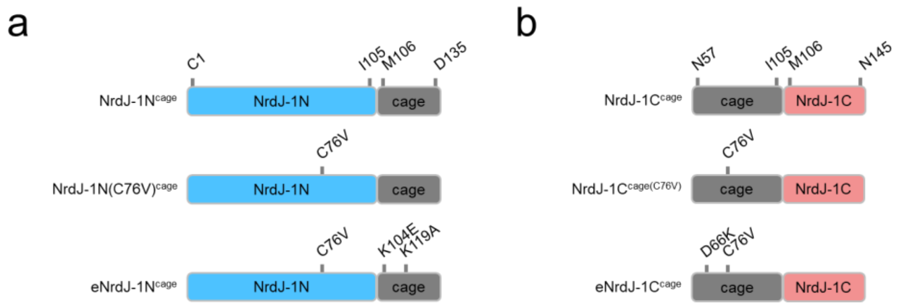


**Supplementary Figure 4. Development of an optimal NrdJ-1 actuator. a**-**b**, Summary of the mutations introduced to optimize NrdJ-1$^{cage}$. The N- and C-terminal residues for each caged fragment and their respective cages are displayed along with relevant mutations introduced: NrdJ-1N spans residues Cys1 to Ile105, its cage from Met106 to Asp135; NrdJ-1C spans residues Met106 to Asn145, its cage from Asn57 to Ile105.

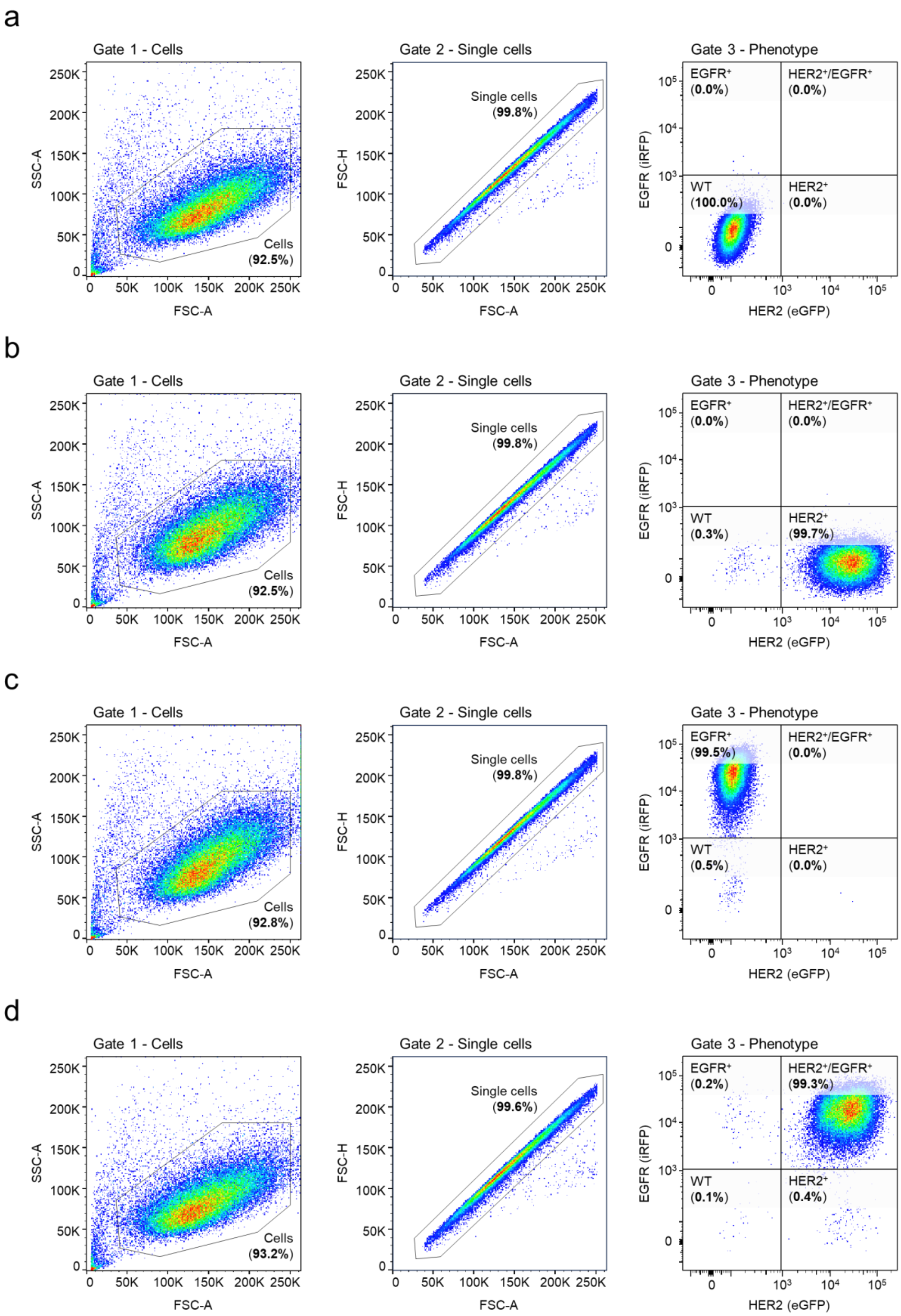


**Supplementary Figure 5. Flow cytometry of individual K562 cell lines.** Displayed is the flow cytometry data for **a**, K562 (WT), **b**, K562$^{HER2+}$, **c**, K562$^{EGFR+}$, and **d**, K562$^{HER2+/EGFR+}$ cell lines validating their HER2-eGFP/EGFR-iRFP phenotypes. Three gates were used to define the cell population: Gate 1 was set as forward scatter area (FSC-A) versus side scatter area (SSC-A) to identify cells; gate 2 was set as forward scatter area (FSC-A) versus forward scatter height (FSC-H) to isolate single cells; gate 3 was set as eGFP fluorescence versus iRFP fluorescence to categorize K562 cells by their HER2-eGFP and EGFR-iRFP phenotypes (their distribution is shown in percentages).

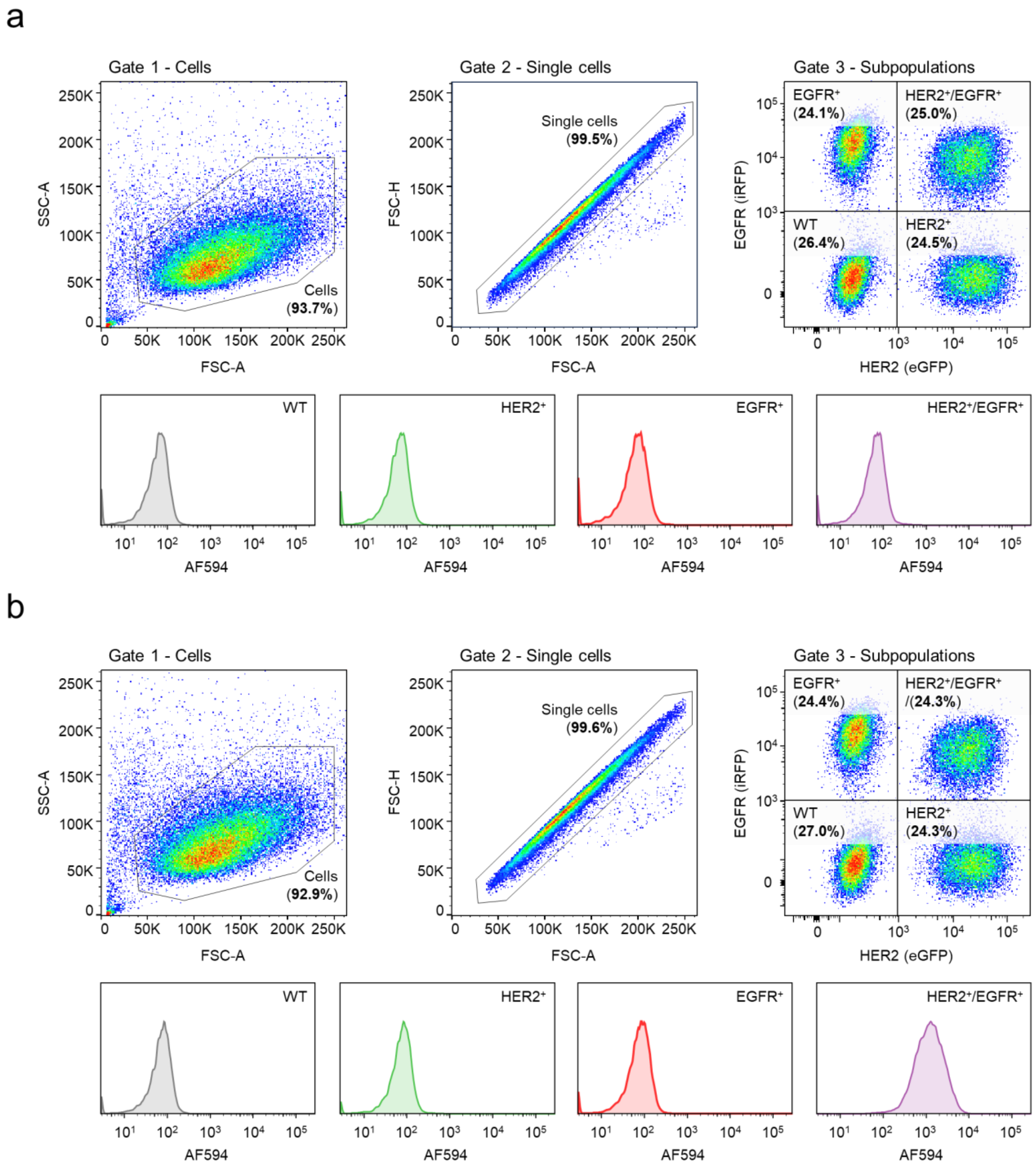


**Supplementary Figure 6. Flow cytometry gating strategy for mixed K562 population 1 flow cytometry experiments.** Example of flow cytometry gating strategy for mixed K562 population 1 (equal amounts of K562 (wildtype), K562$^{HER2+}$, K562$^{EGFR+}$, and K562$^{HER2+/EGFR+}$ cells). **a,** The mixed population was, treated with SpyTag003-AF594 (100 nM) for 20 min and analyzed by flow cytometry. Three gates were used to define the cell population: Gate 1 was set as forward scatter area (FSC-A) versus side scatter area (SSC-A) to identify cells; gate 2 was set as forward scatter area (FSC-A) versus forward scatter height (FSC-H) to isolate single cells; gate 3 was set as eGFP fluorescence versus iRFP fluorescence to categorize K562 subpopulations by their HER2-eGFP and EGFR-iRFP phenotypes (their distribution is shown in percentages). Below the three gating plots are shown the AF594 histograms associated with each individual subpopulation. **b,** The mixed population was treated with αHER2-SpyN (100 nM) and SpyC-αEGFR (100 nM) for 2 hr, followed by SpyTag003 labeled with Alexa Fluor 594 (SpyTag003-AF594) for 20 min. The cell mixture was then analyzed by flow cytometry as described in panel **a**.

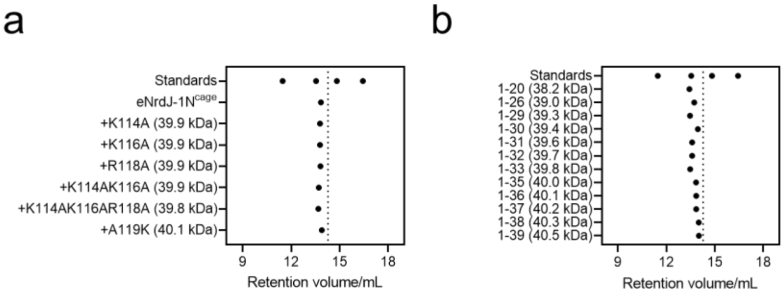


**Supplementary Fig. 7. The elution profiles of αHER2-SpyN employing tuned eNrdJ-1N$^{cage}$ variants.** The retention volume profiles of all tested tuned variants were determined by size-exclusion chromatography. All variants were analyzed in 100 mM sodium phosphate (pH 7.2), 150 mM NaCl, 1 mM EDTA at 1 mg/mL which equates to 25-26 µM (>250-fold higher than the concentrations used in the standard cell assay). The determined hydrodynamic radii correspond to monomeric statuses when compared to a known set of size-standards (from left to right: BSA-dimer (133 kDa); β-amylase (56 kDa); Ovalbumin (43 kDa); Myoglobulin (17 kDa); the dotted line indicates the theoretical retention volume of 14.27 mL for a globular 40 kDa protein. The retention volume profiles for the αHER2-SpyN variants employing **a**, electrostatically engineered cages, or **b**, toehold engineered cages as indicated. Standard eNrdJ-1N$^{cage}$ employs the cage length of 1-35.

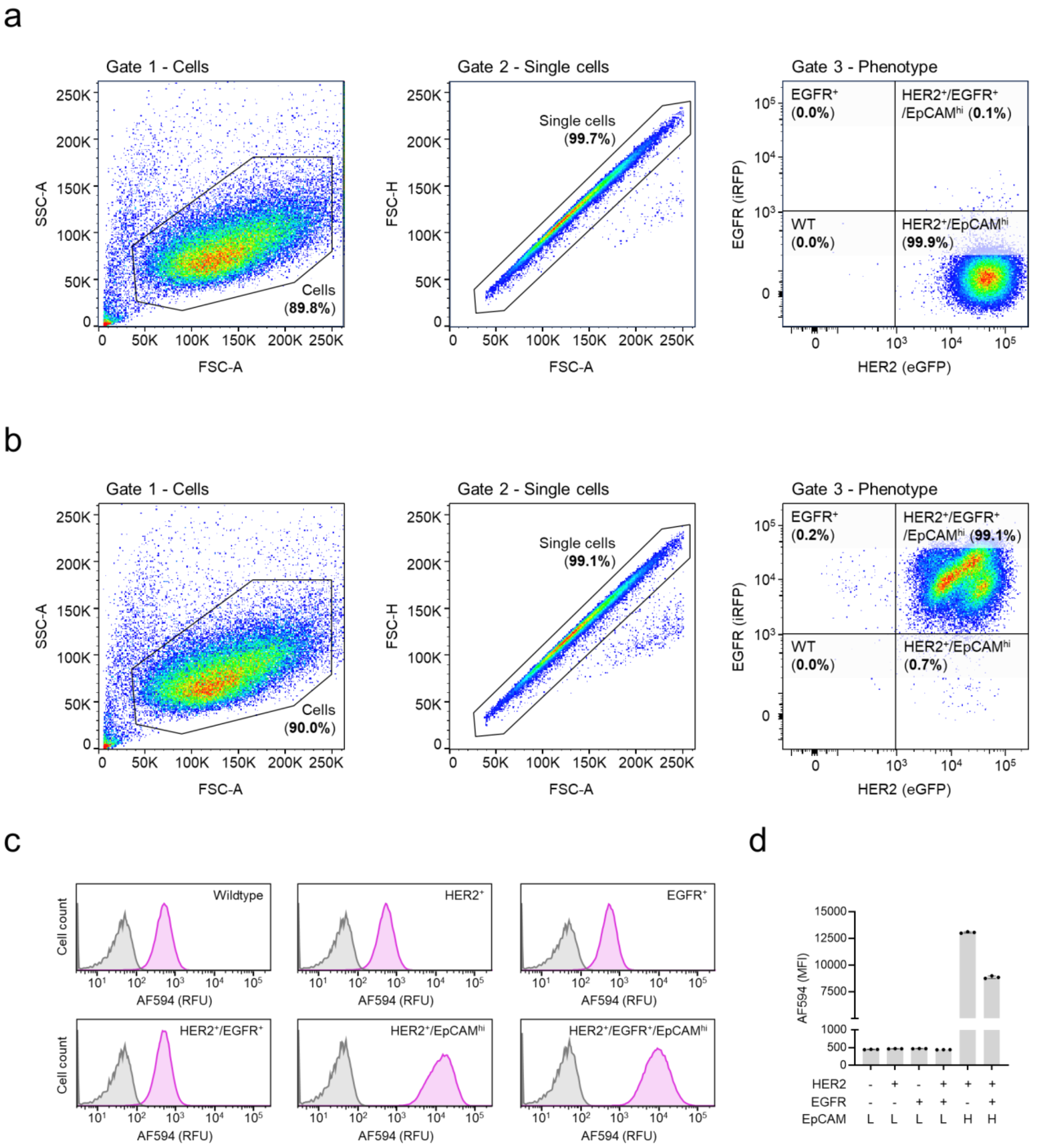


**Supplementary Figure 8. Phenotyping K562 cell lines with high ectopic levels of EpCAM.** Displayed is the flow cytometry data for **a**, K562$^{HER2+/EpCAMhi}$ and **b**, K562$^{HER2+/EGFR+/EpCAMhi}$ cell lines validating their HER2-eGFP/EGFR-iRFP phenotypes. Gating as in Supplemental figure 5. **c**, K562 cell lines were profiled individually for their relative surface level of EpCAM. Cells were treated with anti-EpCAM DARPin labeled with AF594 followed by flow cytometry analysis. Unstained cells are shown in grey, while stained cells are in magenta. **d**, Quantification of the data shown in panel **c**. The antigen profile of each cell line is indicated below each bar plot (L and H designates low endogenous (MFI < 1000) and high ectopic levels (MFI ≥ 1000) respectively for EpCAM). Flow cytometry data is presented as the mean of the AF594 median fluorescence intensities (MFI) with error bars signifying the standard error mean (n = 3 independent biological replicates).

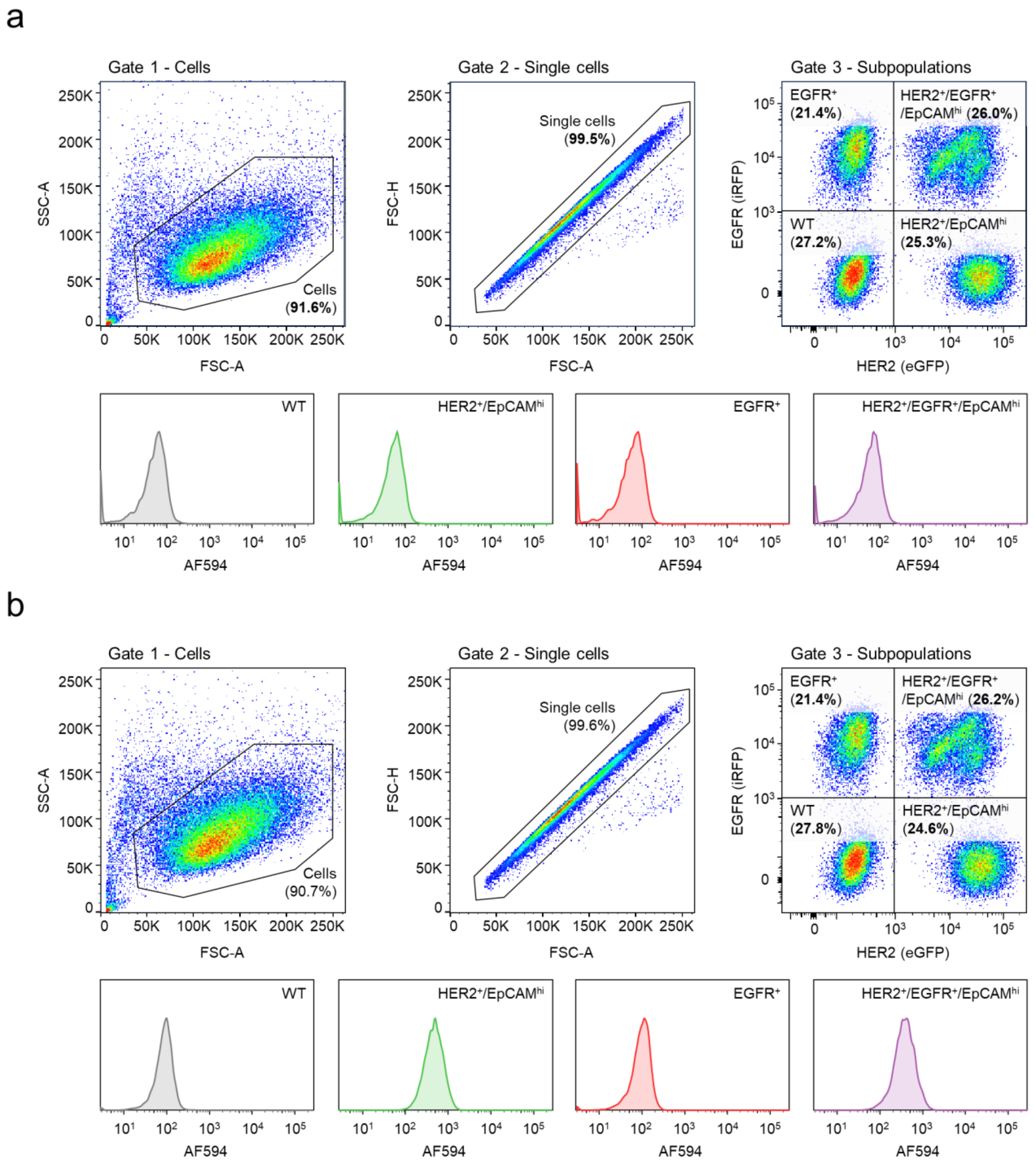


**Supplementary Figure 9. Flow cytometry gating strategy for mixed K562 population 2 flow cytometry experiments.** Example of flow cytometry gating strategy for mixed K562 population 2 (equal amounts of K562 (WT), K562$^{HER2+/EpCAMhi}$, K562$^{EGFR+}$, and K562$^{HER2+/EGFR+/EpCAMhi}$ cells). **a,** The mixed population was treated with SpyTag003-AF594 (100 nM) for 20 min and analyzed by flow cytometry. Three gates were used to define the cell population: Gate 1 was set as forward scatter area (FSC-A) versus side scatter area (SSC-A) to identify cells; gate 2 was set as forward scatter area (FSC-A) versus forward scatter height (FSC-H) to isolate single cells; gate 3 was set as eGFP fluorescence versus iRFP fluorescence to categorize K562 subpopulations by their HER2-eGFP and EGFR-iRFP phenotypes (their distribution is shown in percentages). Below the three gating plots are shown the AF594 histograms associated with each individual subpopulation. **b,** The mixed population was treated with αHER2-SpyN (100 nM) and SpyC-αEpCAM (100 nM) for 2 hr, followed by SpyTag003 labeled with Alexa Fluor 594 (SpyTag003-AF594) for 20 min. The cell mixture was then analyzed by flow cytometry as described in panel **a**.

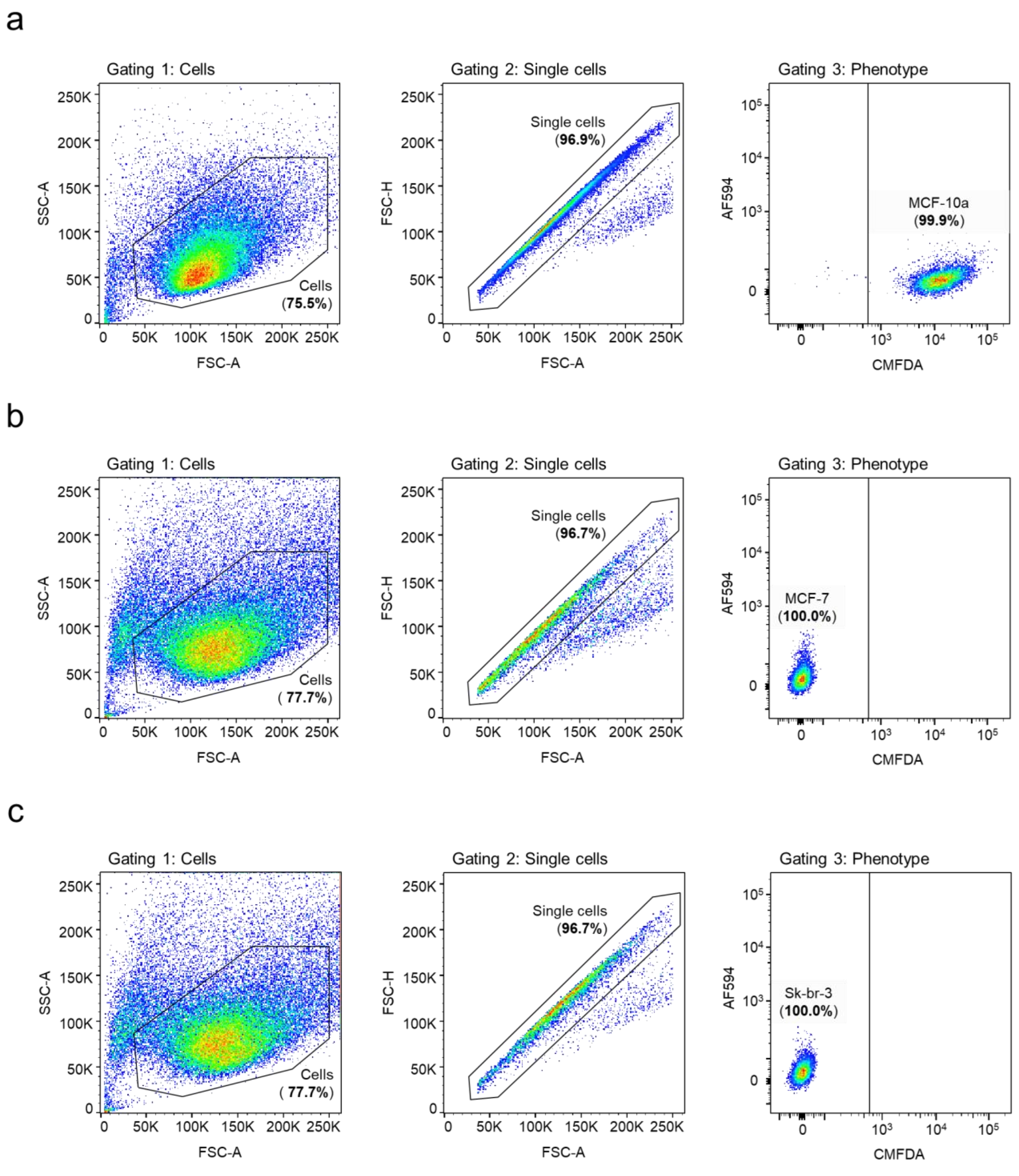


**Supplementary Figure 10. Flow cytometry of individual mammary cell lines.** Displayed is the flow cytometry data for **a**, MCF-10a (stained with CMFDA), **b**, MCF-7, and **c**, Sk-br-3 validating their CMFDA/AF594 phenotype. Three gates were used to define the cell population: Gate 1 was set as forward scatter area (FSC-A) versus side scatter area (SSC-A) to identify cells; gate 2 was set as forward scatter area (FSC-A) versus forward scatter height (FSC-H) to isolate single cells; gate 3 was set as CMFDA fluorescence versus AF594 fluorescence to categorize mammary subpopulations by their CMFDA/AF594 phenotype (their distribution is shown in percentages).

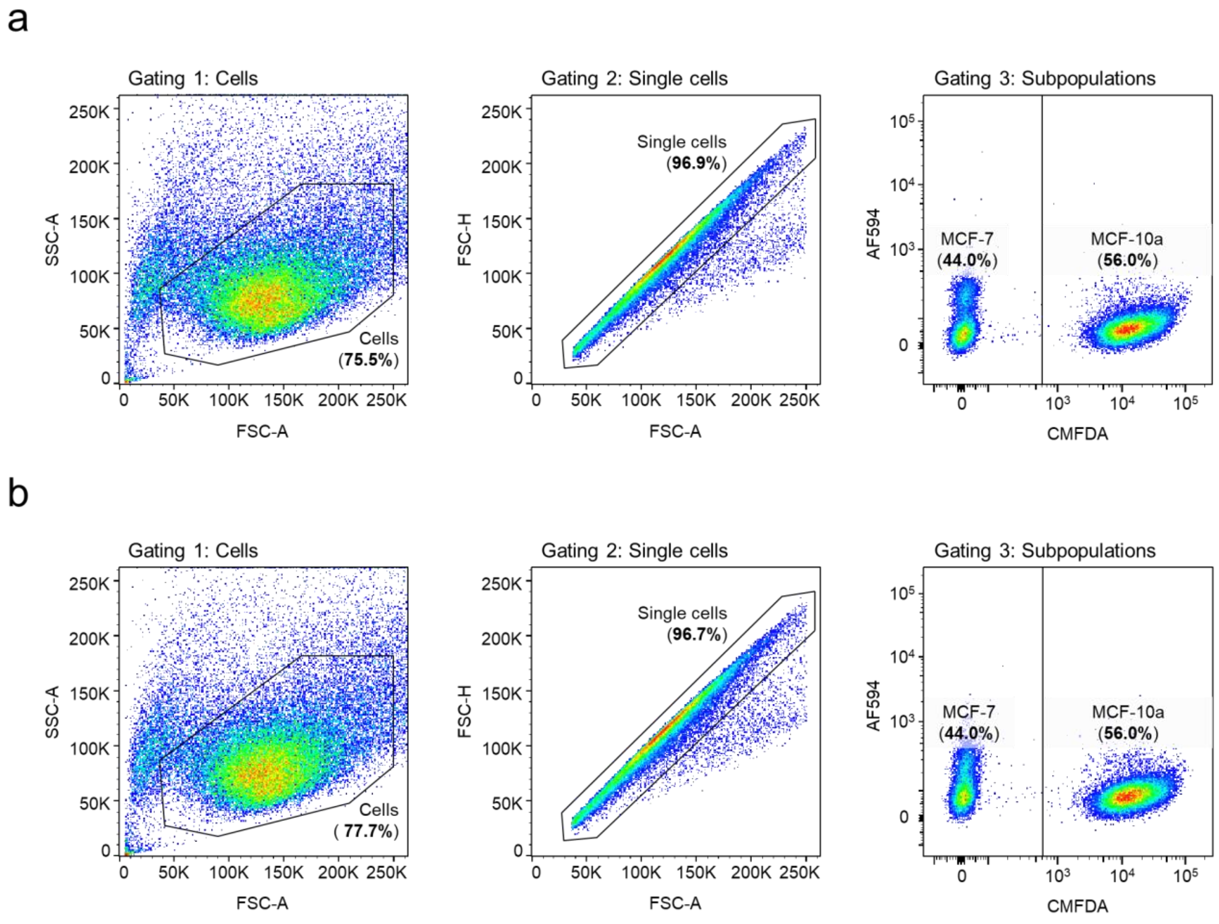


**Supplementary Figure 11. Flow cytometry gating strategy for mixed mammary population 1 flow cytometry experiments.** Example of flow cytometry gating strategy for mixed mammary population 1 (equal amounts of MCF-10a and MCF-7 cells). **a**, The mixed population was treated with SpyTag003-AF594 (100 nM) for 20 min and analyzed by flow cytometry. Gating as in Supplemental figure 10. **b,** The mixed population was treated with αHER2-SpyN (100 nM) and SpyC-αEpCAM (100 nM) for 2 hr, followed by SpyTag003 labeled with Alexa Fluor 594 (SpyTag003-AF594) for 20 min. The cell mixture was then analyzed by flow cytometry as described in panel **a**.

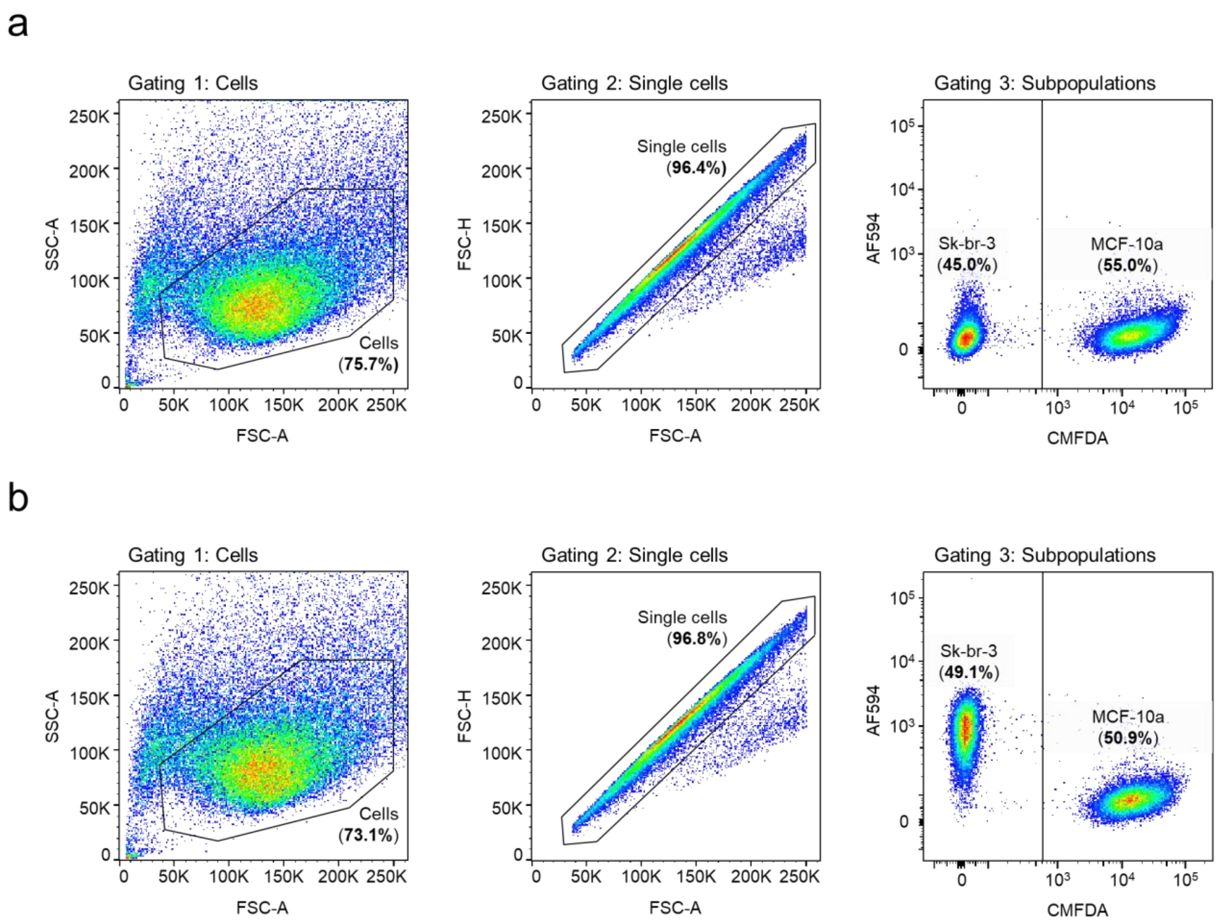


**Supplementary Figure 12. Flow cytometry gating strategy for mixed mammary population 2 flow cytometry experiments.** Example of flow cytometry gating strategy for mixed mammary population 2 (equal amounts of MCF-10a and Sk-br-3 cells). **a**, The mixed population was treated with SpyTag003-AF594 (100 nM) for 20 min and analyzed by flow cytometry. Gating as in Supplemental figure 10. **b,** The mixed population was treated with αHER2-SpyN (100 nM) and SpyC-αEpCAM (100 nM) for 2 hr, followed by SpyTag003 labeled with Alexa Fluor 594 (SpyTag003-AF594) for 20 min. The cell mixture was then analyzed by flow cytometry as described in panel **a**.

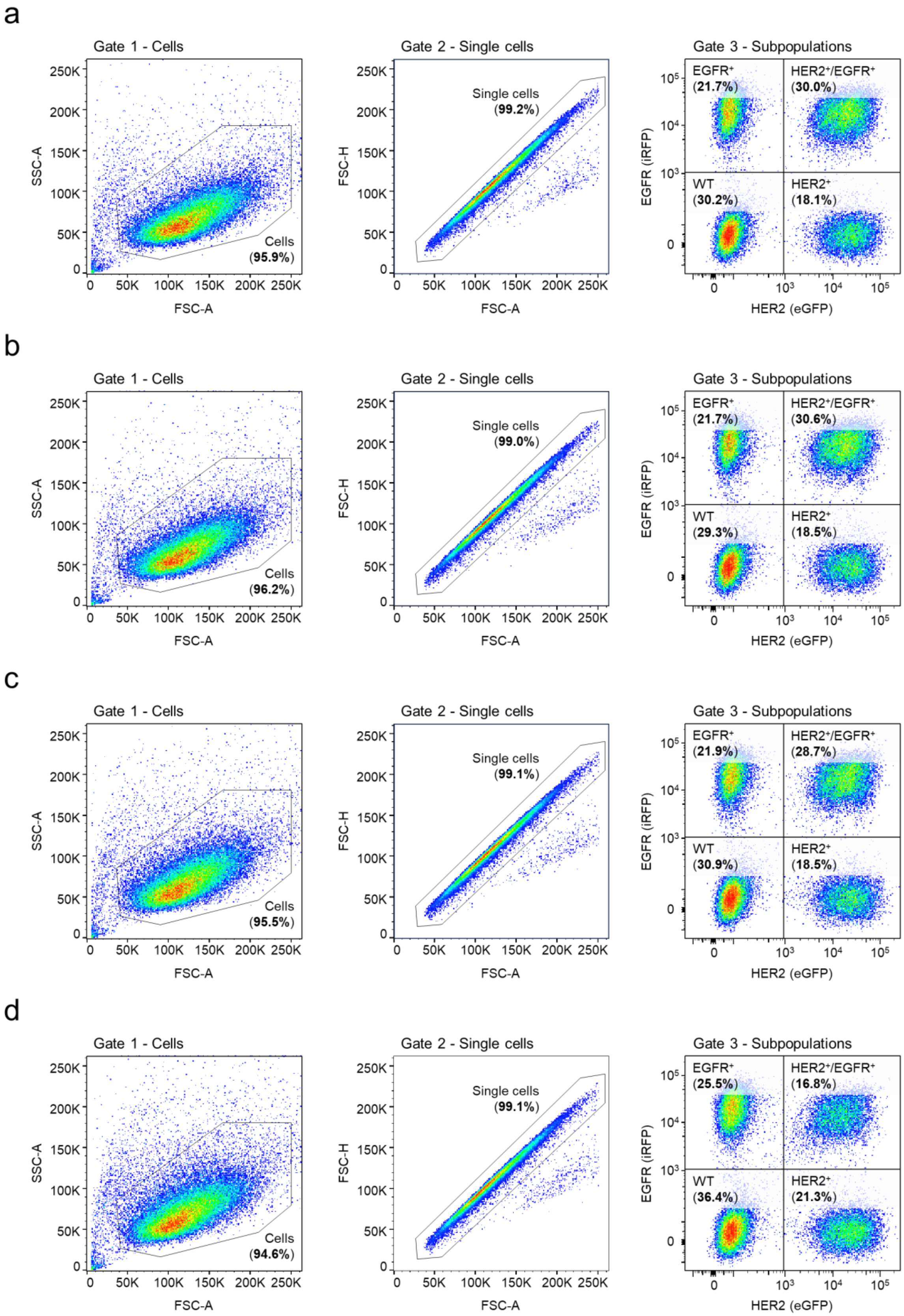


**Supplementary Figure 13. K562 depletion experiments with a one-dose regimen.** Flow cytometry analysis of mixed K562 population 1 (K562 (wildtype), K562$^{HER2+}$, K562$^{EGFR+}$, and K562$^{HER2+/EGFR+}$). Gating as in Supplemental figure 5. **a**, The cell mixture was cultured in complete media for 72 hr and analyzed by flow cytometry. **b**, The cell mixture was treated with SpyTag003-biotin (100 nM) for 20 min, washed before and then cultured in complete media supplemented with a Streptavidin-Saporin disulfide conjugate (Strep-Saporin, 20 nM) for 72 hr. The cells were subsequently analyzed by flow cytometry to determine the subpopulation distribution. **c**, The cell mixture was treated with SMART-SpyCatcher (eNrdJ-1, 100 nM) operating by [HER2 AND EGFR]. Following washing, the cell mixture was cultured in complete media for 72 hr and then analyzed as described above. **d**, The cell mixture was treated with SMART-SpyCatcher (eNrdJ-1, 100 nM) operating by [HER2 AND EGFR] for 2 hr, followed by SpyTag003-biotin for 20 min. Following washing, the cell mixture was cultured in complete media supplemented with Strep-Saporin (20 nM) for 72 hr and then analyzed by flow cytometry.

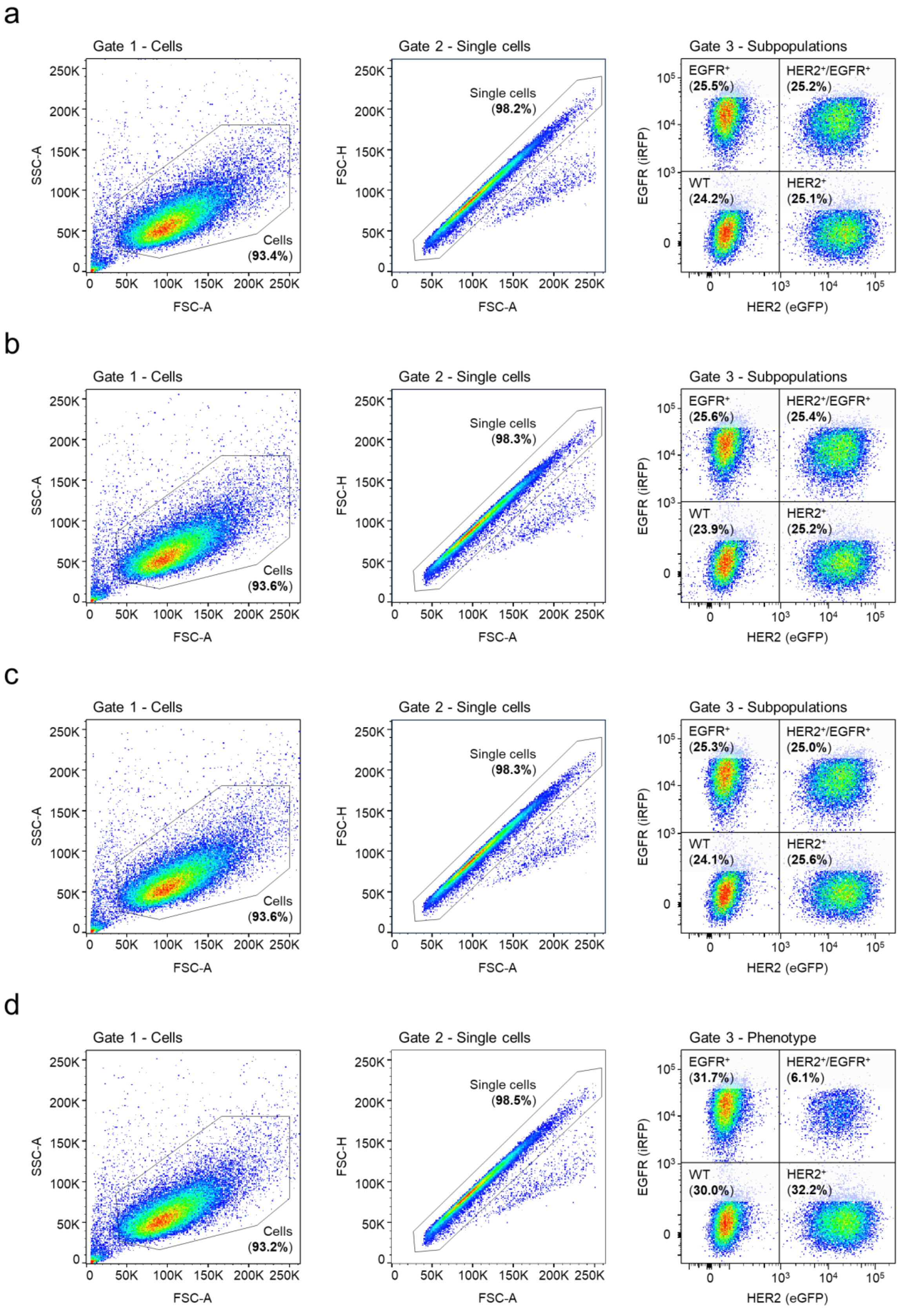


**Supplementary Figure 14. K562 depletion experiments with a two-dose regimen.** Flow cytometry analysis of mixed K562 population 1 (K562 (wildtype), K562$^{HER2+}$, K562$^{EGFR+}$, and K562$^{HER2+/EGFR+}$). Gating as in Supplemental figure 5. **a**, The cell mixture was cultured in complete media and analyzed by flow cytometry. **b**, The cell mixture was treated with SpyTag003-biotin (100 nM) for 20 min, washed before and then cultured in complete media supplemented with a Streptavidin-Saporin disulfide conjugate (Strep-Saporin, 20 nM). After 24 hr the treatment was repeated, and the cells then cultured for 72 hr. The cells were then analyzed by flow cytometry to determine the subpopulation distribution. **c**, The cell mixture was treated with SMART-SpyCatcher (eNrdJ-1, 100 nM) operating by [HER2 AND EGFR]. Following washing, the cell mixture was cultured in complete media 24 hr. The treatment was then repeated, and the cells cultured for an additional 72 hr before being analyzed as described above. **d**, The cell mixture was treated with SMART-SpyCatcher (eNrdJ-1, 100 nM) operating by [HER2 AND EGFR] for 2 hr, followed by SpyTag003-biotin for 20 min. Following washing, the cell mixture was cultured in complete media supplemented with Strep-Saporin (20 nM) for 24 hr. The treatment was then repeated, and the cells cultured for an additional 72 hr before being analyzed by flow cytometry.

**Supplementary Table 1. Proteins used in this study.** Tabulated are the domain structures, calculated masses (Cal. mass) and the experimentally determined molar masses (Det. mass) for all proteins used in this study.

| **Protein** | **Domain structure** | **Cal. mass** | **Det. mass** |
|---|---|---:|---:|
| NrdJ-1 | SGG-NrdJ-1-SEI | 16.918.0 | 16.917.5 |
| SpyCatcher003 | FLAG-αHER2-SpyCatcher003-αEGFR-Myc | 45,316.0 | 45,316.0 |
| SpyTag003-Cys | $His_6$-SpyTag003-SUMO-Cys | 15,591.0 | 15,591.6 |
| SpyTag003-AF594 | $His_6$-SpyTag003-SUMO-Cys Alexa Fluor 594 conjugate | 16,478.5 | 16,478.5 |
| SpyTag003-biotin | $His_6$-SpyTag003-SUMO-Cys biotin conjugate | 16,043.0 | 16,043.0 |
| SpyTag003-DNP | $His_6$-SpyTag003-SUMO-Cys dinitrophenol conjugate | 16,087.7 | 16,087.0 |
| SpyTag003$^{D117A}$-Cys | $His_6$-SpyTag003$^{D117A}$-SUMO-Cys | 15,547.5 | 15,547.1 |
| SpyTag003$^{D117A}$-AF594 | $His_6$-SpyTag003$^{D117A}$-SUMO-Cys Alexa Fluor 594 conjugate | 16,435.0 | 16,434.8 |
| SpyTag003-APEX2 | $His_6$-SpyTag003-HA-APEX2 | 32,493.0 | 32,492.0 |
| αHER2-Cys DARPin | αHER2-HA-Cys | 14,700.5 | 14,699.5 |
| αHER2-AF594 DARPin | αHER2-HA-Cys Alexa Fluor 594 conjugate | 15,587.5 | 15,586.0 |
| αEGFR-Cys DARPin | αEGFR-HA-Cys | 18,397.5 | 18,396.2 |
| αEGFR-AF594 DARPin | αEGFR-HA-Cys Alexa Fluor 594 conjugate | 19,284.5 | 19,284.0 |
| αEpCAM-Cys DARPin | αEpCAM-HA-Cys | 18,452.7 | 18,451.5 |
| αEpCAM-AF594 DARPin | αEpCAM-HA-Cys Alexa Fluor 594 conjugate | 19,339.7 | 19,339.0 |
| FLAG-SpyN$^{1-24}$-NpuN$^{cage}$-FKBP | FLAG-αHER2-SpyN$^{1-24}$-NpuN$^{cage}$-FKBP | 50,637.8 | 50,637.0 |
| FLAG-SpyN$^{1-42}$-NpuN$^{cage}$-FKBP | FLAG-αHER2-SpyN$^{1-42}$-NpuN$^{cage}$-FKBP | 52,650.1 | 52,648.3 |
| FLAG-SpyN$^{1-55}$-NpuN$^{cage}$-FKBP | FLAG-αHER2-SpyN$^{1-55}$-NpuN$^{cage}$-FKBP | 54,056.6 | 54,055.2 |
| FLAG-SpyN$^{1-73}$-NpuN$^{cage}$-FKBP | FLAG-αHER2-SpyN$^{1-73}$-NpuN$^{cage}$-FKBP | 56,228.1 | 56,227.8 |
| FLAG-SpyN$^{1-82}$-NpuN$^{cage}$-FKBP | FLAG-αHER2-SpyN$^{1-82}$-NpuN$^{cage}$-FKBP | 57,160.1 | 57,158.6 |
| FLAG-SpyN$^{1-90}$-NpuN$^{cage}$-FKBP | FLAG-αHER2-SpyN$^{1-90}$-NpuN$^{cage}$-FKBP | 57,991.0 | 57,989.9 |
| FRB-NpuC$^{cage}$-SpyC$^{25-113}$-Myc | FRB-NpuC$^{cage}$-SpyC$^{25-113}$-αEGFR-Myc | 52,518.6 | 52,517.0 |
| FRB-NpuC$^{cage}$-SpyC$^{43-113}$-Myc | FRB-NpuC$^{cage}$-SpyC$^{43-113}$-αEGFR-Myc | 50,506.4 | 50,504.1 |
| FRB-NpuC$^{cage}$-SpyC$^{56-113}$-Myc | FRB-NpuC$^{cage}$-SpyC$^{56-113}$-αEGFR-Myc | 49,099.8 | 49,101.1 |
| FRB-NpuC$^{cage}$-SpyC$^{74-113}$-Myc | FRB-NpuC$^{cage}$-SpyC$^{74-113}$-αEGFR-Myc | 46,928.4 | 46,927.1 |
| FRB-NpuC$^{cage}$-SpyC$^{83-113}$-Myc | FRB-NpuC$^{cage}$-SpyC$^{83-113}$-αEGFR-Myc | 46,040.4 | 46,038.6 |
| FRB-NpuC$^{cage}$-SpyC$^{91-113}$-Myc | FRB-NpuC$^{cage}$-SpyC$^{91-113}$-αEGFR-Myc | 45,137.4 | 45,135.0 |
| FLAG-SpyN$^{1-73}$-NrdJ-1N$^{cage}$-FKBP | FLAG-αHER2-SpyN$^{1-73}$-NrdJ-1N$^{cage}$-FKBP | 52,197.4 | 52,196.6 |
| FLAG-SpyN$^{1-73}$-NrdJ-1N(C1A)$^{cage}$-FKBP | FLAG-αHER2-SpyN$^{1-73}$-NrdJ-1N(C1A)$^{cage}$-FKBP | 52,161.3 | 52,159.4 |
| FLAG-SpyN$^{1-73}$-NrdJ-1N(C76V)$^{cage}$-FKBP | FLAG-αHER2-SpyN$^{1-73}$-NrdJ-1N(C76V)$^{cage}$-FKBP | 52,193.3 | 52,191.0 |
| FRB-NrdJ-1C$^{cage}$-SpyC$^{74-113}$-Myc | FRB-NrdJ-1C$^{cage}$-SpyC$^{74-113}$-αEGFR-Myc | 46,364.5 | 46,364.6 |
| FRB-NrdJ-1C$^{cage(C76V)}$-SpyC$^{74-113}$-Myc | FRB-NrdJ-1C$^{cage(C76V)}$-SpyC$^{74-113}$-αEGFR-Myc | 46,360.5 | 46,359.0 |
| αHER2-SpyN-NrdJ-1N$^{cage(C76V)}$ | FLAG-αHER2-SpyN$^{1-73}$-NrdJ-1N(C76V)$^{cage}$ | 40,062.5 | 40,062.0 |
| αHER2-SpyN-eNrdJ-1N$^{cage}$ | FLAG-αHER2-SpyN$^{1-73}$-NrdJ-1N(C76V)$^{cage(K104EK119A)}$ | 40,005.4 | 40.004.4 |
| αHER2-SpyN-eNrdJ-1N$^{cage}$ Cys1-alkylated | FLAG-αHER2-SpyN$^{1-73}$-NrdJ-1N(C76V)$^{cage(K104EK119A)}$ Cys1-alkylated | 40,063.0 | 40,062.0 |
| αEGFR-SpyN-eNrdJ-1N$^{cage}$ | FLAG-αEGFR-SpyN$^{1-73}$-NrdJ-1N(C76V)$^{cage(K104EK119A)}$ | 43,702.3 | 43,701.5 |
| αEpCAM-SpyN-eNrdJ-1N$^{cage}$ | FLAG-αEpCAM-SpyN$^{1-73}$- NrdJ-1N(C76V)$^{cage(K104EK119A)}$ | 43,757.6 | 43,756.4 |
| NrdJ-1C$^{cage(C76V)}$-SpyC-αEGFR | NrdJ-1C$^{cage(C76V)}$-SpyC$^{74-113}$-αEGFR-Myc | 34,437.9 | 34,437.0 |
| eNrdJ-1C$^{cage}$-SpyC-αHER2 | NrdJ-1C$^{cage(D66KC76V)}$-SpyC$^{74-113}$-αHER2-Myc | 30,753.1 | 30,752.0 |
| eNrdJ-1C$^{cage}$-SpyC-αEGFR | NrdJ-1C$^{cage(D66KC76V)}$-SpyC$^{74-113}$-αEGFR-Myc | 34,450.0 | 34,449.8 |
| eNrdJ-1C$^{cage}$-SpyC-αEpCAM | NrdJ-1C$^{cage(D66KC76V)}$-SpyC$^{74-113}$-αEpCAM-Myc | 34,505.3 | 34,505.2 |
| αHER2-SpyN-eNrdJ-1N$^{cage(K114A)}$ | FLAG-αHER2-SpyN$^{1-42}$-NrdJ-1N(C76V)$^{cage(K104EK114AK119A)}$ | 39,948.3 | 39,947.3 |
| αHER2-SpyN-eNrdJ-1N$^{cage(K116A)}$ | FLAG-αHER2-SpyN$^{1-42}$-NrdJ-1N(C76V)$^{cage(K104EK116AK119A)}$ | 39,948.3 | 39,946.8 |
| αHER2-SpyN-eNrdJ-1N$^{cage(R118A)}$ | FLAG-αHER2-SpyN$^{1-42}$-NrdJ-1N(C76V)$^{cage(K104ER118AK119A)}$ | 39,920.3 | 39,921.2 |
| αHER2-SpyN-eNrdJ-1N$^{cage(K114AK116A)}$ | FLAG-αHER2-SpyN$^{1-42}$-NrdJ-1N(C76V)$^{cage(K104EK114AK116AK119A)}$ | 39,891.1 | 39,892.2 |

| | | | |
|---|---|---|---|
| αHER2-SpyN-eNrdJ-1N$^{cage(K114AK116AR118A)}$ | FLAG-αHER2-SpyN$^{1-42}$-NrdJ-1N(C76V)$^{cage(K104EK114AK116AR118AK119A)}$ | 39,806.1 | 39,807.1 |
| αHER2-SpyN-eNrdJ-1N$^{cage(A119K)}$ | FLAG-αHER2-SpyN$^{1-42}$-NrdJ-1N(C76V)$^{cage(K104E)}$ | 40,062.4 | 40,063.0 |
| αEGFR-SpyN-eNrdJ-1N$^{cage(K114AK116A)}$ | FLAG-αEGFR-SpyN$^{1-42}$-NrdJ-1N(C76V)$^{cage(K104EK114AK116AK119A)}$ | 43,588.1 | 43,587.1 |
| αEpCAM-SpyN-eNrdJ-1N$^{cage(K114AK116A)}$ | FLAG-αEpCAM-SpyN$^{1-42}$-NrdJ-1N(C76V)$^{cage(K104EK114AK116AK119A)}$ | 43,643.4 | 43,641.8 |
| αHER2-SpyN-eNrdJ-1N$^{cage(1-39)}$ | FLAG-αHER2-SpyN$^{1-73}$-NrdJ-1N(C76V)$^{cage(K104EK119A_1-39)}$ | 40,468.0 | 40,467.0 |
| αHER2-SpyN-eNrdJ-1N$^{cage(1-38)}$ | FLAG-αHER2-SpyN$^{1-73}$-NrdJ-1N(C76V)$^{cage(K104EK119A_1-38)}$ | 40,330.8 | 40,331.4 |
| αHER2-SpyN-eNrdJ-1N$^{cage(1-37)}$ | FLAG-αHER2-SpyN$^{1-73}$-NrdJ-1N(C76V)$^{cage(K104EK119A_1-37)}$ | 40,231.7 | 40,231.2 |
| αHER2-SpyN-eNrdJ-1N$^{cage(1-36)}$ | FLAG-αHER2-SpyN$^{1-73}$-NrdJ-1N(C76V)$^{cage(K104EK119A_1-36)}$ | 40,118.5 | 40,118.3 |
| αHER2-SpyN-eNrdJ-1N$^{cage(1-33)}$ | FLAG-αHER2-SpyN$^{1-73}$-NrdJ-1N(C76V)$^{cage(K104EK119A_1-33)}$ | 39,776.2 | 39,775.3 |
| αHER2-SpyN-eNrdJ-1N$^{cage(1-32)}$ | FLAG-αHER2-SpyN$^{1-73}$-NrdJ-1N(C76V)$^{cage(K104EK119A_1-32)}$ | 39,705.1 | 39,704.7 |
| αHER2-SpyN-eNrdJ-1N$^{cage(1-31)}$ | FLAG-αHER2-SpyN$^{1-73}$-NrdJ-1N(C76V)$^{cage(K104EK119A_1-31)}$ | 39,557.9 | 39,556.8 |
| αHER2-SpyN-eNrdJ-1N$^{cage(1-30)}$ | FLAG-αHER2-SpyN$^{1-73}$-NrdJ-1N(C76V)$^{cage(K104EK119A_1-30)}$ | 39,410.7 | 39,410.2 |
| αHER2-SpyN-eNrdJ-1N$^{cage(1-29)}$ | FLAG-αHER2-SpyN$^{1-73}$-NrdJ-1N(C76V)$^{cage(K104EK119A_1-29)}$ | 39,296.6 | 39,295.6 |
| αHER2-SpyN-eNrdJ-1N$^{cage(1-26)}$ | FLAG-αHER2-SpyN$^{1-73}$-NrdJ-1N(C76V)$^{cage(K104EK119A_1-26)}$ | 38,971.3 | 38,970.1 |
| αHER2-SpyN-eNrdJ-1N$^{cage(1-20)}$ | FLAG-αHER2-SpyN$^{1-73}$-NrdJ-1N(C76V)$^{cage(K104EK119A_1-20)}$ | 38,249.5 | 38,248.6 |
| αHER2-SpyN-NrdJ-1N | FLAG-αHER2 -SpyN$^{1-73}$-NrdJ-1N | 34,426.4 | 34,425.2 |
| NrdJ-1C-SpyC-αEGFR | NrdJ-1C-SpyC$^{74-113}$-αEGFR-Myc | 27,334.1 | 27,333.7 |
| αEGFR-Decoy | FLAG-αEGFR-SpyN$^{1-74(K31E)}$-NrdJ-1N(C76V)$^{cage(K104EK114AK116AK119A)_1-29}$ | 42,880.3 | 42,881.0 |

**Supplementary Table 2. Crystallization data collection and refinement statistics.** Highest-resolution shell shown in parentheses. $R_{free}$ represents the R-factor calculated from 5% of the reflections not used during refinement. PDB ID: 8UBS.

| | |
|---|---|
| **Data collection** | |
| Wavelength (Å) | 0.97242 |
| Space group | P 21 21 21 |
| **Cell dimensions** | |
| a, b, c (Å) | 48.79, 84.38, 174.71 |
| β (degree) | 90 |
| Resolution (Å) | 29.12-1.95 |
| $R_{pim}$ | 0.053 (0.352) |
| I/σI | 1.66 |
| CC1/2 (%) | 99.7 (80.8) |
| Completeness (%) | 99.8 (98.1) |
| Anomalous completeness (%) | 98.4 (95.6) |
| Redundancy (%) | 6.8 (6.6) |
| Anomalous redundancy (%) | 3.5 (3.1) |
| Wilson B factor (Å2) | 26.5 |
| **Refinement** | |
| Resolution (Å) | 1.95 |
| Reflections/unique | 362053/53184 |
| $R_{work}$/$Rf_{ree}$ | 0.1801/0.2210 |
| **No. atoms / ions** | |
| Protein | 4800 |
| Other | 423 |
| Average B-factor (Å2) | 36 |
| **R.M.S. deviations** | |
| Bond lengths (Å) | 0.006 |
| Bond angles (degree) | 0.802 |
| **Ramachandran (%)** | |
| Favored | 98.2 |
| Allowed | 1.8 |

**Supplementary Table 3. Statistical analysis of flow cytometry data for AND gate experiments.** A one-way ANOVA followed by Dunnett's test was used to evaluate SMART-SpyCatcher targeting in complex mixtures of K562 cell lines. The control cell line used in each test is noted by ctrl, whereas ns denotes not significant; * denotes P < 0.05, ** denotes P < 0.01, *** denotes P < 0.001, **** denotes P < 0.0001. Cell lines not included in the specific mixed population are crossed out. K562 (wildtype), K562$^{EGFR+}$, K562$^{HER2+}$, and K562$^{HER2+/EGFR+}$ all have low endogenous levels of EpCAM.

| Panel | SMART-SpyCatcher | eNrdJ-1N$^{cage}$ variant | eNrdJ-1C$^{cage}$ variant | Mixed K562 population | K562 WT | K562 $^{EGFR+}$ | K562 $^{HER2+}$ | K562 $^{HER2+/EGFR+}$ | K562 $^{HER2+/EpCAMhi}$ | K562 $^{HER2/EGFR+/EpCAMhi}$ |
|---|---|---|---|---|---|---|---|---|---|---|
| Fig. 2b | αHER2-SpyN/SpyC-αEpCAM | Standard | Standard | 1 | ctrl | ns | **** | **** | | |
| Fig. 2b | αHER2-SpyN/SpyC-αEpCAM | Standard | Standard | 2 | ctrl | ns | | | **** | **** |
| Fig. 2b | αEpCAM-SpyN/SpyC-αHER2 | Standard | Standard | 1 | ctrl | ns | **** | **** | | |
| Fig. 2b | αEpCAM-SpyN/SpyC-αHER2 | Standard | Standard | 2 | ctrl | ns | | | **** | **** |
| Fig. 2b | αEGFR-SpyN/SpyC-αEpCAM | Standard | Standard | 1 | ctrl | **** | ns | **** | | |
| Fig. 2b | αEGFR-SpyN/SpyC-αEpCAM | Standard | Standard | 2 | ctrl | ** | | | ns | **** |
| Fig. 2b | αEpCAM-SpyN/SpyC-αEGFR | Standard | Standard | 1 | ctrl | **** | ns | **** | | |
| Fig. 2b | αEpCAM-SpyN/SpyC-αEGFR | Standard | Standard | 2 | ctrl | ** | | | ns | **** |
| Fig. 4d | αHER2-SpyN/SpyC-αEGFR | Standard | Standard | 1 | ctrl | ns | ns | **** | | |
| Ext. Fig. 3e | αHER2-SpyN/SpyC-αEGFR | Standard | Standard | 1 | ctrl | ns | ns | **** | | |
| Ext. Fig. 3e | αEGFR-SpyN/SpyC-αHER2 | Standard | Standard | 1 | ctrl | ns | ns | **** | | |
| Ext. Fig. 3e | αHER2-SpyN/SpyC-αHER2 | Standard | Standard | 1 | ctrl | ns | **** | **** | | |
| Ext. Fig. 3e | αEGFR-SpyN/SpyC-αEGFR | Standard | Standard | 1 | ctrl | **** | ns | **** | | |
| Ext. Fig. 4g | αHER2-SpyN/SpyC-αEGFR | +A119K | Standard | 1 | ctrl | ns | ns | **** | | |
| Ext. Fig. 4g | αHER2-SpyN/SpyC-αEGFR | Standard | Standard | 1 | ctrl | ns | ns | **** | | |
| Ext. Fig. 4g | αHER2-SpyN/SpyC-αEGFR | +K114A | Standard | 1 | ctrl | ns | ns | **** | | |
| Ext. Fig. 4g | αHER2-SpyN/SpyC-αEGFR | +K116A | Standard | 1 | ctrl | ns | ns | **** | | |
| Ext. Fig. 4g | αHER2-SpyN/SpyC-αEGFR | +R118A | Standard | 1 | ctrl | ns | ns | **** | | |
| Ext. Fig. 4g | αHER2-SpyN/SpyC-αEGFR | +K114A K116A | Standard | 1 | ctrl | ns | ns | **** | | |
| Ext. Fig. 4g | αHER2-SpyN/SpyC-αEGFR | +K114A K116A R118A | Standard | 1 | ctrl | ns | ns | **** | | |
| Ext. Fig. 5e | αHER2-SpyN/SpyC-αEGFR | 1-39 | Standard | 1 | ctrl | ns | ns | **** | | |
| Ext. Fig. 5e | αHER2-SpyN/SpyC-αEGFR | 1-38 | Standard | 1 | ctrl | ns | ns | **** | | |
| Ext. Fig. 5e | αHER2-SpyN/SpyC-αEGFR | 1-37 | Standard | 1 | ctrl | ns | ns | **** | | |
| Ext. Fig. 5e | αHER2-SpyN/SpyC-αEGFR | 1-36 | Standard | 1 | ctrl | ns | ns | **** | | |
| Ext. Fig. 5e | αHER2-SpyN/SpyC-αEGFR | 1-35 | Standard | 1 | ctrl | ns | ns | **** | | |
| Ext. Fig. 5e | αHER2-SpyN/SpyC-αEGFR | 1-33 | Standard | 1 | ctrl | ns | ns | **** | | |
| Ext. Fig. 5e | αHER2-SpyN/SpyC-αEGFR | 1-32 | Standard | 1 | ctrl | ns | ns | **** | | |
| Ext. Fig. 5e | αHER2-SpyN/SpyC-αEGFR | 1-31 | Standard | 1 | ctrl | ns | ns | **** | | |
| Ext. Fig. 5e | αHER2-SpyN/SpyC-αEGFR | 1-30 | Standard | 1 | ctrl | ns | ns | **** | | |
| Ext. Fig. 5e | αHER2-SpyN/SpyC-αEGFR | 1-29 | Standard | 1 | ctrl | ns | ns | **** | | |
| Ext. Fig. 5e | αHER2-SpyN/SpyC-αEGFR | 1-26 | Standard | 1 | ctrl | ns | ns | **** | | |
| Ext. Fig. 5e | αHER2-SpyN/SpyC-αEGFR | 1-20 | Standard | 1 | ctrl | ns | ns | **** | | |
| Ext. Fig. 8b | αHER2-SpyN/SpyC-αEpCAM | Standard | Standard | 1 | ctrl | ns | **** | **** | | |
| Ext. Fig. 8b | αHER2-SpyN/SpyC-αEpCAM | Standard | Standard | 2 | ctrl | ns | | | **** | **** |
| Ext. Fig. 9c | αHER2-SpyN/SpyC-αEGFR | Standard | Standard | 1 | ctrl | ns | **** | **** | | |

**Supplementary Table 4. Flow cytometry data for electrostatically tuned eNrdJ-1cage.** The mean AF594 median fluorescence intensity (MFI) was normalized to that collected for K562HER2+/EGFR+ using SMART-SpyCatcher employing the standard eNrdJ-1cage pair (n = 3 independent biological replicates; SEM denotes standard error mean).

| Panel | SMART-SpyCatcher | eNrdJ-1Ncage variant | eNrdJ-1Ccage variant | K562 WT | | K562HER2+ | | K562EGFR+ | | K562HER2+/EGFR+ | |
|---|---|---|---|---|---|---|---|---|---|---|---|
| | | | | Mean MFI (%) | ± SEM (%) | Mean MFI (%) | ± SEM (%) | Mean MFI (%) | ± SEM (%) | Mean MFI (%) | ± SEM (%) |
| Ext. Fig. 4g | αHER2-SpyN /SpyC-αEGFR | +A119K | Standard | 0 | 0 | 0 | 0 | 0 | 0 | 78 | 2 |
| Ext. Fig. 4g | αHER2-SpyN /SpyC-αEGFR | Standard | Standard | 0 | 0 | 0 | 0 | 0 | 0 | 100 | 1 |
| Ext. Fig. 4g | αHER2-SpyN /SpyC-αEGFR | +K114A | Standard | 0 | 0 | 0 | 0 | 1 | 0 | 187 | 4 |
| Ext. Fig. 4g | αHER2-SpyN /SpyC-αEGFR | +K116A | Standard | 0 | 0 | 0 | 0 | 1 | 0 | 189 | 7 |
| Ext. Fig. 4g | αHER2-SpyN /SpyC-αEGFR | +R118A | Standard | 1 | 0 | 1 | 0 | 1 | 0 | 171 | 2 |
| Ext. Fig. 4g | αHER2-SpyN /SpyC-αEGFR | +K114A K116A | Standard | 1 | 0 | 1 | 0 | 2 | 0 | 249 | 3 |
| Ext. Fig. 4g | αHER2-SpyN /SpyC-αEGFR | +K114A K116A R118 | Standard | 1 | 0 | 1 | 0 | 2 | 0 | 241 | 5 |

**Supplementary Table 5. Flow cytometry data for toehold tuned eNrdJ-1cage.** The mean AF594 median fluorescence intensity (MFI) was normalized to that collected for K562HER2+/EGFR+ using SMART-SpyCatcher employing the standard eNrdJ-1cage pair (n = 3 independent biological replicates; SEM denotes standard error mean).

| Panel | SMART-SpyCatcher | eNrdJ-1Ncage variant | eNrdJ-1Ccage variant | K562 WT | | K562HER2+ | | K562EGFR+ | | K562HER2+/EGFR+ | |
|---|---|---|---|---|---|---|---|---|---|---|---|
| | | | | Mean MFI (%) | ± SEM (%) | Mean MFI (%) | ± SEM (%) | Mean MFI (%) | ± SEM (%) | Mean MFI (%) | ± SEM (%) |
| Ext. Fig. 5e | αHER2-SpyN /SpyC-αEGFR | 1-39 | Standard | 0 | 0 | 0 | 0 | 0 | 0 | 14 | 1 |
| Ext. Fig. 5e | αHER2-SpyN /SpyC-αEGFR | 1-38 | Standard | 0 | 0 | 0 | 0 | 0 | 0 | 40 | 2 |
| Ext. Fig. 5e | αHER2-SpyN /SpyC-αEGFR | 1-37 | Standard | 0 | 0 | 0 | 0 | 0 | 0 | 77 | 1 |
| Ext. Fig. 5e | αHER2-SpyN /SpyC-αEGFR | 1-36 | Standard | 0 | 0 | 0 | 0 | 0 | 0 | 82 | 4 |
| Ext. Fig. 5e | αHER2-SpyN /SpyC-αEGFR | 1-35 | Standard | 0 | 0 | 0 | 0 | 0 | 0 | 100 | 1 |
| Ext. Fig. 5e | αHER2-SpyN /SpyC-αEGFR | 1-33 | Standard | 0 | 0 | 0 | 0 | 0 | 0 | 94 | 3 |
| Ext. Fig. 5e | αHER2-SpyN /SpyC-αEGFR | 1-32 | Standard | 0 | 0 | 0 | 0 | 0 | 0 | 93 | 4 |

| | | | | | | | | | | | |
|---|---|---|---|---|---|---|---|---|---|---|---|
| Ext. Fig. 5e | αHER2-SpyN /SpyC-αEGFR | 1-31 | Standard | 0 | 0 | 0 | 0 | 0 | 0 | 111 | 4 |
| Ext. Fig. 5e | αHER2-SpyN /SpyC-αEGFR | 1-30 | Standard | 0 | 0 | 0 | 0 | 0 | 0 | 116 | 6 |
| Ext. Fig. 5e | αHER2-SpyN /SpyC-αEGFR | 1-29 | Standard | 0 | 1 | 0 | 1 | 0 | 1 | 155 | 10 |
| Ext. Fig. 5e | αHER2-SpyN /SpyC-αEGFR | 1-26 | Standard | 0 | 0 | 0 | 0 | 0 | 0 | 148 | 2 |
| Ext. Fig. 5e | αHER2-SpyN /SpyC-αEGFR | 1-20 | Standard | 0 | 0 | 0 | 0 | 0 | 0 | 150 | 7 |

**Supplementary Table 6. Statistical analysis of flow cytometry data for OR gate experiments.** A one-way ANOVA followed by Dunnett's test was used to evaluate SMART-SpyCatcher targeting in complex mixtures of K562 cell lines. The control cell line used in each test is noted by ctrl, whereas ns denotes not significant; * denotes P < 0.05, ** denotes P < 0.01, *** denotes P < 0.001, **** denotes P < 0.0001.

| Panel | SMART-SpyCatcher | eNrdJ-1N$^{cage}$ variant | eNrdJ-1C$^{cage}$ variant | Mixed K562 population | K562 WT | K562 $^{HER2+}$ | K562 $^{EGFR+}$ | K562 $^{HER2+/EGFR+}$ |
|---|---|---|---|---|---|---|---|---|
| Ext. Fig. 6c | αHER2-SpyN/αEGFR-SpyN /SpyC-αHER2/SpyC-αEGFR | Standard | Standard | 1 | ctrl | *** | **** | **** |

**Supplementary Table 7. Statistical analysis of flow cytometry data for AND/OR gate experiments.** A one-way ANOVA followed by Dunnett's test was used to evaluate SMART-SpyCatcher targeting in complex mixtures of K562 cell lines. The control cell line used in each test is noted by ctrl, whereas ns denotes not significant; * denotes P < 0.05, ** denotes P < 0.01, *** denotes P < 0.001, **** denotes P < 0.0001. Cell lines not included in the specific mixed population are crossed out. K562 (wildtype), $K562^{EGFR+}$, $K562^{HER2+}$, and $K562^{HER2+/EGFR+}$ all have low endogenous levels of EpCAM.

| Panel | SMART-SpyCatcher | eNrdJ-1N$^{cage}$ variant | eNrdJ-1C$^{cage}$ variant | Mixed K562 population | K562 WT | K562 EGFR+ | K562 HER2+ | K562 HER2+ /EGFR+ | K562 HER2+ /EpCAMhi | K562 HER2/EGFR+ /EpCAMhi |
|---|---|---|---|---|---|---|---|---|---|---|
| Fig. 2d | αHER2-SpyN /SpyC-αEGFR /SpyC-αEpCAM | Standard | Standard | 1 | ctrl | ns | ** | **** | X | X |
| Fig. 2d | αHER2-SpyN /SpyC-αEGFR /SpyC-αEpCAM | Standard | Standard | 2 | ctrl | ns | X | X | **** | **** |
| Fig. 2d | αEGFR-SpyN /SpyC-αHER2 /SpyC-αEpCAM | Standard | Standard | 1 | ctrl | ns | ns | **** | X | X |
| Fig. 2d | αEGFR-SpyN /SpyC-αHER2 /SpyC-αEpCAM | Standard | Standard | 2 | ctrl | * | X | X | ns | **** |
| Fig. 2d | αEpCAM-SpyN /SpyC-αEGFR /SpyC-αEpCAM | Standard | Standard | 1 | ctrl | ** | ns | * | X | X |
| Fig. 2d | αEpCAM-SpyN /SpyC-αEGFR /SpyC-αEpCAM | Standard | Standard | 2 | ctrl | *** | X | X | **** | **** |

**Supplementary Table 8. Flow cytometry data for antigen phenotyping.** Tabulated is the mean AF594 median fluorescence intensity (MFI) for each specified cell line using the specified DARPin AF594 conjugate (n = 3 independent biological replicates; SEM denotes standard error mean).

| Panel | Cell line | DARPin-AF594 | Mean AF594 (MFI) | ± SEM |
|---|---|---|---|---|
| Fig. 3a | OE19 | αHER2 | 14224 | 108 |
| Fig. 3a | OE19 | αEGFR | 402 | 5 |
| Fig. 3a | OE19 | αEpCAM | 12261 | 174 |
| Fig. 3a | A549 | αHER2 | 48 | 4 |
| Fig. 3a | A549 | αEGFR | 1345 | 6 |
| Fig. 3a | A549 | αEpCAM | 30 | 5 |
| Ext. Fig. 7a | A431 | αHER2 | 325 | 25 |
| Ext. Fig. 7a | A431 | αEGFR | 8148 | 36 |
| Ext. Fig. 7a | A431 | αEpCAM | 1992 | 59 |
| Ext. Fig. 7a | LoVo | αHER2 | 138 | 12 |
| Ext. Fig. 7a | LoVo | αEGFR | 428 | 9 |
| Ext. Fig. 7a | LoVo | αEpCAM | 2072 | 45 |
| Ext. Fig. 7a | Sk-br-3 | αHER2 | 13745 | 232 |

| | | | | |
|---|---|---|---|---|
| Ext. Fig. 7a | Sk-br-3 | αEGFR | 554 | 1 |
| Ext. Fig. 7a | Sk-br-3 | αEpCAM | 4147 | 14 |
| Ext. Fig. 7a | HCT-116 | αHER2 | 89 | 6 |
| Ext. Fig. 7a | HCT-116 | αEGFR | 383 | 10 |
| Ext. Fig. 7a | HCT-116 | αEpCAM | 12819 | 60 |

**Supplementary Table 9. Flow cytometry data for SpyTag003-AF594 recruitment.** Tabulated is the mean AF594 mean fluorescence intensity (MFI) obtained for SMART-SpyCatcher mediated SpyTag003-AF594 recruitment to the specified cell line when operating through the specified AND gate (n = 3 independent biological replicates; SEM denotes standard error mean).

| Panel | Cell line | Logic function | SMART-SpyCatcher (eNrdJ-1$^{cage}$) | Mean AF594 (MFI) | ± SEM |
|---|---|---|---|---|---|
| Fig. 3c | OE19 | [HER2 AND HER2] | αHER2-SpyN/SpyC-αHER2 | 871 | 43 |
| Fig. 3c | OE19 | [HER2 AND EGFR] | αHER2-SpyN/SpyC-αEGFR | 117 | 20 |
| Fig. 3c | OE19 | [HER2 AND EpCAM] | αHER2-SpyN/SpyC-αEpCAM | 1528 | 16 |
| Fig. 3c | OE19 | [EGFR AND HER2] | αEGFR-SpyN/SpyC-αHER2 | 98 | 1 |
| Fig. 3c | OE19 | [EGFR AND EGFR] | αEGFR-SpyN/SpyC-αEGFR | 72 | 3 |
| Fig. 3c | OE19 | [EGFR AND EpCAM] | αEGFR-SpyN/SpyC-αEpCAM | 143 | 5 |
| Fig. 3c | OE19 | [EpCAM AND HER2] | αEpCAM-SpyN/SpyC-αHER2 | 1217 | 12 |
| Fig. 3c | OE19 | [EpCAM AND EGFR] | αEpCAM-SpyN/SpyC-αEGFR | 78 | 3 |
| Fig. 3c | OE19 | [EpCAM AND EpCAM] | αEpCAM-SpyN/SpyC-αEpCAM | 462 | 5 |
| Fig. 3c | A549 | [HER2 AND HER2] | αHER2-SpyN/SpyC-αHER2 | 3 | 4 |
| Fig. 3c | A549 | [HER2 AND EGFR] | αHER2-SpyN/SpyC-αEGFR | 50 | 4 |
| Fig. 3c | A549 | [HER2 AND EpCAM] | αHER2-SpyN/SpyC-αEpCAM | 18 | 11 |
| Fig. 3c | A549 | [EGFR AND HER2] | αEGFR-SpyN/SpyC-αHER2 | 9 | 6 |
| Fig. 3c | A549 | [EGFR AND EGFR] | αEGFR-SpyN/SpyC-αEGFR | 142 | 28 |
| Fig. 3c | A549 | [EGFR AND EpCAM] | αEGFR-SpyN/SpyC-αEpCAM | 18 | 2 |
| Fig. 3c | A549 | [EpCAM AND HER2] | αEpCAM-SpyN/SpyC-αHER2 | 45 | 55 |
| Fig. 3c | A549 | [EpCAM AND EGFR] | αEpCAM-SpyN/SpyC-αEGFR | 43 | 40 |
| Fig. 3c | A549 | [EpCAM AND EpCAM] | αEpCAM-SpyN/SpyC-αEpCAM | 1 | 2 |
| Ext. Fig. 7a | A431 | [HER2 AND HER2] | αHER2-SpyN/SpyC-αHER2 | 4 | 5 |
| Ext. Fig. 7a | A431 | [HER2 AND EGFR] | αHER2-SpyN/SpyC-αEGFR | 53 | 54 |
| Ext. Fig. 7a | A431 | [HER2 AND EpCAM] | αHER2-SpyN/SpyC-αEpCAM | 30 | 2 |
| Ext. Fig. 7a | A431 | [EGFR AND HER2] | αEGFR-SpyN/SpyC-αHER2 | 103 | 19 |
| Ext. Fig. 7a | A431 | [EGFR AND EGFR] | αEGFR-SpyN/SpyC-αEGFR | 480 | 177 |
| Ext. Fig. 7a | A431 | [EGFR AND EpCAM] | αEGFR-SpyN/SpyC-αEpCAM | 473 | 23 |
| Ext. Fig. 7a | A431 | [EpCAM AND HER2] | αEpCAM-SpyN/SpyC-αHER2 | 48 | 22 |
| Ext. Fig. 7a | A431 | [EpCAM AND EGFR] | αEpCAM-SpyN/SpyC-αEGFR | 420 | 29 |
| Ext. Fig. 7a | A431 | [EpCAM AND EpCAM] | αEpCAM-SpyN/SpyC-αEpCAM | 65 | 4 |
| Ext. Fig. 7a | LoVo | [HER2 AND HER2] | αHER2-SpyN/SpyC-αHER2 | 0 | 0 |
| Ext. Fig. 7a | LoVo | [HER2 AND EGFR] | αHER2-SpyN/SpyC-αEGFR | 0 | 0 |
| Ext. Fig. 7a | LoVo | [HER2 AND EpCAM] | αHER2-SpyN/SpyC-αEpCAM | 0 | 0 |
| Ext. Fig. 7a | LoVo | [EGFR AND HER2] | αEGFR-SpyN/SpyC-αHER2 | 0 | 0 |
| Ext. Fig. 7a | LoVo | [EGFR AND EGFR] | αEGFR-SpyN/SpyC-αEGFR | 0 | 0 |
| Ext. Fig. 7a | LoVo | [EGFR AND EpCAM] | αEGFR-SpyN/SpyC-αEpCAM | 7 | 3 |
| Ext. Fig. 7a | LoVo | [EpCAM AND HER2] | αEpCAM-SpyN/SpyC-αHER2 | 2 | 3 |
| Ext. Fig. 7a | LoVo | [EpCAM AND EGFR] | αEpCAM-SpyN/SpyC-αEGFR | 14 | 2 |
| Ext. Fig. 7a | LoVo | [EpCAM AND EpCAM] | αEpCAM-SpyN/SpyC-αEpCAM | 18 | 2 |
| Ext. Fig. 7a | Sk-br-3 | [HER2 AND HER2] | αHER2-SpyN/SpyC-αHER2 | 1096 | 121 |
| Ext. Fig. 7a | Sk-br-3 | [HER2 AND EGFR] | αHER2-SpyN/SpyC-αEGFR | 206 | 7 |
| Ext. Fig. 7a | Sk-br-3 | [HER2 AND EpCAM] | αHER2-SpyN/SpyC-αEpCAM | 738 | 4 |
| Ext. Fig. 7a | Sk-br-3 | [EGFR AND HER2] | αEGFR-SpyN/SpyC-αHER2 | 200 | 19 |
| Ext. Fig. 7a | Sk-br-3 | [EGFR AND EGFR] | αEGFR-SpyN/SpyC-αEGFR | 92 | 6 |
| Ext. Fig. 7a | Sk-br-3 | [EGFR AND EpCAM] | αEGFR-SpyN/SpyC-αEpCAM | 228 | 6 |
| Ext. Fig. 7a | Sk-br-3 | [EpCAM AND HER2] | αEpCAM-SpyN/SpyC-αHER2 | 761 | 21 |

| | | | | | |
|---|---|---|---|---|---|
| Ext. Fig. 7a | Sk-br-3 | [EpCAM AND EGFR] | αEpCAM-SpyN/SpyC-αEGFR | 95 | 13 |
| Ext. Fig. 7a | Sk-br-3 | [EpCAM AND EpCAM] | αEpCAM-SpyN/SpyC-αEpCAM | 143 | 21 |
| Ext. Fig. 7a | HCT-116 | [HER2 AND HER2] | αHER2-SpyN/SpyC-αHER2 | 2 | 1 |
| Ext. Fig. 7a | HCT-116 | [HER2 AND EGFR] | αHER2-SpyN/SpyC-αEGFR | 2 | 1 |
| Ext. Fig. 7a | HCT-116 | [HER2 AND EpCAM] | αHER2-SpyN/SpyC-αEpCAM | 16 | 2 |
| Ext. Fig. 7a | HCT-116 | [EGFR AND HER2] | αEGFR-SpyN/SpyC-αHER2 | 6 | 2 |
| Ext. Fig. 7a | HCT-116 | [EGFR AND EGFR] | αEGFR-SpyN/SpyC-αEGFR | 2 | 2 |
| Ext. Fig. 7a | HCT-116 | [EGFR AND EpCAM] | αEGFR-SpyN/SpyC-αEpCAM | 26 | 5 |
| Ext. Fig. 7a | HCT-116 | [EpCAM AND HER2] | αEpCAM-SpyN/SpyC-αHER2 | 23 | 1 |
| Ext. Fig. 7a | HCT-116 | [EpCAM AND EGFR] | αEpCAM-SpyN/SpyC-αEGFR | 48 | 3 |
| Ext. Fig. 7a | HCT-116 | [EpCAM AND EpCAM] | αEpCAM-SpyN/SpyC-αEpCAM | 275 | 8 |

**Supplementary Table 10. Primary structures of expressed proteins.** The fully annotated amino acid sequences of overexpressed constructs used in this study are given (here shown with unprocessed N-terminal methionine and $His_6$-SUMO tag when relevant).

$His_6$-SUMO-NrdJ-1

SGGALVGSSEIITRNYGKTTIKEVVEIFDNDKNIQVLAFNTHTDNIEWAPIKAAQLTRPNAELVELEIDTLH
GVKTIRCTPDHPVYTKNRGYVRADELTDDDELVVAIMEAKTYIGKLKSRKIVSNEDTYDIQTSTHNFFAN
DILVHASEI

$His_6$-SUMO-SpyCatcher003

MGSSHHHHHHGSGLVPRGSASMSDSEVNQEAKPEVKPEVKPETHINLKVSDGSSEIFFKIKKTTPLRR
LMEAFAKRQGKEMDSLRFLYDGIRIQADQTPEDLDMEDNDIIEAHREQIGGDYKDDDDKDLGKKLLEAA
RAGQDDEVRILMANGADVNAKDEYGLTPLYLATAHGHLEIVEVLLKNGADVNAVDAIGFTPLHLAAFIGH
LEIAEVLLKHGADVNAQDKFGKTAFDISIGNGNEDLAEILQKLNGSTSGSVTTLSGLSGEQGPSGDMTT
EEDSATHIKFSKRDEDGRELAGATMELRDSSGKTISTWISDGHVKDFYLYPGKYTFVETAAPDGYEVAT
PIEFTVNEDGQVTVDGEATEGDAHTGSTSGSDLGKKLLEAARAGQDDEVRILMANGADVNADDTWG
WTPLHLAAYQGHLEIVEVLLKNGADVNAYDYIGWTPLHLAADGHLEIVEVLLKNGADVNASDYIGDTPL
HLAAHNGHLEIVEVLLKHGADVNAQDKFGKTAFDISIDNGNEDLAEILQKLNEQKLISEEDL

SpyTag003-Cys

MGSSHHHHHHSSGLVPRGSRGVPHIVMVDAYKRYKGSGESGMSDSEVNQEAKPEVKPEVKPETHIN
LKVSDGSSEIFFKIKKTTPLRRLMEAFAKRQGKEMDSLRFLYDGIRIQADQTPEDLDMEDNDIIEAHREQI
GC

SpyTag003$^{D117A}$-Cys

MGSSHHHHHHSSGLVPRGSRGVPHIVMVAAYKRYKGSGESGMSDSEVNQEAKPEVKPEVKPETHINL
KVSDGSSEIFFKIKKTTPLRRLMEAFAKRQGKEMDSLRFLYDGIRIQADQTPEDLDMEDNDIIEAHREQI
GC

SpyTag003-APEX2

MGSSHHHHHHSSGLVPRGSRGVPHIVMVDAYKRYKGSGESGYPYDVPDYASSGGKSYPTVSADYQD
AVEKAKKKLRGFIAEKRCAPLMLRLAFHSAGTFDKGTKTGGPFGTIKHPAELAHSANNGLDIAVRLLEPL
KAEFPILSYADFYQLAGVVAVEVTGGPKVPFHPGREDKPEPPPEGRLPDPTKGSDHLRDVFGKAMGLT
DQDIVALSGGHTIGAAHKERSGFEGPWTSNPLIFDNSYFTELLSGEKEGLLQLPSDKALLSDPVFRPLV
DKYAADEDAFFADYAEAHQKLSELGFADA

His$_6$-SUMO-αHER2-Cys DARPin

MGSSHHHHHHGSGLVPRGSASMSDSEVNQEAKPEVKPEVKPETHINLKVSDGSSEIFFKIKKTTPLRR
LMEAFAKRQGKEMDSLRFLYDGIRIQADQTPEDLDMEDNDIIEAHREQIGGDLGKKLLEAARAGQDDEV
RILMANGADVNAKDEYGLTPLYLATAHGHLEIVEVLLKNGADVNAVDAIGFTPLHLAAFIGHLEIAEVLLK
HGADVNAQDKFGKTAFDISIGNGNEDLAEILQKLNGGYPYDVPDYAGGSC

His$_6$-SUMO-αEGFR-Cys DARPin

MGSSHHHHHHGSGLVPRGSASMSDSEVNQEAKPEVKPEVKPETHINLKVSDGSSEIFFKIKKTTPLRR
LMEAFAKRQGKEMDSLRFLYDGIRIQADQTPEDLDMEDNDIIEAHREQIGGDLGKKLLEAARAGQDDEV
RILMANGADVNADDTWGWTPLHLAAYQGHLEIVEVLLKNGADVNAYDYIGWTPLHLAADGHLEIVEVLL
KNGADVNASDYIGDTPLHLAAHNGHLEIVEVLLKHGADVNAQDKFGKTAFDISIDNGNEDLAEILQKLNG
GYPYDVPDYAGGSC

His$_6$-SUMO-αEpCAM-Cys DARPin

MGSSHHHHHHGSGLVPRGSASMSDSEVNQEAKPEVKPEVKPETHINLKVSDGSSEIFFKIKKTTPLRR
LMEAFAKRQGKEMDSLRFLYDGIRIQADQTPEDLDMEDNDIIEAHREQIGGDLGKKLLEAARAGQDDEV
RILVANGADVNAYFGTTPLHLAAAHGRLEIVEVLLKNGADVNAQDVWGITPLHLAAYNGHLEIVEVLLKY
GADVNAHDTRGWTPLHLAAINGHLEIVEVLLKNVADVNAQDRSGKTPFDLAIDNGNEDIAEVLQKAAKL
NGGYPYDVPDYAGGSC

His$_6$-SUMO-FLAG-SpyN$^{1\text{-}24}$-NpuN$^{cage}$-FKBP

MGSSHHHHHHGSGLVPRGSASMSDSEVNQEAKPEVKPEVKPETHINLKVSDGSSEIFFKIKKTTPLRR
LMEAFAKRQGKEMDSLRFLYDGIRIQADQTPEDLDMEDNDIIEAHREQIGGDYKDDDDKDLGKKLLEAA
RAGQDDEVRILMANGADVNAKDEYGLTPLYLATAHGHLEIVEVLLKNGADVNAVDAIGFTPLHLAAFIGH
LEIAEVLLKHGADVNAQDKFGKTAFDISIGNGNEDLAEILQKLNGSTSGSVTTLSGLSGEQGPSGDMTT
EEDSACLSYETEILTVEYGLLPIGKIVEKRIECTVYSVDNNGNIYTQPVAQWHDRGKQKVFEYCLEDGSL
IRATKDHKFMTVDGQMLPIKEIFRRKLDLMRVDNLPNGSGGENLYFQGENLYFQGGSGGIEIATEKYLG
EQNVYDIGVERDHNFALKNGGYFQGIEIATEKYLGEQNVYDIGVERDHNFALKNGNSAGGVQVETISP
GDGRTFPKRGQTCVVHYTGMLEDGKKFDSSRDRNKPFKFMLGKQEVIRGWEEGVAQMSVGQRAKLT
ISPDYAYGATGHPGIIPPHATLVFDVELLKLE

His$_6$-SUMO-FLAG-SpyN$^{1\text{-}42}$-NpuN$^{cage}$-FKBP

MGSSHHHHHHGSGLVPRGSASMSDSEVNQEAKPEVKPEVKPETHINLKVSDGSSEIFFKIKKTTPLRR
LMEAFAKRQGKEMDSLRFLYDGIRIQADQTPEDLDMEDNDIIEAHREQIGGDYKDDDDKDLGKKLLEAA
RAGQDDEVRILMANGADVNAKDEYGLTPLYLATAHGHLEIVEVLLKNGADVNAVDAIGFTPLHLAAFIGH
LEIAEVLLKHGADVNAQDKFGKTAFDISIGNGNEDLAEILQKLNGSTSGSVTTLSGLSGEQGPSGDMTT

EEDSATHIKFSKRDEDGRELAGACLSYETEILTVEYGLLPIGKIVEKRIECTVYSVDNNGNIYTQPVAQW
HDRGKQKVFEYCLEDGSLIRATKDHKFMTVDGQMLPIKEIFRRKLDLMRVDNLPNGSGGENLYFQGEN
LYFQGGSGGIEIATEKYLGEQNVYDIGVERDHNFALKNGGYFQGIEIATEKYLGEQNVYDIGVERDHNF
ALKNGNSAGGVQVETISPGDGRTFPKRGQTCVVHYTGMLEDGKKFDSSRDRNKPFKFMLGKQEVIRG
WEEGVAQMSVGQRAKLTISPDYAYGATGHPGIIPPHATLVFDVELLKLE

His$_6$-SUMO-FLAG-SpyN$^{1-55}$-NpuN$^{cage}$-FKBP
MGSSHHHHHHGSGLVPRGSASMSDSEVNQEAKPEVKPEVKPETHINLKVSDGSSEIFFKIKKTTPLRR
LMEAFAKRQGKEMDSLRFLYDGIRIQADQTPEDLDMEDNDIIEAHREQIGGDYKDDDDKDLGKKLLEAA
RAGQDDEVRILMANGADVNAKDEYGLTPLYLATAHGHLEIVEVLLKNGADVNAVDAIGFTPLHLAAFIGH
LEIAEVLLKHGADVNAQDKFGKTAFDISIGNGNEDLAEILQKLNGSTSGSVTTLSGLSGEQGPSGDMTT
EEDSATHIKFSKRDEDGRELAGATMELRDSSGKTISCLSYETEILTVEYGLLPIGKIVEKRIECTVYSVDN
NGNIYTQPVAQWHDRGKQKVFEYCLEDGSLIRATKDHKFMTVDGQMLPIKEIFRRKLDLMRVDNLPNG
SGGENLYFQGENLYFQGGSGGIEIATEKYLGEQNVYDIGVERDHNFALKNGGYFQGIEIATEKYLGEQN
VYDIGVERDHNFALKNGNSAGGVQVETISPGDGRTFPKRGQTCVVHYTGMLEDGKKFDSSRDRNKPF
KFMLGKQEVIRGWEEGVAQMSVGQRAKLTISPDYAYGATGHPGIIPPHATLVFDVELLKLE

His$_6$-SUMO-FLAG-SpyN$^{1-73}$-NpuN$^{cage}$-FKBP
MGSSHHHHHHGSGLVPRGSASMSDSEVNQEAKPEVKPEVKPETHINLKVSDGSSEIFFKIKKTTPLRR
LMEAFAKRQGKEMDSLRFLYDGIRIQADQTPEDLDMEDNDIIEAHREQIGGDYKDDDDKDLGKKLLEAA
RAGQDDEVRILMANGADVNAKDEYGLTPLYLATAHGHLEIVEVLLKNGADVNAVDAIGFTPLHLAAFIGH
LEIAEVLLKHGADVNAQDKFGKTAFDISIGNGNEDLAEILQKLNGSTSGSVTTLSGLSGEQGPSGDMTT
EEDSATHIKFSKRDEDGRELAGATMELRDSSGKTISTWISDGHVKDFYLYPGKYCLSYETEILTVEYGLL
PIGKIVEKRIECTVYSVDNNGNIYTQPVAQWHDRGKQKVFEYCLEDGSLIRATKDHKFMTVDGQMLPIK
EIFRRKLDLMRVDNLPNGSGGENLYFQGENLYFQGGSGGIEIATEKYLGEQNVYDIGVERDHNFALKN
GGYFQGIEIATEKYLGEQNVYDIGVERDHNFALKNGNSAGGVQVETISPGDGRTFPKRGQTCVVHYTG
MLEDGKKFDSSRDRNKPFKFMLGKQEVIRGWEEGVAQMSVGQRAKLTISPDYAYGATGHPGIIPPHAT
LVFDVELLKLE

His$_6$-SUMO-FLAG-SpyN$^{1-82}$-NpuN$^{cage}$-FKBP
MGSSHHHHHHGSGLVPRGSASMSDSEVNQEAKPEVKPEVKPETHINLKVSDGSSEIFFKIKKTTPLRR
LMEAFAKRQGKEMDSLRFLYDGIRIQADQTPEDLDMEDNDIIEAHREQIGGDYKDDDDKDLGKKLLEAA
RAGQDDEVRILMANGADVNAKDEYGLTPLYLATAHGHLEIVEVLLKNGADVNAVDAIGFTPLHLAAFIGH
LEIAEVLLKHGADVNAQDKFGKTAFDISIGNGNEDLAEILQKLNGSTSGSVTTLSGLSGEQGPSGDMTT
EEDSATHIKFSKRDEDGRELAGATMELRDSSGKTISTWISDGHVKDFYLYPGKYTFVETAAPDCLSYET
EILTVEYGLLPIGKIVEKRIECTVYSVDNNGNIYTQPVAQWHDRGKQKVFEYCLEDGSLIRATKDHKFMT

VDGQMLPIKEIFRRKLDLMRVDNLPNGSGGENLYFQGENLYFQGGSGGIEIATEKYLGEQNVYDIGVER
DHNFALKNGGYFQGIEIATEKYLGEQNVYDIGVERDHNFALKNGNSAGGVQVETISPGDGRTFPKRGQ
TCVVHYTGMLEDGKKFDSSRDRNKPFKFMLGKQEVIRGWEEGVAQMSVGQRAKLTISPDYAYGATG
HPGIIPPHATLVFDVELLKLE

$His_6$-SUMO-FLAG-$SpyN^{1-90}$-$NpuN^{cage}$-FKBP

MGSSHHHHHHGSGLVPRGSASMSDSEVNQEAKPEVKPEVKPETHINLKVSDGSSEIFFKIKKTTPLRR
LMEAFAKRQGKEMDSLRFLYDGIRIQADQTPEDLDMEDNDIIEAHREQIGGDYKDDDDKDLGKKLLEAA
RAGQDDEVRILMANGADVNAKDEYGLTPLYLATAHGHLEIVEVLLKNGADVNAVDAIGFTPLHLAAFIGH
LEIAEVLLKHGADVNAQDKFGKTAFDISIGNGNEDLAEILQKLNGSTSGSVTTLSGLSGEQGPSGDMTT
EEDSATHIKFSKRDEDGRELAGATMELRDSSGKTISTWISDGHVKDFYLYPGKYTFVETAAPDGYEVAT
PICLSYETEILTVEYGLLPIGKIVEKRIECTVYSVDNNGNIYTQPVAQWHDRGKQKVFEYCLEDGSLIRAT
KDHKFMTVDGQMLPIKEIFRRKLDLMRVDNLPNGSGGENLYFQGENLYFQGGSGGIEIATEKYLGEQN
VYDIGVERDHNFALKNGGYFQGIEIATEKYLGEQNVYDIGVERDHNFALKNGNSAGGVQVETISPGDG
RTFPKRGQTCVVHYTGMLEDGKKFDSSRDRNKPFKFMLGKQEVIRGWEEGVAQMSVGQRAKLTISP
DYAYGATGHPGIIPPHATLVFDVELLKLE

$His_6$-SUMO-FRB-$NpuC^{cage}$-$SpyC^{25-113}$-Myc

MGSSHHHHHHGSGLVPRGSASMSDSEVNQEAKPEVKPEVKPETHINLKVSDGSSEIFFKIKKTTPLRR
LMEAFAKRQGKEMDSLRFLYDGIRIQADQTPEDLDMEDNDIIEAHREQIGGGRVAILWHEMWHEGLEE
ASRLYFGERNVKGMFEVLEPLHAMMERGPQTLKETSFNQAYGRDLMEAQEWCRKYMKSGNVKDLLQ
AWDLYYHVFRRISGNGNGGEQEVFEYCLEDGSLIRATKDHKFMTVDGQMLPIDEIFERELDLMRVDNL
PNGSGGENLYFQGENLYFQGGSGGIKIATRKYLGKQNVYDIGVERDHNFALKNGFIASNCHIKFSKRDE
DGRELAGATMELRDSSGKTISTWISDGHVKDFYLYPGKYTFVETAAPDGYEVATPIEFTVNEDGQVTV
DGEATEGDAHTGSTSGSDLGKKLLEAARAGQDDEVRILMANGADVNADDTWGWTPLHLAAYQGHLEI
VEVLLKNGADVNAYDYIGWTPLHLAADGHLEIVEVLLKNGADVNASDYIGDTPLHLAAHNGHLEIVEVLL
KHGADVNAQDKFGKTAFDISIDNGNEDLAEILQKLNEQKLISEEDL

$His_6$-SUMO-FRB-$NpuC^{cage}$-$SpyC^{43-113}$-Myc

MGSSHHHHHHGSGLVPRGSASMSDSEVNQEAKPEVKPEVKPETHINLKVSDGSSEIFFKIKKTTPLRR
LMEAFAKRQGKEMDSLRFLYDGIRIQADQTPEDLDMEDNDIIEAHREQIGGGRVAILWHEMWHEGLEE
ASRLYFGERNVKGMFEVLEPLHAMMERGPQTLKETSFNQAYGRDLMEAQEWCRKYMKSGNVKDLLQ
AWDLYYHVFRRISGNGNGGEQEVFEYCLEDGSLIRATKDHKFMTVDGQMLPIDEIFERELDLMRVDNL
PNGSGGENLYFQGENLYFQGGSGGIKIATRKYLGKQNVYDIGVERDHNFALKNGFIASNCMELRDSSG
KTISTWISDGHVKDFYLYPGKYTFVETAAPDGYEVATPIEFTVNEDGQVTVDGEATEGDAHTGSTSGS
DLGKKLLEAARAGQDDEVRILMANGADVNADDTWGWTPLHLAAYQGHLEIVEVLLKNGADVNAYDYIG

WTPLHLAADGHLEIVEVLLKNGADVNASDYIGDTPLHLAAHNGHLEIVEVLLKHGADVNAQDKFGKTAF
DISIDNGNEDLAEILQKLNEQKLISEEDL

$His_6$-SUMO-FRB-NpuC$^{cage}$-SpyC$^{56-113}$-Myc

MGSSHHHHHHGSGLVPRGSASMSDSEVNQEAKPEVKPEVKPETHINLKVSDGSSEIFFKIKKTTPLRR
LMEAFAKRQGKEMDSLRFLYDGIRIQADQTPEDLDMEDNDIIEAHREQIGGGRVAILWHEMWHEGLEE
ASRLYFGERNVKGMFEVLEPLHAMMERGPQTLKETSFNQAYGRDLMEAQEWCRKYMKSGNVKDLLQ
AWDLYYHVFRRISGNGNGGEQEVFEYCLEDGSLIRATKDHKFMTVDGQMLPIDEIFERELDLMRVDNL
PNGSGGENLYFQGENLYFQGGSGGIKIATRKYLGKQNVYDIGVERDHNFALKNGFIASNCWISDGHVK
DFYLYPGKYTFVETAAPDGYEVATPIEFTVNEDGQVTVDGEATEGDAHTGSTSGSDLGKKLLEAARAG
QDDEVRILMANGADVNADDTWGWTPLHLAAYQGHLEIVEVLLKNGADVNAYDYIGWTPLHLAADGHL
EIVEVLLKNGADVNASDYIGDTPLHLAAHNGHLEIVEVLLKHGADVNAQDKFGKTAFDISIDNGNEDLAEI
LQKLNEQKLISEEDL

$His_6$-SUMO-FRB-NpuC$^{cage}$-SpyC$^{74-113}$-Myc

MGSSHHHHHHGSGLVPRGSASMSDSEVNQEAKPEVKPEVKPETHINLKVSDGSSEIFFKIKKTTPLRR
LMEAFAKRQGKEMDSLRFLYDGIRIQADQTPEDLDMEDNDIIEAHREQIGGGRVAILWHEMWHEGLEE
ASRLYFGERNVKGMFEVLEPLHAMMERGPQTLKETSFNQAYGRDLMEAQEWCRKYMKSGNVKDLLQ
AWDLYYHVFRRISGNGNGGEQEVFEYCLEDGSLIRATKDHKFMTVDGQMLPIDEIFERELDLMRVDNL
PNGSGGENLYFQGENLYFQGGSGGIKIATRKYLGKQNVYDIGVERDHNFALKNGFIASNCFVETAAPD
GYEVATPIEFTVNEDGQVTVDGEATEGDAHTGSTSGSDLGKKLLEAARAGQDDEVRILMANGADVNA
DDTWGWTPLHLAAYQGHLEIVEVLLKNGADVNAYDYIGWTPLHLAADGHLEIVEVLLKNGADVNASDYI
GDTPLHLAAHNGHLEIVEVLLKHGADVNAQDKFGKTAFDISIDNGNEDLAEILQKLNEQKLISEEDL

$His_6$-SUMO-FRB-NpuC$^{cage}$-SpyC$^{83-113}$-Myc

MGSSHHHHHHGSGLVPRGSASMSDSEVNQEAKPEVKPEVKPETHINLKVSDGSSEIFFKIKKTTPLRR
LMEAFAKRQGKEMDSLRFLYDGIRIQADQTPEDLDMEDNDIIEAHREQIGGGRVAILWHEMWHEGLEE
ASRLYFGERNVKGMFEVLEPLHAMMERGPQTLKETSFNQAYGRDLMEAQEWCRKYMKSGNVKDLLQ
AWDLYYHVFRRISGNGNGGEQEVFEYCLEDGSLIRATKDHKFMTVDGQMLPIDEIFERELDLMRVDNL
PNGSGGENLYFQGENLYFQGGSGGIKIATRKYLGKQNVYDIGVERDHNFALKNGFIASNCYEVATPIEF
TVNEDGQVTVDGEATEGDAHTGSTSGSDLGKKLLEAARAGQDDEVRILMANGADVNADDTWGWTPL
HLAAYQGHLEIVEVLLKNGADVNAYDYIGWTPLHLAADGHLEIVEVLLKNGADVNASDYIGDTPLHLAAH
NGHLEIVEVLLKHGADVNAQDKFGKTAFDISIDNGNEDLAEILQKLNEQKLISEEDL

$His_6$-SUMO-FRB-NpuC$^{cage}$-SpyC$^{91-113}$-Myc

MGSSHHHHHHGSGLVPRGSASMSDSEVNQEAKPEVKPEVKPETHINLKVSDGSSEIFFKIKKTTPLRR

LMEAFAKRQGKEMDSLRFLYDGIRIQADQTPEDLDMEDNDIIEAHREQIGGGRVAILWHEMWHEGLEE
ASRLYFGERNVKGMFEVLEPLHAMMERGPQTLKETSFNQAYGRDLMEAQEWCRKYMKSGNVKDLLQ
AWDLYYHVFRRISGNGNGGEQEVFEYCLEDGSLIRATKDHKFMTVDGQMLPIDEIFERELDLMRVDNL
PNGSGGENLYFQGENLYFQGGSGGIKIATRKYLGKQNVYDIGVERDHNFALKNGFIASNCFTVNEDGQ
VTVDGEATEGDAHTGSTSGSDLGKKLLEAARAGQDDEVRILMANGADVNADDTWGWTPLHLAAYQG
HLEIVEVLLKNGADVNAYDYIGWTPLHLAADGHLEIVEVLLKNGADVNASDYIGDTPLHLAAHNGHLEIV
EVLLKHGADVNAQDKFGKTAFDISIDNGNEDLAEILQKLNEQKLISEEDL

His$_6$-SUMO-FLAG-SpyN$^{1-73}$-NrdJ-1N$^{cage}$-FKBP

MGSSHHHHHHGSGLVPRGSASMSDSEVNQEAKPEVKPEVKPETHINLKVSDGSSEIFFKIKKTTPLRR
LMEAFAKRQGKEMDSLRFLYDGIRIQADQTPEDLDMEDNDIIEAHREQIGGDYKDDDDKDLGKKLLEAA
RAGQDDEVRILMANGADVNAKDEYGLTPLYLATAHGHLEIVEVLLKNGADVNAVDAIGFTPLHLAAFIGH
LEIAEVLLKHGADVNAQDKFGKTAFDISIGNGNEDLAEILQKLNGSTSGSVTTLSGLSGEQGPSGDMTT
EEDSATHIKFSKRDEDGRELAGATMELRDSSGKTISTWISDGHVKDFYLYPGKYCLVGSSEIITRNYGK
TTIKEVVEIFDNDKNIQVLAFNTHTDNIEWAPIKAAQLTRPNAELVELEIDTLHGVKTIRCTPDHPVYTKNR
GYVRADELTDDDELVVAIENLYGDEVDGYFQGMEAKTYIGKLKSRKIVSNEDTYDIQTSTHNFFANDNS
AGGVQVETISPGDGRTFPKRGQTCVVHYTGMLEDGKKFDSSRDRNKPFKFMLGKQEVIRGWEEGVA
QMSVGQRAKLTISPDYAYGATGHPGIIPPHATLVFDVELLKLE

His$_6$-SUMO-FLAG-SpyN$^{1-73}$-NrdJ-1N$^{cage(C1A)}$-FKBP

MGSSHHHHHHGSGLVPRGSASMSDSEVNQEAKPEVKPEVKPETHINLKVSDGSSEIFFKIKKTTPLRR
LMEAFAKRQGKEMDSLRFLYDGIRIQADQTPEDLDMEDNDIIEAHREQIGGDYKDDDDKDLGKKLLEAA
RAGQDDEVRILMANGADVNAKDEYGLTPLYLATAHGHLEIVEVLLKNGADVNAVDAIGFTPLHLAAFIGH
LEIAEVLLKHGADVNAQDKFGKTAFDISIGNGNEDLAEILQKLNGSTSGSVTTLSGLSGEQGPSGDMTT
EEDSATHIKFSKRDEDGRELAGATMELRDSSGKTISTWISDGHVKDFYLYPGKYALVGSSEIITRNYGKT
TIKEVVEIFDNDKNIQVLAFNTHTDNIEWAPIKAAQLTRPNAELVELEIDTLHGVKTIRVTPDHPVYTKNR
GYVRADELTDDDELVVAIENLYGDEVDGYFQGMEAKTYIGKLKSRKIVSNEDTYDIQTSTHNFFANDNS
AGGVQVETISPGDGRTFPKRGQTCVVHYTGMLEDGKKFDSSRDRNKPFKFMLGKQEVIRGWEEGVA
QMSVGQRAKLTISPDYAYGATGHPGIIPPHATLVFDVELLKLE

His$_6$-SUMO-FLAG-SpyN$^{1-73}$-NrdJ-1N$^{cage(C76V)}$-FKBP

MGSSHHHHHHGSGLVPRGSASMSDSEVNQEAKPEVKPEVKPETHINLKVSDGSSEIFFKIKKTTPLRR
LMEAFAKRQGKEMDSLRFLYDGIRIQADQTPEDLDMEDNDIIEAHREQIGGDYKDDDDKDLGKKLLEAA
RAGQDDEVRILMANGADVNAKDEYGLTPLYLATAHGHLEIVEVLLKNGADVNAVDAIGFTPLHLAAFIGH
LEIAEVLLKHGADVNAQDKFGKTAFDISIGNGNEDLAEILQKLNGSTSGSVTTLSGLSGEQGPSGDMTT
EEDSATHIKFSKRDEDGRELAGATMELRDSSGKTISTWISDGHVKDFYLYPGKYCLVGSSEIITRNYGK

TTIKEVVEIFDNDKNIQVLAFNTHTDNIEWAPIKAAQLTRPNAELVELEIDTLHGVKTIRVTPDHPVYTKNR
GYVRADELTDDDELVVAIENLYGDEVDGYFQGMEAKTYIGKLKSRKIVSNEDTYDIQTSTHNFFANDNS
AGGVQVETISPGDGRTFPKRGQTCVVHYTGMLEDGKKFDSSRDRNKPFKFMLGKQEVIRGWEEGVA
QMSVGQRAKLTISPDYAYGATGHPGIIPPHATLVFDVELLKLE

$His_6$-SUMO-FRB-NrdJ-1$C^{cage}$-Spy$C^{74\text{-}113}$-Myc

MGSSHHHHHHGSGLVPRGSASMSDSEVNQEAKPEVKPEVKPETHINLKVSDGSSEIFFKIKKTTPLRR
LMEAFAKRQGKEMDSLRFLYDGIRIQADQTPEDLDMEDNDIIEAHREQIGGGRVAILWHEMWHEGLEE
ASRLYFGERNVKGMFEVLEPLHAMMERGPQTLKETSFNQAYGRDLMEAQEWCRKYMKSGNVKDLLQ
AWDLYYHVFRRISGNGNGNAELVELEIDTLHGVKTIRCTPDHPVYTKNRGYVRADELTDDDELVVAIEN
LYGDEVDGYFQGMEAKTYIGKLKSRKIVSNEDTYDIQTSTHNFFANDILVHNSFVETAAPDGYEVATPIE
FTVNEDGQVTVDGEATEGDAHTGSTSGSDLGKKLLEAARAGQDDEVRILMANGADVNADDTWGWTP
LHLAAYQGHLEIVEVLLKNGADVNAYDYIGWTPLHLAADGHLEIVEVLLKNGADVNASDYIGDTPLHLAA
HNGHLEIVEVLLKHGADVNAQDKFGKTAFDISIDNGNEDLAEILQKLNEQKLISEEDL

$His_6$-SUMO-FRB-NrdJ-1$C^{cage(C76V)}$-Spy$C^{74\text{-}113}$-Myc

MGSSHHHHHHGSGLVPRGSASMSDSEVNQEAKPEVKPEVKPETHINLKVSDGSSEIFFKIKKTTPLRR
LMEAFAKRQGKEMDSLRFLYDGIRIQADQTPEDLDMEDNDIIEAHREQIGGGRVAILWHEMWHEGLEE
ASRLYFGERNVKGMFEVLEPLHAMMERGPQTLKETSFNQAYGRDLMEAQEWCRKYMKSGNVKDLLQ
AWDLYYHVFRRISGNGNGNAELVELEIDTLHGVKTIRVTPDHPVYTKNRGYVRADELTDDDELVVAIEN
LYGDEVDGYFQGMEAKTYIGKLKSRKIVSNEDTYDIQTSTHNFFANDILVHNSFVETAAPDGYEVATPIE
FTVNEDGQVTVDGEATEGDAHTGSTSGSDLGKKLLEAARAGQDDEVRILMANGADVNADDTWGWTP
LHLAAYQGHLEIVEVLLKNGADVNAYDYIGWTPLHLAADGHLEIVEVLLKNGADVNASDYIGDTPLHLAA
HNGHLEIVEVLLKHGADVNAQDKFGKTAFDISIDNGNEDLAEILQKLNEQKLISEEDL

$His_6$-SUMO-αHER2-SpyN-eNrdJ-1$N^{cage(C76V)}$

MGSSHHHHHHGSGLVPRGSASMSDSEVNQEAKPEVKPEVKPETHINLKVSDGSSEIFFKIKKTTPLRR
LMEAFAKRQGKEMDSLRFLYDGIRIQADQTPEDLDMEDNDIIEAHREQIGGDYKDDDDKDLGKKLLEAA
RAGQDDEVRILMANGADVNAKDEYGLTPLYLATAHGHLEIVEVLLKNGADVNAVDAIGFTPLHLAAFIGH
LEIAEVLLKHGADVNAQDKFGKTAFDISIGNGNEDLAEILQKLNGSTSGSVTTLSGLSGEQGPSGDMTT
EEDSATHIKFSKRDEDGRELAGATMELRDSSGKTISTWISDGHVKDFYLYPGKYCLVGSSEIITRNYGK
TTIKEVVEIFDNDKNIQVLAFNTHTDNIEWAPIKAAQLTRPNAELVELEIDTLHGVKTIRVTPDHPVYTKNR
GYVRADELTDDDELVVAIENLYGDEVDGYFQGMEAKTYIGKLKSRKIVSNEDTYDIQTSTHNFFAND

$His_6$-SUMO-αHER2-SpyN-eNrdJ-1$N^{cage}$

MGSSHHHHHHGSGLVPRGSASMSDSEVNQEAKPEVKPEVKPETHINLKVSDGSSEIFFKIKKTTPLRR

LMEAFAKRQGKEMDSLRFLYDGIRIQADQTPEDLDMEDNDIIEAHREQIGGDYKDDDDKDLGKKLLEAA
RAGQDDEVRILMANGADVNAKDEYGLTPLYLATAHGHLEIVEVLLKNGADVNAVDAIGFTPLHLAAFIGH
LEIAEVLLKHGADVNAQDKFGKTAFDISIGNGNEDLAEILQKLNGSTSGSVTTLSGLSGEQGPSGDMTT
EEDSATHIKFSKRDEDGRELAGATMELRDSSGKTISTWISDGHVKDFYLYPGKYCLVGSSEIITRNYGK
TTIKEVVEIFDNDKNIQVLAFNTHTDNIEWAPIKAAQLTRPNAELVELEIDTLHGVKTIRVTPDHPVYTKNR
GYVRADELTDDDELVVAIENLYGDEVNGYFQGMEAETYIGKLKSRAIVSNEDTYDIQTSTHNFFAND

$His_6$-SUMO-αEGFR-SpyN-eNrdJ-1N$^{cage}$

MGSSHHHHHHGSGLVPRGSASMSDSEVNQEAKPEVKPEVKPETHINLKVSDGSSEIFFKIKKTTPLRR
LMEAFAKRQGKEMDSLRFLYDGIRIQADQTPEDLDMEDNDIIEAHREQIGGDYKDDDDKDLGKKLLEAA
RAGQDDEVRILMANGADVNADDTWGWTPLHLAAYQGHLEIVEVLLKNGADVNAYDYIGWTPLHLAAD
GHLEIVEVLLKNGADVNASDYIGDTPLHLAAHNGHLEIVEVLLKHGADVNAQDKFGKTAFDISIDNGNED
LAEILQKLNGSTSGSVTTLSGLSGEQGPSGDMTTEEDSATHIKFSKRDEDGRELAGATMELRDSSGKTI
STWISDGHVKDFYLYPGKYCLVGSSEIITRNYGKTTIKEVVEIFDNDKNIQVLAFNTHTDNIEWAPIKAAQ
LTRPNAELVELEIDTLHGVKTIRVTPDHPVYTKNRGYVRADELTDDDELVVAIENLYGDEVNGYFQGME
AETYIGKLKSRAIVSNEDTYDIQTSTHNFFAND

$His_6$-SUMO-αEpCAM-SpyN-eNrdJ-1N$^{cage}$

MGSSHHHHHHGSGLVPRGSASMSDSEVNQEAKPEVKPEVKPETHINLKVSDGSSEIFFKIKKTTPLRR
LMEAFAKRQGKEMDSLRFLYDGIRIQADQTPEDLDMEDNDIIEAHREQIGGDYKDDDDKDLGKKLLEAA
RAGQDDEVRILVANGADVNAYFGTTPLHLAAAHGRLEIVEVLLKNGADVNAQDVWGITPLHLAAYNGH
LEIVEVLLKYGADVNAHDTRGWTPLHLAAINGHLEIVEVLLKNVADVNAQDRSGKTPFDLAIDNGNEDIA
EVLQKAAKLNGSTSGSVTTLSGLSGEQGPSGDMTTEEDSATHIKFSKRDEDGRELAGATMELRDSSG
KTISTWISDGHVKDFYLYPGKYCLVGSSEIITRNYGKTTIKEVVEIFDNDKNIQVLAFNTHTDNIEWAPIKA
AQLTRPNAELVELEIDTLHGVKTIRVTPDHPVYTKNRGYVRADELTDDDELVVAIENLYGDEVNGYFQG
MEAETYIGKLKSRAIVSNEDTYDIQTSTHNFFAND

$His_6$-SUMO-NrdJ-1C$^{cage(C76V)}$-SpyC-αEGFR

MGSSHHHHHHGSGLVPRGSASMSDSEVNQEAKPEVKPEVKPETHINLKVSDGSSEIFFKIKKTTPLRR
LMEAFAKRQGKEMDSLRFLYDGIRIQADQTPEDLDMEDNDIIEAHREQIGGNAELVELEIDTLHGVKTIR
VTPDHPVYTKNRGYVRADELTDDDELVVAIENLYGDEVDGYFQGMEAKTYIGKLKSRKIVSNEDTYDIQ
TSTHNFFANDILVHNSFVETAAPDGYEVATPIEFTVNEDGQVTVDGEATEGDAHTGSTSGSDLGKKLLE
AARAGQDDEVRILMANGADVNADDTWGWTPLHLAAYQGHLEIVEVLLKNGADVNAYDYIGWTPLHLA
ADGHLEIVEVLLKNGADVNASDYIGDTPLHLAAHNGHLEIVEVLLKHGADVNAQDKFGKTAFDISIDNGN
EDLAEILQKLNEQKLISEEDL

His$_6$-SUMO-eNrdJ-1C$^{cage}$-SpyC-αHER2

MGSSHHHHHHGSGLVPRGSASMSDSEVNQEAKPEVKPEVKPETHINLKVSDGSSEIFFKIKKTTPLRR
LMEAFAKRQGKEMDSLRFLYDGIRIQADQTPEDLDMEDNDIIEAHREQIGGNAELVELEIKTLHGVKTIR
VTPDHPVYTKNRGYVRADELTDDDELVVAIENLYGDEVNGYFQGMEAKTYIGKLKSRKIVSNEDTYDIQ
TSTHNFFANDILVHNSFVETAAPDGYEVATPIEFTVNEDGQVTVDGEATEGDAHTGSTSGSDLGKKLLE
AARAGQDDEVRILMANGADVNAKDEYGLTPLYLATAHGHLEIVEVLLKNGADVNAVDAIGFTPLHLAAFI
GHLEIAEVLLKHGADVNAQDKFGKTAFDISIGNGNEDLAEILQKLNEQKLISEEDL

His$_6$-SUMO-eNrdJ-1C$^{cage}$-SpyC-αEGFR

MGSSHHHHHHGSGLVPRGSASMSDSEVNQEAKPEVKPEVKPETHINLKVSDGSSEIFFKIKKTTPLRR
LMEAFAKRQGKEMDSLRFLYDGIRIQADQTPEDLDMEDNDIIEAHREQIGGNAELVELEIKTLHGVKTIR
VTPDHPVYTKNRGYVRADELTDDDELVVAIENLYGDEVNGYFQGMEAKTYIGKLKSRKIVSNEDTYDIQ
TSTHNFFANDILVHNSFVETAAPDGYEVATPIEFTVNEDGQVTVDGEATEGDAHTGSTSGSDLGKKLLE
AARAGQDDEVRILMANGADVNADDTWGWTPLHLAAYQGHLEIVEVLLKNGADVNAYDYIGWTPLHLA
ADGHLEIVEVLLKNGADVNASDYIGDTPLHLAAHNGHLEIVEVLLKHGADVNAQDKFGKTAFDISIDNGN
EDLAEILQKLNEQKLISEEDL

His$_6$-SUMO-eNrdJ-1C$^{cage}$-SpyC-αEpCAM

MGSSHHHHHHGSGLVPRGSASMSDSEVNQEAKPEVKPEVKPETHINLKVSDGSSEIFFKIKKTTPLRR
LMEAFAKRQGKEMDSLRFLYDGIRIQADQTPEDLDMEDNDIIEAHREQIGGNAELVELEIKTLHGVKTIR
VTPDHPVYTKNRGYVRADELTDDDELVVAIENLYGDEVNGYFQGMEAKTYIGKLKSRKIVSNEDTYDIQ
TSTHNFFANDILVHNSFVETAAPDGYEVATPIEFTVNEDGQVTVDGEATEGDAHTGSTSGSDLGKKLLE
AARAGQDDEVRILVANGADVNAYFGTTPLHLAAAHGRLEIVEVLLKNGADVNAQDVWGITPLHLAAYN
GHLEIVEVLLKYGADVNAHDTRGWTPLHLAAINGHLEIVEVLLKNVADVNAQDRSGKTPFDLAIDNGNE
DIAEVLQKAAKLNEQKLISEEDL

His$_6$-SUMO-αHER2-SpyN-eNrdJ-1N$^{cage(K114A)}$

MGSSHHHHHHGSGLVPRGSASMSDSEVNQEAKPEVKPEVKPETHINLKVSDGSSEIFFKIKKTTPLRR
LMEAFAKRQGKEMDSLRFLYDGIRIQADQTPEDLDMEDNDIIEAHREQIGGDYKDDDDKDLGKKLLEAA
RAGQDDEVRILMANGADVNAKDEYGLTPLYLATAHGHLEIVEVLLKNGADVNAVDAIGFTPLHLAAFIGH
LEIAEVLLKHGADVNAQDKFGKTAFDISIGNGNEDLAEILQKLNGSTSGSVTTLSGLSGEQGPSGDMTT
EEDSATHIKFSKRDEDGRELAGATMELRDSSGKTISTWISDGHVKDFYLYPGKYCLVGSSEIITRNYGK
TTIKEVVEIFDNDKNIQVLAFNTHTDNIEWAPIKAAQLTRPNAELVELEIDTLHGVKTIRVTPDHPVYTKNR
GYVRADELTDDDELVVAIENLYGDEVNGYFQGMEAETYIGALKSRAIVSNEDTYDIQTSTHNFFAND

His$_6$-SUMO-αHER2-SpyN-eNrdJ-1N$^{cage(K116A)}$

MGSSHHHHHHGSGLVPRGSASMSDSEVNQEAKPEVKPEVKPETHINLKVSDGSSEIFFKIKKTTPLRR
LMEAFAKRQGKEMDSLRFLYDGIRIQADQTPEDLDMEDNDIIEAHREQIGGDYKDDDDKDLGKKLLEAA
RAGQDDEVRILMANGADVNAKDEYGLTPLYLATAHGHLEIVEVLLKNGADVNAVDAIGFTPLHLAAFIGH
LEIAEVLLKHGADVNAQDKFGKTAFDISIGNGNEDLAEILQKLNGSTSGSVTTLSGLSGEQGPSGDMTT
EEDSATHIKFSKRDEDGRELAGATMELRDSSGKTISTWISDGHVKDFYLYPGKYCLVGSSEIITRNYGK
TTIKEVVEIFDNDKNIQVLAFNTHTDNIEWAPIKAAQLTRPNAELVELEIDTLHGVKTIRVTPDHPVYTKNR
GYVRADELTDDDELVVAIENLYGDEVNGYFQGMEAETYIGKLASRAIVSNEDTYDIQTSTHNFFAND

His$_6$-SUMO-αHER2-SpyN-eNrdJ-1N$^{cage(R118A)}$

MGSSHHHHHHGSGLVPRGSASMSDSEVNQEAKPEVKPEVKPETHINLKVSDGSSEIFFKIKKTTPLRR
LMEAFAKRQGKEMDSLRFLYDGIRIQADQTPEDLDMEDNDIIEAHREQIGGDYKDDDDKDLGKKLLEAA
RAGQDDEVRILMANGADVNAKDEYGLTPLYLATAHGHLEIVEVLLKNGADVNAVDAIGFTPLHLAAFIGH
LEIAEVLLKHGADVNAQDKFGKTAFDISIGNGNEDLAEILQKLNGSTSGSVTTLSGLSGEQGPSGDMTT
EEDSATHIKFSKRDEDGRELAGATMELRDSSGKTISTWISDGHVKDFYLYPGKYCLVGSSEIITRNYGK
TTIKEVVEIFDNDKNIQVLAFNTHTDNIEWAPIKAAQLTRPNAELVELEIDTLHGVKTIRVTPDHPVYTKNR
GYVRADELTDDDELVVAIENLYGDEVNGYFQGMEAETYIGKLKSAAIVSNEDTYDIQTSTHNFFAND

His$_6$-SUMO-αHER2-SpyN-eNrdJ-1N$^{cage(K114AK116A)}$

MGSSHHHHHHGSGLVPRGSASMSDSEVNQEAKPEVKPEVKPETHINLKVSDGSSEIFFKIKKTTPLRR
LMEAFAKRQGKEMDSLRFLYDGIRIQADQTPEDLDMEDNDIIEAHREQIGGDYKDDDDKDLGKKLLEAA
RAGQDDEVRILMANGADVNAKDEYGLTPLYLATAHGHLEIVEVLLKNGADVNAVDAIGFTPLHLAAFIGH
LEIAEVLLKHGADVNAQDKFGKTAFDISIGNGNEDLAEILQKLNGSTSGSVTTLSGLSGEQGPSGDMTT
EEDSATHIKFSKRDEDGRELAGATMELRDSSGKTISTWISDGHVKDFYLYPGKYCLVGSSEIITRNYGK
TTIKEVVEIFDNDKNIQVLAFNTHTDNIEWAPIKAAQLTRPNAELVELEIDTLHGVKTIRVTPDHPVYTKNR
GYVRADELTDDDELVVAIENLYGDEVNGYFQGMEAETYIGALASRAIVSNEDTYDIQTSTHNFFAND

His$_6$-SUMO-αHER2-SpyN-eNrdJ-1N$^{cage(K114AK116AR118A)}$

MGSSHHHHHHGSGLVPRGSASMSDSEVNQEAKPEVKPEVKPETHINLKVSDGSSEIFFKIKKTTPLRR
LMEAFAKRQGKEMDSLRFLYDGIRIQADQTPEDLDMEDNDIIEAHREQIGGDYKDDDDKDLGKKLLEAA
RAGQDDEVRILMANGADVNAKDEYGLTPLYLATAHGHLEIVEVLLKNGADVNAVDAIGFTPLHLAAFIGH
LEIAEVLLKHGADVNAQDKFGKTAFDISIGNGNEDLAEILQKLNGSTSGSVTTLSGLSGEQGPSGDMTT
EEDSATHIKFSKRDEDGRELAGATMELRDSSGKTISTWISDGHVKDFYLYPGKYCLVGSSEIITRNYGK
TTIKEVVEIFDNDKNIQVLAFNTHTDNIEWAPIKAAQLTRPNAELVELEIDTLHGVKTIRVTPDHPVYTKNR
GYVRADELTDDDELVVAIENLYGDEVNGYFQGMEAETYIGALASAAIVSNEDTYDIQTSTHNFFAND

His$_6$-SUMO-αHER2-SpyN-eNrdJ-1N$^{cage(A119K)}$

MGSSHHHHHHGSGLVPRGSASMSDSEVNQEAKPEVKPEVKPETHINLKVSDGSSEIFFKIKKTTPLRR
LMEAFAKRQGKEMDSLRFLYDGIRIQADQTPEDLDMEDNDIIEAHREQIGGDYKDDDDKDLGKKLLEAA
RAGQDDEVRILMANGADVNAKDEYGLTPLYLATAHGHLEIVEVLLKNGADVNAVDAIGFTPLHLAAFIGH
LEIAEVLLKHGADVNAQDKFGKTAFDISIGNGNEDLAEILQKLNGSTSGSVTTLSGLSGEQGPSGDMTT
EEDSATHIKFSKRDEDGRELAGATMELRDSSGKTISTWISDGHVKDFYLYPGKYCLVGSSEIITRNYGK
TTIKEVVEIFDNDKNIQVLAFNTHTDNIEWAPIKAAQLTRPNAELVELEIDTLHGVKTIRVTPDHPVYTKNR
GYVRADELTDDDELVVAIENLYGDEVNGYFQGMEAETYIGKLKSRKIVSNEDTYDIQTSTHNFFAND

His$_6$-SUMO-αEGFR-SpyN-eNrdJ-1N$^{cage(K114AK116A)}$

MGSSHHHHHHGSGLVPRGSASMSDSEVNQEAKPEVKPEVKPETHINLKVSDGSSEIFFKIKKTTPLRR
LMEAFAKRQGKEMDSLRFLYDGIRIQADQTPEDLDMEDNDIIEAHREQIGGDYKDDDDKDLGKKLLEAA
RAGQDDEVRILMANGADVNADDTWGWTPLHLAAYQGHLEIVEVLLKNGADVNAYDYIGWTPLHLAAD
GHLEIVEVLLKNGADVNASDYIGDTPLHLAAHNGHLEIVEVLLKHGADVNAQDKFGKTAFDISIDNGNED
LAEILQKLNGSTSGSVTTLSGLSGEQGPSGDMTTEEDSATHIKFSKRDEDGRELAGATMELRDSSGKTI
STWISDGHVKDFYLYPGKYCLVGSSEIITRNYGKTTIKEVVEIFDNDKNIQVLAFNTHTDNIEWAPIKAAQ
LTRPNAELVELEIDTLHGVKTIRVTPDHPVYTKNRGYVRADELTDDDELVVAIENLYGDEVNGYFQGME
AETYIGALASRAIVSNEDTYDIQTSTHNFFAND

His$_6$-SUMO-αEpCAM-SpyN-eNrdJ-1N$^{cage(K114AK116A)}$

MGSSHHHHHHGSGLVPRGSASMSDSEVNQEAKPEVKPEVKPETHINLKVSDGSSEIFFKIKKTTPLRR
LMEAFAKRQGKEMDSLRFLYDGIRIQADQTPEDLDMEDNDIIEAHREQIGGDYKDDDDKDLGKKLLEAA
RAGQDDEVRILVANGADVNAYFGTTPLHLAAAHGRLEIVEVLLKNGADVNAQDVWGITPLHLAAYNGH
LEIVEVLLKYGADVNAHDTRGWTPLHLAAINGHLEIVEVLLKNVADVNAQDRSGKTPFDLAIDNGNEDIA
EVLQKAAKLNGSTSGSVTTLSGLSGEQGPSGDMTTEEDSATHIKFSKRDEDGRELAGATMELRDSSG
KTISTWISDGHVKDFYLYPGKYCLVGSSEIITRNYGKTTIKEVVEIFDNDKNIQVLAFNTHTDNIEWAPIKA
AQLTRPNAELVELEIDTLHGVKTIRVTPDHPVYTKNRGYVRADELTDDDELVVAIENLYGDEVNGYFQG
MEAETYIGALASRAIVSNEDTYDIQTSTHNFFAND

His$_6$-SUMO-αHER2-SpyN-eNrdJ-1N$^{cage(1-39)}$

MGSSHHHHHHGSGLVPRGSASMSDSEVNQEAKPEVKPEVKPETHINLKVSDGSSEIFFKIKKTTPLRR
LMEAFAKRQGKEMDSLRFLYDGIRIQADQTPEDLDMEDNDIIEAHREQIGGDYKDDDDKDLGKKLLEAA
RAGQDDEVRILMANGADVNAKDEYGLTPLYLATAHGHLEIVEVLLKNGADVNAVDAIGFTPLHLAAFIGH
LEIAEVLLKHGADVNAQDKFGKTAFDISIGNGNEDLAEILQKLNGSTSGSVTTLSGLSGEQGPSGDMTT
EEDSATHIKFSKRDEDGRELAGATMELRDSSGKTISTWISDGHVKDFYLYPGKYCLVGSSEIITRNYGK
TTIKEVVEIFDNDKNIQVLAFNTHTDNIEWAPIKAAQLTRPNAELVELEIDTLHGVKTIRVTPDHPVYTKNR

GYVRADELTDDDELVVAIENLYGDEVNGYFQGMEAETYIGKLKSRAIVSNEDTYDIQTSTHNFFANDILV
H

His$_6$-SUMO-αHER2-SpyN-eNrdJ-1N$^{cage(1-38)}$

MGSSHHHHHHGSGLVPRGSASMSDSEVNQEAKPEVKPEVKPETHINLKVSDGSSEIFFKIKKTTPLRR
LMEAFAKRQGKEMDSLRFLYDGIRIQADQTPEDLDMEDNDIIEAHREQIGGDYKDDDDKDLGKKLLEAA
RAGQDDEVRILMANGADVNAKDEYGLTPLYLATAHGHLEIVEVLLKNGADVNAVDAIGFTPLHLAAFIGH
LEIAEVLLKHGADVNAQDKFGKTAFDISIGNGNEDLAEILQKLNGSTSGSVTTLSGLSGEQGPSGDMTT
EEDSATHIKFSKRDEDGRELAGATMELRDSSGKTISTWISDGHVKDFYLYPGKYCLVGSSEIITRNYGK
TTIKEVVEIFDNDKNIQVLAFNTHTDNIEWAPIKAAQLTRPNAELVELEIDTLHGVKTIRVTPDHPVYTKNR
GYVRADELTDDDELVVAIENLYGDEVNGYFQGMEAETYIGKLKSRAIVSNEDTYDIQTSTHNFFANDILV

His$_6$-SUMO-αHER2-SpyN-eNrdJ-1N$^{cage(1-37)}$

MGSSHHHHHHGSGLVPRGSASMSDSEVNQEAKPEVKPEVKPETHINLKVSDGSSEIFFKIKKTTPLRR
LMEAFAKRQGKEMDSLRFLYDGIRIQADQTPEDLDMEDNDIIEAHREQIGGDYKDDDDKDLGKKLLEAA
RAGQDDEVRILMANGADVNAKDEYGLTPLYLATAHGHLEIVEVLLKNGADVNAVDAIGFTPLHLAAFIGH
LEIAEVLLKHGADVNAQDKFGKTAFDISIGNGNEDLAEILQKLNGSTSGSVTTLSGLSGEQGPSGDMTT
EEDSATHIKFSKRDEDGRELAGATMELRDSSGKTISTWISDGHVKDFYLYPGKYCLVGSSEIITRNYGK
TTIKEVVEIFDNDKNIQVLAFNTHTDNIEWAPIKAAQLTRPNAELVELEIDTLHGVKTIRVTPDHPVYTKNR
GYVRADELTDDDELVVAIENLYGDEVNGYFQGMEAETYIGKLKSRAIVSNEDTYDIQTSTHNFFANDIL

His$_6$-SUMO-αHER2-SpyN-eNrdJ-1N$^{cage(1-36)}$

MGSSHHHHHHGSGLVPRGSASMSDSEVNQEAKPEVKPEVKPETHINLKVSDGSSEIFFKIKKTTPLRR
LMEAFAKRQGKEMDSLRFLYDGIRIQADQTPEDLDMEDNDIIEAHREQIGGDYKDDDDKDLGKKLLEAA
RAGQDDEVRILMANGADVNAKDEYGLTPLYLATAHGHLEIVEVLLKNGADVNAVDAIGFTPLHLAAFIGH
LEIAEVLLKHGADVNAQDKFGKTAFDISIGNGNEDLAEILQKLNGSTSGSVTTLSGLSGEQGPSGDMTT
EEDSATHIKFSKRDEDGRELAGATMELRDSSGKTISTWISDGHVKDFYLYPGKYCLVGSSEIITRNYGK
TTIKEVVEIFDNDKNIQVLAFNTHTDNIEWAPIKAAQLTRPNAELVELEIDTLHGVKTIRVTPDHPVYTKNR
GYVRADELTDDDELVVAIENLYGDEVNGYFQGMEAETYIGKLKSRAIVSNEDTYDIQTSTHNFFANDI

His$_6$-SUMO-αHER2-SpyN-eNrdJ-1N$^{cage(1-33)}$

MGSSHHHHHHGSGLVPRGSASMSDSEVNQEAKPEVKPEVKPETHINLKVSDGSSEIFFKIKKTTPLRR
LMEAFAKRQGKEMDSLRFLYDGIRIQADQTPEDLDMEDNDIIEAHREQIGGDYKDDDDKDLGKKLLEAA
RAGQDDEVRILMANGADVNAKDEYGLTPLYLATAHGHLEIVEVLLKNGADVNAVDAIGFTPLHLAAFIGH
LEIAEVLLKHGADVNAQDKFGKTAFDISIGNGNEDLAEILQKLNGSTSGSVTTLSGLSGEQGPSGDMTT
EEDSATHIKFSKRDEDGRELAGATMELRDSSGKTISTWISDGHVKDFYLYPGKYCLVGSSEIITRNYGK

TTIKEVVEIFDNDKNIQVLAFNTHTDNIEWAPIKAAQLTRPNAELVELEIDTLHGVKTIRVTPDHPVYTKNR
GYVRADELTDDDELVVAIENLYGDEVNGYFQGMEAETYIGKLKSRAIVSNEDTYDIQTSTHNFFA

$His_6$-SUMO-αHER2-SpyN-eNrdJ-1N$^{cage(1-32)}$

MGSSHHHHHHGSGLVPRGSASMSDSEVNQEAKPEVKPEVKPETHINLKVSDGSSEIFFKIKKTTPLRR
LMEAFAKRQGKEMDSLRFLYDGIRIQADQTPEDLDMEDNDIIEAHREQIGGDYKDDDDKDLGKKLLEAA
RAGQDDEVRILMANGADVNAKDEYGLTPLYLATAHGHLEIVEVLLKNGADVNAVDAIGFTPLHLAAFIGH
LEIAEVLLKHGADVNAQDKFGKTAFDISIGNGNEDLAEILQKLNGSTSGSVTTLSGLSGEQGPSGDMTT
EEDSATHIKFSKRDEDGRELAGATMELRDSSGKTISTWISDGHVKDFYLYPGKYCLVGSSEIITRNYGK
TTIKEVVEIFDNDKNIQVLAFNTHTDNIEWAPIKAAQLTRPNAELVELEIDTLHGVKTIRVTPDHPVYTKNR
GYVRADELTDDDELVVAIENLYGDEVNGYFQGMEAETYIGKLKSRAIVSNEDTYDIQTSTHNFF

$His_6$-SUMO-αHER2-SpyN-eNrdJ-1N$^{cage(1-31)}$

MGSSHHHHHHGSGLVPRGSASMSDSEVNQEAKPEVKPEVKPETHINLKVSDGSSEIFFKIKKTTPLRR
LMEAFAKRQGKEMDSLRFLYDGIRIQADQTPEDLDMEDNDIIEAHREQIGGDYKDDDDKDLGKKLLEAA
RAGQDDEVRILMANGADVNAKDEYGLTPLYLATAHGHLEIVEVLLKNGADVNAVDAIGFTPLHLAAFIGH
LEIAEVLLKHGADVNAQDKFGKTAFDISIGNGNEDLAEILQKLNGSTSGSVTTLSGLSGEQGPSGDMTT
EEDSATHIKFSKRDEDGRELAGATMELRDSSGKTISTWISDGHVKDFYLYPGKYCLVGSSEIITRNYGK
TTIKEVVEIFDNDKNIQVLAFNTHTDNIEWAPIKAAQLTRPNAELVELEIDTLHGVKTIRVTPDHPVYTKNR
GYVRADELTDDDELVVAIENLYGDEVNGYFQGMEAETYIGKLKSRAIVSNEDTYDIQTSTHNF

$His_6$-SUMO-αHER2-SpyN-eNrdJ-1N$^{cage(1-30)}$

MGSSHHHHHHGSGLVPRGSASMSDSEVNQEAKPEVKPEVKPETHINLKVSDGSSEIFFKIKKTTPLRR
LMEAFAKRQGKEMDSLRFLYDGIRIQADQTPEDLDMEDNDIIEAHREQIGGDYKDDDDKDLGKKLLEAA
RAGQDDEVRILMANGADVNAKDEYGLTPLYLATAHGHLEIVEVLLKNGADVNAVDAIGFTPLHLAAFIGH
LEIAEVLLKHGADVNAQDKFGKTAFDISIGNGNEDLAEILQKLNGSTSGSVTTLSGLSGEQGPSGDMTT
EEDSATHIKFSKRDEDGRELAGATMELRDSSGKTISTWISDGHVKDFYLYPGKYCLVGSSEIITRNYGK
TTIKEVVEIFDNDKNIQVLAFNTHTDNIEWAPIKAAQLTRPNAELVELEIDTLHGVKTIRVTPDHPVYTKNR
GYVRADELTDDDELVVAIENLYGDEVNGYFQGMEAETYIGKLKSRAIVSNEDTYDIQTSTHN

$His_6$-SUMO-αHER2-SpyN-eNrdJ-1N$^{cage(1-29)}$

MGSSHHHHHHGSGLVPRGSASMSDSEVNQEAKPEVKPEVKPETHINLKVSDGSSEIFFKIKKTTPLRR
LMEAFAKRQGKEMDSLRFLYDGIRIQADQTPEDLDMEDNDIIEAHREQIGGDYKDDDDKDLGKKLLEAA
RAGQDDEVRILMANGADVNAKDEYGLTPLYLATAHGHLEIVEVLLKNGADVNAVDAIGFTPLHLAAFIGH
LEIAEVLLKHGADVNAQDKFGKTAFDISIGNGNEDLAEILQKLNGSTSGSVTTLSGLSGEQGPSGDMTT
EEDSATHIKFSKRDEDGRELAGATMELRDSSGKTISTWISDGHVKDFYLYPGKYCLVGSSEIITRNYGK

TTIKEVVEIFDNDKNIQVLAFNTHTDNIEWAPIKAAQLTRPNAELVELEIDTLHGVKTIRVTPDHPVYTKNR
GYVRADELTDDDELVVAIENLYGDEVNGYFQGMEAETYIGKLKSRAIVSNEDTYDIQTSTH

$His_6$-SUMO-αHER2-SpyN-eNrdJ-1N$^{cage(1-26)}$

MGSSHHHHHHGSGLVPRGSASMSDSEVNQEAKPEVKPEVKPETHINLKVSDGSSEIFFKIKKTTPLRR
LMEAFAKRQGKEMDSLRFLYDGIRIQADQTPEDLDMEDNDIIEAHREQIGGDYKDDDDKDLGKKLLEAA
RAGQDDEVRILMANGADVNAKDEYGLTPLYLATAHGHLEIVEVLLKNGADVNAVDAIGFTPLHLAAFIGH
LEIAEVLLKHGADVNAQDKFGKTAFDISIGNGNEDLAEILQKLNGSTSGSVTTLSGLSGEQGPSGDMTT
EEDSATHIKFSKRDEDGRELAGATMELRDSSGKTISTWISDGHVKDFYLYPGKYCLVGSSEIITRNYGK
TTIKEVVEIFDNDKNIQVLAFNTHTDNIEWAPIKAAQLTRPNAELVELEIDTLHGVKTIRVTPDHPVYTKNR
GYVRADELTDDDELVVAIENLYGDEVNGYFQGMEAETYIGKLKSRAIVSNEDTYDIQT

$His_6$-SUMO-αHER2-SpyN-eNrdJ-1N$^{cage(1-20)}$

MGSSHHHHHHGSGLVPRGSASMSDSEVNQEAKPEVKPEVKPETHINLKVSDGSSEIFFKIKKTTPLRR
LMEAFAKRQGKEMDSLRFLYDGIRIQADQTPEDLDMEDNDIIEAHREQIGGDYKDDDDKDLGKKLLEAA
RAGQDDEVRILMANGADVNAKDEYGLTPLYLATAHGHLEIVEVLLKNGADVNAVDAIGFTPLHLAAFIGH
LEIAEVLLKHGADVNAQDKFGKTAFDISIGNGNEDLAEILQKLNGSTSGSVTTLSGLSGEQGPSGDMTT
EEDSATHIKFSKRDEDGRELAGATMELRDSSGKTISTWISDGHVKDFYLYPGKYCLVGSSEIITRNYGK
TTIKEVVEIFDNDKNIQVLAFNTHTDNIEWAPIKAAQLTRPNAELVELEIDTLHGVKTIRVTPDHPVYTKNR
GYVRADELTDDDELVVAIENLYGDEVNGYFQGMEAETYIGKLKSRAIVSNED

$His_6$-SUMO-αHER2-SpyN-NrdJ-1N

MGSSHHHHHHGSGLVPRGSASMSDSEVNQEAKPEVKPEVKPETHINLKVSDGSSEIFFKIKKTTPLRR
LMEAFAKRQGKEMDSLRFLYDGIRIQADQTPEDLDMEDNDIIEAHREQIGGDYKDDDDKDLGKKLLEAA
RAGQDDEVRILMANGADVNAKDEYGLTPLYLATAHGHLEIVEVLLKNGADVNAVDAIGFTPLHLAAFIGH
LEIAEVLLKHGADVNAQDKFGKTAFDISIGNGNEDLAEILQKLNGSTSGSVTTLSGLSGEQGPSGDMTT
EEDSATHIKFSKRDEDGRELAGATMELRDSSGKTISTWISDGHVKDFYLYPGKYCLVGSSEIITRNYGK
TTIKEVVEIFDNDKNIQVLAFNTHTDNIEWAPIKAAQLTRPNAELVELEIDTLHGVKTIRVTPDHPVYTKNR
GYVRADELTDDDELVVAI

$His_6$-SUMO-NrdJ-1C-SpyC-αEGFR

MGSSHHHHHHGSGLVPRGSASMSDSEVNQEAKPEVKPEVKPETHINLKVSDGSSEIFFKIKKTTPLRR
LMEAFAKRQGKEMDSLRFLYDGIRIQADQTPEDLDMEDNDIIEAHREQIGGMEAKTYIGKLKSRKIVSNE
DTYDIQTSTHNFFANDILVHNSFVETAAPDGYEVATPIEFTVNEDGQVTVDGEATEGDAHTGSTSGSDL
GKKLLEAARAGQDDEVRILMANGADVNADDTWGWTPLHLAAYQGHLEIVEVLLKNGADVNAYDYIGW

TPLHLAADGHLEIVEVLLKNGADVNASDYIGDTPLHLAAHNGHLEIVEVLLKHGADVNAQDKFGKTAFDI
SIDNGNEDLAEILQKLNEQKLISEEDL

His$_6$-SUMO-αEGFR-Decoy

MGSSHHHHHHGSGLVPRGSASMSDSEVNQEAKPEVKPEVKPETHINLKVSDGSSEIFFKIKKTTPLRR
LMEAFAKRQGKEMDSLRFLYDGIRIQADQTPEDLDMEDNDIIEAHREQIGGDYKDDDDKDLGKKLLEAA
RAGQDDEVRILMANGADVNADDTWGWTPLHLAAYQGHLEIVEVLLKNGADVNAYDYIGWTPLHLAAD
GHLEIVEVLLKNGADVNASDYIGDTPLHLAAHNGHLEIVEVLLKHGADVNAQDKFGKTAFDISIDNGNED
LAEILQKLNGSTSGSVTTLSGLSGEQGPSGDMTTEEDSATHIKFSERDEDGRELAGATMELRDSSGKTI
STWISDGHVKDFYLYPGKYCLVGSSEIITRNYGKTTIKEVVEIFDNDKNIQVLAFNTHTDNIEWAPIKAAQ
LTRPNAELVELEIDTLHGVKTIRVTPDHPVYTKNRGYVRADELTDDDELVVAIENLYGDEVNGYFQGME
AETYIGALASRAIVSNEDTYDIQTSTH

**Supplementary Table 11. Antibodies used for in this study.** Tabulated are epitopes and antibodies used in the study. All dilutions were done in TBS-T (25 mM Tris-HCl (pH 7.7), 150 mM NaCl, 0.1% v/v Tween-20) supplemented with 3% w/v BSA and 0.02% w/v sodium azide.

| Epitope | Antibody | Vendor (product) | Western blotting |
|---|---|---|---|
| FLAG | Rabbit anti-FLAG | Sigma (F7425) | 1:2,000 |
| Myc | Mouse anti-Myc | CST (2276S) | 1:2,000 |
| HA | Mouse anti-HA | Invitrogen (26183) | 1:2,000 |
| HER2 | Rabbit anti-HER2 | CST (4290) | 1:1,000 |
| EGFR | Rabbit anti-EGFR | CST (4267) | 1:1,000 |
| EpCAM | Mouse anti-EpCAM | Abcam (ab216136) | 1:2,000 |
| Dinitrophenol | Rabbit anti-DNP | CST (14681) | 1:10,000 |
| Rabbit IgG | Goat Anti-Rabbit IgG H&L (Alexa Fluor® 594) | Abcam (ab150080) | 1:10,000 |
| Rabbit IgG | IRDye 680RD Goat anti-Rabbit IgG (H + L) | LI-COR (926-68071) | 1:10,000 |
| Rabbit IgG | IRDye 800CW Goat anti-Rabbit IgG (H + L) | LI-COR (926-32211) | 1:10,000 |
| Mouse IgG | IRDye 680RD Goat anti-Mouse IgG (H + L) | LI-COR (926-68070) | 1:10,000 |
| Mouse IgG | IRDye 800CW Goat anti-Mouse IgG (H + L) | LI-COR (926-32210) | 1:10,000 |
| Biotin | IRDye 800CW Streptavidin | LI-COR (926-32230) | 1:10,000 |

**Supplementary Table 12. Mammalian cell culture media conditions.** All media were prepared with 100 U/mL penicillin/streptomycin and stored at 4 °C.

| Cell line | Medium | FBS | Supplements |
|---|---|---|---|
| K562 (WT) | RPMI-1640 | 5% | 25 mM HEPES |
| $K562^{HER2+}$ | RPMI-1640 | 5% | 25 mM HEPES |
| $K562^{EGFR+}$ | RPMI-1640 | 5% | 25 mM HEPES |
| $K562^{HER2+/EGFR+}$ | RPMI-1640 | 5% | 25 mM HEPES |
| $K562^{HER2+/EpCAMhi}$ | RPMI-1640 | 5% | 25 mM HEPES |
| $K562^{HER2+/EGFR+/EpCAMhi}$ | RPMI-1640 | 5% | 25 mM HEPES |
| OE19 | RPMI-1640 | 10% | 2 mM L-glutamine |
| A431 | DMEM | 10% | - |
| MCF-7 | DMEM | 10% | - |
| HCT-116 | McCoy's 5a Medium Modified | 10% | - |
| SK-BR-3 | McCoy's 5a Medium Modified | 10% | - |
| A549 | F-12K Medium | 10% | - |
| LoVo | F-12K Medium | 10% | - |
| MCF-10a | Mammary Epithelial Cell Growth Medium | 5% | Bovine Pituitary Extract (0.004 ml/ml)<br>Epidermal Growth Factor (recombinant human, 10 ng/ml)<br>Insulin (recombinant human, 5 µg/ml)<br>Hydrocortisone (0.5 µg/ml) |

### Supplementary Synthetic Methods

### Synthesis of DNP-maleimide

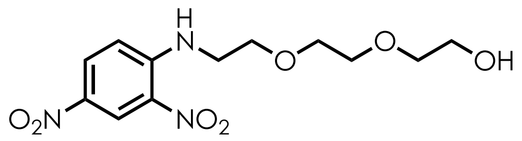


**2-(2-(2-((2,4-Dinitrophenyl)amino)ethoxy)ethoxy)ethan-1-ol (1).** To a clean, dry, round-bottomed flask equipped with a stir bar was added 2,4-dinitrochlorobenzene (1.02 g, 1 equiv., 5 mmol), 2-[2-(2-aminoethoxy)ethoxy]ethanol (746 mg, 1 equiv., 5 mmol), and ethanol (20 mL). Triethylamine (2 mL, 2.8 equiv, 14.3 mmol) was then added to the stirring flask at room temperature for 5 min. Next, the flask was equipped with a condenser and refluxed overnight. Upon completion of the reaction, the flask was cooled to room temperature, concentrated under reduced pressure, and reconstituted in EtOAc. The crude organic mixture was then sequentially washed with water and brine in a separatory funnel, dried over $Na_2SO_4$, before being concentrated under reduced pressure. The mixture was then purified using flash column chromatography on silica gel (gradient of 10–50% EtOAc:Hex, then gradient of 0–10% MeOH:EtOAc) to provide the title compound as a yellow gum in 81% yield (1.28 g).

**$^1$H NMR (500 MHz, $CDCl_3$)** δ 9.14 (d, *J* = 2.7 Hz, 1H), 8.83 (t, *J* = 5.1 Hz, 1H), 8.27 (dd, *J* = 9.5, 2.7 Hz, 1H), 6.94 (d, *J* = 9.5 Hz, 1H), 3.84 (t, *J* = 5.2 Hz, 2H), 3.78 – 3.69 (m, 6H), 3.61 (dt, *J* = 10.4, 4.8 Hz, 4H), 1.96 (bs, 1H).

**$^{13}$C NMR (126 MHz, $CDCl_3$)** δ 148.5, 136.3, 130.7, 130.5, 124.5, 114.1, 72.7, 71.0, 70.5, 68.6, 62.0, 43.3.

**HRMS (ESI):** Calculated for $C_{12}H_{17}N_3O_7$ $[M+Na]^+$: 338.09642; found: 338.09868.

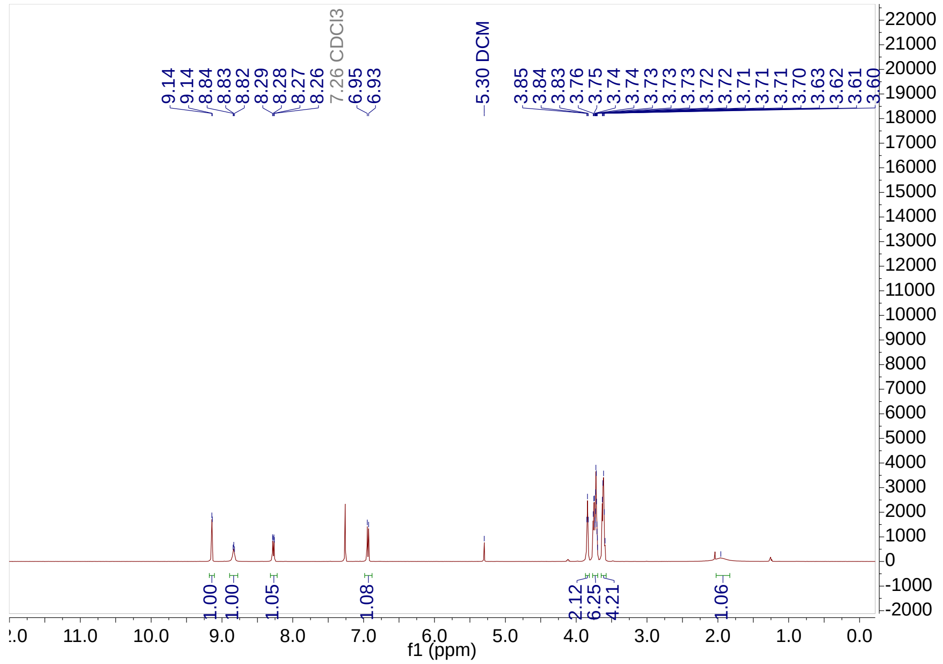

9.14
9.14
8.84
8.83
8.82
8.29
8.28
8.27
8.26
7.26 CDCl3
6.95
6.93
5.30 DCM
3.85
3.84
3.83
3.76
3.75
3.74
3.74
3.73
3.73
3.73
3.72
3.71
3.71
3.71
3.70
3.63
3.62
3.61
3.60
1.00
1.00
1.05
1.08
2.12
6.25
4.21
1.06
f1 (ppm)


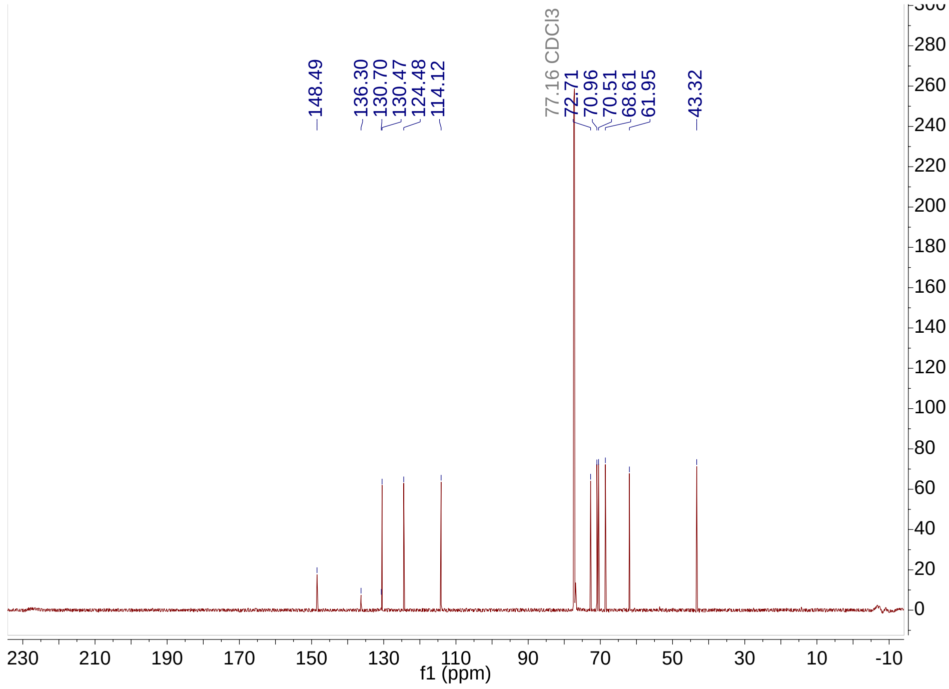

148.49
136.30
130.70
130.47
124.48
114.12
77.16 CDCl3
72.71
70.96
70.51
68.61
61.95
43.32
f1 (ppm)

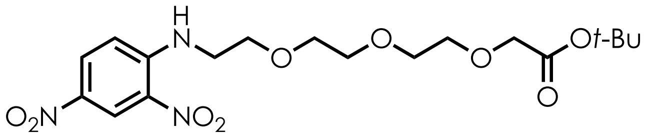

**Tert-butyl 2-(2-(2-(2-((2,4-dinitrophenyl)amino)ethoxy)ethoxy)ethoxy)acetate (2).** To a clean, dry, round-bottomed flask equipped with a stir bar was added anhydrous DMF (19 mL) and the flask was cooled to 0 °C. Then, sodium hydride (60% in mineral oil, 324.8 mg, 2 equiv., 8.12 mmol) was added portion-wise. Next, a solution of **1** (1.28 g, 1 equiv., 4.06 mmol) in DMF (5 mL) was added dropwise to the stirring NaH/DMF slurry and deprotonation was confirmed by observation of $H_2$ effervescence. The reaction mixture was stirred for 30 min under a positive $N_2$ pressure before dropwise addition of *tert*-butyl bromoacetate (0.72 mL, 1.2 equiv., 4.9 mmol) via syringe. The reaction was then warmed to room temperature and stirred overnight. Upon completion of the reaction, the mixture was quenched with saturated ammonium chloride solution and diluted with diethyl ether in a separatory funnel. The aqueous layer was washed twice with diethyl ether and the organic fractions collected. The organic layer was then washed with brine, dried over $MgSO_4$, and subsequently concentrated under reduced pressure. The crude produce was then purified using flash column chromatography on silica gel (gradient of 10–80% EtOAc:Hex) to provide the title compound as a yellow oil in 47% yield (0.82 g).

**$^1$H NMR (500 MHz, $CDCl_3$)** δ 9.14 (d, *J* = 2.5 Hz, 1H), 8.80 (t, *J* = 5.4 Hz, 1H), 8.26 (dd, *J* = 9.5, 2.7 Hz, 1H), 6.95 (d, *J* = 9.5 Hz, 1H), 4.00 (s, 2H), 3.84 (t, *J* = 5.3 Hz, 2H), 3.74 – 3.66 (m, 8H), 3.60 (q, *J* = 5.2 Hz, 2H), 1.46 (s, 9H).

**$^{13}$C NMR (126 MHz, $CDCl_3$)** δ 169.7, 148.6, 136.2, 130.7, 130.4, 124.4, 114.2, 81.7, 70.9, 70.9, 70.8, 70.8, 69.1, 68.7, 43.4, 28.2.

**HRMS (ESI):** Calculated for $C_{18}H_{27}N_3O_9$ $[M+Na]^+$: 452.16459; found: 452.16763.

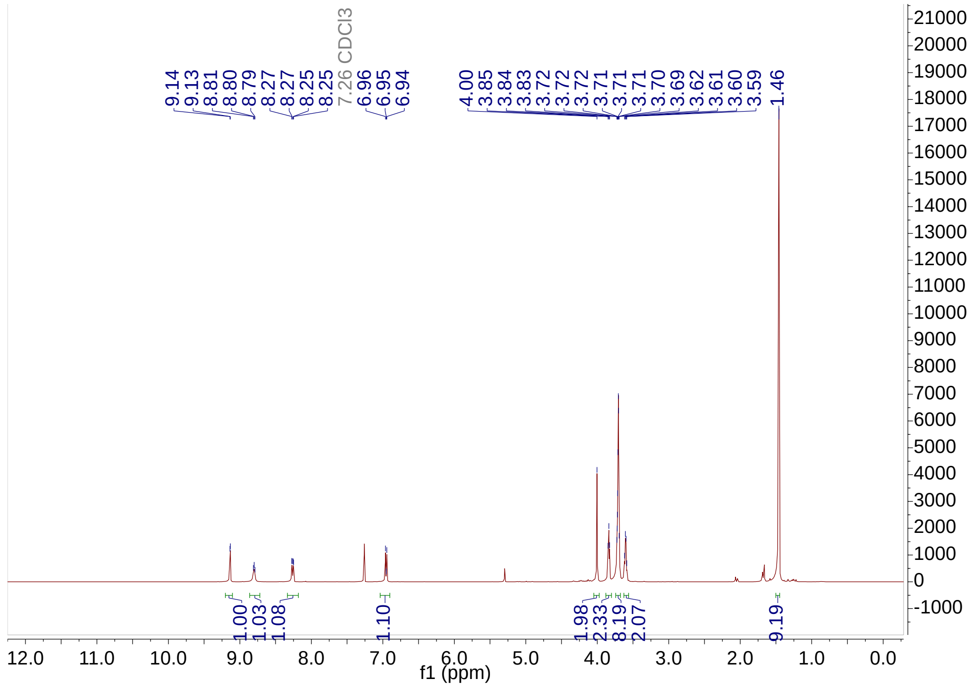
9.14
9.13
8.81
8.80
8.79
8.27
8.27
8.25
8.25
7.26 CDCl3
6.96
6.95
6.94
4.00
3.85
3.84
3.83
3.72
3.72
3.72
3.71
3.71
3.71
3.70
3.69
3.62
3.61
3.60
3.59
1.46
1.00
1.03
1.08
1.10
1.98
2.33
8.19
2.07
9.19
f1 (ppm)

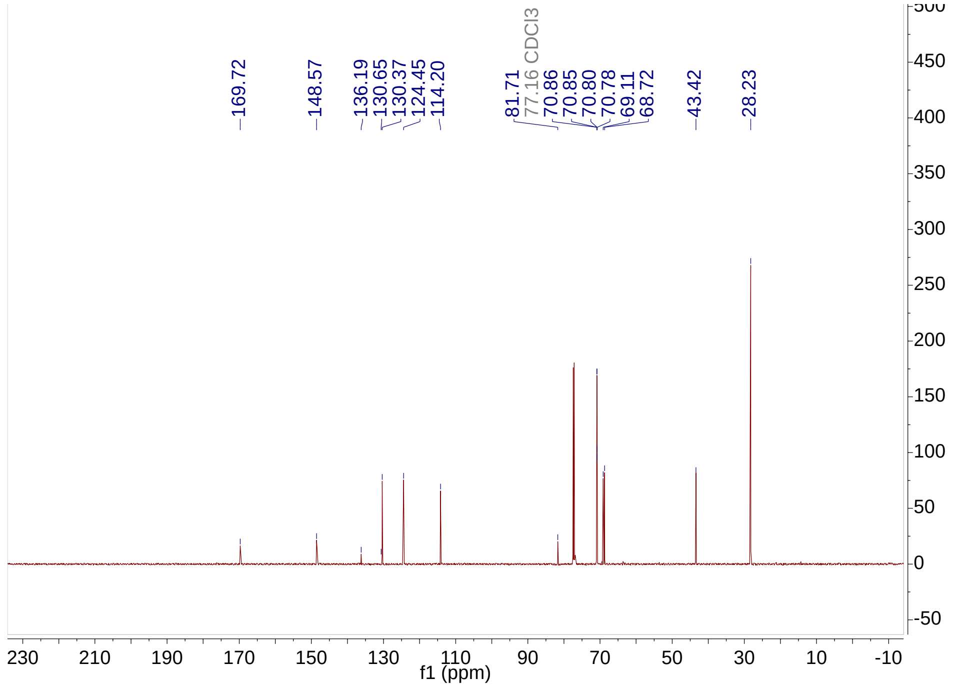
169.72
148.57
136.19
130.65
130.37
124.45
114.20
81.71
77.16 CDCl3
70.86
70.85
70.80
70.78
69.11
68.72
43.42
28.23
f1 (ppm)

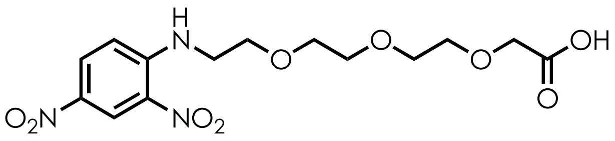

**2-(2-(2-(2-((2,4-Dinitrophenyl)amino)ethoxy)ethoxy)ethoxy)acetic acid (3).** To a clean, dry, round-bottomed flask equipped with a stir bar was added **2** (714.9 mg, 1 equiv., 1.92 mmol) and DCM (10 mL). The flask was then cooled to 0 °C and TFA was added to the stirring mixture. The reaction mixture was stirred at 0 °C for 10 min before being warmed to room temperature and stirred overnight. Conversion of the starting material was detected by TLC and upon complete conversion of starting material, the mixture was concentrated under reduced pressure, using 2 x 10 mL of MeOH to form a TFA-azeotrope leading to removal of residual TFA. The resulting product was carried to the next step without further purification (681 mg, 95%).

**$^1$H NMR (500 MHz, $CDCl_3$)** δ 9.13 (d, *J* = 2.7 Hz, 1H), 8.80 (t, *J* = 5.3 Hz, 1H), 8.26 (dd, *J* = 9.5, 2.7 Hz, 1H), 6.95 (d, *J* = 9.6 Hz, 1H), 4.15 (s, 2H), 3.83 (t, *J* = 5.3 Hz, 2H), 3.70 (tdd, *J* = 5.0, 3.5, 1.7 Hz, 8H), 3.60 (q, *J* = 5.2 Hz, 2H).

**$^{13}$C NMR (126 MHz, $CDCl_3$)** δ 171.0, 148.5, 136.1, 130.6, 130.4, 124.4, 114.2, 71.0, 70.8, 70.8, 70.7, 68.7, 68.7, 43.4.

**HRMS (ESI):** Calculated for $C_{14}H_{19}N_3O_9$ $[M+Na]^+$: 396.10190; found: 396.10217.

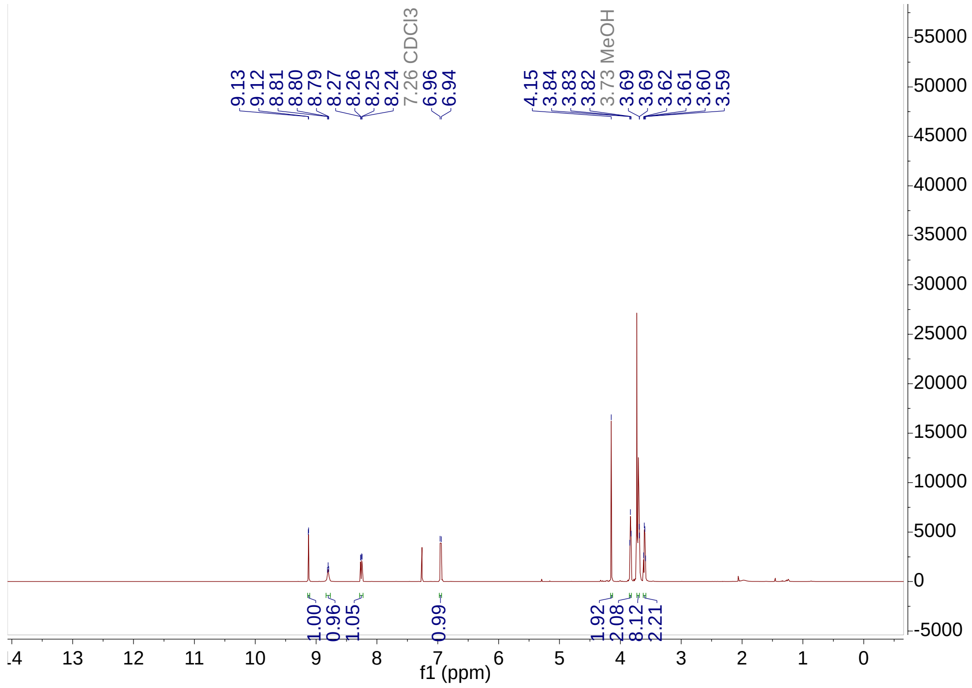
9.13
9.12
8.81
8.80
8.79
8.27
8.26
8.25
8.24
7.26 CDCl3
6.96
6.94
4.15
3.84
3.83
3.82
3.73 MeOH
3.69
3.69
3.62
3.61
3.60
3.59
1.00
0.96
1.05
0.99
1.92
2.08
8.12
2.21
f1 (ppm)

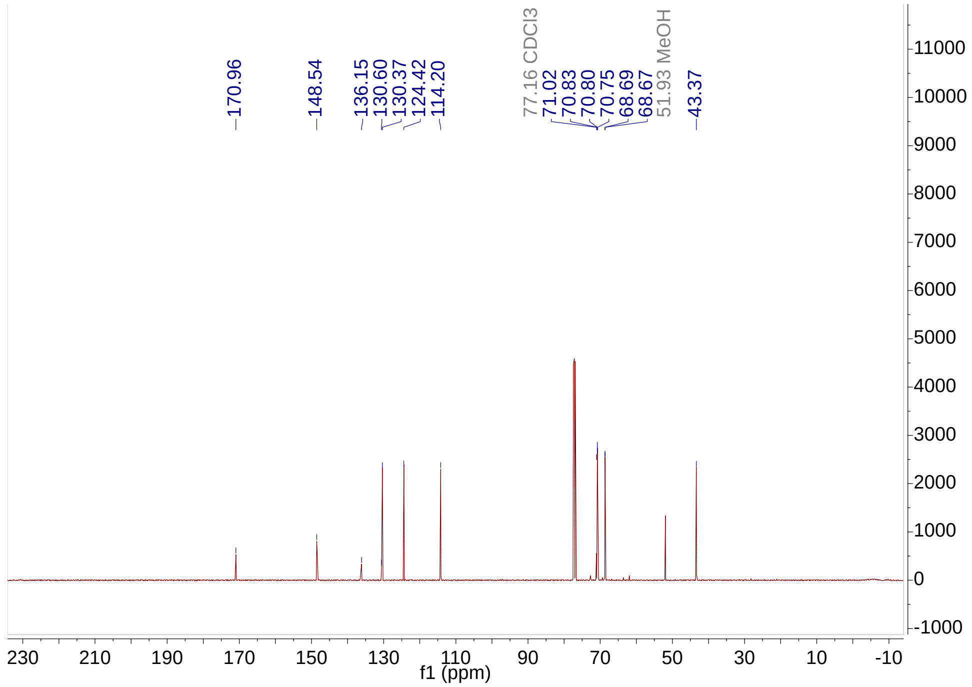
170.96
148.54
136.15
130.60
130.37
124.42
114.20
77.16 CDCl3
71.02
70.83
70.80
70.75
68.69
68.67
51.93 MeOH
43.37
f1 (ppm)

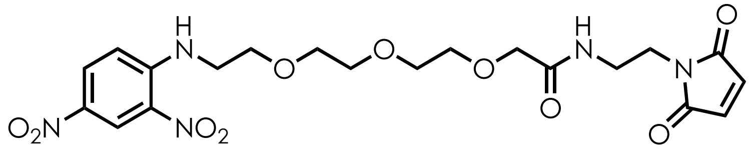

**2-(2-(2-(2-((2,4-Dinitrophenyl)amino)ethoxy)ethoxy)ethoxy)-N-(2-(2,5-dioxo-2,5-dihydro-1H-pyrrol-1-yl)ethyl)acetamide (4).** To a clean, dry, round-bottomed flask equipped with a stir bar was added **3** (190.3 mg, 1 equiv., 0.509 mmol), *N*-hydroxysuccinimide (58.7 mg, 1 equiv., 0.510 mmol) and DCM (10 mL). *N*,*N*'-Dicyclohexylcarbodiimide (115.8 mg, 1.1 equiv., 0.561 mmol) was then added to the reaction solution and the formation of dicyclohexylurea is observed within 5 min. The reaction was stirred for at least 6 h before being filtered over Celite® in a glass frit (medium porosity). The collected filtrate containing the NHS ester was concentrated under reduced pressure and used in the next step without further purification. The crude product was then reconstituted in 10 mL DCM within a clean, dry, round-bottomed flask equipped with a stir bar. To the NHS-ester solution was added 1-(2-aminoethyl)maleimide hydrochloride (90 mg, 1.0 equiv., 0.509 mmol), followed by triethylamine (0.36 mL, 5 equiv., 2.55 mmol). The reaction mixture was then monitored by TLC until full conversion of the NHS-ester was observed. The reaction was then quenched with DI water and diluted with DCM in a separatory funnel. The aqueous layer was extracted 2x with DCM, followed by combination of the organic layers, which were then dried over $MgSO_4$. The crude mixture was then concentrated under reduced pressure and purified using flash column chromatography on silica gel (gradient of 10–100% EtOAc:Hex) to provide the title compound as a yellow oil in 65% yield (163.8 mg).

**$^1$H NMR (500 MHz, $CDCl_3$)** δ 9.12 (d, *J* = 2.7 Hz, 1H), 8.80 (t, *J* = 5.3 Hz, 1H), 8.27 (dd, *J* = 9.5, 2.7 Hz, 1H), 7.23 – 7.15 (m, 1H), 6.95 (d, *J* = 9.6 Hz, 1H), 6.70 (s, 2H), 3.94 (s, 2H), 3.83 (t, *J* = 5.2 Hz, 2H), 3.73 – 3.65 (m, 10H), 3.61 (q, *J* = 5.2 Hz, 2H), 3.47 (q, *J* = 5.9 Hz, 2H).

**$^{13}$C NMR (126 MHz, $CDCl_3$)** δ 170.9, 170.8, 148.5, 136.2, 134.3, 130.6, 130.4, 124.4, 114.2, 71.0, 70.9, 70.7, 70.6, 70.5, 68.7, 43.3, 38.0, 37.6.

**HRMS (ESI):** Calculated for $C_{20}H_{25}N_5O_{10}$ $[M+H]^+$: 496.16797; found: 496.16863.

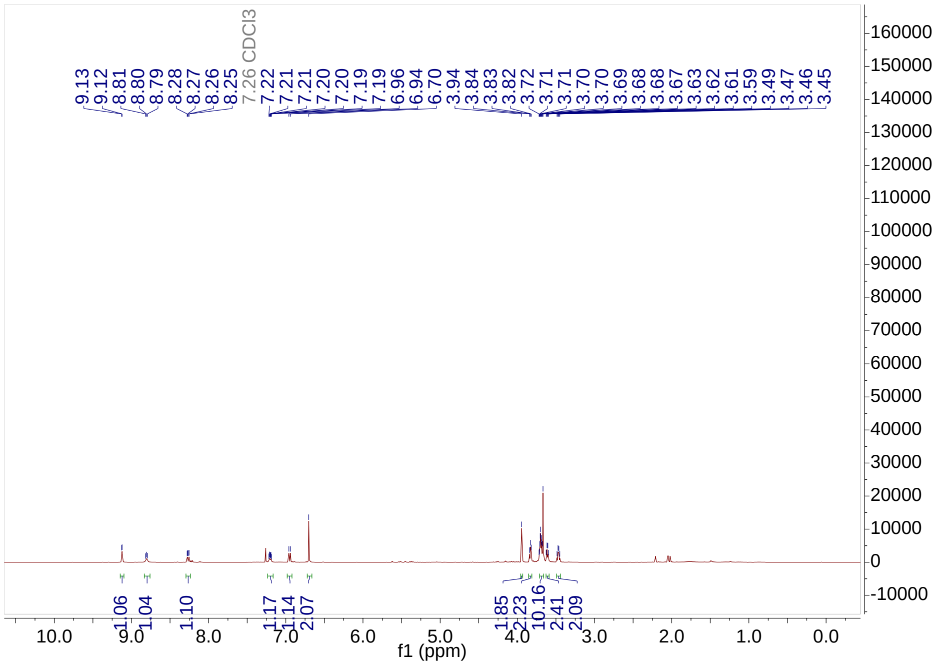
CDCl3
f1 (ppm)

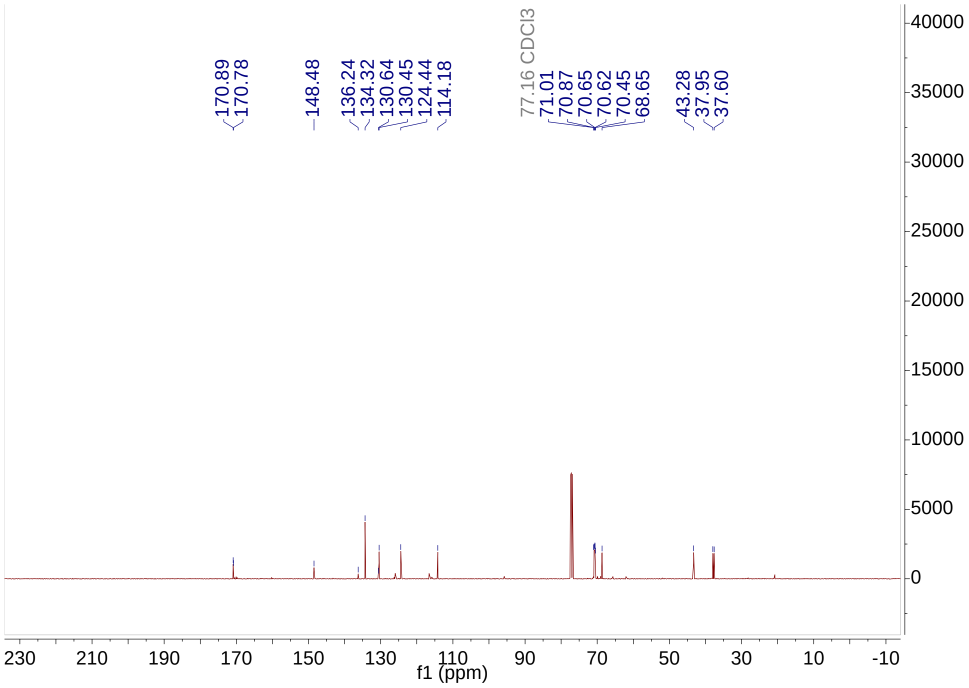
CDCl3
f1 (ppm)

**Synthesis of biotin-phenol**

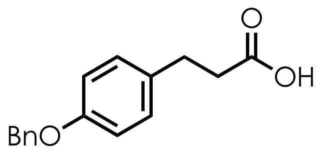


**3-(4-(benzyloxy)phenyl)propanoic acid (5).** The title compound was prepared in accordance with literature reference[46].

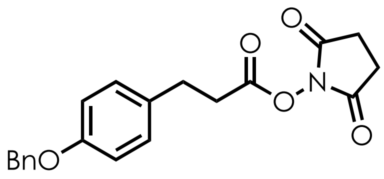

**2,5-dioxopyrrolidin-1-yl 3-(4-(benzyloxy)phenyl)propanoate (6).** To a clean, dry, round-bottomed flask equipped with a stir bar was added **5** (2.40 g, 1 equiv., 9.4 mmol), *N*-hydroxysuccinimide (1.08 g, 1 equiv., 9.4 mmol) and DCM (20 mL). *N*,*N*'-Dicyclohexylcarbodiimide (2.02 g, 1.05 equiv., 9.8 mmol) was then added to the reaction solution and the formation of dicyclohexylurea is observed within 5 min. The reaction is stirred overnight before being filtered over Celite® in a glass frit (medium porosity). The collected filtrate containing the NHS ester is concentrated under reduced pressure and used in the next step without further purification.

**$^{1}$H NMR (500 MHz, $CDCl_3$)** $^{1}$H NMR (500 MHz, $CDCl_3$) δ 7.46 – 7.36 (m, 4H), 7.35 – 7.29 (m, 1H), 7.15 (d, *J* = 8.6 Hz, 2H), 6.93 (d, *J* = 8.7 Hz, 2H), 5.04 (s, 2H), 3.00 (t, *J* = 7.8 Hz, 2H), 2.93 – 2.86 (m, 2H), 2.85 – 2.81 (m, 4H)
**$^{13}$C NMR (126 MHz, $CDCl_3$)** δ 169.3, 168.1, 157.7, 137.2, 131.6, 129.4, 128.7, 128.1, 127.6, 115.2, 70.2, 33.1, 29.8, 25.7.

**HRMS (ESI):** Calculated for $C_{20}H_{19}NO_5$ $[M+Na]^+$: 376.11609; found: 376.11649.

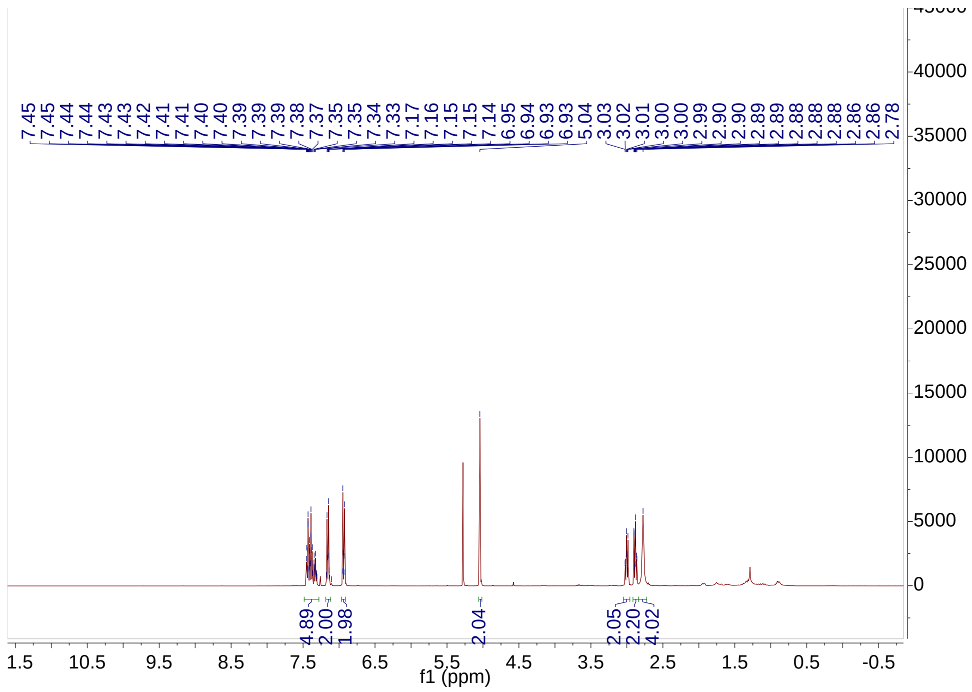
7.45
7.45
7.44
7.44
7.43
7.43
7.42
7.41
7.41
7.40
7.40
7.39
7.39
7.39
7.38
7.37
7.35
7.35
7.34
7.33
7.17
7.16
7.15
7.15
7.14
6.95
6.94
6.93
6.93
5.04
3.03
3.02
3.01
3.00
3.00
2.99
2.90
2.90
2.89
2.89
2.88
2.88
2.88
2.88
2.86
2.78
4.89
2.00
1.98
2.04
2.05
2.20
4.02
f1 (ppm)

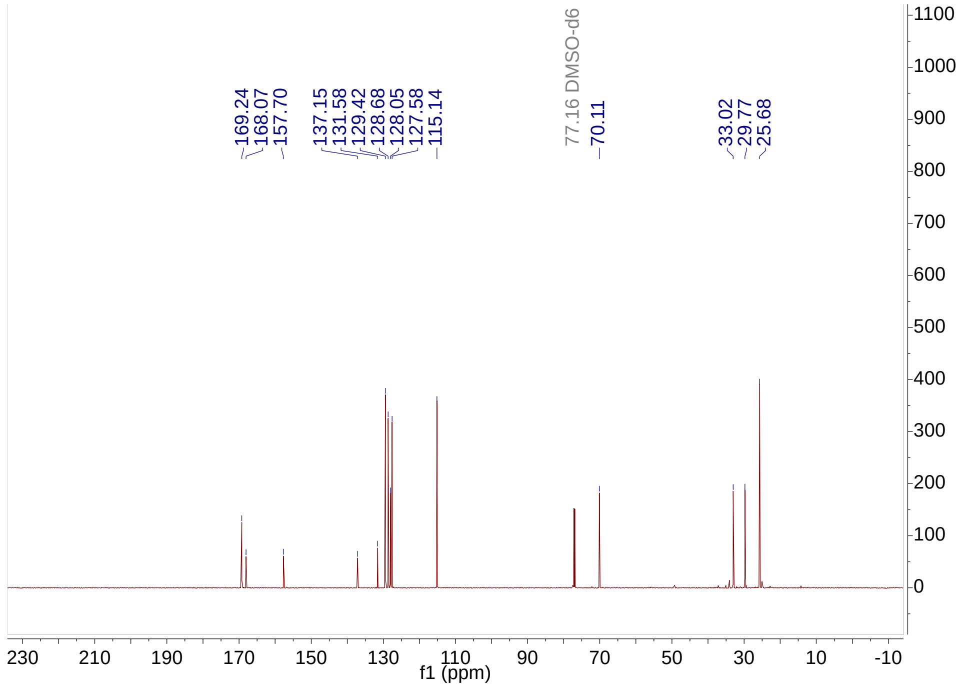
169.24
168.07
157.70
137.15
131.58
129.42
128.68
128.05
127.58
115.14
77.16 DMSO-d6
70.11
33.02
29.77
25.68
f1 (ppm)

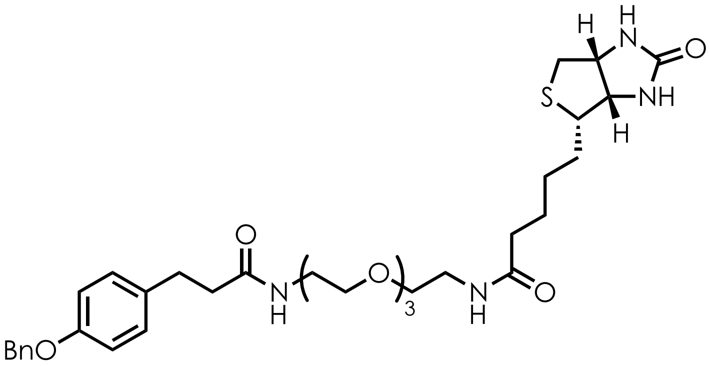

***N*-(15-(4-(benzyloxy)phenyl)-13-oxo-3,6,9-trioxa-12-azapentadecyl)-5-((3aS,4S,6aR)-2-oxohexahydro-1H-thieno[3,4-d]imidazol-4-yl)pentanamide (7).** To a dry, clean 20 mL vial was added NHS ester **6** (84.8 mg, 1.0 equiv., 0.24 mmol), biotin-PEG3-amine (100 mg, 1.0 equiv., 0.24 mmol) and DCM (2 mL). Triethylamine (0.10 mL, 3.0 equiv., 0.72 mmol) was then added to the reaction mixture and the reaction stirred overnight at room temperature. Upon completion of the reaction, the mixture is concentrated under reduced pressure and purified using flash column chromatography on silica gel (gradient of 100% DCM to 20% MeOH:DCM) to provide the title compound as a colorless solid in 67% yield (106.2 mg).

**$^{1}$H NMR (500 MHz, $CDCl_3$)** δ 7.52 – 7.26 (m, 5H), 7.12 (d, *J* = 8.3 Hz, 2H), 6.93 – 6.81 (m, 2H), 6.68 (d, *J* = 6.0 Hz, 1H), 6.52 (s, 1H), 6.43 (d, *J* = 5.7 Hz, 1H), 5.51 (s, 1H), 5.02 (s, 2H), 4.45 (dd, *J* = 7.9, 4.9 Hz, 1H), 4.25 (dd, *J* = 8.0, 4.7 Hz, 1H), 3.63 – 3.55 (m, 6H), 3.53 (t, *J* = 5.1 Hz, 2H), 3.50 (t, *J* = 5.1 Hz, 2H), 3.41 (tt, *J* = 7.7, 4.0 Hz, 4H), 3.10 (td, *J* = 7.4, 4.6 Hz, 1H), 2.93 – 2.80 (m, 3H), 2.71 (d, *J* = 12.8 Hz, 1H), 2.45 (dd, *J* = 8.7, 6.9 Hz, 2H), 2.19 (t, *J* = 7.4 Hz, 2H), 1.77 – 1.52 (m, 3H), 1.41 (p, *J* = 7.5 Hz, 2H).

**$^{13}$C NMR (126 MHz, $CDCl_3$)** δ 173.4, 172.6, 164.2, 157.3, 137.2, 133.5, 129.5, 129.4, 128.7, 128.7, 128.1, 128.0, 127.6, 114.9, 70.4, 70.4, 70.2, 70.1, 70.1, 70.1, 61.9, 60.3, 55.7, 40.6, 39.3, 39.2, 38.6, 36.0, 31.0, 28.3, 28.2, 25.7.

**HRMS (ESI):** Calculated for $C_{34}H_{48}N_4O_7S$ $[M+Na]^+$: 679.31414; found: 679.31600.

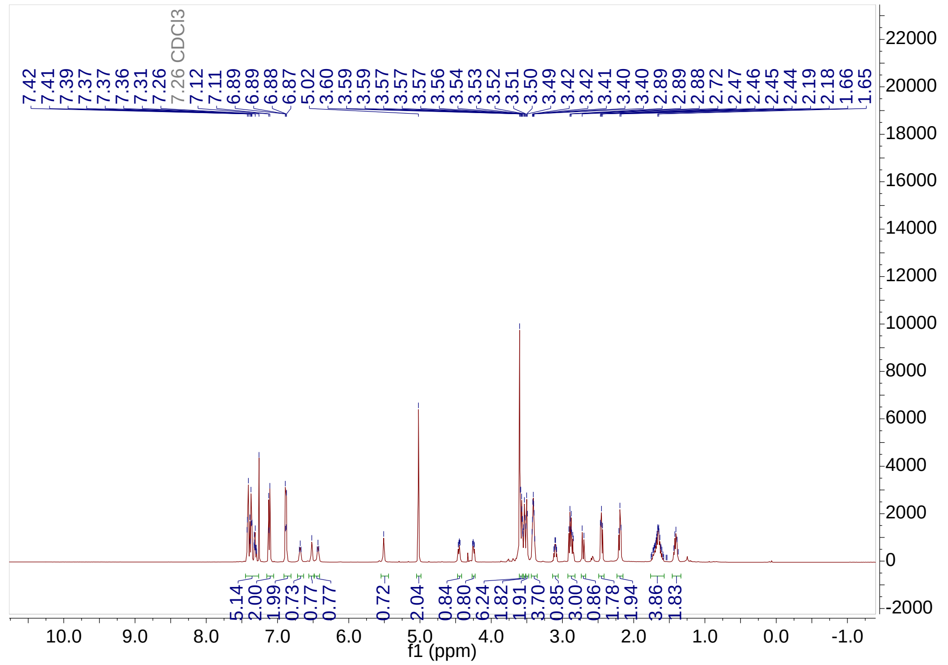
7.42 7.41 7.39 7.37 7.37 7.36 7.31 7.26 CDCl3 7.12 7.11 6.89 6.89 6.88 6.87 5.02 3.60 3.59 3.59 3.57 3.57 3.57 3.56 3.54 3.53 3.52 3.51 3.50 3.49 3.42 3.42 3.41 3.40 3.40 2.89 2.89 2.88 2.72 2.47 2.46 2.45 2.44 2.19 2.18 1.66 1.65
5.14 2.00 1.99 0.73 0.77 0.77 0.72 2.04 0.84 0.80 6.24 1.82 1.91 3.70 0.85 3.00 0.86 1.78 1.94 3.86 1.83
f1 (ppm)

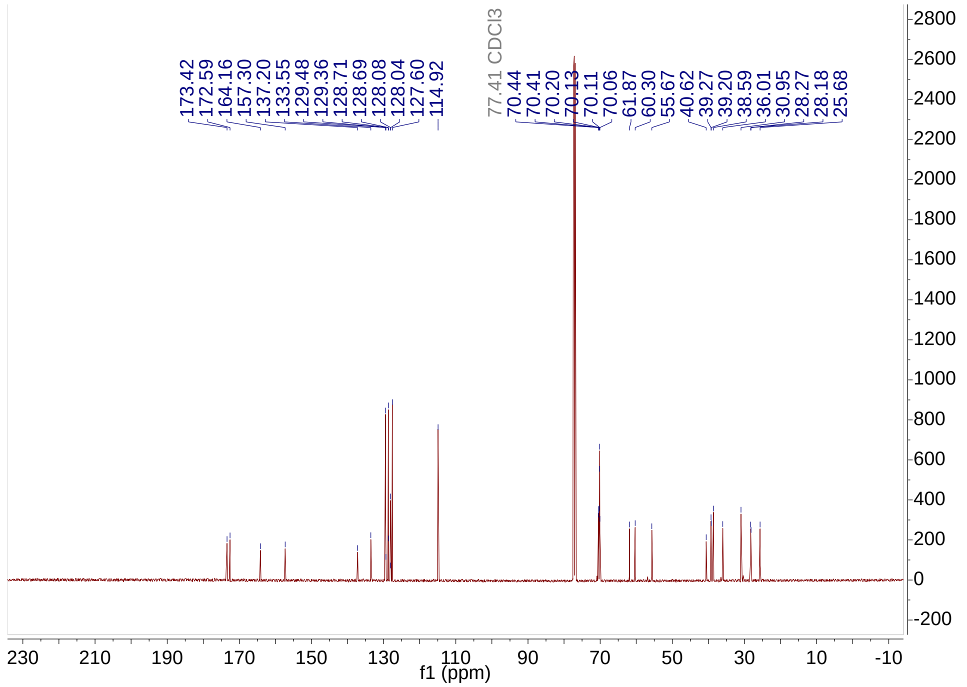
173.42 172.59 164.16 157.30 137.20 133.55 129.48 129.36 128.71 128.69 128.08 128.04 127.60 114.92
77.41 CDCl3 70.44 70.41 70.20 70.13 70.11 70.06 61.87 60.30 55.67 40.62 39.27 39.20 38.59 36.01 30.95 28.27 28.18 25.68
f1 (ppm)

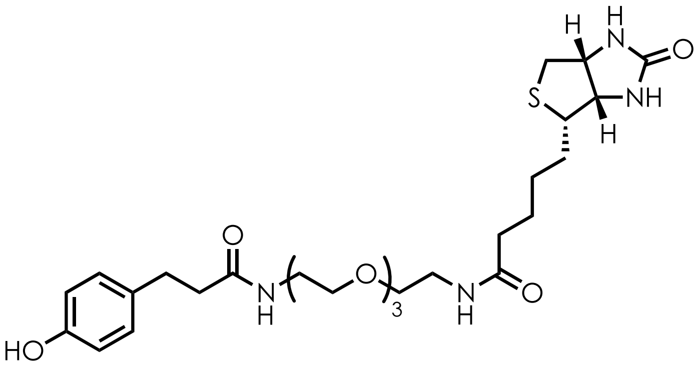

**Biotin-PEG3-desaminotyrosine (8).** To a clean, dry round-bottomed flask equipped with a stir bar was added substrate **7** (106.20 mg, 1.0 equiv., 0.16 mmol). The solid was dissolved in a 1:1 MeOH:THF mixture. 10% Pd/C (100 mg) was added to the flask, and the flask was sealed with a rubber septum and immediately placed under a positive $N_2$ pressure. The flask was purged and backfilled twice with $N_2$. The reaction flask was then purged and refilled with a balloon containing an atmosphere of $H_2$ (1 atm) and left to stir overnight at room temperature. At the end of the reaction, the reaction was filtered through Celite® over a filter frit (medium porosity) and concentrated under reduced pressure. The crude mixture is purified using flash column chromatography on silica gel (gradient of 100% DCM to 20% MeOH:DCM) to provide the title compound as a colorless solid in 37% yield (33.7 mg).

**$^{1}$H NMR (500 MHz, MeOD)** δ 7.00 – 6.93 (m, 2H), 6.67 – 6.59 (m, 2H), 4.44 (dd, *J* = 7.9, 4.8 Hz, 1H), 4.24 (dd, *J* = 7.9, 4.5 Hz, 1H), 3.58 (ddp, *J* = 8.0, 4.9, 2.5 Hz, 5H), 3.48 (t, *J* = 5.5 Hz, 2H), 3.41 (d, *J* = 5.5 Hz, 2H), 3.31 (t, *J* = 5.5 Hz, 2H), 3.29 – 3.24 (m, 4H), 3.14 (ddd, *J* = 9.0, 5.9, 4.4 Hz, 1H), 2.87 (dd, *J* = 12.8, 4.9 Hz, 1H), 2.76 (t, *J* = 7.6 Hz, 2H), 2.65 (d, *J* = 12.7 Hz, 1H), 2.39 (dd, *J* = 8.2, 7.0 Hz, 2H), 2.16 (t, *J* = 7.3 Hz, 2H), 1.75 – 1.48 (m, 3H), 1.38 (p, *J* = 7.8 Hz, 2H).

**$^{13}$C NMR (126 MHz, MeOD)** δ 176.1, 175.5, 166.1, 156.8, 132.9, 130.4, 116.2, 71.6, 71.2, 71.2, 70.6, 70.6, 63.3, 61.6, 57.0, 41.0, 40.3, 40.3, 39.3, 36.7, 32.1, 29.8, 29.5, 26.8.

**HRMS (ESI):** Calculated for $C_{27}H_{42}N_4O_7S$ $[M+Na]^+$: 589.26719; found: 589.26959.

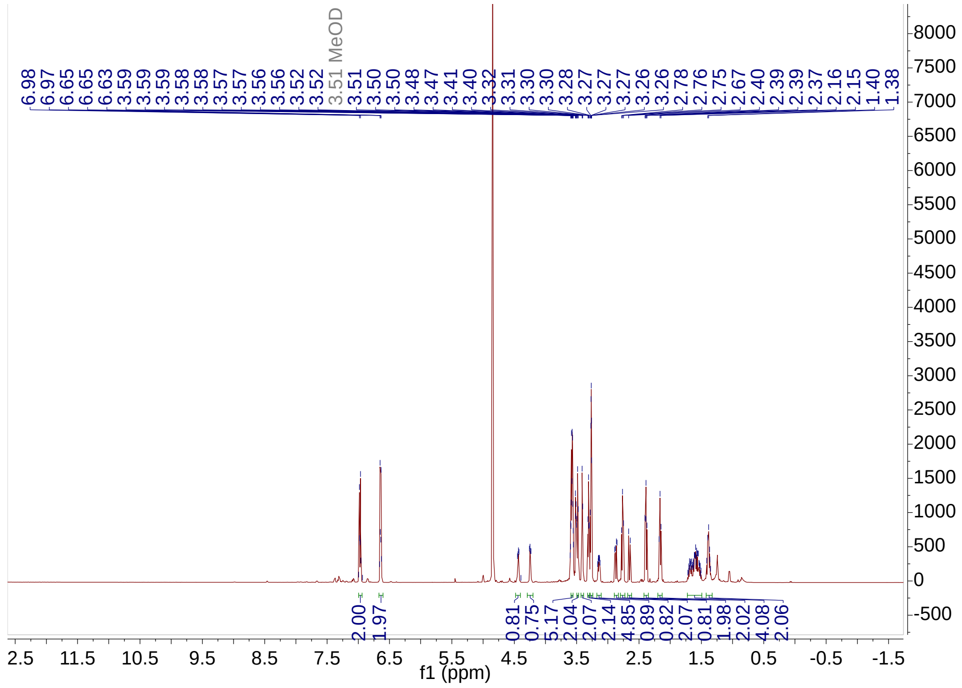
6.98 6.97 6.65 6.65 6.63 3.59 3.59 3.59 3.58 3.58 3.57 3.57 3.56 3.52 3.52 3.51 MeOD 3.51 3.50 3.50 3.48 3.47 3.41 3.40 3.32 3.31 3.30 3.28 3.27 3.27 3.27 3.26 3.26 2.78 2.76 2.75 2.67 2.40 2.39 2.37 2.16 2.15 1.40 1.38
2.00 1.97 0.81 0.75 5.17 2.04 2.07 2.14 4.85 0.89 0.82 2.07 0.81 1.98 2.02 4.08 2.06
f1 (ppm)

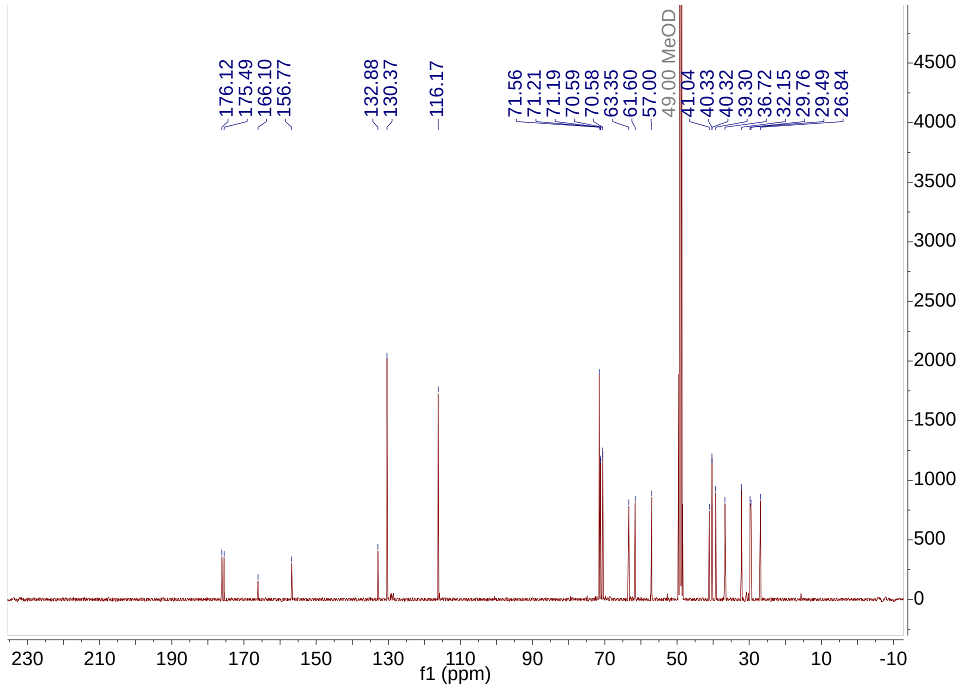
176.12 175.49 166.10 156.77 132.88 130.37 116.17 71.56 71.21 71.19 70.59 70.58 63.35 61.60 57.00 49.00 MeOD 41.04 40.33 40.32 39.30 36.72 32.15 29.76 29.49 26.84
f1 (ppm)
